# Transposable elements have contributed human regulatory regions that are activated upon bacterial infection

**DOI:** 10.1101/707612

**Authors:** Lucia Bogdan, Luis Barreiro, Guillaume Bourque

## Abstract

Transposable elements (TEs) are increasingly recognized as important contributors to mammalian regulatory systems. For instance, they have been shown to play a role in the human interferon response. However, their involvement in other mechanisms of immune cell activation remains poorly understood. We investigated the profile of accessible chromatin enhanced in stimulated human macrophages using ATAC-Seq to assess the role of different TE subfamilies in regulating the immune response. We found that both previously identified and new repeats belonging to the MER44, THE1, Tigger3 and MLT1 families provide 14 subfamilies that are enriched in differentially accessible chromatin and found near differentially expressed genes. These TEs also harbour binding motifs for several candidate transcription factors, including important immune regulators AP-1 and NF-kB, present in 96% of accessible MER44B and 83% of THE1C instances, respectively. To more directly assess their regulatory potential, we evaluated their presence in regions putatively affecting gene expression, as defined by quantitative trait locus (QTL) analysis, and find that repeats are also contributing to accessible elements near QTLs. Together, these results suggest that a number of TE families have contributed to the regulation of the immunogenomic response to infection in humans.

## INTRODUCTION

The regulatory networks responsible for immune function are particularly susceptible to evolutionary pressures as organisms adapt to withstand continuous assaults from a variety of infectious agents. Through their ability to move and replicate,^1^ be bound by transcription factors,^2,3^ and produce non-coding transcripts with regulatory potential,^4,5^ there is increasing evidence that transposable elements (TEs) have been co-opted throughout evolution and have contributed to the generation of regulatory diversity in mammalian genomes.^6^ As such, they present interesting targets for the discovery of novel functional elements contributing to gene regulation in different human cell types, particularly immune cells. A notable example is the MER41 repeat and its associated subfamilies, which includes an element in the AIM2 promoter with immune regulatory function confirmed through CRISPR-Cas9 knockouts.^7^ In that analysis, Chuong *et al.* also described the enrichment of several other TE subfamilies within STAT1 and IRF1 transcription factor binding sites putatively involved in the interferon pathway of macrophages.

To gain a more comprehensive understanding of the TE-derived non-coding elements implicated in the immune response, we explored the landscape of accessible chromatin in macrophages during immune cell activation. More specifically, we investigated the association between repetitive elements and non-coding regions activated upon infection in the datasets published by Nédélec *et al.*,^8^ which evaluated population differences in the response to infection. From that study, we obtained assays for transposase-accessible chromatin (ATAC-Seq) which were derived from macrophages challenged *in vitro* with *Listeria monocytogenes* and *Salmonella typhimurium*, two intracellular pathogens. The regulatory contribution of TEs has been previously described in immune cells,^7^ but analyses in that study were limited by targeting specific inflammatory pathways and transcription factor binding sites. In that respect, ATAC-Seq data offers the advantage of targeting all regions of accessible chromatin genome-wide without the need for a pull-down experiment that would restrict the analysis to a pre-selected set of transcription factors.^9^ Moreover, these data allowed us to explore the presence of TEs in the functional regions identified by expression quantitative trait locus (eQTL). Overall, we show that TEs contribute to non-coding sequences activated upon infection, as they are overrepresented both in accessible chromatin and in proximity to differentially expressed genes, harbour transcription factor motifs and are enriched near QTLs belonging to their haplotype block.

## RESULTS

### TEs contribute to accessible chromatin upon infection

We were interested in the contributions of TEs to the accessible chromatin that was specific to the immune response upon infection. For this purpose, we obtained ATAC-Seq datasets from Nédélec *et al.* and focused our analysis on regions of the genome where the chromatin becomes more accessible following bacterial exposure (see Methods). We processed the datasets separately for macrophages challenged with *Salmonella typhimurium* (S12) and *Listeria monocytogenes* (L12), 12 hours post infection. By overlapping the peak summits with repetitive element annotations from the *RepeatMasker* track,^10^ we found that 25.6% (7,995/31,256) and 23.6% (4,678/19,788) of infection-induced accessible regions contain TEs in the S12 and L12 samples, respectively. That being said, given that nearly half of the genome is derived from repeats^6^ and consistent with previous reports^11^, we find that overall TEs are underrepresented in regions for which the chromatin becomes more accessible upon infection.

Next, we sought to identify specific TE families recurrently overrepresented in the S12 and L12 infected samples. We therefore compared the presence of peak-associated repeats (PARs) with their expected distribution and computed the statistical enrichment of TEs at three levels of repeat organization: individual subfamilies, families and the four main TE classes (LTR, DNA, LINE and SINE) (Methods). This analysis revealed 34 “immune” subfamilies significantly enriched in both conditions (Tables S1-S2), including the MER41B repeat previously shown to have a regulatory role in the immune activation of the AIM2 gene.^7^ The strongest enrichment observed was for the MER44B subfamily, which was found 16 more times than expected in both conditions (*p* = 3.36e- 75 and *p* = 1.29e-38 for S12 and L12, respectively) (Figure 1a-b). Notably, the related subfamilies MER44C and MER44D were also highly enriched, suggesting the potential presence of conserved activating elements within these related TEs. The THE1, Tigger3 and MLT1 subfamilies were also overrepresented, providing respectively 2, 4 and 9 related subfamilies enriched in at least one condition. However, the MER44B repeats were unique in their contribution, as they represent a small subfamily with only 2131 instances, but still provide as many PARs as the MLT1K subfamily, which is 9 times larger in size (Figure 1c).

**Figure 1.**
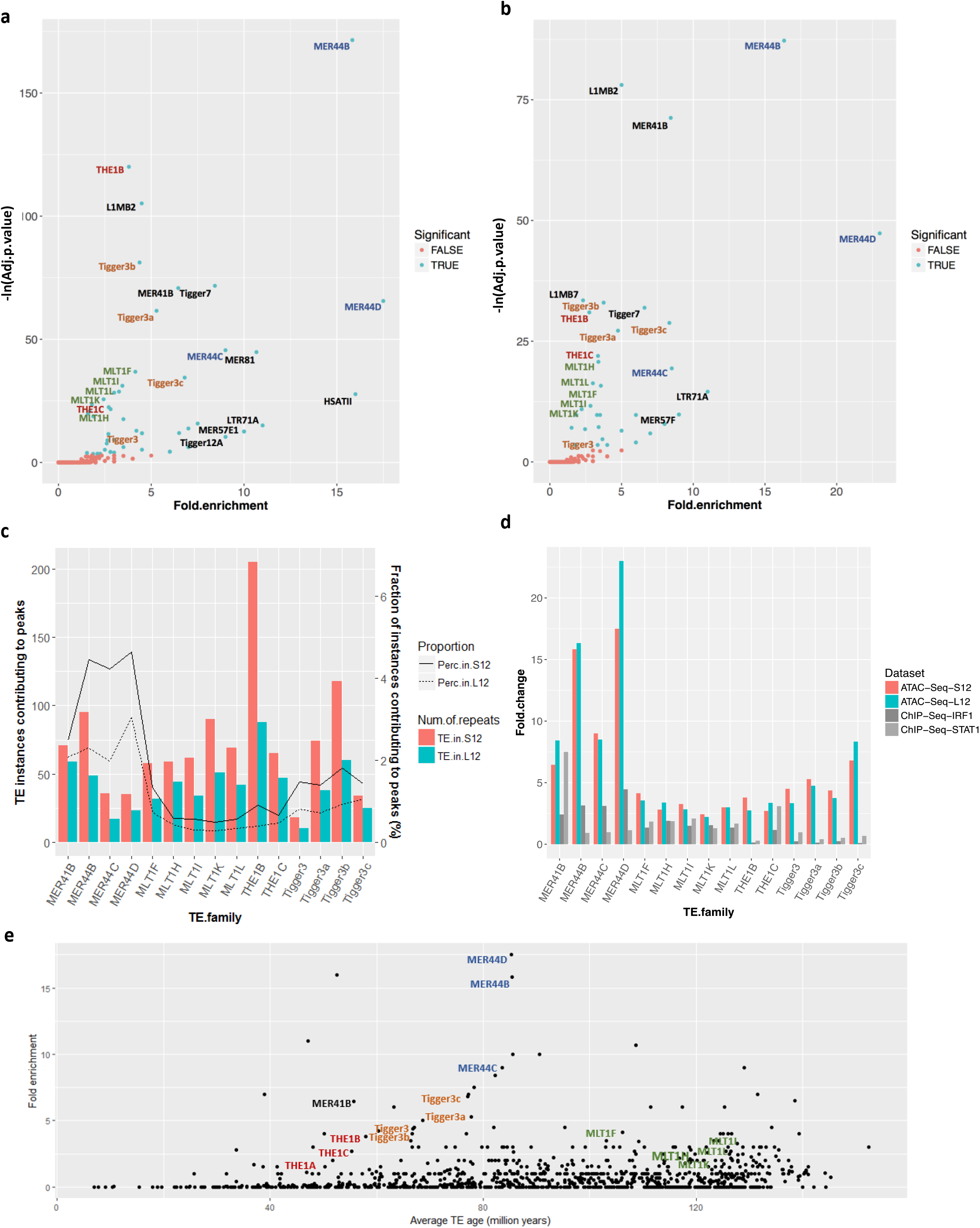
(a) TE enrichment in ATAC peaks by repeat subfamily (S12 samples). (b) TE enrichment in L12 samples. The label colors match the related family they belong to (blue for MER44, red for THE1, orange for Tigger3 and green for MLT1). (c) The absolute number of TE instances (bars) and the fraction of all instances in each subfamily (lines) contributing to accessible chromatin. (d) Comparison of TE enrichment results in our ATAC-Seq peaks and the ChIP-Seq datasets published by Chuong *et al.* (e) Age of TE instances in accessible and inaccessible chromatin for each repeat family.

From this analysis, we therefore identified 14 TE subfamilies belonging to 4 groups of related repeats belonging to the MER44, THE1, Tigger3 and MLT1 subfamilies, as well as the previously described MER41B subfamily (Figure 1c-e). A comparison with Chuong *et al.* shows an overlapping set of TE subfamilies enriched in the STAT1 and IRF1 ChIP-Seq datasets, which map the transcription factor binding sites required for the regulation of interferon-stimulated genes in CD14+ cells. However, there are several notable differences between our findings and the previously described data (Figure 1d). In particular, the MER44, Tigger3 and THE1B repeats are several times more enriched in the overall landscape of accessible chromatin upon bacterial infection than in the previously defined STAT1 and IRF1 binding sites. Next, while these individual subfamilies appear to be contributing more than expected by chance to putative regulatory regions, the core TE classes they belong to, DNA and LTR, are comparatively poorly enriched overall, with a fold enrichment of only 1.25 and 1.03 (Figure S1). However, the Tigger3 and MER44 subfamilies all belong to the larger TcMar-Tigger family, a subdivision of DNA transposons which is among the most highly enriched TE families in its category.

Finally, we wanted to characterize the TE subfamilies according to their approximate age of insertion in the genome to determine whether the immune repeats have inserted early or late in evolution. We thus estimated the age of each repeat instance based on its similarity to the original sequence and compared the results between accessible and inaccessible repeats within each subfamily (Methods). Among the 14 grouped TEs, several subfamilies show a moderate difference in age between accessible and inaccessible instances (Table S3, Figure S2), as is the case for Tigger3b (*p* = 6.60E-05), and to a lesser extent, MER44B and THE1B (*p* = 0.028 and *p* = 0.035), where the accessible instances are slightly younger on average. Notably, we found that the four groups have inserted in the genome at different points in time (Figure 1e). Where the younger THE1 and Tigger3 subfamilies have inserted on average 57.2 and 72.2 million years ago, the MER44 and MLT1 repeats have been present for 84.9 and 124 million years, respectively. As such, immune TEs may have contributed immune regulatory functions multiple times in the course of evolution.

### Selected TE subfamilies harbour motifs for master regulators of the immune response

While binding sites of the IRF1 and STAT1 transcription factors were previously described in repeats in immune cells,^7^ the use of ATAC-Seq data enables the detection of multiple transcription factors by scanning for the presence of TF motifs within regions that become more open after infection using *HOMER.*^12^ Overall, the most significantly enriched motifs in our ATAC-Seq data belong to the AP-1 and NF-kB transcription factors, found respectively in 30.8% and 12.4% of all peaks enhanced upon infection (vs. 6.6% and 4.0% genome-wide, p=1e-2112 and p=1e-325).

We specifically wanted to assess the presence of TF motifs within the immune TE families identified, comparing TF motifs found in repeat instances that are present in differentially accessible chromatin more often than in instances found in the rest of the genome (see Methods). We selected the top 30 motifs that were found in over 1/3 of accessible instances in at least one repeat subfamily and showed at least a 20% increase compared with inaccessible repeats (Figure 2a). Several motifs showed a subfamily-specific enrichment (e.g. GATA in MER44 and NFkB in THE1), while other motifs were found to be shared among TE subfamilies of the same type (e.g. Fox/Foxo motifs in MER44D and Tigger3-type repeats). Notably, we observed that the motifs for the AP-1 family of transcription factors (including BATF, Fra1, Atf3 and JunB), are enriched across all 14 subfamilies. Actually, 65.1% of accessible instances contain at least one of these motifs. It is comparatively depleted in activated repeats which do not belong to the immune TEs, where it is found in only 48.5% of instances. The enrichment is particularly significant for MER44 and THE1 (Figure 2b), where the motifs are found in over 93% of MER44B and THE1C peaks, with less than 42% and 65% of non-accessible instances harbouring the motifs in each family, respectively. While the AP-1 motifs are present overall in accessible chromatin upon infection, their overrepresentation in these particular TE subfamilies suggests an association with immune-specific repeats. The second most significant enrichment is observed for the NF-kB motif, which is found almost exclusively in THE1 repeats (Figure 2b), where it is found in 67% and 83% of THE1B and THE1C instances. The enrichment of these motifs is reasonable overall within immune-specific accessible chromatin, as the significant role of these TFs in immunity is well described in the literature, both for AP-1^13–15^ and NF-kB.^16–18^ Interestingly, functional NF-kB and AP-1 binding sites have both been previously validated in THE1B instances involved in gene reactivation of Hodgkin’s lymphoma.^19^

**Figure 2.**
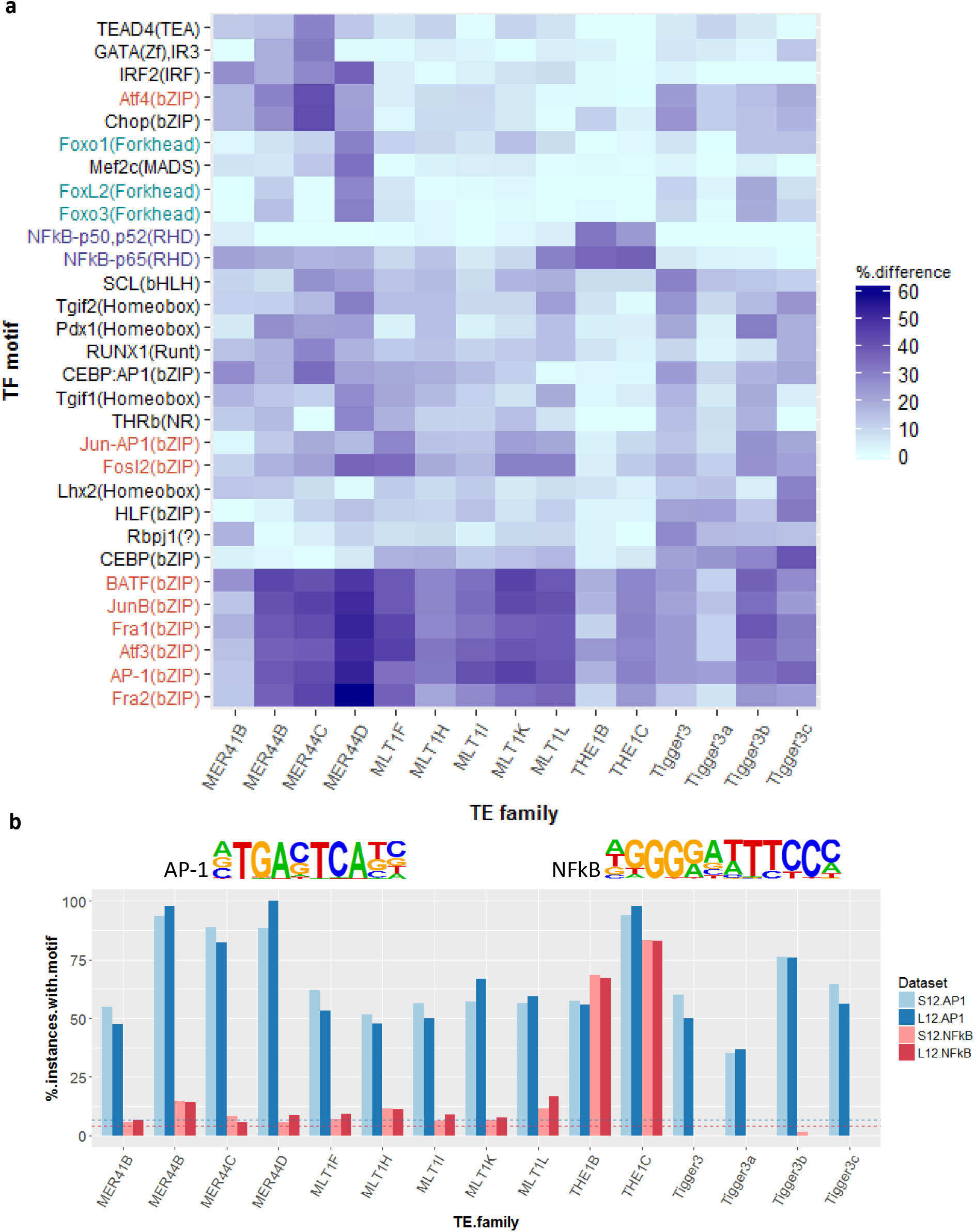
(a) Absolute difference between proportion of accessible repeats containing the top most enriched TF motifs compared to their non-accessible counterparts within the same subfamily. (b) Proportion of accessible repeats with the AP-1 and NF-kB motifs. Dashed lines show proportion of background genome sequences containing the AP-1 (blue) and NFkB (red) motifs.

### A subset of TEs are enriched near differentially expressed genes

To clarify the role of TEs in the regulation of immunity, we are particularly interested in PARs found in proximity to genes which are up- or down-regulated following infection. To determine the presence of PARs specifically near differentially expressed genes (DEGs), we obtained lists of DEGs from Nédélec *et al.*^8^ for both *Salmonella* and *Listeria* infected cells. We were able to confirm that accessible chromatin is enriched overall within 100kb of DEGs (*p*=4.94e-324, see Methods). We then compared the presence of accessible and inaccessible repeats near DEGs and computed the enrichment of all TE subfamilies contributing at least 10 instances to accessible chromatin. This analysis identified a new set of 85 and 28 subfamilies enriched in the S12 and L12 samples, with 23 of these subfamilies enriched in both conditions (Figure S3, Table S4-S5). Notably, the previously identified THE1B repeat is the second most significantly enriched subfamily, with 57 instances near DEGs found in the S12 samples (we would have only expected 16, *p*-value = 3.77e-16). The THE1C repeats are also observed, which further supports the possibility of a shared activating element within these TE subfamilies.

Similarly, the MLT1 repeats are also enriched near DEGs, with the MLT1K family presenting as the most highly enriched subfamily (*p* = 8.06e-5 and *p* = 6.38e-4). For example, an interesting MLT1G instance, which is accessible in S12 but not in L12, can be found within 3 kb of the DAPP1 gene, which is upregulated only in S12 cells. Another example is an MLT1K instance 2 kb upstream of the TXN gene, which has been described as a player in redox reactions in stimulated macrophages.^20,21^ As there are over 200 such PARs found within 100 kb of DEGs, this large group of repeats offers several other candidates for contribution to genetic regulation (see Tables 1 and S6-S7 for top PARs nearest DEGs).

**Table 1.** Top 3 PARs nearest differentially expressed genes’ transcription start sites for each TE group, present in both S12 and L12 samples. See Table S4 for complete list.

| TE.family | Locus | Gene | Distance to TSS (bp) |
| --- | --- | --- | --- |
| MER44D | chr8:90,794,231-90,794,648 | RIPK2 | 1,915 |
| MER44B | chr8:22,216,286-22,216,553 | SLC39A14 | 8,207 |
| MER44B | chr10:26,970,664-26,971,209 | PDSS1 | 15,377 |
| MLT1H | chr1:167,754,291-167,754,845 | MPZL1 | 1,457 |
| MLT1K | chr7:89,786,435-89,786,988 | STEAP1 | 2,665 |
| MLT1K | chr6:160,086,744-160,087,114 | SOD2 | 2,973 |
| THE1B | chr2:113,814,070-113,814,437 | IL36RN | 1,776 |
| THE1B | chr2:6,998,590-6,998,932 | CMPK2 | 2,455 |
| THE1B | chr12:113,426,846-113,427,200 | OAS2 | 8,848 |
| Tigger3c | chr10:90,595,119-90,595,493 | ANKRD22 | 12,561 |
| Tigger3b | chr9:117,583,234-117,584,446 | TNFSF15 | 31,634 |
| Tigger3b | chr3:172,279,540-172,279,935 | TNFSF10 | 44,395 |

Although THE1 and MLT1 subfamilies appear enriched both in accessible chromatin and near DEGs, it is surprising that the most significantly enriched TE subfamily, MER44B, is only moderately enriched near DEGs, with 19 and 5 notable instances within 100 kb of such genes for the two conditions. However, MER44 repeats may still be acting distally or contributing to gene regulation at earlier or later time points or through alternative mechanisms.

### A number of peak-associated repeats are found in QTL regions

To further associate PARs with the genes they potentially regulate, we evaluated their presence in regions where genetic variation affects gene expression in response to infection, as defined by quantitative trait locus (QTL) analysis. These regions, termed reQTLs, were previously established by Nédélec *et al.* through mapping of common SNP genotypes from 175 individuals to the magnitude of change in expression levels upon infection. We retrieved the most significant reQTLs for the S12 and L12 samples, 503 and 244 respectively. As SNPs in linkage disequilibrium (LD) are commonly grouped within haplotype blocks,^22,23^ we expanded each reQTL to encompass a larger interval based on its LD block. We used this method as an alternative to direct overlap with SNPs, as the latter returns no TEs overlapping the QTL themselves, or 53 and 15 TEs overlapping any SNP in the corresponding LD block for S12 and L12, respectively. This is insufficient to perform statistical analysis. We therefore used the entire interval between the two most distal SNPs belonging to each LD block but extending no farther than 50 kb away from the original SNP, and we defined these as QTL-regions (see Methods). We first assessed the enrichment of all ATAC-Seq peaks in these regions, as we expect accessible chromatin to localize to regions with a functional impact on gene expression. We found that accessible chromatin does overlap the defined QTL-regions more than expected by chance (1.78 and 2.1-fold, *p*=1.24e-41 and *p*=1.31e-22 for S12 and L12, respectively). We then reproduced the analysis by limiting the accessible chromatin to repeat-associated peaks and confirmed that these elements are also enriched in the QTL-regions (2.37 and 2.85-fold, *p*=5.66e-23 and *p*=9.63e-12).

As we have already defined above a set of 34 TE subfamilies enriched in accessible chromatin (Tables S1-S2), we hypothesize that these immune-specific repeats are more likely contributing to regulatory networks and thus expect them to be overrepresented in QTL-regions. We therefore classified PARs by level of enrichment in accessible chromatin to create 2 distinct TE subgroups: immune (belonging to the 34 enriched subfamilies described above) and non-immune (belonging to the remaining subfamilies). We then compared the enrichment of PARs in QTL-regions separately for each subgroup (Figure S4). We observe a higher enrichment for immune TEs in the S12 sample (3.17-fold*, p*=1.34e-13) than for non-immune TEs (2.1-fold, *p*=1.91e-12). Conversely, there appears to be a slightly greater enrichment for non-immune TEs (2.93-fold, *p*=8.52e-10) than immune TEs (2.6-fold, *p*=1.97e-3) in the L12 sample, although the small number of immune TEs overlapping QTL-regions is insufficient to confirm significance for this condition. To further quantify the association between significantly enriched TE families and their presence in the S12 QTL-regions, we performed a chi-square test to evaluate the correlation between the type of TE family (immune vs. non-immune) and the type of peak (overlapping or not overlapping a QTL-region). We confirm a moderate association between TE category and peak type (*p*=0.0239), which further supports that the immune-specific subfamilies we identified associate with reQTLs in their haplotype block in the S12 sample.

Finally, we observe interesting instances of PARs overlapping QTL-regions, including several of the 14 TE subfamilies identified above from the MER44, THE1, Tigger3 and MLT1 subfamilies (Figure 3, Tables S8-S9). Notably, the most interesting MER44B instance, falling 15 kb upstream of PDSS1, also overlaps its corresponding QTL-region (Figure S5). We also observed more distal examples, such as a THE1B instance 150 kb downstream of BASP1, and an MLT1F2 instance 70 kb upstream of GADD45A, both repeats overlapping the corresponding QTL-region for those genes. Together, these results suggest that the significant TE subfamilies which we have identified are not only enriched in accessible chromatin but also contributing to regions that are genetically affecting gene expression based on a QTL analysis.

**Figure 3.**
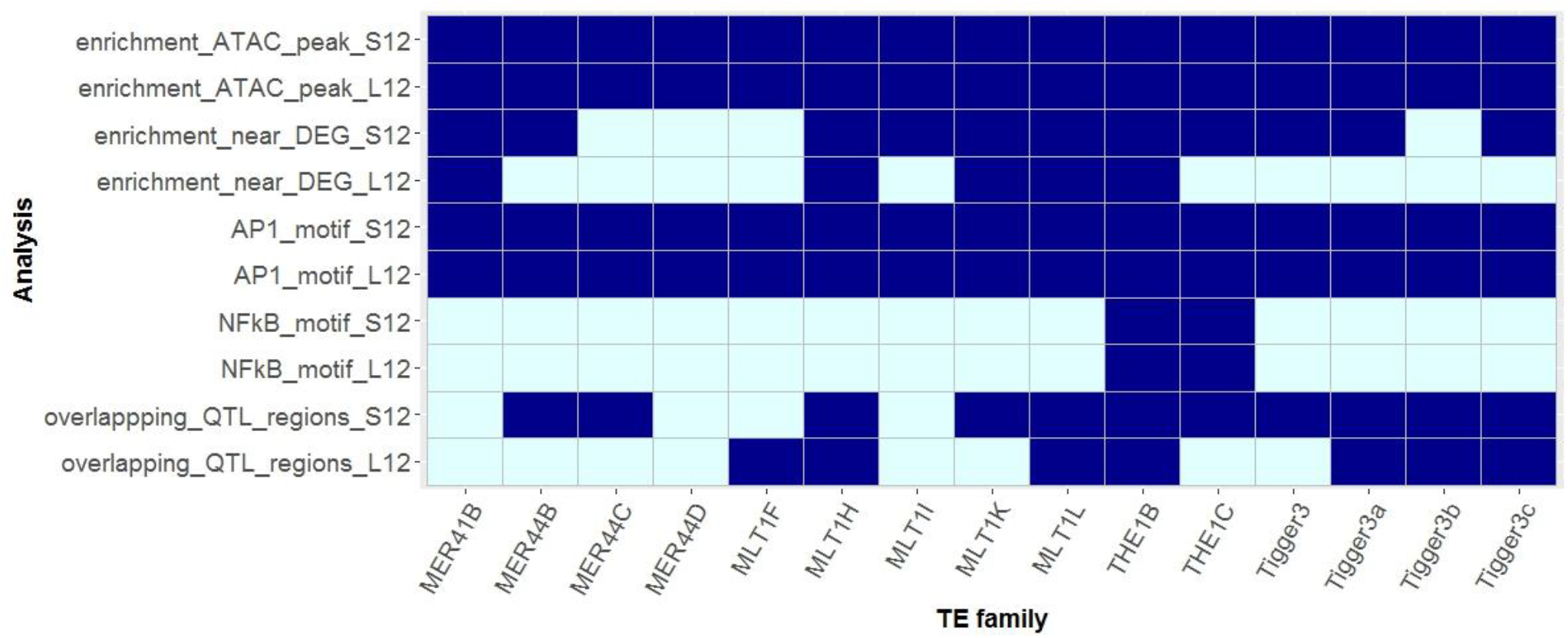
Summary of all results for MER41B and the 14 recurrently overrepresented TE subfamilies. Enrichment indicated in dark blue. Enrichment of the AP-1 and NF-kB motifs is defined as p-value < 0.05 in *Homer*, motif found in at least 30% of PARs and a difference of at least 20% in accessible vs. inaccessible repeats. Bottom 2 rows indicate if the subfamily has at least 1 PAR overlapping a QTL region.

## CONCLUSION

In this study, we advanced our understanding of the TEs contributing to the accessible chromatin enhanced by infection in macrophages stimulated *in-vitro*. We specifically found a subset of families that are enriched in differentially accessible chromatin and near differentially accessible genes, with multiple related instances from the MER44, THE1, Tigger3 and MLT1 subfamilies. We identified binding motifs for the AP-1 immune transcription factor that were enriched in these subfamilies, and most particularly in the MER44 and THE1C subfamilies. NF-kB, another master regulator of immunity, also has enriched binding motifs specifically in the THE1B and THE1C subfamilies, which are the most highly enriched subfamilies near differentially expressed genes.

We also used a set of previously defined regulatory QTLs as a complementary analysis to define potential regions of influence on gene regulation. Although we do not have sufficient power to find statistically significant overlap between peak-associated repeats and the QTLs themselves, using linkage disequilibrium to define regions of influence flanking the QTLs show that peak-associated repeats contribute to putative regulatory regions more than expected. This also identifies a number of specific TE instances that would be interesting targets for biological validation.

While the THE1B subfamily has a known role as an activated set of repeats in Hodgkin’s lymphoma,^19^ it is interesting to note here that our most highly enriched subfamilies, MER44B, C and D, have not been previously identified. It would therefore be interesting to further explore these findings to determine if these small yet significant subfamilies have a regulatory role in other cell types and conditions.

## METHODS

### ATAC-Seq dataset processing

We obtained raw ATAC-Seq datasets from Nédélec *et al.*,^8^ with two replicates for each of three conditions: macrophages infected with *L. monocytogenes* (i), infected with *S. typhimurium* (ii), and non-infected (iii) at 12 hours. The FASTQ files were processed using the MUGQIC ChIP-Seq pipeline v.2.2.1 to obtain narrow peaks. Briefly, the pipeline pools the replicates for each condition and was adapted to retain reads longer than 37 bp for subsequent processing. Sequencing adaptors were trimmed using *Trimmomatic v0.36*^24^ before aligning to the hg19 human reference with *BWA v0.7.12*.^25^ Next, unique reads were filtered by mapping quality using *Samtools v1.3.1*^26^ and duplicates were marked with *Picard v2.0.1*.^27^ Narrow peaks were called with *MACS2 v2.1.0*,^28^ using profiles from non-infected cells as background (input) to obtain differentially accessible peaks. Peaks with a q-value of less than 0.0005 and shorter than 1 kb in width were kept for downstream analyses.

### TE enrichment in accessible chromatin

Enrichment of each repeat family was computed within accessible chromatin. We obtained repeat annotations from the RepeatMasker track in the UCSC Table Browser and removed coordinates corresponding to tRNAs, simple repeats and tandem repeats. We intersected the remaining 4,506,876 repetitive sequences with S12 and L12 peak summits (center 1 bp) using *intersectBed* from the *BEDtools* suite, with the –u option specified to avoid duplicates.^29^ Strand information was not considered for any of the analyses, as it was unavailable for the peak calls.

We then computed the enrichment in accessible chromatin of each TE subfamily using a one-sided binomial test, comparing the number of TE instances overlapping peak summits with its expected counterpart. The expected distribution was obtained by shuffling the true peaks randomly across the genome for 1,000 iterations, while maintaining a comparable distribution of peak locations. More specifically, the original peaks were annotated by categories based on their distance to RefSeq genes: 5’UTR, exon, intron, TSS (< 1 kb upstream), promoter (1-5 kb upstream), proximal (5-10 kb upstream or < 10 kb downstream), distal (10-100 kb upstream or downstream) and desert (> 100 kb upstream or downstream). We then separated the peaks based on their annotation and shuffled them separately with *shuffleBed*, using the –incl and –excl parameters to restrict the randomization within the corresponding genomic regions defined above. Each of the 1,000 shuffled peak sets were overlapped with the RepeatMasker annotations and the number of peaks overlapping instances in each TE subfamily, large family and class was obtained.

The mean of the expected counts was taken and compared with the observed counts for each TE family using the binom.test() function in R and resulting p-values were adjusted for multiple testing using the Benjamini-Hochberg method with the p.adjust() function from the *multtest* R package.^30^ TE families with a q-value of less than 0.05 were kept for further consideration.

### Estimation of TE age

We obtained an estimate of the age of each TE instance based on the sequence divergence (base mismatches in parts per thousand as defined by the milliDiv value from RepeatMasker). The milliDiv value of each instance was divided by 2.2e-9, the substitution rate for the human genome, to obtain the final age. Each TE instance in RepeatMasker was then classified as either accessible (overlapping ATAC-seq peaks) or inaccessible and the mean age was taken for each group and TE subfamily separately.

### Transcription factor motif analysis

We scanned the PARs for known TF motifs using *findMotifsGenome.pl* from *HOMER v4.9.1.^12^* This tool uses a hypergeometric test for each TF to compare the number of motifs found in a target set of genomic regions with that found in a specified set of background regions. We first compared the motifs detected in all ATAC-Seq peaks to the STAT1 and IRF1 ChIP-Seq datasets published by Chuong *et al*. We restricted the width of all peaks to the center 200 bp and separated them in two categories: (1) regions shared with the ChIP-Seq datasets (overlapping the STAT1 and IRF1 summits), and (2) regions which are specific to the ATAC-Seq datasets. *findMotifsGenome.pl* was run twice using alternatively the shared regions and the ATAC-Seq-specific regions as target and background sets, respectively, to detect the most overrepresented motifs in each dataset.

Next, we sought TF motifs which were enriched specifically in PARs belonging to the subset of 14 most significant TE families we identified. We defined the 14 target sets separately and ran *findMotifsGenome.pl* using a custom background set of all TEs in each subfamily not overlapping accessible chromatin. The –size parameter was set to *given* to include the entire TE sequence both for the target and background regions for all analyses. Other parameters were left as default.

Finally, we used *HOMER’*s *scanMotifGenomeWide.pl* to extract all loci containing the motifs for AP-1 and NF-kB, the most significant TFs detected. The motifs were specified using HOMER’s pre-defined motif files for AP-1(bZIP), Atf3(bZIP), BATF(bZIP), Fra1(bZIP), Fra2(bZIP) and NFkB-p65(RHD). The motifs were subsequently overlapped with all repeats and PARs separately using intersectBED, with the –s option specified to intersect regions found on the same strand.

### TE enrichment near differentially expressed genes

The enrichment of TEs near differentially expressed genes (DEGs) was computed separately for each repeat subfamily. We defined DEGs as genes with a log2 fold-change in expression greater than 2 or lesser than −2. Lists of 1574 and 667 DEGs were thus obtained from the Nédélec *et al.* dataset for S12 and L12 samples, respectively. For each TE subfamily, the number of RAPs within 100 kb upstream or downstream of a DEG was obtained using the *distanceToNearest()* function from the *GenomicRanges* R package.^31^ TE subfamilies with less than 10 instances in accessible chromatin were removed from the analysis. A random set of the same size as the number of corresponding RAPs was taken 1,000 times from all TEs not overlapping peaks and processed similarly to obtain an average expected number of TEs near DEGs. A binomial test was then performed comparing observed and expected counts, and p-values were adjusted as previously described. Individual repeat instances were visualized manually using the Integrative Genomics Viewer (IGV, v1.4.2).^32^

### Definition of QTL-regions and overlap with PARs

We defined QTL-regions using *rAggr*^33^ for all reQTLs published by Nedelec *et al.* for both S12 and L12 samples. *rAggr* was adapted by Edlund *et al.* from the Haploview software^34^ to provide proxy markers for SNPs in linkage-disequilibrium based on the 1000 Genomes (Phase 3) genotype data. The rsIDs for all 503 and 244 top SNPs were provided as input directly in the web browser. The software was run with the CEU and YRI populations selected, a maximal distance of 50 kb, and all other options left as default (minimum mean allele frequency = 0.001, r^2^ range of 0.8 to 1.0, maximum number of Mendelian errors = 1, HW p-value cutoff = 0, and minimum genotype percentage = 75). The interval between the resulting two most distant proxy SNPs was taken to define the QTL-regions for each original reQTL. The reQTLs with no proxy SNPs returned by *rAggr* were extended to encompass a 1 kb interval flanking each reQTL, thus creating intervals between 1-100 kb in width across all QTLs.

We then computed the enrichment of all ATAC-Seq peaks in QTL-regions, as well as the subset of peaks overlapping TEs (corresponding to PARs). The statistical method is described above for the enrichment of repeats, comparing the true overlaps with an expected distribution obtained by shuffling the peaks across the genome while respecting their corresponding annotations. We also compared the categories of PARs according to the subfamilies they belong to (with immune-specific TEs belonging to the 34 families enriched in accessible chromatin) and the type of peak they overlap (either peaks found in or absent from the QTL-regions). The association between these groups was evaluated using the chisq.test() function in R.

## AUTHOR CONTRIBUTIONS

**L.B**. participated in the design of the study, conceived and performed the computational analyses, and drafted the manuscript. **L.B.B.** provided the data, suggested analyses and critically revised the manuscript. **G.B.** conceived, designed and directed the study, and edited the manuscript. All authors revised the paper and gave final approval for publication.

## ACKNOWLEDGEMENTS

We thank all members of the Bourque Lab for helpful comments and discussion, and David Venuto for advice on the peak enrichment analysis. We also thank Yohann Nédélec and Alain Pacis from the Barreiro Lab for providing access and support for the ATAC-Seq data. This work was supported by funding from the Canadian Institute for Health Research [CIHR-MOP-115090]; Fonds de Recherche Santé Québec [FRSQ-25348 to G.B.]. Data analyses were enabled by compute and storage resources provided by Compute Canada and Calcul Québec.

