## Supplementary Figures for "Transposable elements have contributed human regulatory regions that are activated upon bacterial infection"

a

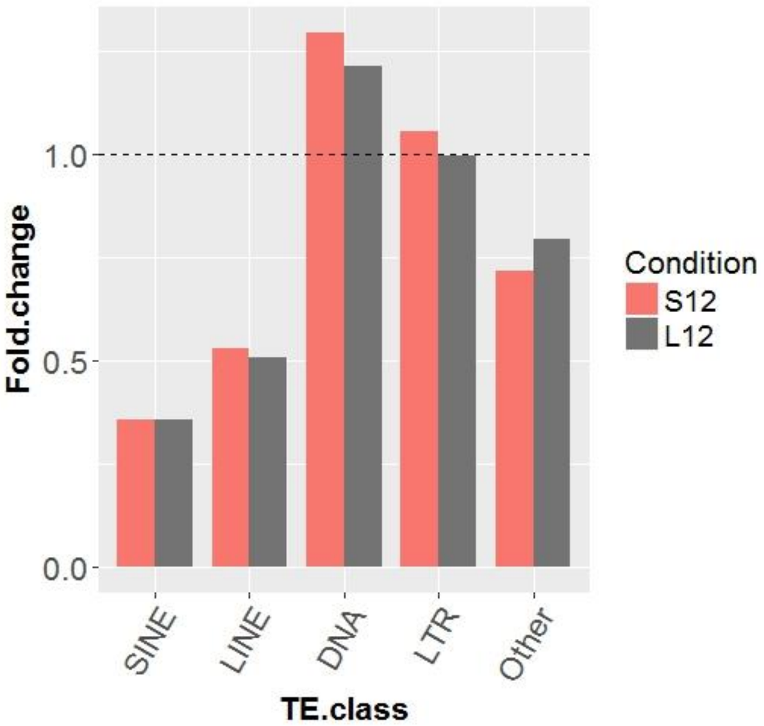


b

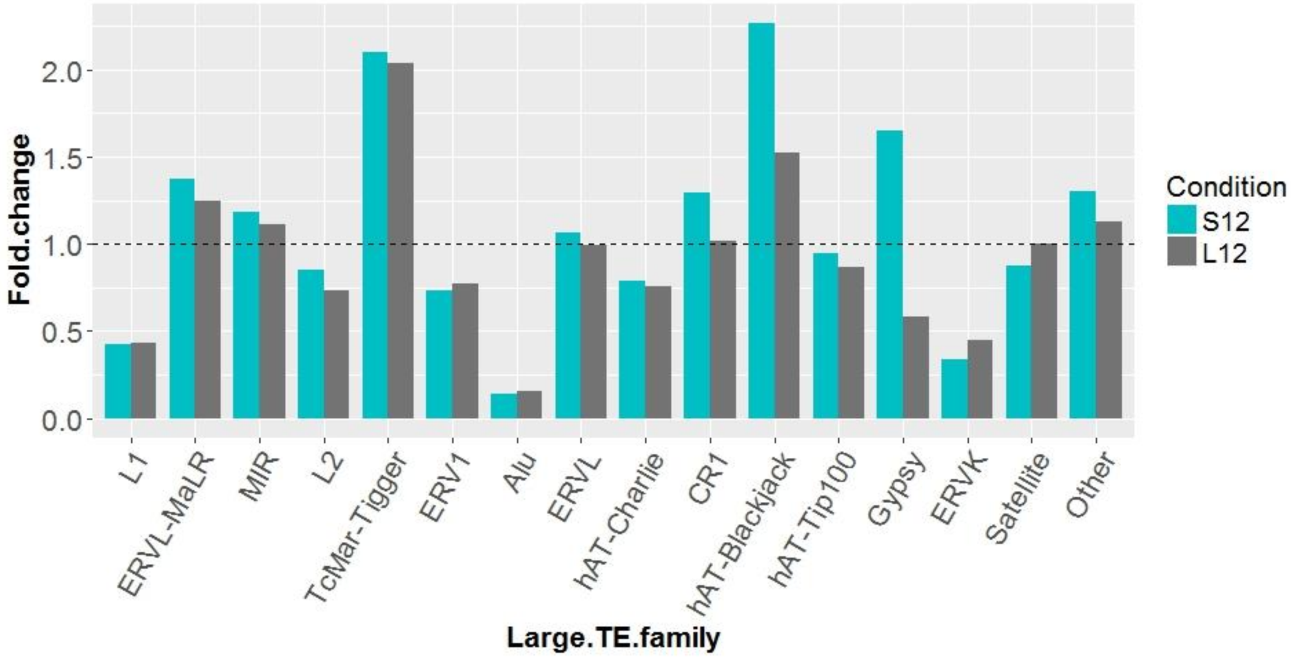


**Figure S1.** Enrichment of the 4 major TE classes (a) and large TE families/categories (b) in accessible chromatin (expected/observed number of TEs in ATAC-Seq peaks).

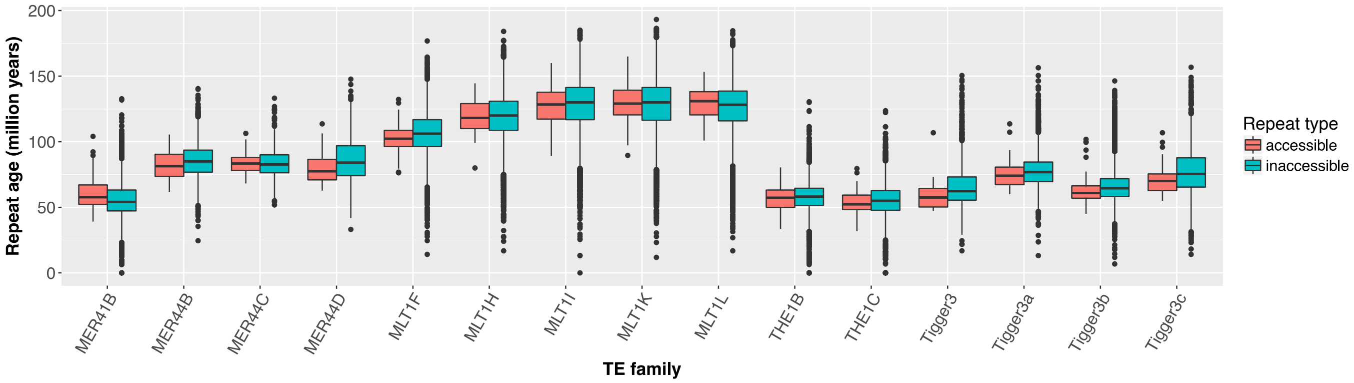


**Figure S2.** Age of TE instances in accessible and inaccessible chromatin for each repeat family.

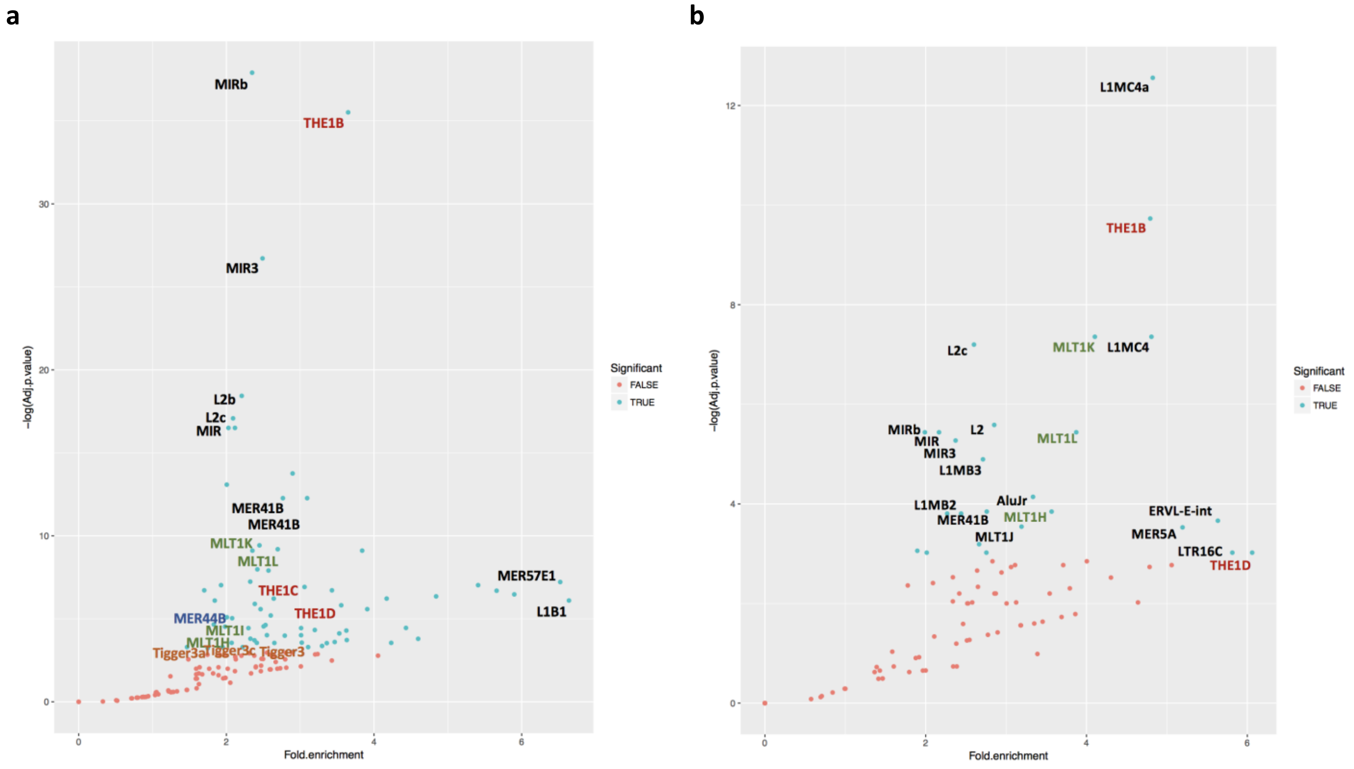


**Figure S3.** (a-b) Enrichment of PARs near differentially expressed genes in S12 and L12 samples.

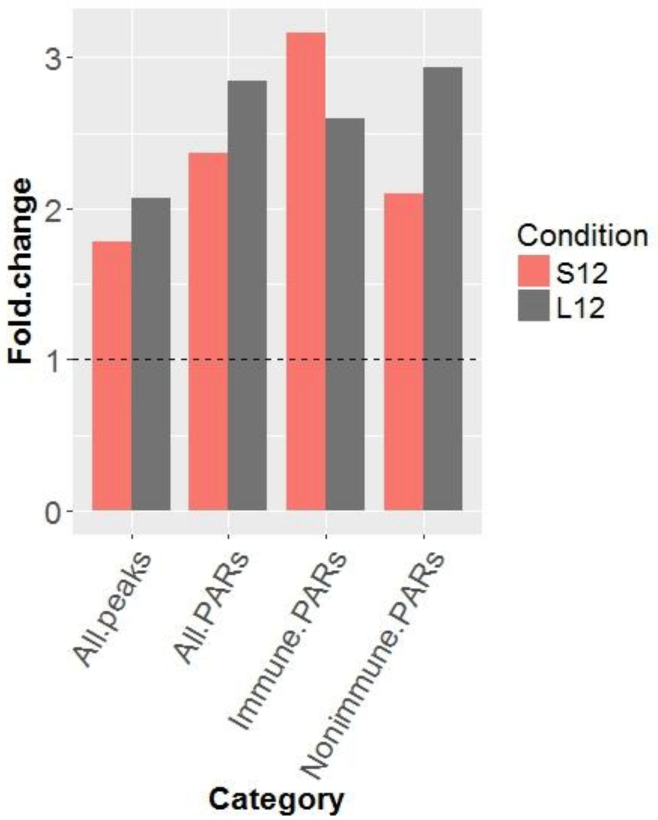


**Figure S4.** Enrichment of all ATAC-Seq peaks, PARs, immune PARs and non-immune PARs in QTL-regions.

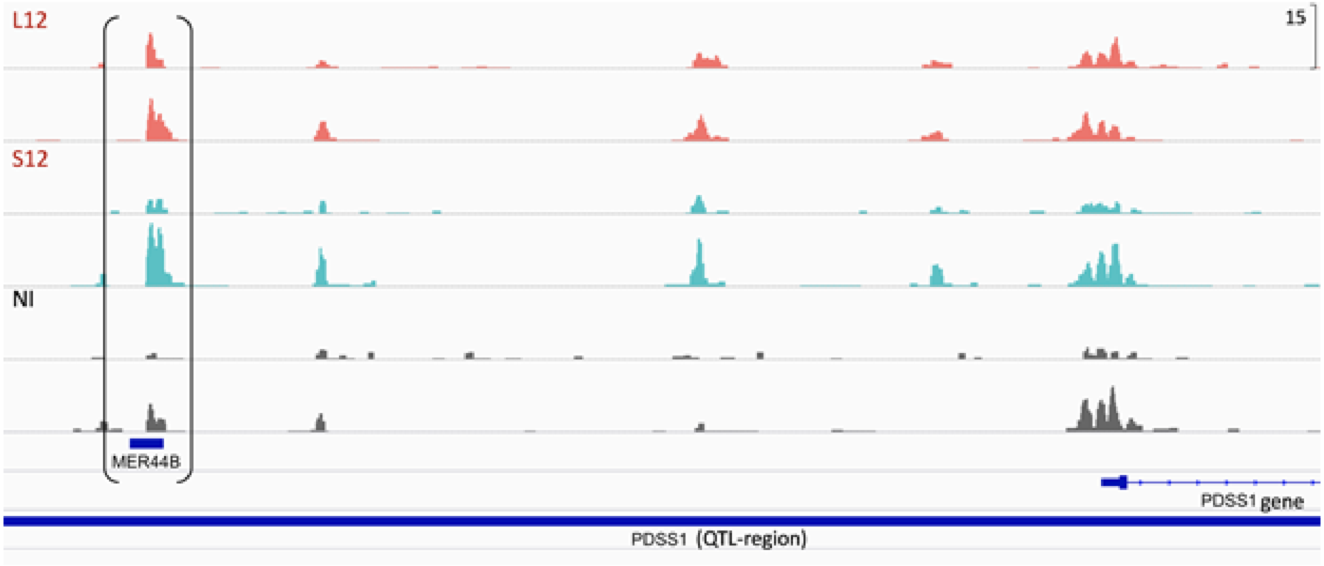


**Figure S5.** Example of accessible PAR overlapping its corresponding QTL-region near a DEG. In this case, the MER44B instance is 15kb upstream of the PDSS1 gene.
