## Supplementary Tables 1 and 2 for "Transposable elements have contributed human regulatory regions that are activated upon bacterial infection"

Table S1. Enriched TE families in accessible chromatin after infection for the S12 samples

| TE.family | Observed.instances | Expected.instances | Fold.enrichment | Prob.success | P.value | Q.value |
|---|---|---|---|---|---|---|
| MER44B | 95 | 6 | 15.83333333 | 0.00281558 | 3.31E-78 | 3.36E-75 |
| THE1B | 205 | 54 | 3.796296296 | 0.002407168 | 1.46E-55 | 7.42E-53 |
| L1MB2 | 148 | 33 | 4.484848485 | 0.003625975 | 6.21E-49 | 2.10E-46 |
| Tigger3b | 118 | 27 | 4.37037037 | 0.0041392 | 2.19E-38 | 5.54E-36 |
| Tigger7 | 59 | 7 | 8.428571429 | 0.002245749 | 3.51E-34 | 7.13E-32 |
| MER41B | 71 | 11 | 6.454545455 | 0.003856942 | 1.07E-33 | 1.81E-31 |
| MER44D | 35 | 2 | 17.5 | 0.002659574 | 2.33E-31 | 3.37E-29 |
| Tigger3a | 74 | 14 | 5.285714286 | 0.002627135 | 1.44E-29 | 1.82E-27 |
| MER44C | 36 | 4 | 9 | 0.004683841 | 1.44E-22 | 1.62E-20 |
| MER81 | 32 | 3 | 10.66666667 | 0.000817661 | 3.45E-22 | 3.50E-20 |
| MLT1F | 58 | 14 | 4.142857143 | 0.003258087 | 1.11E-18 | 1.02E-16 |
| Tigger3c | 34 | 5 | 6.8 | 0.002110595 | 1.30E-17 | 1.10E-15 |
| MLT1F2 | 62 | 18 | 3.444444444 | 0.002982107 | 3.87E-16 | 3.02E-14 |
| MLT1I | 62 | 19 | 3.263157895 | 0.00171341 | 4.49E-15 | 3.25E-13 |
| MLT1L | 69 | 23 | 3 | 0.00190492 | 7.43E-15 | 5.02E-13 |
| HSATII | 16 | 1 | 16 | 0.002506266 | 1.43E-14 | 9.05E-13 |
| MLT1K | 90 | 37 | 2.432432432 | 0.002035987 | 1.23E-13 | 7.31E-12 |
| L1MB7 | 171 | 95 | 1.8 | 0.004095887 | 1.35E-12 | 7.60E-11 |
| THE1C | 65 | 24 | 2.708333333 | 0.002430626 | 3.40E-12 | 1.82E-10 |
| MLT1H | 59 | 21 | 2.80952381 | 0.002080444 | 8.00E-12 | 4.06E-10 |
| MIR3 | 197 | 118 | 1.669491525 | 0.001298973 | 1.94E-11 | 9.37E-10 |
| MIRc | 226 | 142 | 1.591549296 | 0.001372087 | 5.01E-11 | 2.31E-09 |
| L1MB3 | 122 | 65 | 1.876923077 | 0.003743377 | 1.71E-10 | 7.54E-09 |
| MLT1F1 | 35 | 10 | 3.5 | 0.00304971 | 5.52E-10 | 2.33E-08 |
| LTR26 | 15 | 2 | 7.5 | 0.003246753 | 3.40E-09 | 1.38E-07 |
| LTR71A | 11 | 1 | 11 | 0.006024096 | 7.59E-09 | 2.96E-07 |
| AmnSINE1 | 14 | 2 | 7 | 0.001792115 | 2.76E-08 | 1.04E-06 |
| MER51A | 21 | 5 | 4.2 | 0.00522466 | 7.13E-08 | 2.58E-06 |
| MER57F | 10 | 1 | 10 | 0.002320186 | 1.02E-07 | 3.50E-06 |
| MER57E1 | 10 | 1 | 10 | 0.001996008 | 1.04E-07 | 3.50E-06 |
| MamSINE1 | 13 | 2 | 6.5 | 0.001199041 | 2.01E-07 | 6.56E-06 |
| MER45B | 18 | 4 | 4.5 | 0.003041825 | 2.31E-07 | 7.10E-06 |
| Tigger3 | 18 | 4 | 4.5 | 0.003292181 | 2.30E-07 | 7.10E-06 |
| MLT1J2 | 35 | 13 | 2.692307692 | 0.001877256 | 3.19E-07 | 9.52E-06 |
| Tigger12A | 9 | 1 | 9 | 0.001461988 | 1.08E-06 | 3.13E-05 |
| MLT1H2 | 24 | 8 | 3 | 0.001697073 | 3.63E-06 | 0.0001 |
| LTR8A | 29 | 11 | 2.636363636 | 0.003878702 | 4.43E-06 | 0.00012 |
| MLT1H1 | 26 | 10 | 2.6 | 0.002747253 | 1.71E-05 | 0.00046 |
| MLT1M | 14 | 4 | 3.5 | 0.00135318 | 7.51E-05 | 0.00195 |
| LTR44 | 7 | 1 | 7 | 0.004587156 | 7.75E-05 | 0.00196 |
| MER107 | 7 | 1 | 7 | 0.002785515 | 7.97E-05 | 0.00197 |
| Tigger12c | 9 | 2 | 4.5 | 0.001583531 | 0.00023 | 0.00563 |
| MLT1G | 20 | 8 | 2.5 | 0.002803083 | 0.00025 | 0.00582 |
| MER121 | 6 | 1 | 6 | 0.001081081 | 0.00059 | 0.01268 |
| Kanga1b | 6 | 1 | 6 | 0.002659574 | 0.00058 | 0.01268 |

| | | | | | | |
|---|---|---|---|---|---|---|
| MER41E | 6 | 1 | 6 | 0.002785515 | 0.00058 | 0.01268 |
| Charlie13b | 6 | 1 | 6 | 0.00245098 | 0.00058 | 0.01268 |
| LTR13 | 14 | 5 | 2.8 | 0.010183299 | 0.00065 | 0.01363 |
| Plat_L3 | 14 | 5 | 2.8 | 0.001350257 | 0.00069 | 0.0143 |
| MER77B | 12 | 4 | 3 | 0.00317965 | 0.00089 | 0.01813 |
| L1MB5 | 60 | 39 | 1.538461538 | 0.003950967 | 0.00105 | 0.02086 |
| MamRep6( | 18 | 8 | 2.25 | 0.001805869 | 0.00158 | 0.02961 |
| MLT1G1 | 21 | 10 | 2.1 | 0.002783964 | 0.00156 | 0.02961 |
| MLT1N2 | 21 | 10 | 2.1 | 0.001699524 | 0.00157 | 0.02961 |
| LTR16A | 28 | 15 | 1.866666667 | 0.002153316 | 0.0017 | 0.03125 |
| LTR8 | 27 | 15 | 1.8 | 0.0042337 | 0.00325 | 0.05876 |
| MER119 | 9 | 3 | 3 | 0.002487562 | 0.00375 | 0.06162 |
| MER117 | 14 | 6 | 2.333333333 | 0.00133452 | 0.0036 | 0.06162 |
| CR1_Mam | 9 | 3 | 3 | 0.001734104 | 0.00377 | 0.06162 |
| LTR54 | 9 | 3 | 3 | 0.002676182 | 0.00375 | 0.06162 |
| LTR84b | 9 | 3 | 3 | 0.00231303 | 0.00376 | 0.06162 |
| LTR72B | 5 | 1 | 5 | 0.005882353 | 0.00353 | 0.06162 |
| HERVP71A- | 7 | 2 | 3.5 | 0.007968127 | 0.00434 | 0.06991 |
| Charlie20a | 7 | 2 | 3.5 | 0.002277904 | 0.00448 | 0.07097 |
| MLT1G3 | 15 | 7 | 2.142857143 | 0.002583979 | 0.00565 | 0.08819 |
| MLT1J | 50 | 34 | 1.470588235 | 0.002226588 | 0.00592 | 0.09093 |
| MER21C | 35 | 22 | 1.590909091 | 0.003999273 | 0.00626 | 0.09478 |
| Tigger13a | 16 | 8 | 2 | 0.002565747 | 0.00815 | 0.12153 |
| LTR67B | 13 | 6 | 2.166666667 | 0.001614205 | 0.00877 | 0.12892 |
| Tigger15a | 17 | 9 | 1.888888889 | 0.001777953 | 0.01104 | 0.15989 |
| MER21B | 21 | 12 | 1.75 | 0.004290311 | 0.01143 | 0.16327 |
| Charlie25 | 8 | 3 | 2.666666667 | 0.002782931 | 0.01178 | 0.16376 |
| Charlie10 | 8 | 3 | 2.666666667 | 0.002659574 | 0.01179 | 0.16376 |
| MER112 | 14 | 7 | 2 | 0.0016 | 0.01274 | 0.17462 |
| MER68 | 11 | 5 | 2.2 | 0.003333333 | 0.01354 | 0.18312 |
| LTR40c | 6 | 2 | 3 | 0.003322259 | 0.01638 | 0.20911 |
| LTR45C | 6 | 2 | 3 | 0.004705882 | 0.01631 | 0.20911 |
| LTR88b | 6 | 2 | 3 | 0.001215805 | 0.0165 | 0.20911 |
| Looper | 6 | 2 | 3 | 0.003738318 | 0.01636 | 0.20911 |
| LTR43 | 6 | 2 | 3 | 0.004987531 | 0.01629 | 0.20911 |
| LTR13_ | 4 | 1 | 4 | 0.022727273 | 0.01759 | 0.21802 |
| X3_LINE | 4 | 1 | 4 | 0.001526718 | 0.01889 | 0.21802 |
| LTR24 | 4 | 1 | 4 | 0.003246753 | 0.01879 | 0.21802 |
| MLT2E | 4 | 1 | 4 | 0.002557545 | 0.01883 | 0.21802 |
| LTR88a | 4 | 1 | 4 | 0.001090513 | 0.01892 | 0.21802 |
| MamGypLT | 4 | 1 | 4 | 0.001331558 | 0.01891 | 0.21802 |
| LTR90A | 4 | 1 | 4 | 0.002457002 | 0.01884 | 0.21802 |
| LTR72 | 4 | 1 | 4 | 0.006535948 | 0.01859 | 0.21802 |
| L3 | 100 | 81 | 1.234567901 | 0.0017994 | 0.02255 | 0.25693 |
| MER63A | 10 | 5 | 2 | 0.001421666 | 0.03172 | 0.35743 |
| HERV9-int | 20 | 13 | 1.538461538 | 0.012560386 | 0.04164 | 0.46403 |
| LTR16A1 | 11 | 6 | 1.833333333 | 0.002136752 | 0.04244 | 0.46781 |

| | | | | | | |
|---|---|---|---|---|---|---|
| MER20 | 39 | 29 | 1.344827586 | 0.001728969 | 0.04363 | 0.4757 |
| Kanga1a | 5 | 2 | 2.5 | 0.002369668 | 0.05244 | 0.53787 |
| Tigger14a | 5 | 2 | 2.5 | 0.00154202 | 0.05251 | 0.53787 |
| MamGypLT | 5 | 2 | 2.5 | 0.001647446 | 0.0525 | 0.53787 |
| LTR81A | 5 | 2 | 2.5 | 0.002059732 | 0.05247 | 0.53787 |
| LTR87 | 5 | 2 | 2.5 | 0.00286123 | 0.05239 | 0.53787 |
| L1MEg1 | 5 | 2 | 2.5 | 0.002136752 | 0.05246 | 0.53787 |
| MER65D | 3 | 1 | 3 | 0.002109705 | 0.08011 | 0.67247 |
| MLT2B3 | 14 | 9 | 1.555555556 | 0.002716571 | 0.07358 | 0.67247 |
| LTR81C | 3 | 1 | 3 | 0.001375516 | 0.08017 | 0.67247 |
| MER96 | 3 | 1 | 3 | 0.000784314 | 0.08023 | 0.67247 |
| Tigger3d | 3 | 1 | 3 | 0.001666667 | 0.08015 | 0.67247 |
| HERVL66-ir | 3 | 1 | 3 | 0.009433962 | 0.07943 | 0.67247 |
| MamRep56 | 3 | 1 | 3 | 0.001257862 | 0.08019 | 0.67247 |
| MER94 | 9 | 5 | 1.8 | 0.001042101 | 0.06799 | 0.67247 |
| MER91C | 3 | 1 | 3 | 0.000614251 | 0.08024 | 0.67247 |
| LTR26E | 3 | 1 | 3 | 0.003289474 | 0.08 | 0.67247 |
| MLT2B5 | 3 | 1 | 3 | 0.003030303 | 0.08002 | 0.67247 |
| MLT2F | 9 | 5 | 1.8 | 0.002608242 | 0.06784 | 0.67247 |
| MER6B | 3 | 1 | 3 | 0.001472754 | 0.08017 | 0.67247 |
| Tigger12 | 3 | 1 | 3 | 0.002159827 | 0.0801 | 0.67247 |
| LTR16B | 3 | 1 | 3 | 0.001445087 | 0.08017 | 0.67247 |
| MER57E2 | 3 | 1 | 3 | 0.013157895 | 0.07908 | 0.67247 |
| MER51D | 3 | 1 | 3 | 0.006451613 | 0.0797 | 0.67247 |
| MamGyp-ir | 3 | 1 | 3 | 0.002923977 | 0.08003 | 0.67247 |
| MER99 | 3 | 1 | 3 | 0.003378378 | 0.07999 | 0.67247 |
| MER72B | 3 | 1 | 3 | 0.005524862 | 0.07979 | 0.67247 |
| Eulor6D | 3 | 1 | 3 | 0.013513514 | 0.07904 | 0.67247 |
| LTR73 | 3 | 1 | 3 | 0.004608295 | 0.07988 | 0.67247 |
| Charlie1 | 10 | 6 | 1.666666667 | 0.003120125 | 0.0836 | 0.68477 |
| MARNA | 10 | 6 | 1.666666667 | 0.001791045 | 0.08374 | 0.68477 |
| Ricksha_c | 6 | 3 | 2 | 0.002423263 | 0.08367 | 0.68477 |
| MER66B | 7 | 4 | 1.75 | 0.002787456 | 0.11038 | 0.87475 |
| LTR33A_ | 7 | 4 | 1.75 | 0.002411091 | 0.11042 | 0.87475 |
| Cheshire | 7 | 4 | 1.75 | 0.003940887 | 0.11026 | 0.87475 |
| Tigger6a | 7 | 4 | 1.75 | 0.00516129 | 0.11013 | 0.87475 |
| MLT1J1 | 12 | 8 | 1.5 | 0.001624365 | 0.11175 | 0.87839 |
| L2b | 210 | 193 | 1.088082902 | 0.001970675 | 0.11814 | 0.91741 |
| L1MB8 | 74 | 64 | 1.15625 | 0.00384893 | 0.11852 | 0.91741 |
| L1MD3 | 20 | 15 | 1.333333333 | 0.002828054 | 0.12447 | 0.95612 |
| MamRep15 | 4 | 2 | 2 | 0.001586043 | 0.14273 | 0.99815 |
| LTR89 | 4 | 2 | 2 | 0.001731602 | 0.14272 | 0.99815 |
| L1MB4 | 41 | 34 | 1.205882353 | 0.003658668 | 0.13317 | 0.99815 |
| ERV3-16A3 | 4 | 2 | 2 | 0.001851852 | 0.14271 | 0.99815 |
| LTR40a | 8 | 5 | 1.6 | 0.002544529 | 0.13311 | 0.99815 |
| MER74A | 8 | 5 | 1.6 | 0.003160556 | 0.13304 | 0.99815 |
| LTR52 | 4 | 2 | 2 | 0.001850139 | 0.14271 | 0.99815 |

| | | | | | | |
|---|---|---|---|---|---|---|
| LTR33C | 4 | 2 | 2 | 0.002159827 | 0.14268 | 0.99815 |
| LTR16D | 4 | 2 | 2 | 0.001805054 | 0.14271 | 0.99815 |
| MLT1D-int | 4 | 2 | 2 | 0.00312989 | 0.14259 | 0.99815 |
| Kanga11a | 4 | 2 | 2 | 0.002259887 | 0.14267 | 0.99815 |
| LTR54B | 4 | 2 | 2 | 0.002364066 | 0.14266 | 0.99815 |
| FordPrefec | 4 | 2 | 2 | 0.003322259 | 0.14258 | 0.99815 |
| TAR1 | 3 | 2 | 1.5 | 0.01242236 | 0.32332 | 1 |
| L1MC | 10 | 27 | 0.37037037 | 0.002256393 | 0.99994 | 1 |
| MER5B | 12 | 29 | 0.413793103 | 0.001163397 | 0.99988 | 1 |
| L2a | 243 | 411 | 0.591240876 | 0.00240261 | 1 | 1 |
| MIR | 264 | 282 | 0.936170213 | 0.001605868 | 0.8654 | 1 |
| L2c | 269 | 278 | 0.967625899 | 0.001976495 | 0.71349 | 1 |
| AluSp | 18 | 174 | 0.103448276 | 0.003464686 | 1 | 1 |
| MER33 | 4 | 20 | 0.2 | 0.002085941 | 1 | 1 |
| MIRb | 374 | 355 | 1.053521127 | 0.001576692 | 0.16279 | 1 |
| MER53 | 1 | 8 | 0.125 | 0.001364024 | 0.99967 | 1 |
| MLT1A | 15 | 21 | 0.714285714 | 0.002315325 | 0.92865 | 1 |
| AluJo | 41 | 194 | 0.211340206 | 0.002683153 | 1 | 1 |
| AluYc | 1 | 13 | 0.076923077 | 0.001526001 | 1 | 1 |
| L1PA6 | 2 | 84 | 0.023809524 | 0.014053873 | 1 | 1 |
| L1P1 | 0 | 19 | 0 | 0.006054812 | 1 | 1 |
| AluJr | 49 | 215 | 0.227906977 | 0.002784794 | 1 | 1 |
| Charlie5 | 3 | 6 | 0.5 | 0.002347418 | 0.93824 | 1 |
| MLT1E1A-i | 0 | 1 | 0 | 0.011363636 | 1 | 1 |
| MLT1E1A | 5 | 9 | 0.555555556 | 0.002676978 | 0.94526 | 1 |
| AluSx | 37 | 423 | 0.087470449 | 0.002928612 | 1 | 1 |
| AluSz6 | 27 | 133 | 0.203007519 | 0.002906913 | 1 | 1 |
| LTR16C | 10 | 14 | 0.714285714 | 0.002111295 | 0.89085 | 1 |
| ERVL-E-int | 34 | 30 | 1.133333333 | 0.003360968 | 0.25525 | 1 |
| L1MA8 | 9 | 44 | 0.204545455 | 0.003926818 | 1 | 1 |
| L1M5 | 62 | 154 | 0.402597403 | 0.00239227 | 1 | 1 |
| L1MA9 | 21 | 58 | 0.362068966 | 0.00346248 | 1 | 1 |
| LTR12F | 2 | 2 | 1 | 0.003144654 | 0.59442 | 1 |
| MER45A | 2 | 5 | 0.4 | 0.001443001 | 0.95967 | 1 |
| MER58A | 14 | 21 | 0.666666667 | 0.001555786 | 0.95676 | 1 |
| L1PA14 | 5 | 19 | 0.263157895 | 0.006188925 | 0.99996 | 1 |
| AluY | 21 | 372 | 0.056451613 | 0.003084142 | 1 | 1 |
| L2 | 151 | 147 | 1.027210884 | 0.002588712 | 0.38149 | 1 |
| FLAM_A | 2 | 22 | 0.090909091 | 0.001357019 | 1 | 1 |
| MER47A | 1 | 7 | 0.142857143 | 0.002317881 | 0.9991 | 1 |
| AluSc | 9 | 108 | 0.083333333 | 0.003128893 | 1 | 1 |
| L1PA4 | 0 | 131 | 0 | 0.010989011 | 1 | 1 |
| L1MC4a | 69 | 84 | 0.821428571 | 0.003027136 | 0.95831 | 1 |
| L1PA7 | 6 | 138 | 0.043478261 | 0.010580388 | 1 | 1 |
| L1PA16 | 8 | 71 | 0.112676056 | 0.005053021 | 1 | 1 |
| L1PA2 | 1 | 65 | 0.015384615 | 0.013219443 | 1 | 1 |
| AluSz | 67 | 320 | 0.209375 | 0.003253917 | 1 | 1 |

| | | | | | | |
|---|---|---|---|---|---|---|
| L1M2 | 9 | 38 | 0.236842105 | 0.004158914 | 1 | 1 |
| FLAM_C | 6 | 36 | 0.166666667 | 0.001581931 | 1 | 1 |
| L1PREC2 | 7 | 43 | 0.162790698 | 0.00546726 | 1 | 1 |
| L1PB | 1 | 6 | 0.166666667 | 0.003376477 | 0.99755 | 1 |
| Tigger5 | 0 | 3 | 0 | 0.001139818 | 1 | 1 |
| AluSc5 | 1 | 20 | 0.05 | 0.002911208 | 1 | 1 |
| L1M4c | 9 | 29 | 0.310344828 | 0.004685733 | 1 | 1 |
| L1MC5 | 23 | 55 | 0.418181818 | 0.002638902 | 1 | 1 |
| MER4D1 | 0 | 7 | 0 | 0.004602235 | 1 | 1 |
| MER4A | 1 | 5 | 0.2 | 0.004420866 | 0.99334 | 1 |
| L5 | 1 | 1 | 1 | 0.001355014 | 0.63237 | 1 |
| AluSg4 | 3 | 24 | 0.125 | 0.003201708 | 1 | 1 |
| AluSx3 | 12 | 95 | 0.126315789 | 0.003207942 | 1 | 1 |
| LTR33 | 17 | 20 | 0.85 | 0.002159827 | 0.7792 | 1 |
| MSTB1 | 0 | 14 | 0 | 0.002759708 | 1 | 1 |
| AluSq | 11 | 75 | 0.146666667 | 0.003424658 | 1 | 1 |
| L1ME4a | 38 | 93 | 0.408602151 | 0.002151284 | 1 | 1 |
| MLT1C | 36 | 54 | 0.666666667 | 0.002723971 | 0.99616 | 1 |
| AluJb | 110 | 400 | 0.275 | 0.002759667 | 1 | 1 |
| FAM | 4 | 9 | 0.444444444 | 0.001862583 | 0.97886 | 1 |
| AluSq2 | 14 | 183 | 0.076502732 | 0.003307607 | 1 | 1 |
| L1ME3E | 4 | 16 | 0.25 | 0.003146509 | 0.99991 | 1 |
| LTR33B | 3 | 3 | 1 | 0.001889169 | 0.57702 | 1 |
| MLT1D | 13 | 57 | 0.228070175 | 0.00274818 | 1 | 1 |
| L1MA5 | 4 | 21 | 0.19047619 | 0.004680187 | 1 | 1 |
| AluSg | 10 | 141 | 0.070921986 | 0.003388039 | 1 | 1 |
| MER1A | 2 | 13 | 0.153846154 | 0.004014824 | 0.99997 | 1 |
| LTR41 | 2 | 5 | 0.4 | 0.002880184 | 0.95977 | 1 |
| LTR37A | 1 | 4 | 0.25 | 0.002033554 | 0.98176 | 1 |
| AluSg7 | 4 | 27 | 0.148148148 | 0.003207413 | 1 | 1 |
| MER11B | 0 | 4 | 0 | 0.00729927 | 1 | 1 |
| AluJr4 | 12 | 47 | 0.255319149 | 0.002657619 | 1 | 1 |
| U6 | 1 | 2 | 0.5 | 0.001168907 | 0.86482 | 1 |
| MER8 | 0 | 3 | 0 | 0.001540832 | 1 | 1 |
| AluSx1 | 29 | 373 | 0.077747989 | 0.003365484 | 1 | 1 |
| L1MC4 | 50 | 92 | 0.543478261 | 0.003106639 | 1 | 1 |
| MSTD | 22 | 19 | 1.157894737 | 0.0024788 | 0.27431 | 1 |
| Charlie4z | 7 | 8 | 0.875 | 0.001344312 | 0.68679 | 1 |
| MER31-int | 3 | 8 | 0.375 | 0.003889159 | 0.98637 | 1 |
| MER34A | 1 | 3 | 0.333333333 | 0.002031144 | 0.95036 | 1 |
| L1MC3 | 20 | 51 | 0.392156863 | 0.003809382 | 1 | 1 |
| MER4A1 | 0 | 8 | 0 | 0.003577818 | 1 | 1 |
| MER4E | 0 | 3 | 0 | 0.004070556 | 1 | 1 |
| L1M2c | 1 | 3 | 0.333333333 | 0.006944444 | 0.95073 | 1 |
| THE1D | 33 | 29 | 1.137931034 | 0.002293941 | 0.25188 | 1 |
| LTR64 | 1 | 2 | 0.5 | 0.005376344 | 0.86539 | 1 |
| MLT1H1-in | 2 | 1 | 2 | 0.002325581 | 0.26424 | 1 |

| | | | | | | |
|---|---|---|---|---|---|---|
| THE1B-int | 3 | 32 | 0.09375 | 0.007565012 | 1 | 1 |
| L1M7 | 12 | 12 | 1 | 0.002726034 | 0.53856 | 1 |
| L1MEc | 19 | 71 | 0.267605634 | 0.0037776 | 1 | 1 |
| L1MA4 | 6 | 46 | 0.130434783 | 0.004575749 | 1 | 1 |
| L1M4 | 35 | 60 | 0.583333333 | 0.003277793 | 0.99982 | 1 |
| L1MA7 | 7 | 30 | 0.233333333 | 0.003461006 | 1 | 1 |
| ERVL-int | 2 | 3 | 0.666666667 | 0.003797468 | 0.80142 | 1 |
| L1PA5 | 3 | 116 | 0.025862069 | 0.010231081 | 1 | 1 |
| L1MEd | 16 | 28 | 0.571428571 | 0.002606834 | 0.99462 | 1 |
| L1PA10 | 5 | 48 | 0.104166667 | 0.006688963 | 1 | 1 |
| L1PA8 | 2 | 47 | 0.042553191 | 0.00578248 | 1 | 1 |
| L1PA8A | 2 | 19 | 0.105263158 | 0.007630522 | 1 | 1 |
| L1P4 | 2 | 12 | 0.166666667 | 0.003214573 | 0.99992 | 1 |
| L1PA15-16 | 1 | 9 | 0.111111111 | 0.006382979 | 0.99988 | 1 |
| LTR22C | 0 | 1 | 0 | 0.002538071 | 1 | 1 |
| L1ME3D | 4 | 14 | 0.285714286 | 0.003072197 | 0.99953 | 1 |
| L1MA2 | 8 | 49 | 0.163265306 | 0.006521161 | 1 | 1 |
| L1ME3B | 7 | 27 | 0.259259259 | 0.003192999 | 1 | 1 |
| LTR39 | 1 | 4 | 0.25 | 0.003154574 | 0.9818 | 1 |
| MER103C | 14 | 12 | 1.166666667 | 0.001341682 | 0.31839 | 1 |
| MSTA | 20 | 48 | 0.416666667 | 0.002426448 | 1 | 1 |
| LTR37B | 0 | 4 | 0 | 0.001966568 | 1 | 1 |
| MLT1E3 | 3 | 6 | 0.5 | 0.002944063 | 0.93829 | 1 |
| MER4E1 | 0 | 4 | 0 | 0.003813155 | 1 | 1 |
| MER5A | 28 | 45 | 0.622222222 | 0.001291359 | 0.99735 | 1 |
| LTR9B | 0 | 3 | 0 | 0.003957784 | 1 | 1 |
| LTR83 | 1 | 2 | 0.5 | 0.002735978 | 0.86504 | 1 |
| L1ME1 | 89 | 123 | 0.723577236 | 0.003875969 | 0.99946 | 1 |
| MER39B | 3 | 4 | 0.75 | 0.003392706 | 0.76239 | 1 |
| Tigger4b | 0 | 6 | 0 | 0.002380008 | 1 | 1 |
| Ricksha_0 | 2 | 2 | 1 | 0.004950495 | 0.59467 | 1 |
| Charlie9 | 4 | 3 | 1.333333333 | 0.002215657 | 0.35277 | 1 |
| MER31B | 4 | 4 | 1 | 0.00268998 | 0.56679 | 1 |
| MER124 | 1 | 1 | 1 | 0.001340483 | 0.63237 | 1 |
| MER3 | 8 | 16 | 0.5 | 0.001461988 | 0.99004 | 1 |
| MER5A1 | 15 | 20 | 0.75 | 0.001255256 | 0.89528 | 1 |
| L1ME3 | 7 | 29 | 0.24137931 | 0.003083794 | 1 | 1 |
| L1MEg | 14 | 54 | 0.259259259 | 0.002978982 | 1 | 1 |
| BLACKJACK | 2 | 5 | 0.4 | 0.002345216 | 0.95973 | 1 |
| L1M3 | 3 | 19 | 0.157894737 | 0.002773723 | 1 | 1 |
| LTR7 | 0 | 8 | 0 | 0.003412969 | 1 | 1 |
| MER67C | 0 | 5 | 0 | 0.003045067 | 1 | 1 |
| LTR79 | 5 | 6 | 0.833333333 | 0.00148002 | 0.71514 | 1 |
| Charlie1b | 10 | 7 | 1.428571429 | 0.002298851 | 0.16927 | 1 |
| MER113A | 2 | 3 | 0.666666667 | 0.001878522 | 0.80113 | 1 |
| MER102c | 3 | 5 | 0.6 | 0.001436369 | 0.87553 | 1 |
| MER41D | 0 | 3 | 0 | 0.006315789 | 1 | 1 |

| | | | | | | |
|---|---|---|---|---|---|---|
| HAL1 | 70 | 82 | 0.853658537 | 0.003002783 | 0.9194 | 1 |
| MLT1A1 | 7 | 16 | 0.4375 | 0.002364765 | 0.99602 | 1 |
| LTR78 | 13 | 11 | 1.181818182 | 0.002282631 | 0.31118 | 1 |
| L1M4b | 8 | 24 | 0.333333333 | 0.003673095 | 0.99995 | 1 |
| MLT1B | 15 | 43 | 0.348837209 | 0.002388358 | 1 | 1 |
| L1MC1 | 26 | 58 | 0.448275862 | 0.004415011 | 1 | 1 |
| L1PA3 | 0 | 130 | 0 | 0.012212306 | 1 | 1 |
| Charlie19a | 2 | 3 | 0.666666667 | 0.001866833 | 0.80113 | 1 |
| LTR2C | 0 | 2 | 0 | 0.006779661 | 1 | 1 |
| FRAM | 4 | 15 | 0.266666667 | 0.001733102 | 0.99979 | 1 |
| HAL1-3A_M | 0 | 2 | 0 | 0.001040583 | 1 | 1 |
| BSR/Beta | 9 | 12 | 0.75 | 0.006048387 | 0.84577 | 1 |
| AluSc8 | 5 | 69 | 0.072463768 | 0.003144511 | 1 | 1 |
| LOR1-int | 6 | 6 | 1 | 0.004325883 | 0.55467 | 1 |
| LTR57-int | 0 | 2 | 0 | 0.004576659 | 1 | 1 |
| LTR49-int | 3 | 8 | 0.375 | 0.003673095 | 0.98636 | 1 |
| AluYk4 | 0 | 6 | 0 | 0.003210273 | 1 | 1 |
| HERVK9-int | 1 | 9 | 0.111111111 | 0.013452915 | 0.99988 | 1 |
| MER9a2 | 0 | 1 | 0 | 0.003236246 | 1 | 1 |
| MER102a | 2 | 5 | 0.4 | 0.001753771 | 0.95969 | 1 |
| MER67A | 1 | 1 | 1 | 0.002087683 | 0.6325 | 1 |
| MER2 | 1 | 21 | 0.047619048 | 0.002265617 | 1 | 1 |
| HERV16-int | 5 | 6 | 0.833333333 | 0.003194888 | 0.71537 | 1 |
| MER63B | 6 | 4 | 1.5 | 0.001667361 | 0.21474 | 1 |
| MER4A1_ | 0 | 2 | 0 | 0.004975124 | 1 | 1 |
| AluYa5 | 0 | 10 | 0 | 0.002552323 | 1 | 1 |
| L1MEe | 6 | 34 | 0.176470588 | 0.002852828 | 1 | 1 |
| LTR12C | 3 | 36 | 0.083333333 | 0.013138686 | 1 | 1 |
| LTR13A | 2 | 2 | 1 | 0.010638298 | 0.59544 | 1 |
| L1MD | 7 | 25 | 0.28 | 0.002826775 | 0.99999 | 1 |
| L1MEf | 20 | 54 | 0.37037037 | 0.003783102 | 1 | 1 |
| MER49 | 3 | 6 | 0.5 | 0.004382761 | 0.93842 | 1 |
| L1ME2 | 26 | 46 | 0.565217391 | 0.003535198 | 0.99948 | 1 |
| MER50 | 0 | 9 | 0 | 0.003499222 | 1 | 1 |
| L1PB1 | 10 | 92 | 0.108695652 | 0.006816835 | 1 | 1 |
| MER30 | 0 | 8 | 0 | 0.001927246 | 1 | 1 |
| Arthur1B | 4 | 5 | 0.8 | 0.002158895 | 0.73528 | 1 |
| LTR46 | 0 | 1 | 0 | 0.003861004 | 1 | 1 |
| MER9a1 | 0 | 2 | 0 | 0.005830904 | 1 | 1 |
| MER54A | 5 | 3 | 1.666666667 | 0.004424779 | 0.18436 | 1 |
| MER4B | 0 | 7 | 0 | 0.00404157 | 1 | 1 |
| MER4B-int | 5 | 6 | 0.833333333 | 0.004081633 | 0.71549 | 1 |
| AluYf4 | 0 | 4 | 0 | 0.002900653 | 1 | 1 |
| LTR16B1 | 4 | 3 | 1.333333333 | 0.002516779 | 0.35277 | 1 |
| AluSx4 | 1 | 18 | 0.055555556 | 0.00311311 | 1 | 1 |
| MER63D | 3 | 3 | 1 | 0.001737116 | 0.577 | 1 |
| MER66C | 1 | 2 | 0.5 | 0.003395586 | 0.86512 | 1 |

| | | | | | | |
|---|---|---|---|---|---|---|
| MER66-int | 1 | 5 | 0.2 | 0.00669344 | 0.99337 | 1 |
| AluYb8 | 0 | 7 | 0 | 0.002452698 | 1 | 1 |
| AluYk11 | 0 | 1 | 0 | 0.000957854 | 1 | 1 |
| 7SLRNA | 1 | 3 | 0.333333333 | 0.002025658 | 0.95036 | 1 |
| MER74B | 3 | 3 | 1 | 0.003215434 | 0.57717 | 1 |
| LTR5_Hs | 0 | 7 | 0 | 0.010852713 | 1 | 1 |
| L1M3e | 0 | 3 | 0 | 0.003703704 | 1 | 1 |
| LTR2B | 5 | 3 | 1.666666667 | 0.009202454 | 0.18396 | 1 |
| Harlequin-i | 5 | 8 | 0.625 | 0.014678899 | 0.90205 | 1 |
| LTR75 | 0 | 1 | 0 | 0.003436426 | 1 | 1 |
| LTR10A | 0 | 2 | 0 | 0.006389776 | 1 | 1 |
| HERVIP10F | 3 | 6 | 0.5 | 0.012684989 | 0.93916 | 1 |
| HERVI-int | 2 | 3 | 0.666666667 | 0.020833333 | 0.80399 | 1 |
| HUERS-P1- | 0 | 4 | 0 | 0.007920792 | 1 | 1 |
| LTR7B | 0 | 4 | 0 | 0.004716981 | 1 | 1 |
| Charlie22a | 1 | 2 | 0.5 | 0.001796945 | 0.86491 | 1 |
| MER65A | 5 | 4 | 1.25 | 0.002830856 | 0.37116 | 1 |
| MER65-int | 2 | 5 | 0.4 | 0.005213764 | 0.95992 | 1 |
| MER52A | 4 | 15 | 0.266666667 | 0.008210181 | 0.9998 | 1 |
| MER52C | 1 | 3 | 0.333333333 | 0.008 | 0.95081 | 1 |
| MER83 | 0 | 2 | 0 | 0.004301075 | 1 | 1 |
| MER113 | 9 | 9 | 1 | 0.002356021 | 0.5445 | 1 |
| LTR26B | 1 | 1 | 1 | 0.002178649 | 0.63252 | 1 |
| LTR42 | 2 | 1 | 2 | 0.003236246 | 0.26424 | 1 |
| MER4D | 0 | 5 | 0 | 0.003753754 | 1 | 1 |
| Tigger1a_A | 0 | 1 | 0 | 0.003289474 | 1 | 1 |
| MER65B | 1 | 1 | 1 | 0.004854369 | 0.63302 | 1 |
| L1ME2z | 6 | 22 | 0.272727273 | 0.002868318 | 0.99999 | 1 |
| L1P5 | 0 | 1 | 0 | 0.001436782 | 1 | 1 |
| ORSL-2b | 0 | 2 | 0 | 0.002797203 | 1 | 1 |
| Tigger1 | 40 | 76 | 0.526315789 | 0.006276323 | 1 | 1 |
| L1MCa | 11 | 29 | 0.379310345 | 0.003806274 | 0.99996 | 1 |
| MamRep11 | 2 | 3 | 0.666666667 | 0.001722158 | 0.80111 | 1 |
| Arthur1A | 0 | 2 | 0 | 0.001733102 | 1 | 1 |
| L1MEb | 5 | 5 | 1 | 0.002748763 | 0.55975 | 1 |
| L3b | 10 | 9 | 1.111111111 | 0.001302083 | 0.41259 | 1 |
| AluSq4 | 0 | 4 | 0 | 0.002795248 | 1 | 1 |
| AluYc3 | 0 | 2 | 0 | 0.003508772 | 1 | 1 |
| Charlie15a | 3 | 4 | 0.75 | 0.001170275 | 0.76207 | 1 |
| AluSq10 | 1 | 6 | 0.166666667 | 0.002437043 | 0.99754 | 1 |
| Tigger2 | 5 | 13 | 0.384615385 | 0.004761905 | 0.99632 | 1 |
| MER44A | 7 | 6 | 1.166666667 | 0.002554278 | 0.3937 | 1 |
| X7A_LINE | 0 | 1 | 0 | 0.001078749 | 1 | 1 |
| MER1B | 1 | 15 | 0.066666667 | 0.002792256 | 1 | 1 |
| Charlie1a | 23 | 22 | 1.045454545 | 0.003785923 | 0.44362 | 1 |
| LSU-rRNA_ | 2 | 2 | 1 | 0.004830918 | 0.59465 | 1 |
| LTR16E1 | 6 | 5 | 1.2 | 0.002020202 | 0.38404 | 1 |

| | | | | | | |
|---|---|---|---|---|---|---|
| ERV3-16A3 | 24 | 30 | 0.8 | 0.004638219 | 0.88591 | 1 |
| MLT2D | 2 | 9 | 0.222222222 | 0.00198895 | 0.99877 | 1 |
| Charlie2b | 3 | 10 | 0.3 | 0.002710762 | 0.99726 | 1 |
| MER89 | 1 | 4 | 0.25 | 0.002875629 | 0.98179 | 1 |
| MER91B | 0 | 2 | 0 | 0.001244555 | 1 | 1 |
| MER31A | 0 | 5 | 0 | 0.003700962 | 1 | 1 |
| MLT1A0 | 33 | 47 | 0.70212766 | 0.002276801 | 0.9867 | 1 |
| MSTB | 12 | 22 | 0.545454545 | 0.002569493 | 0.99243 | 1 |
| LTR18B | 0 | 2 | 0 | 0.004987531 | 1 | 1 |
| MER102b | 5 | 8 | 0.625 | 0.001962227 | 0.90059 | 1 |
| MER46C | 4 | 6 | 0.666666667 | 0.002130682 | 0.84908 | 1 |
| Charlie8 | 4 | 6 | 0.666666667 | 0.002046385 | 0.84907 | 1 |
| LTR81B | 0 | 2 | 0 | 0.001725626 | 1 | 1 |
| LTR81 | 2 | 1 | 2 | 0.00308642 | 0.26424 | 1 |
| MER58B | 4 | 14 | 0.285714286 | 0.001995155 | 0.99953 | 1 |
| Kanga1 | 3 | 2 | 1.5 | 0.00244798 | 0.32332 | 1 |
| MLT-int | 1 | 1 | 1 | 0.002832861 | 0.63264 | 1 |
| MLT2C1 | 5 | 5 | 1 | 0.001908397 | 0.55967 | 1 |
| L1ME3F | 7 | 16 | 0.4375 | 0.003234283 | 0.99604 | 1 |
| MER91A | 2 | 5 | 0.4 | 0.001375516 | 0.95966 | 1 |
| HAL1-2a_M | 0 | 2 | 0 | 0.001344086 | 1 | 1 |
| MLT2B1 | 10 | 10 | 1 | 0.002232143 | 0.54221 | 1 |
| MER47B | 1 | 2 | 0.5 | 0.002214839 | 0.86496 | 1 |
| THE1A | 13 | 12 | 1.083333333 | 0.002834869 | 0.42403 | 1 |
| Tigger2a | 1 | 9 | 0.111111111 | 0.002697842 | 0.99988 | 1 |
| MER34C_ | 2 | 2 | 1 | 0.003053435 | 0.59441 | 1 |
| MER58C | 0 | 3 | 0 | 0.001148106 | 1 | 1 |
| MER34 | 0 | 3 | 0 | 0.002259036 | 1 | 1 |
| LTR19A | 2 | 2 | 1 | 0.004464286 | 0.5946 | 1 |
| L1MCc | 3 | 10 | 0.3 | 0.0029274 | 0.99726 | 1 |
| MSTA-int | 4 | 19 | 0.210526316 | 0.006014562 | 0.99999 | 1 |
| HERVL-int | 17 | 24 | 0.708333333 | 0.010544815 | 0.94466 | 1 |
| LTR50 | 4 | 6 | 0.666666667 | 0.002290076 | 0.8491 | 1 |
| LTR27 | 1 | 2 | 0.5 | 0.004854369 | 0.86532 | 1 |
| L4 | 22 | 31 | 0.709677419 | 0.001838671 | 0.96218 | 1 |
| MLT1H-int | 0 | 2 | 0 | 0.002487562 | 1 | 1 |
| MER75 | 0 | 1 | 0 | 0.002105263 | 1 | 1 |
| MLT2B4 | 12 | 10 | 1.2 | 0.002180074 | 0.3031 | 1 |
| MER82 | 13 | 10 | 1.3 | 0.003050641 | 0.20815 | 1 |
| L1ME3A | 33 | 49 | 0.673469388 | 0.003044613 | 0.99362 | 1 |
| LTR40b | 2 | 3 | 0.666666667 | 0.00261324 | 0.80124 | 1 |
| MLT1E2 | 10 | 12 | 0.833333333 | 0.003003003 | 0.758 | 1 |
| ERVL-B4-in | 19 | 19 | 1 | 0.004893124 | 0.53074 | 1 |
| Arthur1C | 2 | 1 | 2 | 0.001398601 | 0.26424 | 1 |
| MER45C | 2 | 3 | 0.666666667 | 0.002506266 | 0.80123 | 1 |
| THE1C-int | 2 | 12 | 0.166666667 | 0.00660066 | 0.99992 | 1 |
| L1PA12 | 3 | 11 | 0.272727273 | 0.006176305 | 0.99882 | 1 |

| | | | | | | |
|---|---|---|---|---|---|---|
| AluYg6 | 0 | 2 | 0 | 0.00250941 | 1 | 1 |
| MSTB-int | 2 | 6 | 0.333333333 | 0.006841505 | 0.9829 | 1 |
| L1MDa | 8 | 33 | 0.242424242 | 0.004532344 | 1 | 1 |
| L1PA15 | 7 | 52 | 0.134615385 | 0.00615749 | 1 | 1 |
| UCON16 | 0 | 1 | 0 | 0.027027027 | 1 | 1 |
| Eulor10 | 0 | 1 | 0 | 0.029411765 | 1 | 1 |
| L1MA10 | 6 | 12 | 0.5 | 0.002350637 | 0.97976 | 1 |
| L1MC2 | 15 | 28 | 0.535714286 | 0.004347826 | 0.99731 | 1 |
| L1ME3C | 10 | 36 | 0.277777778 | 0.002856463 | 1 | 1 |
| L1MA5A | 4 | 14 | 0.285714286 | 0.004092371 | 0.99953 | 1 |
| LTR10B1 | 0 | 1 | 0 | 0.004115226 | 1 | 1 |
| LTR21B | 0 | 1 | 0 | 0.016666667 | 1 | 1 |
| LTR62 | 2 | 1 | 2 | 0.003174603 | 0.26424 | 1 |
| L1PA11 | 3 | 29 | 0.103448276 | 0.006931166 | 1 | 1 |
| THE1D-int | 2 | 12 | 0.166666667 | 0.006389776 | 0.99992 | 1 |
| Charlie4a | 4 | 8 | 0.5 | 0.002296871 | 0.95778 | 1 |
| MER34C2 | 0 | 2 | 0 | 0.003558719 | 1 | 1 |
| MER34B-in | 1 | 3 | 0.333333333 | 0.00443787 | 0.95054 | 1 |
| MER39 | 1 | 9 | 0.111111111 | 0.002697033 | 0.99988 | 1 |
| MADE2 | 1 | 2 | 0.5 | 0.000749906 | 0.86477 | 1 |
| MSR1 | 0 | 2 | 0 | 0.006329114 | 1 | 1 |
| Helitron3N | 2 | 2 | 1 | 0.001795332 | 0.59424 | 1 |
| LTR5B | 0 | 4 | 0 | 0.009280742 | 1 | 1 |
| LTR38 | 0 | 1 | 0 | 0.003225806 | 1 | 1 |
| LTR38-int | 0 | 1 | 0 | 0.008130081 | 1 | 1 |
| HERVK13-ii | 0 | 1 | 0 | 0.014705882 | 1 | 1 |
| LTR3B_ | 1 | 1 | 1 | 0.003816794 | 0.63282 | 1 |
| MLT1E1 | 1 | 3 | 0.333333333 | 0.002876318 | 0.95043 | 1 |
| LTR78B | 8 | 6 | 1.333333333 | 0.001828711 | 0.25589 | 1 |
| L1MB1 | 11 | 22 | 0.5 | 0.003523382 | 0.9965 | 1 |
| Tigger4a | 1 | 6 | 0.166666667 | 0.001718705 | 0.99753 | 1 |
| L1MD2 | 29 | 44 | 0.659090909 | 0.00396004 | 0.99337 | 1 |
| MER115 | 3 | 7 | 0.428571429 | 0.002656546 | 0.97051 | 1 |
| MER77 | 6 | 4 | 1.5 | 0.003088803 | 0.21463 | 1 |
| LTR3A | 0 | 1 | 0 | 0.006134969 | 1 | 1 |
| LTR85c | 3 | 2 | 1.5 | 0.001686341 | 0.32332 | 1 |
| Kanga1d | 2 | 1 | 2 | 0.001501502 | 0.26424 | 1 |
| MLT2B2 | 1 | 5 | 0.2 | 0.002263468 | 0.9933 | 1 |
| MER85 | 0 | 1 | 0 | 0.001104972 | 1 | 1 |
| MER92B | 2 | 2 | 1 | 0.002239642 | 0.5943 | 1 |
| LTR10E | 0 | 1 | 0 | 0.003436426 | 1 | 1 |
| MER63C | 5 | 3 | 1.666666667 | 0.003367003 | 0.18445 | 1 |
| LTR16E2 | 4 | 5 | 0.8 | 0.002158895 | 0.73528 | 1 |
| L1PA13 | 7 | 51 | 0.137254902 | 0.00560871 | 1 | 1 |
| polypurine | 0 | 1 | 0 | 0.00073692 | 1 | 1 |
| L1ME5 | 8 | 9 | 0.888888889 | 0.002610209 | 0.67641 | 1 |
| MamRep4( | 1 | 3 | 0.333333333 | 0.002358491 | 0.95039 | 1 |

| | | | | | | |
|---|---|---|---|---|---|---|
| L1P2 | 0 | 16 | 0 | 0.010403121 | 1 | 1 |
| polypyrimi | 0 | 1 | 0 | 0.000766284 | 1 | 1 |
| LTR85a | 2 | 3 | 0.666666667 | 0.001576458 | 0.80109 | 1 |
| LTR24C | 1 | 1 | 1 | 0.002617801 | 0.6326 | 1 |
| L1MA1 | 2 | 25 | 0.08 | 0.005787037 | 1 | 1 |
| MSTC | 5 | 8 | 0.625 | 0.002524456 | 0.90066 | 1 |
| MER21A | 3 | 8 | 0.375 | 0.004164498 | 0.98638 | 1 |
| MSTB2 | 4 | 4 | 1 | 0.002495321 | 0.56677 | 1 |
| MLT2A2 | 8 | 11 | 0.727272727 | 0.00282196 | 0.85717 | 1 |
| MER48 | 0 | 1 | 0 | 0.005181347 | 1 | 1 |
| HERVH-int | 1 | 44 | 0.022727273 | 0.007446268 | 1 | 1 |
| LTR7A | 0 | 1 | 0 | 0.066666667 | 1 | 1 |
| L1PA17 | 5 | 23 | 0.217391304 | 0.004726675 | 1 | 1 |
| L1PB4 | 5 | 28 | 0.178571429 | 0.003727866 | 1 | 1 |
| LTR53 | 3 | 2 | 1.5 | 0.002998501 | 0.32332 | 1 |
| MER61B | 0 | 1 | 0 | 0.002320186 | 1 | 1 |
| MLT1A0-in | 0 | 4 | 0 | 0.004319654 | 1 | 1 |
| LOR1a | 1 | 2 | 0.5 | 0.002478315 | 0.865 | 1 |
| MER57B1 | 2 | 2 | 1 | 0.001974334 | 0.59426 | 1 |
| LTR16A2 | 2 | 4 | 0.5 | 0.002171553 | 0.90866 | 1 |
| MER90 | 3 | 3 | 1 | 0.004273504 | 0.57729 | 1 |
| LTR1 | 5 | 7 | 0.714285714 | 0.004834254 | 0.82767 | 1 |
| MER11A | 1 | 8 | 0.125 | 0.008298755 | 0.99968 | 1 |
| LTR81AB | 0 | 1 | 0 | 0.003663004 | 1 | 1 |
| LTR1B | 0 | 5 | 0 | 0.004258944 | 1 | 1 |
| MER61D | 0 | 1 | 0 | 0.003676471 | 1 | 1 |
| MER61A | 0 | 2 | 0 | 0.002463054 | 1 | 1 |
| Arthur1 | 7 | 6 | 1.166666667 | 0.002574003 | 0.3937 | 1 |
| X7B_LINE | 1 | 1 | 1 | 0.001057082 | 0.63232 | 1 |
| L1MA6 | 6 | 23 | 0.260869565 | 0.004206291 | 0.99999 | 1 |
| MamRep43 | 0 | 3 | 0 | 0.0013947 | 1 | 1 |
| L1PB2 | 2 | 16 | 0.125 | 0.005582694 | 1 | 1 |
| L1P3 | 3 | 19 | 0.157894737 | 0.005565319 | 1 | 1 |
| MER45R | 1 | 1 | 1 | 0.001607717 | 0.63242 | 1 |
| Tigger11a | 2 | 1 | 2 | 0.002314815 | 0.26424 | 1 |
| MER110 | 0 | 1 | 0 | 0.00128041 | 1 | 1 |
| LTR66 | 0 | 1 | 0 | 0.00265252 | 1 | 1 |
| MER54B | 1 | 1 | 1 | 0.002304147 | 0.63254 | 1 |
| MER70A | 0 | 1 | 0 | 0.001869159 | 1 | 1 |
| MLT1E | 0 | 2 | 0 | 0.002202643 | 1 | 1 |
| L1MD1 | 24 | 35 | 0.685714286 | 0.005036696 | 0.97947 | 1 |
| LTR90B | 1 | 1 | 1 | 0.001858736 | 0.63246 | 1 |
| MER20B | 11 | 10 | 1.1 | 0.002407318 | 0.41696 | 1 |
| LTR24B | 2 | 2 | 1 | 0.004273504 | 0.59457 | 1 |
| MER57A1 | 4 | 3 | 1.333333333 | 0.002827521 | 0.35277 | 1 |
| LTR10C | 0 | 3 | 0 | 0.005791506 | 1 | 1 |
| MLT1J-int | 4 | 3 | 1.333333333 | 0.002079002 | 0.35277 | 1 |

| | | | | | | |
|---|---|---|---|---|---|---|
| MADE1 | 1 | 4 | 0.25 | 0.000511313 | 0.9817 | 1 |
| MLT2A1 | 1 | 8 | 0.125 | 0.002116402 | 0.99967 | 1 |
| LTR48B | 3 | 3 | 1 | 0.002938296 | 0.57714 | 1 |
| MER34-int | 1 | 2 | 0.5 | 0.005050505 | 0.86535 | 1 |
| MER70B | 0 | 1 | 0 | 0.002087683 | 1 | 1 |
| LFSINE_Ve | 3 | 2 | 1.5 | 0.002079002 | 0.32332 | 1 |
| Tigger9b | 1 | 2 | 0.5 | 0.002617801 | 0.86502 | 1 |
| Kanga2_a | 4 | 4 | 1 | 0.002602472 | 0.56678 | 1 |
| MamGypLT | 3 | 2 | 1.5 | 0.001833181 | 0.32332 | 1 |
| MamRep38 | 2 | 3 | 0.666666667 | 0.001411101 | 0.80106 | 1 |
| L1PB3 | 2 | 18 | 0.111111111 | 0.004971002 | 1 | 1 |
| LTR23 | 0 | 2 | 0 | 0.002355713 | 1 | 1 |
| LTR23-int | 0 | 1 | 0 | 0.003584229 | 1 | 1 |
| MER34B | 1 | 2 | 0.5 | 0.002989537 | 0.86507 | 1 |
| MER34C | 0 | 1 | 0 | 0.005434783 | 1 | 1 |
| MER75A | 0 | 1 | 0 | 0.010869565 | 1 | 1 |
| Tigger2b_P | 7 | 5 | 1.4 | 0.002436647 | 0.23764 | 1 |
| L1M6 | 7 | 16 | 0.4375 | 0.002321869 | 0.99602 | 1 |
| MER34A1 | 0 | 2 | 0 | 0.002604167 | 1 | 1 |
| MamRep18 | 2 | 2 | 1 | 0.001349528 | 0.59418 | 1 |
| LTR16B2 | 2 | 4 | 0.5 | 0.002469136 | 0.90869 | 1 |
| Zaphod3 | 0 | 2 | 0 | 0.001612903 | 1 | 1 |
| MER58D | 0 | 1 | 0 | 0.000866551 | 1 | 1 |
| LTR33A | 6 | 4 | 1.5 | 0.001967536 | 0.21472 | 1 |
| MER41A | 10 | 10 | 1 | 0.003921569 | 0.54232 | 1 |
| MER5C | 2 | 3 | 0.666666667 | 0.002232143 | 0.80119 | 1 |
| MLT1C-int | 1 | 2 | 0.5 | 0.004338395 | 0.86525 | 1 |
| LTR16D2 | 0 | 1 | 0 | 0.003558719 | 1 | 1 |
| ORSL | 0 | 2 | 0 | 0.001647446 | 1 | 1 |
| HAL1b | 5 | 11 | 0.454545455 | 0.00227932 | 0.98498 | 1 |
| MLT1G1-in | 1 | 1 | 1 | 0.003257329 | 0.63272 | 1 |
| LTR1C | 1 | 1 | 1 | 0.004032258 | 0.63286 | 1 |
| THE1A-int | 0 | 13 | 0 | 0.008879781 | 1 | 1 |
| LTR41B | 1 | 3 | 0.333333333 | 0.003021148 | 0.95044 | 1 |
| MLT1B-int | 0 | 2 | 0 | 0.003597122 | 1 | 1 |
| MER105 | 1 | 1 | 1 | 0.001385042 | 0.63238 | 1 |
| LTR19B | 0 | 1 | 0 | 0.003558719 | 1 | 1 |
| HERVIP10F | 4 | 5 | 0.8 | 0.007342144 | 0.73601 | 1 |
| Charlie16a | 0 | 3 | 0 | 0.001515152 | 1 | 1 |
| FordPrefec | 0 | 1 | 0 | 0.003546099 | 1 | 1 |
| MER11C | 0 | 7 | 0 | 0.008083141 | 1 | 1 |
| LTR71B | 2 | 1 | 2 | 0.004201681 | 0.26424 | 1 |
| LTR16D1 | 0 | 1 | 0 | 0.004878049 | 1 | 1 |
| L1MA3 | 6 | 60 | 0.1 | 0.006643047 | 1 | 1 |
| Charlie18a | 0 | 4 | 0 | 0.001610954 | 1 | 1 |
| MER51E | 0 | 1 | 0 | 0.004016064 | 1 | 1 |
| LTR86A1 | 2 | 1 | 2 | 0.001904762 | 0.26424 | 1 |

| | | | | | | |
|---|---|---|---|---|---|---|
| U2 | 1 | 1 | 1 | 0.000918274 | 0.63229 | 1 |
| LTR75_1 | 0 | 1 | 0 | 0.0078125 | 1 | 1 |
| MER72 | 2 | 3 | 0.666666667 | 0.003836317 | 0.80143 | 1 |
| SVA_D | 0 | 21 | 0 | 0.014716188 | 1 | 1 |
| AluYa8 | 0 | 1 | 0 | 0.00295858 | 1 | 1 |
| MER52D | 3 | 5 | 0.6 | 0.007656968 | 0.87632 | 1 |
| LTR12E | 3 | 2 | 1.5 | 0.017391304 | 0.32332 | 1 |
| HERVFH21- | 0 | 1 | 0 | 0.008474576 | 1 | 1 |
| LTR14B | 0 | 2 | 0 | 0.005509642 | 1 | 1 |
| LTR12 | 0 | 4 | 0 | 0.005167959 | 1 | 1 |
| MER101 | 1 | 1 | 1 | 0.002724796 | 0.63262 | 1 |
| X7C_LINE | 0 | 1 | 0 | 0.001607717 | 1 | 1 |
| LTR36 | 0 | 2 | 0 | 0.00408998 | 1 | 1 |
| MER126 | 0 | 1 | 0 | 0.004694836 | 1 | 1 |
| UCON22 | 0 | 1 | 0 | 0.028571429 | 1 | 1 |
| U13_ | 1 | 1 | 1 | 0.003257329 | 0.63272 | 1 |
| Charlie7a | 0 | 2 | 0 | 0.001253133 | 1 | 1 |
| MamRep13 | 3 | 4 | 0.75 | 0.001697793 | 0.76215 | 1 |
| Charlie17a | 2 | 2 | 1 | 0.001365188 | 0.59418 | 1 |
| MER94B | 1 | 1 | 1 | 0.001607717 | 0.63242 | 1 |
| Eulor2A | 0 | 1 | 0 | 0.014925373 | 1 | 1 |
| X9_LINE | 0 | 1 | 0 | 0.011904762 | 1 | 1 |
| UCON20 | 0 | 1 | 0 | 0.022727273 | 1 | 1 |
| LTR86A2 | 0 | 1 | 0 | 0.003194888 | 1 | 1 |
| X5B_LINE | 0 | 1 | 0 | 0.01010101 | 1 | 1 |
| U1 | 0 | 1 | 0 | 0.004878049 | 1 | 1 |
| Mam_R4 | 2 | 1 | 2 | 0.001798561 | 0.26424 | 1 |
| MLT2C2 | 2 | 3 | 0.666666667 | 0.002267574 | 0.80119 | 1 |
| MER96B | 0 | 4 | 0 | 0.001867414 | 1 | 1 |
| MER131 | 0 | 1 | 0 | 0.002352941 | 1 | 1 |
| Charlie24 | 3 | 2 | 1.5 | 0.001506024 | 0.32332 | 1 |
| Tigger16a | 0 | 1 | 0 | 0.001414427 | 1 | 1 |
| MER75B | 0 | 1 | 0 | 0.008849558 | 1 | 1 |
| 5S | 1 | 1 | 1 | 0.000784314 | 0.63226 | 1 |
| MER90a | 4 | 4 | 1 | 0.003207698 | 0.56684 | 1 |
| L1MA4A | 6 | 36 | 0.166666667 | 0.005727924 | 1 | 1 |
| Zaphod | 7 | 6 | 1.166666667 | 0.002831524 | 0.3937 | 1 |
| SVA_E | 0 | 2 | 0 | 0.008474576 | 1 | 1 |
| MER84 | 2 | 1 | 2 | 0.002754821 | 0.26424 | 1 |
| MER84-int | 0 | 1 | 0 | 0.006896552 | 1 | 1 |
| HY1 | 0 | 1 | 0 | 0.002252252 | 1 | 1 |
| LTR9 | 4 | 8 | 0.5 | 0.00397812 | 0.9579 | 1 |
| Charlie26a | 0 | 1 | 0 | 0.001941748 | 1 | 1 |
| Charlie21a | 3 | 2 | 1.5 | 0.00269179 | 0.32332 | 1 |
| MamGypLT | 3 | 2 | 1.5 | 0.001897533 | 0.32332 | 1 |
| LTR29 | 0 | 2 | 0 | 0.002849003 | 1 | 1 |
| Charlie10a | 0 | 1 | 0 | 0.003322259 | 1 | 1 |

| | | | | | | |
|---|---|---|---|---|---|---|
| LTR86C | 1 | 1 | 1 | 0.003508772 | 0.63277 | 1 |
| U5 | 0 | 1 | 0 | 0.004651163 | 1 | 1 |
| L1M | 0 | 1 | 0 | 0.001004016 | 1 | 1 |
| Charlie2a | 4 | 6 | 0.666666667 | 0.002899952 | 0.84918 | 1 |
| SVA_F | 0 | 9 | 0 | 0.008780488 | 1 | 1 |
| MER104 | 2 | 3 | 0.666666667 | 0.001361162 | 0.80106 | 1 |
| MER5C1 | 1 | 1 | 1 | 0.001623377 | 0.63242 | 1 |
| Charlie23a | 1 | 2 | 0.5 | 0.001345895 | 0.86485 | 1 |
| Tigger9a | 0 | 2 | 0 | 0.00194742 | 1 | 1 |
| L1MCb | 3 | 7 | 0.428571429 | 0.003272557 | 0.97055 | 1 |
| MER51B | 3 | 2 | 1.5 | 0.002793296 | 0.32332 | 1 |
| LTR47A | 0 | 2 | 0 | 0.002207506 | 1 | 1 |
| MER9a3 | 0 | 2 | 0 | 0.003448276 | 1 | 1 |
| LTR82A | 1 | 2 | 0.5 | 0.001919386 | 0.86492 | 1 |
| LTR21A | 1 | 1 | 1 | 0.010752688 | 0.63411 | 1 |
| HERVL40-ir | 6 | 4 | 1.5 | 0.003046458 | 0.21463 | 1 |
| LTR77 | 1 | 1 | 1 | 0.006756757 | 0.63337 | 1 |
| LTR61 | 1 | 1 | 1 | 0.005181347 | 0.63308 | 1 |
| MER106B | 0 | 2 | 0 | 0.00241838 | 1 | 1 |
| Charlie7 | 11 | 9 | 1.222222222 | 0.002436383 | 0.29387 | 1 |
| LTR56 | 1 | 3 | 0.333333333 | 0.003061224 | 0.95044 | 1 |
| MER4C | 4 | 4 | 1 | 0.003184713 | 0.56684 | 1 |
| AluYf5 | 0 | 1 | 0 | 0.005555556 | 1 | 1 |
| HERVFH19- | 0 | 1 | 0 | 0.008547009 | 1 | 1 |
| MER110A | 1 | 1 | 1 | 0.001572327 | 0.63241 | 1 |
| AluYd8 | 0 | 1 | 0 | 0.004444444 | 1 | 1 |
| MER11D | 1 | 1 | 1 | 0.003846154 | 0.63283 | 1 |
| MamRep48 | 0 | 1 | 0 | 0.002087683 | 1 | 1 |
| LTR85b | 1 | 2 | 0.5 | 0.001702128 | 0.8649 | 1 |
| LTR60 | 1 | 1 | 1 | 0.003649635 | 0.63279 | 1 |
| LTR2 | 1 | 4 | 0.25 | 0.004509583 | 0.98185 | 1 |
| MER57C1 | 0 | 1 | 0 | 0.005263158 | 1 | 1 |
| MER21-int | 2 | 5 | 0.4 | 0.004578755 | 0.95988 | 1 |
| Tigger4 | 5 | 6 | 0.833333333 | 0.004123711 | 0.7155 | 1 |
| MER47C | 0 | 1 | 0 | 0.001303781 | 1 | 1 |
| ORSL-2a | 0 | 1 | 0 | 0.002747253 | 1 | 1 |
| 7SK | 1 | 1 | 1 | 0.001371742 | 0.63237 | 1 |
| LTR1D | 0 | 5 | 0 | 0.005030181 | 1 | 1 |
| LTR51 | 2 | 1 | 2 | 0.002392344 | 0.26424 | 1 |
| Kanga1c | 0 | 1 | 0 | 0.00120919 | 1 | 1 |
| X7D_LINE | 0 | 1 | 0 | 0.017857143 | 1 | 1 |
| Tigger10 | 2 | 2 | 1 | 0.001953125 | 0.59426 | 1 |
| MER106A | 0 | 1 | 0 | 0.001338688 | 1 | 1 |
| LTR84a | 0 | 1 | 0 | 0.001506024 | 1 | 1 |
| Tigger16b | 0 | 1 | 0 | 0.000951475 | 1 | 1 |
| AmnSINE2 | 0 | 1 | 0 | 0.006711409 | 1 | 1 |
| UCON14 | 0 | 1 | 0 | 0.010204082 | 1 | 1 |

| | | | | | | |
|---|---|---|---|---|---|---|
| UCON29 | 0 | 1 | 0 | 0.004149378 | 1 | 1 |
| Charlie13a | 3 | 2 | 1.5 | 0.003603604 | 0.32332 | 1 |
| Zaphod2 | 0 | 1 | 0 | 0.001360544 | 1 | 1 |
| MER2B | 0 | 3 | 0 | 0.001876173 | 1 | 1 |
| AluYh9 | 0 | 1 | 0 | 0.003676471 | 1 | 1 |
| SVA_C | 0 | 4 | 0 | 0.013333333 | 1 | 1 |
| LTR43B | 0 | 1 | 0 | 0.004310345 | 1 | 1 |
| LTR47B | 1 | 1 | 1 | 0.002421308 | 0.63257 | 1 |
| HY3 | 0 | 1 | 0 | 0.001834862 | 1 | 1 |
| HSMAR1 | 0 | 4 | 0 | 0.005457026 | 1 | 1 |
| L1P4a | 1 | 4 | 0.25 | 0.006791171 | 0.98193 | 1 |
| LTR75B | 1 | 1 | 1 | 0.004854369 | 0.63302 | 1 |
| SVA_B | 0 | 5 | 0 | 0.010504202 | 1 | 1 |
| MER67B | 1 | 2 | 0.5 | 0.004132231 | 0.86522 | 1 |
| U3 | 0 | 1 | 0 | 0.004366812 | 1 | 1 |
| MER97c | 0 | 1 | 0 | 0.004608295 | 1 | 1 |
| LTR65 | 2 | 2 | 1 | 0.004357298 | 0.59459 | 1 |
| HSMAR2 | 2 | 9 | 0.222222222 | 0.005070423 | 0.99879 | 1 |
| Tigger8 | 3 | 2 | 1.5 | 0.002132196 | 0.32332 | 1 |
| MER52-int | 1 | 3 | 0.333333333 | 0.006097561 | 0.95067 | 1 |
| HERV4_I-in | 0 | 2 | 0 | 0.006042296 | 1 | 1 |
| MER68-int | 1 | 2 | 0.5 | 0.005319149 | 0.86539 | 1 |
| MER70C | 0 | 1 | 0 | 0.003636364 | 1 | 1 |
| LTR10F | 3 | 2 | 1.5 | 0.004474273 | 0.32332 | 1 |
| MER6 | 0 | 3 | 0 | 0.003541913 | 1 | 1 |
| U13 | 0 | 1 | 0 | 0.005649718 | 1 | 1 |
| Tigger6b | 0 | 1 | 0 | 0.003389831 | 1 | 1 |
| PRIMA4-int | 3 | 3 | 1 | 0.004310345 | 0.57729 | 1 |
| HERVK-int | 1 | 5 | 0.2 | 0.019607843 | 0.99359 | 1 |
| L1M1 | 18 | 61 | 0.295081967 | 0.006396812 | 1 | 1 |
| MER41G | 1 | 1 | 1 | 0.008849558 | 0.63375 | 1 |
| LTR45B | 2 | 1 | 2 | 0.002252252 | 0.26424 | 1 |
| MER57A-in | 0 | 8 | 0 | 0.004214963 | 1 | 1 |
| LTR45 | 0 | 1 | 0 | 0.005649718 | 1 | 1 |
| MER101-in | 2 | 5 | 0.4 | 0.003958828 | 0.95984 | 1 |
| LTR7Y | 0 | 1 | 0 | 0.004016064 | 1 | 1 |
| L1M3f | 3 | 4 | 0.75 | 0.005698006 | 0.76273 | 1 |
| HERV3-int | 3 | 5 | 0.6 | 0.016339869 | 0.87742 | 1 |
| GSATII | 0 | 1 | 0 | 0.005263158 | 1 | 1 |
| HERVE-int | 1 | 6 | 0.166666667 | 0.0234375 | 0.99769 | 1 |
| GSATX | 0 | 1 | 0 | 0.017857143 | 1 | 1 |
| LTR57 | 0 | 1 | 0 | 0.003745318 | 1 | 1 |
| PRIMAX-int | 1 | 2 | 0.5 | 0.00536193 | 0.86539 | 1 |
| MER110-in | 1 | 1 | 1 | 0.00257732 | 0.6326 | 1 |
| LTR12D | 1 | 4 | 0.25 | 0.008179959 | 0.98198 | 1 |
| LOR1b | 3 | 3 | 1 | 0.002719855 | 0.57712 | 1 |
| MER30B | 0 | 1 | 0 | 0.004405286 | 1 | 1 |

| | | | | | | |
|---|---|---|---|---|---|---|
| LTR10B | 0 | 1 | 0 | 0.012658228 | 1 | 1 |
| LTR14A | 0 | 1 | 0 | 0.007633588 | 1 | 1 |
| LTR22B | 0 | 1 | 0 | 0.004291845 | 1 | 1 |
| MER6C | 0 | 1 | 0 | 0.006289308 | 1 | 1 |
| LTR22A | 1 | 1 | 1 | 0.005291005 | 0.6331 | 1 |
| MST-int | 0 | 1 | 0 | 0.002785515 | 1 | 1 |
| Ricksha | 0 | 1 | 0 | 0.005988024 | 1 | 1 |
| Ricksha_a | 0 | 1 | 0 | 0.015873016 | 1 | 1 |
| LTR80A | 1 | 1 | 1 | 0.002544529 | 0.63259 | 1 |
| MER113B | 0 | 1 | 0 | 0.001766784 | 1 | 1 |
| LTR86B1 | 0 | 1 | 0 | 0.003355705 | 1 | 1 |
| MamGypLT | 1 | 2 | 0.5 | 0.001755926 | 0.8649 | 1 |
| LTR82B | 0 | 2 | 0 | 0.001727116 | 1 | 1 |
| Helitron2N | 0 | 1 | 0 | 0.002293578 | 1 | 1 |
| MLT1F-int | 1 | 2 | 0.5 | 0.00390625 | 0.86519 | 1 |
| LTR55 | 0 | 1 | 0 | 0.002457002 | 1 | 1 |
| L1HS | 0 | 23 | 0 | 0.014896373 | 1 | 1 |
| MER6A | 4 | 3 | 1.333333333 | 0.002941176 | 0.35277 | 1 |
| MLT1I-int | 1 | 2 | 0.5 | 0.0025 | 0.865 | 1 |
| HERVL74-ir | 0 | 3 | 0 | 0.005008347 | 1 | 1 |
| MER135 | 0 | 1 | 0 | 0.001221001 | 1 | 1 |
| LTR32 | 0 | 3 | 0 | 0.002291826 | 1 | 1 |
| X1_LINE | 0 | 1 | 0 | 0.011764706 | 1 | 1 |
| MER127 | 0 | 1 | 0 | 0.004739336 | 1 | 1 |
| LTR10D | 0 | 1 | 0 | 0.005235602 | 1 | 1 |
| PRIMA4_LT | 0 | 1 | 0 | 0.002777778 | 1 | 1 |
| DNA1_Mar | 0 | 1 | 0 | 0.005847953 | 1 | 1 |
| U7 | 0 | 1 | 0 | 0.004048583 | 1 | 1 |
| MLT1J1-int | 1 | 1 | 1 | 0.002016129 | 0.63249 | 1 |
| MER129 | 0 | 1 | 0 | 0.008196721 | 1 | 1 |
| MERX | 0 | 1 | 0 | 0.002298851 | 1 | 1 |
| LTR4 | 0 | 1 | 0 | 0.00990099 | 1 | 1 |
| MER87 | 0 | 1 | 0 | 0.004405286 | 1 | 1 |
| LTR49 | 0 | 4 | 0 | 0.002772003 | 1 | 1 |
| PRIMA41-ir | 2 | 3 | 0.666666667 | 0.006818182 | 0.80187 | 1 |
| LTR39-int | 0 | 1 | 0 | 0.00617284 | 1 | 1 |
| MER57B2 | 4 | 3 | 1.333333333 | 0.002717391 | 0.35277 | 1 |
| LTR76 | 0 | 1 | 0 | 0.005319149 | 1 | 1 |
| MamGypLT | 1 | 1 | 1 | 0.001121076 | 0.63233 | 1 |
| MER50C | 1 | 1 | 1 | 0.008474576 | 0.63368 | 1 |
| PABL_B | 0 | 1 | 0 | 0.00265252 | 1 | 1 |
| LTR15 | 0 | 1 | 0 | 0.003759398 | 1 | 1 |
| LTR38B | 1 | 1 | 1 | 0.004366812 | 0.63293 | 1 |
| HERV-Fc2_ | 0 | 1 | 0 | 0.25 | 1 | 1 |
| MER61-int | 1 | 7 | 0.142857143 | 0.006071119 | 0.99911 | 1 |
| Helitron1N | 0 | 1 | 0 | 0.00308642 | 1 | 1 |
| Helitron1N | 0 | 1 | 0 | 0.002762431 | 1 | 1 |

| | | | | | | |
|---|---|---|---|---|---|---|
| UCON11 | 0 | 1 | 0 | 0.022222222 | 1 | 1 |
| X8_LINE | 1 | 1 | 1 | 0.003322259 | 0.63273 | 1 |
| MER4D0 | 1 | 1 | 1 | 0.002309469 | 0.63255 | 1 |
| HUERS-P3- | 3 | 4 | 0.75 | 0.005555556 | 0.76271 | 1 |
| LTR19C | 0 | 1 | 0 | 0.004424779 | 1 | 1 |
| HERV35I-in | 0 | 3 | 0 | 0.005385996 | 1 | 1 |
| U4 | 0 | 1 | 0 | 0.00621118 | 1 | 1 |
| LTR28 | 1 | 2 | 0.5 | 0.003883495 | 0.86519 | 1 |
| Charlie14a | 1 | 1 | 1 | 0.001908397 | 0.63247 | 1 |
| MamRep18 | 0 | 1 | 0 | 0.001457726 | 1 | 1 |
| LTR14 | 0 | 1 | 0 | 0.010309278 | 1 | 1 |
| HERVK14C- | 0 | 2 | 0 | 0.025316456 | 1 | 1 |
| MER66A | 0 | 1 | 0 | 0.00304878 | 1 | 1 |
| MLT1A1-in | 1 | 2 | 0.5 | 0.004395604 | 0.86526 | 1 |
| LTR46-int | 0 | 1 | 0 | 0.012987013 | 1 | 1 |
| L1M2a | 1 | 3 | 0.333333333 | 0.005617978 | 0.95063 | 1 |
| MER4-int | 4 | 14 | 0.285714286 | 0.0047896 | 0.99954 | 1 |
| L1P4e | 0 | 1 | 0 | 0.004975124 | 1 | 1 |
| UCON21 | 0 | 1 | 0 | 0.035714286 | 1 | 1 |
| MLT1A-int | 0 | 2 | 0 | 0.004444444 | 1 | 1 |
| Eulor1 | 0 | 1 | 0 | 0.011627907 | 1 | 1 |
| LTR7C | 1 | 2 | 0.5 | 0.006289308 | 0.86552 | 1 |
| MER73 | 0 | 1 | 0 | 0.003571429 | 1 | 1 |
| L1PBa | 8 | 17 | 0.470588235 | 0.007773205 | 0.9947 | 1 |
| LTR48 | 2 | 3 | 0.666666667 | 0.003076923 | 0.80131 | 1 |
| MER41-int | 0 | 11 | 0 | 0.006406523 | 1 | 1 |
| MLT1E3-int | 0 | 1 | 0 | 0.012048193 | 1 | 1 |
| PABL_A | 0 | 3 | 0 | 0.005736138 | 1 | 1 |
| MLT1K-int | 0 | 1 | 0 | 0.005025126 | 1 | 1 |
| MER57-int | 2 | 8 | 0.25 | 0.00625 | 0.99704 | 1 |
| X6A_LINE | 0 | 1 | 0 | 0.003649635 | 1 | 1 |
| MER50-int | 1 | 3 | 0.333333333 | 0.005825243 | 0.95065 | 1 |
| PABL_B-int | 1 | 1 | 1 | 0.004291845 | 0.63291 | 1 |
| Charlie10b | 0 | 1 | 0 | 0.003676471 | 1 | 1 |
| UCON4 | 0 | 1 | 0 | 0.008130081 | 1 | 1 |
| MER66D | 1 | 1 | 1 | 0.004115226 | 0.63288 | 1 |
| HY4 | 0 | 1 | 0 | 0.00617284 | 1 | 1 |
| HERV1_LTF | 0 | 1 | 0 | 0.0625 | 1 | 1 |
| LTR80B | 2 | 1 | 2 | 0.001956947 | 0.26424 | 1 |
| AluYk12 | 0 | 1 | 0 | 0.004464286 | 1 | 1 |
| MLT1E2-int | 1 | 1 | 1 | 0.007518797 | 0.63351 | 1 |
| MER41C | 2 | 3 | 0.666666667 | 0.003554502 | 0.80138 | 1 |
| L1M2b | 1 | 1 | 1 | 0.004166667 | 0.63289 | 1 |
| MSTD-int | 1 | 4 | 0.25 | 0.005420054 | 0.98188 | 1 |
| LTR3B | 0 | 1 | 0 | 0.010869565 | 1 | 1 |
| Eulor12 | 1 | 1 | 1 | 0.016666667 | 0.63521 | 1 |
| LTR5 | 0 | 1 | 0 | 0.05 | 1 | 1 |

| | | | | | | |
|---|---|---|---|---|---|---|
| LTR6A | 0 | 2 | 0 | 0.006944444 | 1 | 1 |
| MLT1F1-int | 0 | 1 | 0 | 0.00621118 | 1 | 1 |
| MER51C | 0 | 1 | 0 | 0.005586592 | 1 | 1 |
| LTR35A | 1 | 1 | 1 | 0.011111111 | 0.63417 | 1 |
| Charlie4 | 1 | 1 | 1 | 0.008264463 | 0.63365 | 1 |
| MER97b | 1 | 1 | 1 | 0.005494505 | 0.63313 | 1 |
| Eulor9A | 0 | 1 | 0 | 0.008130081 | 1 | 1 |
| MLT1J2-int | 1 | 1 | 1 | 0.001686341 | 0.63243 | 1 |
| MER34D | 0 | 1 | 0 | 0.004 | 1 | 1 |
| MER51-int | 0 | 3 | 0 | 0.005376344 | 1 | 1 |
| MER65C | 2 | 2 | 1 | 0.002928258 | 0.59439 | 1 |
| LTR12B | 0 | 1 | 0 | 0.004739336 | 1 | 1 |
| MER57D | 2 | 1 | 2 | 0.002183406 | 0.26424 | 1 |
| U17 | 0 | 1 | 0 | 0.111111111 | 1 | 1 |
| LTR35 | 0 | 1 | 0 | 0.008474576 | 1 | 1 |
| MER50B | 0 | 3 | 0 | 0.00433526 | 1 | 1 |
| MSTB1-int | 0 | 2 | 0 | 0.007751938 | 1 | 1 |
| UCON19 | 0 | 1 | 0 | 0.035714286 | 1 | 1 |
| L1MDb | 1 | 5 | 0.2 | 0.004612546 | 0.99334 | 1 |
| UCON7 | 0 | 1 | 0 | 0.007874016 | 1 | 1 |
| MLT1G-int | 0 | 1 | 0 | 0.005524862 | 1 | 1 |
| LTR31 | 0 | 1 | 0 | 0.004149378 | 1 | 1 |
| L1M3c | 1 | 4 | 0.25 | 0.003633061 | 0.98182 | 1 |
| HERVK22-ir | 4 | 4 | 1 | 0.009367681 | 0.56745 | 1 |
| LTR17 | 0 | 4 | 0 | 0.00476758 | 1 | 1 |
| MER83B | 0 | 1 | 0 | 0.006944444 | 1 | 1 |
| PrimLTR79 | 0 | 1 | 0 | 0.007092199 | 1 | 1 |
| SVA_A | 0 | 3 | 0 | 0.011152416 | 1 | 1 |
| MER74C | 1 | 1 | 1 | 0.004291845 | 0.63291 | 1 |
| UCON26 | 1 | 1 | 1 | 0.004854369 | 0.63302 | 1 |
| LTR88c | 1 | 1 | 1 | 0.000958773 | 0.6323 | 1 |
| SSU-rRNA_ | 1 | 1 | 1 | 0.0125 | 0.63443 | 1 |
| HUERS-P3b | 0 | 4 | 0 | 0.008733624 | 1 | 1 |
| LTR6B | 0 | 1 | 0 | 0.006493506 | 1 | 1 |
| HERVS71-ir | 1 | 3 | 0.333333333 | 0.017142857 | 0.95149 | 1 |
| HERVK11-ir | 0 | 3 | 0 | 0.010204082 | 1 | 1 |
| MLT1F2-int | 2 | 1 | 2 | 0.003333333 | 0.26424 | 1 |
| X5A_LINE | 0 | 1 | 0 | 0.00729927 | 1 | 1 |
| THE1-int | 0 | 1 | 0 | 0.001531394 | 1 | 1 |
| LTR86B2 | 0 | 1 | 0 | 0.003717472 | 1 | 1 |
| LTR69 | 2 | 1 | 2 | 0.007042254 | 0.26424 | 1 |
| ALINE | 0 | 1 | 0 | 0.003623188 | 1 | 1 |
| L1MEg2 | 0 | 2 | 0 | 0.002538071 | 1 | 1 |
| AluYc5 | 0 | 1 | 0 | 0.022222222 | 1 | 1 |
| HY5 | 0 | 1 | 0 | 0.045454545 | 1 | 1 |
| MER89-int | 1 | 2 | 0.5 | 0.002849003 | 0.86505 | 1 |
| MamGypLT | 3 | 2 | 1.5 | 0.002094241 | 0.32332 | 1 |

| | | | | | | |
|---|---|---|---|---|---|---|
| L1M3de | 0 | 2 | 0 | 0.004329004 | 1 | 1 |
| UCON12A | 0 | 1 | 0 | 0.038461538 | 1 | 1 |
| Eulor2B | 0 | 1 | 0 | 0.01754386 | 1 | 1 |
| BC200 | 0 | 1 | 0 | 0.005988024 | 1 | 1 |
| PABL_A-int | 1 | 3 | 0.333333333 | 0.007125891 | 0.95074 | 1 |
| AluYb9 | 0 | 1 | 0 | 0.003058104 | 1 | 1 |
| MLT1N2-in | 0 | 1 | 0 | 0.013157895 | 1 | 1 |
| MER70-int | 0 | 1 | 0 | 0.003875969 | 1 | 1 |
| L1M3d | 0 | 2 | 0 | 0.003710575 | 1 | 1 |
| LTR5A | 0 | 3 | 0 | 0.011320755 | 1 | 1 |
| LTR34 | 2 | 1 | 2 | 0.002590674 | 0.26424 | 1 |
| HERV17-int | 3 | 7 | 0.428571429 | 0.007981756 | 0.97081 | 1 |
| L1PBb | 2 | 2 | 1 | 0.007575758 | 0.59502 | 1 |
| SUBTEL_sa | 0 | 1 | 0 | 0.029411765 | 1 | 1 |
| MER76 | 2 | 2 | 1 | 0.002785515 | 0.59437 | 1 |
| MER97a | 0 | 1 | 0 | 0.003610108 | 1 | 1 |
| MLT1H2-in | 2 | 1 | 2 | 0.002857143 | 0.26424 | 1 |
| LTR27B | 1 | 2 | 0.5 | 0.003490401 | 0.86514 | 1 |
| LTR19-int | 0 | 1 | 0 | 0.004975124 | 1 | 1 |
| Merlin1_HS | 0 | 1 | 0 | 0.01754386 | 1 | 1 |
| MSTC-int | 1 | 2 | 0.5 | 0.009852217 | 0.866 | 1 |
| MER83B-in | 2 | 1 | 2 | 0.003846154 | 0.26424 | 1 |
| UCON5 | 0 | 1 | 0 | 0.009803922 | 1 | 1 |
| MER61F | 0 | 1 | 0 | 0.00621118 | 1 | 1 |
| GSAT | 0 | 1 | 0 | 0.014925373 | 1 | 1 |
| MLT1-int | 0 | 1 | 0 | 0.004405286 | 1 | 1 |
| Eulor9B | 0 | 1 | 0 | 0.04 | 1 | 1 |
| L1M3a | 0 | 3 | 0 | 0.004709576 | 1 | 1 |
| L1P3b | 0 | 1 | 0 | 0.012195122 | 1 | 1 |
| UCON31 | 0 | 1 | 0 | 0.009090909 | 1 | 1 |
| MER97d | 1 | 1 | 1 | 0.009803922 | 0.63393 | 1 |
| LTR68 | 0 | 1 | 0 | 0.002816901 | 1 | 1 |
| LTR12_ | 0 | 3 | 0 | 0.005405405 | 1 | 1 |
| L1P4d | 0 | 1 | 0 | 0.00625 | 1 | 1 |
| X6B_LINE | 0 | 1 | 0 | 0.002320186 | 1 | 1 |
| UCON6 | 1 | 1 | 1 | 0.011764706 | 0.6343 | 1 |
| LTR40A1 | 0 | 1 | 0 | 0.001760563 | 1 | 1 |
| UCON28b | 0 | 1 | 0 | 0.012987013 | 1 | 1 |
| L1MEa | 1 | 1 | 1 | 0.002645503 | 0.63261 | 1 |
| L1M2a1 | 2 | 1 | 2 | 0.008064516 | 0.26424 | 1 |
| MER132 | 0 | 1 | 0 | 0.027027027 | 1 | 1 |
| Eulor4 | 0 | 1 | 0 | 0.032258065 | 1 | 1 |
| MER125 | 0 | 1 | 0 | 0.007142857 | 1 | 1 |
| MER68B | 1 | 1 | 1 | 0.002222222 | 0.63253 | 1 |
| UCON8 | 0 | 1 | 0 | 0.01010101 | 1 | 1 |
| MER123 | 0 | 1 | 0 | 0.024390244 | 1 | 1 |
| MER95 | 0 | 1 | 0 | 0.006410256 | 1 | 1 |

| | | | | | | |
|---|---|---|---|---|---|---|
| Eulor6A | 0 | 1 | 0 | 0.033333333 | 1 | 1 |
| Eulor5A | 1 | 1 | 1 | 0.007874016 | 0.63357 | 1 |
| MER61C | 0 | 1 | 0 | 0.003184713 | 1 | 1 |
| HERVK14-i | 0 | 2 | 0 | 0.004975124 | 1 | 1 |
| MER61E | 0 | 1 | 0 | 0.002985075 | 1 | 1 |
| MER130 | 0 | 1 | 0 | 0.010309278 | 1 | 1 |
| MER133A | 0 | 1 | 0 | 0.018518519 | 1 | 1 |
| LTR18A | 1 | 1 | 1 | 0.003861004 | 0.63283 | 1 |
| HERVL18-i | 0 | 3 | 0 | 0.005535055 | 1 | 1 |
| Eulor9C | 0 | 1 | 0 | 0.009345794 | 1 | 1 |
| L1M3b | 0 | 3 | 0 | 0.005385996 | 1 | 1 |
| LTR25-int | 0 | 4 | 0 | 0.006791171 | 1 | 1 |
| HUERS-P2- | 0 | 2 | 0 | 0.007874016 | 1 | 1 |
| MER67D | 0 | 2 | 0 | 0.001974334 | 1 | 1 |
| L1P4b | 0 | 1 | 0 | 0.006578947 | 1 | 1 |
| MER83C | 0 | 1 | 0 | 0.006493506 | 1 | 1 |
| Charlie3 | 1 | 3 | 0.333333333 | 0.00877193 | 0.95087 | 1 |
| Eulor3 | 1 | 1 | 1 | 0.027777778 | 0.63729 | 1 |
| UCON18 | 0 | 1 | 0 | 0.055555556 | 1 | 1 |
| MER101B | 2 | 1 | 2 | 0.003861004 | 0.26424 | 1 |
| HERVL32-i | 0 | 1 | 0 | 0.008695652 | 1 | 1 |
| LTR58 | 0 | 1 | 0 | 0.013157895 | 1 | 1 |
| Eulor8 | 0 | 1 | 0 | 0.006993007 | 1 | 1 |
| Ricksha_b | 1 | 1 | 1 | 0.008333333 | 0.63366 | 1 |
| UCON25 | 0 | 1 | 0 | 0.023255814 | 1 | 1 |
| MER92C | 1 | 1 | 1 | 0.003012048 | 0.63268 | 1 |
| UCON13 | 0 | 1 | 0 | 0.019607843 | 1 | 1 |
| MLT1E-int | 0 | 1 | 0 | 0.055555556 | 1 | 1 |
| MER57C2 | 0 | 1 | 0 | 0.002141328 | 1 | 1 |
| UCON28c | 0 | 1 | 0 | 0.015384615 | 1 | 1 |
| Eulor6C | 0 | 1 | 0 | 0.045454545 | 1 | 1 |
| X2_LINE | 0 | 1 | 0 | 0.015384615 | 1 | 1 |
| LTR35B | 0 | 1 | 0 | 0.004830918 | 1 | 1 |
| L1PBa1 | 1 | 5 | 0.2 | 0.012315271 | 0.99347 | 1 |
| UCON2 | 0 | 1 | 0 | 0.008403361 | 1 | 1 |
| UCON15 | 0 | 1 | 0 | 0.022727273 | 1 | 1 |
| UCON27 | 0 | 1 | 0 | 0.009708738 | 1 | 1 |
| UCON10 | 0 | 1 | 0 | 0.015625 | 1 | 1 |
| Eulor2C | 0 | 1 | 0 | 0.026315789 | 1 | 1 |
| Eulor11 | 0 | 1 | 0 | 0.012345679 | 1 | 1 |
| ALR/Alpha | 19 | 17 | 1.117647059 | 0.013066872 | 0.34444 | 1 |
| MLT1G3-in | 0 | 1 | 0 | 0.004166667 | 1 | 1 |
| HERVH48-i | 0 | 2 | 0 | 0.016949153 | 1 | 1 |
| LTR30 | 0 | 1 | 0 | 0.006896552 | 1 | 1 |
| LTR52-int | 0 | 1 | 0 | 0.003731343 | 1 | 1 |
| UCON28a | 0 | 1 | 0 | 0.007751938 | 1 | 1 |
| Eulor7 | 0 | 1 | 0 | 0.090909091 | 1 | 1 |

| | | | | | | |
|---|---|---|---|---|---|---|
| MSTB2-int | 0 | 1 | 0 | 0.010989011 | 1 | 1 |
| MER133B | 0 | 1 | 0 | 0.024390244 | 1 | 1 |
| HERVKC4-ir | 0 | 1 | 0 | 0.02173913 | 1 | 1 |
| UCON1 | 0 | 1 | 0 | 0.034482759 | 1 | 1 |
| Eulor6B | 0 | 1 | 0 | 0.016949153 | 1 | 1 |
| LTR11 | 0 | 1 | 0 | 0.0625 | 1 | 1 |
| HERVE_a-ir | 0 | 4 | 0 | 0.020618557 | 1 | 1 |
| LTR22 | 0 | 1 | 0 | 0.009708738 | 1 | 1 |
| MER134 | 0 | 1 | 0 | 0.014925373 | 1 | 1 |
| MER87B | 0 | 1 | 0 | 0.004098361 | 1 | 1 |
| Charlie11 | 1 | 1 | 1 | 0.009259259 | 0.63383 | 1 |
| MER88 | 0 | 1 | 0 | 0.008849558 | 1 | 1 |
| UCON9 | 0 | 1 | 0 | 0.020408163 | 1 | 1 |
| LTR25 | 1 | 1 | 1 | 0.003546099 | 0.63277 | 1 |
| LTR3 | 0 | 1 | 0 | 0.010752688 | 1 | 1 |
| HERVK3-int | 0 | 4 | 0 | 0.012048193 | 1 | 1 |
| L1P | 0 | 1 | 0 | 0.006289308 | 1 | 1 |
| MER83A-in | 1 | 1 | 1 | 0.009345794 | 0.63385 | 1 |
| MER92A | 1 | 1 | 1 | 0.006578947 | 0.63333 | 1 |
| Charlie6 | 1 | 1 | 1 | 0.003846154 | 0.63283 | 1 |
| CER | 0 | 2 | 0 | 0.027777778 | 1 | 1 |
| HSAT5 | 0 | 1 | 0 | 0.003846154 | 1 | 1 |
| SATR1 | 1 | 7 | 0.142857143 | 0.012867647 | 0.99913 | 1 |
| ACRO1 | 0 | 1 | 0 | 0.016393443 | 1 | 1 |
| MLT1E1-int | 0 | 1 | 0 | 0.022222222 | 1 | 1 |
| UCON23 | 0 | 1 | 0 | 0.032258065 | 1 | 1 |
| Tigger2b | 0 | 1 | 0 | 0.009090909 | 1 | 1 |
| MER76-int | 0 | 1 | 0 | 0.006666667 | 1 | 1 |
| LTR14C | 0 | 1 | 0 | 0.00729927 | 1 | 1 |
| Eulor5B | 0 | 1 | 0 | 0.023255814 | 1 | 1 |
| LTR38C | 0 | 1 | 0 | 0.008064516 | 1 | 1 |
| HERV1_LTF | 0 | 1 | 0 | 0.037037037 | 1 | 1 |
| MLT1L-int | 0 | 1 | 0 | 0.009259259 | 1 | 1 |
| UCON24 | 0 | 1 | 0 | 0.045454545 | 1 | 1 |
| LTR10G | 0 | 1 | 0 | 0.008403361 | 1 | 1 |
| LTR59 | 1 | 1 | 1 | 0.008196721 | 0.63363 | 1 |
| MER136 | 0 | 1 | 0 | 0.035714286 | 1 | 1 |
| UCON17 | 0 | 1 | 0 | 0.03030303 | 1 | 1 |
| SATR2 | 0 | 4 | 0 | 0.015267176 | 1 | 1 |
| MER57E3 | 1 | 1 | 1 | 0.004098361 | 0.63288 | 1 |
| LTR70 | 1 | 1 | 1 | 0.006944444 | 0.6334 | 1 |
| HSAT4 | 0 | 2 | 0 | 0.02020202 | 1 | 1 |
| Eulor6E | 1 | 1 | 1 | 0.032258065 | 0.63814 | 1 |
| U8 | 2 | 1 | 2 | 0.034482759 | 0.2642 | 1 |
| L1P4c | 0 | 1 | 0 | 0.022222222 | 1 | 1 |
| HERVK11D- | 0 | 1 | 0 | 0.02173913 | 1 | 1 |
| UCON12 | 0 | 1 | 0 | 0.022222222 | 1 | 1 |

| | | | | | | |
|---|---|---|---|---|---|---|
| REP522 | 0 | 4 | 0 | 0.016393443 | 1 | 1 |
| HERV1_I-in | 1 | 2 | 0.5 | 0.040816327 | 0.87023 | 1 |
| HSATI | 0 | 1 | 0 | 0.022222222 | 1 | 1 |
| MER9B | 0 | 1 | 0 | 0.025641026 | 1 | 1 |
| Charlie1b_ | 0 | 1 | 0 | 0.5 | 1 | 1 |
| HERV30-int | 0 | 1 | 0 | 0.01369863 | 1 | 1 |
| CheshMITE | 0 | 1 | 0 | 1 | 1 | 1 |
| SST1 | 2 | 6 | 0.333333333 | 0.009803922 | 0.98301 | 1 |
| Tigger1a_M | 0 | 1 | 0 | 0.022727273 | 1 | 1 |
| LTR43-int | 0 | 1 | 0 | 0.006666667 | 1 | 1 |
| LSAU | 0 | 1 | 0 | 0.007751938 | 1 | 1 |
| HERV1_LTF | 0 | 1 | 0 | 0.032258065 | 1 | 1 |
| HERV-Fc1_ | 0 | 1 | 0 | 0.058823529 | 1 | 1 |
| HERV-Fc1_ | 0 | 1 | 0 | 0.333333333 | 1 | 1 |
| HERV-Fc2-i | 0 | 1 | 0 | 0.333333333 | 1 | 1 |
| HERV-Fc1-i | 0 | 1 | 0 | 0.142857143 | 1 | 1 |
| U14 | 0 | 1 | 0 | 0.142857143 | 1 | 1 |
| HAL1N1_M | 0 | 1 | 0 | 0.5 | 1 | 1 |
| HERV1_LTF | 0 | 1 | 0 | 0.029411765 | 1 | 1 |
| HERV15-int | 1 | 2 | 0.5 | 0.021505376 | 0.86759 | 1 |
| HERV-Fc1_ | 0 | 1 | 0 | 0.2 | 1 | 1 |
| HERV1_LTF | 0 | 1 | 0 | 0.1 | 1 | 1 |
| Charlie12 | 0 | 1 | 0 | 0.045454545 | 1 | 1 |
| Cheshire_M | 0 | 1 | 0 | 0.142857143 | 1 | 1 |
| HSAT6 | 0 | 1 | 0 | 0.058823529 | 1 | 1 |
| D20S16 | 0 | 1 | 0 | 0.003937008 | 1 | 1 |
| MLT1M-int | 0 | 1 | 0 | 1 | 1 | 1 |
| SAR | 1 | 1 | 1 | 1 | 1 | 1 |
| Tigger2a_C | 0 | 1 | 0 | 0.5 | 1 | 1 |

Table S2. Enriched TE families in accessible chromatin after infection for the L12 samples

| TE.family | Observed.instances | Expected.instances | Fold.enrichment | Prob.success | P.value | Q.value |
|---|---|---|---|---|---|---|
| MER44B | 49 | 3 | 16.33333333 | 0.00140779 | 1.28E-41 | 1.29E-38 |
| L1MB2 | 100 | 20 | 5 | 0.002197561 | 2.46E-37 | 1.25E-34 |
| MER41B | 59 | 7 | 8.428571429 | 0.002454418 | 3.38E-34 | 1.14E-31 |
| MER44D | 23 | 1 | 23 | 0.001329787 | 1.09E-23 | 2.76E-21 |
| L1MB7 | 132 | 57 | 2.315789474 | 0.002457532 | 1.45E-17 | 2.94E-15 |
| Tigger3b | 60 | 16 | 3.75 | 0.002452859 | 2.79E-17 | 4.72E-15 |
| Tigger7 | 33 | 5 | 6.6 | 0.001604107 | 9.36E-17 | 1.36E-14 |
| THE1B | 88 | 32 | 2.75 | 0.00142647 | 2.81E-16 | 3.57E-14 |
| Tigger3c | 25 | 3 | 8.333333333 | 0.001266357 | 2.78E-15 | 3.14E-13 |
| Tigger3a | 38 | 8 | 4.75 | 0.00150122 | 1.54E-14 | 1.56E-12 |
| THE1C | 47 | 14 | 3.357142857 | 0.001417865 | 3.16E-12 | 2.91E-10 |
| MLT1H | 44 | 13 | 3.384615385 | 0.001287894 | 1.17E-11 | 9.91E-10 |
| MER44C | 17 | 2 | 8.5 | 0.00234192 | 4.95E-11 | 3.86E-09 |
| MLT1L | 42 | 14 | 3 | 0.001159516 | 1.16E-09 | 8.40E-08 |
| MLT1F | 32 | 9 | 3.555555556 | 0.002094485 | 2.08E-09 | 1.40E-07 |
| LTR71A | 11 | 1 | 11 | 0.006024096 | 7.59E-09 | 4.81E-07 |
| MLT1I | 34 | 12 | 2.833333333 | 0.001082153 | 1.52E-07 | 9.04E-06 |
| MLT1K | 51 | 23 | 2.217391304 | 0.001265614 | 3.22E-07 | 1.82E-05 |
| LTR26 | 9 | 1 | 9 | 0.001623377 | 1.07E-06 | 5.45E-05 |
| MER57F | 9 | 1 | 9 | 0.002320186 | 1.05E-06 | 5.45E-05 |
| L1MB3 | 72 | 39 | 1.846153846 | 0.002246026 | 1.39E-06 | 5.95E-05 |
| MER81 | 12 | 2 | 6 | 0.000545108 | 1.35E-06 | 5.95E-05 |
| LTR8A | 23 | 7 | 3.285714286 | 0.002468265 | 1.30E-06 | 5.95E-05 |
| MLT1F1 | 21 | 6 | 3.5 | 0.001829826 | 1.41E-06 | 5.95E-05 |
| MER57E1 | 8 | 1 | 8 | 0.001996008 | 9.82E-06 | 0.00038 |
| HSATII | 8 | 1 | 8 | 0.002506266 | 9.71E-06 | 0.00038 |
| MLT1H2 | 17 | 5 | 3.4 | 0.00106067 | 1.96E-05 | 0.00074 |
| MIR3 | 112 | 74 | 1.513513514 | 0.00081461 | 2.33E-05 | 0.00084 |
| MLT1F2 | 27 | 11 | 2.454545455 | 0.001822399 | 3.20E-05 | 0.00112 |
| MER45B | 10 | 2 | 5 | 0.001520913 | 4.55E-05 | 0.00154 |
| MamSINE1 | 7 | 1 | 7 | 0.00059952 | 8.25E-05 | 0.0027 |
| MER51A | 11 | 3 | 3.666666667 | 0.003134796 | 0.00028 | 0.00898 |
| Tigger12c | 6 | 1 | 6 | 0.000791766 | 0.00059 | 0.01758 |
| LTR40c | 6 | 1 | 6 | 0.00166113 | 0.00058 | 0.01758 |
| MLT1J | 37 | 21 | 1.761904762 | 0.001375246 | 0.00099 | 0.02862 |
| Charlie25 | 8 | 2 | 4 | 0.001855288 | 0.00108 | 0.02967 |
| Tigger3 | 10 | 3 | 3.333333333 | 0.002469136 | 0.00108 | 0.02967 |
| LTR40a | 9 | 3 | 3 | 0.001526718 | 0.00377 | 0.09132 |
| Tigger3d | 5 | 1 | 5 | 0.001666667 | 0.00362 | 0.09132 |
| MLT1M | 9 | 3 | 3 | 0.001014885 | 0.00378 | 0.09132 |
| AmnSINE1 | 5 | 1 | 5 | 0.000896057 | 0.00364 | 0.09132 |
| Charlie13b | 5 | 1 | 5 | 0.00245098 | 0.0036 | 0.09132 |
| LTR84b | 7 | 2 | 3.5 | 0.00154202 | 0.0045 | 0.10604 |
| LTR13 | 8 | 3 | 2.666666667 | 0.00610998 | 0.01164 | 0.26742 |
| MER63A | 8 | 3 | 2.666666667 | 0.000853 | 0.01187 | 0.26742 |

| | | | | | | |
|---|---|---|---|---|---|---|
| THE1A | 14 | 7 | 2 | 0.001653674 | 0.01274 | 0.28086 |
| MLT1G | 11 | 5 | 2.2 | 0.001751927 | 0.01362 | 0.29376 |
| Charlie20a | 4 | 1 | 4 | 0.001138952 | 0.01892 | 0.31485 |
| LTR13_ | 4 | 1 | 4 | 0.022727273 | 0.01759 | 0.31485 |
| LTR26B | 4 | 1 | 4 | 0.002178649 | 0.01885 | 0.31485 |
| MER66B | 6 | 2 | 3 | 0.001393728 | 0.01649 | 0.31485 |
| MER107 | 4 | 1 | 4 | 0.002785515 | 0.01882 | 0.31485 |
| LTR33A_ | 6 | 2 | 3 | 0.001205546 | 0.0165 | 0.31485 |
| LTR16D | 4 | 1 | 4 | 0.000902527 | 0.01893 | 0.31485 |
| LTR24 | 4 | 1 | 4 | 0.003246753 | 0.01879 | 0.31485 |
| Tigger9b | 4 | 1 | 4 | 0.001308901 | 0.01891 | 0.31485 |
| Tigger14a | 4 | 1 | 4 | 0.00077101 | 0.01894 | 0.31485 |
| Charlie21a | 4 | 1 | 4 | 0.001345895 | 0.01891 | 0.31485 |
| Tigger12A | 4 | 1 | 4 | 0.001461988 | 0.0189 | 0.31485 |
| LTR44 | 4 | 1 | 4 | 0.004587156 | 0.01871 | 0.31485 |
| MER57E2 | 4 | 1 | 4 | 0.013157895 | 0.01818 | 0.31485 |
| MLT1N2 | 12 | 6 | 2 | 0.001019714 | 0.02003 | 0.32766 |
| LTR16A1 | 9 | 4 | 2.25 | 0.001424501 | 0.02128 | 0.33735 |
| MARNA | 9 | 4 | 2.25 | 0.00119403 | 0.02129 | 0.33735 |
| L1MB4 | 31 | 21 | 1.476190476 | 0.002259765 | 0.02402 | 0.37468 |
| MER20 | 27 | 18 | 1.5 | 0.001073153 | 0.02816 | 0.4327 |
| Plat_L3 | 7 | 3 | 2.333333333 | 0.000810154 | 0.03345 | 0.49876 |
| Tigger2b_I | 7 | 3 | 2.333333333 | 0.001461988 | 0.0334 | 0.49876 |
| MLT1J2 | 14 | 8 | 1.75 | 0.001155235 | 0.0341 | 0.50105 |
| LTR8 | 15 | 9 | 1.666666667 | 0.00254022 | 0.04126 | 0.59769 |
| MIRc | 107 | 90 | 1.188888889 | 0.000869632 | 0.04385 | 0.62627 |
| L1MB8 | 49 | 38 | 1.289473684 | 0.002285302 | 0.04839 | 0.6815 |
| MamRep1 | 5 | 2 | 2.5 | 0.001586043 | 0.05251 | 0.69449 |
| MER77B | 5 | 2 | 2.5 | 0.001589825 | 0.05251 | 0.69449 |
| MLT1G3 | 8 | 4 | 2 | 0.00147656 | 0.051 | 0.69449 |
| HERV9-int | 12 | 7 | 1.714285714 | 0.006763285 | 0.05274 | 0.69449 |
| HERVL40-i | 5 | 2 | 2.5 | 0.001523229 | 0.05252 | 0.69449 |
| MamRep6 | 9 | 5 | 1.8 | 0.001128668 | 0.06798 | 0.84133 |
| MLT1H1 | 10 | 6 | 1.666666667 | 0.001648352 | 0.08375 | 0.84133 |
| LTR83 | 3 | 1 | 3 | 0.001367989 | 0.08018 | 0.84133 |
| MLT1G1 | 10 | 6 | 1.666666667 | 0.001670379 | 0.08375 | 0.84133 |
| LTR13A | 3 | 1 | 3 | 0.005319149 | 0.07981 | 0.84133 |
| Arthur1B | 6 | 3 | 2 | 0.001295337 | 0.08379 | 0.84133 |
| Tigger15a | 10 | 6 | 1.666666667 | 0.001185302 | 0.0838 | 0.84133 |
| ERV3-16A3 | 3 | 1 | 3 | 0.000925926 | 0.08022 | 0.84133 |
| X3_LINE | 3 | 1 | 3 | 0.001526718 | 0.08016 | 0.84133 |
| MER82 | 10 | 6 | 1.666666667 | 0.001830384 | 0.08373 | 0.84133 |
| MER121 | 3 | 1 | 3 | 0.001081081 | 0.0802 | 0.84133 |
| Kanga1a | 3 | 1 | 3 | 0.001184834 | 0.08019 | 0.84133 |
| Kanga1b | 3 | 1 | 3 | 0.002659574 | 0.08006 | 0.84133 |
| LOR1a | 3 | 1 | 3 | 0.001239157 | 0.08019 | 0.84133 |
| MER57B1 | 3 | 1 | 3 | 0.000987167 | 0.08021 | 0.84133 |

| | | | | | | |
|---|---|---|---|---|---|---|
| HERVL66-i | 3 | 1 | 3 | 0.009433962 | 0.07943 | 0.84133 |
| LTR24B | 3 | 1 | 3 | 0.002136752 | 0.0801 | 0.84133 |
| MLT1D-int | 3 | 1 | 3 | 0.001564945 | 0.08016 | 0.84133 |
| Charlie17a | 3 | 1 | 3 | 0.000682594 | 0.08024 | 0.84133 |
| MER51B | 3 | 1 | 3 | 0.001396648 | 0.08017 | 0.84133 |
| LTR26E | 3 | 1 | 3 | 0.003289474 | 0.08 | 0.84133 |
| MLT2F | 6 | 3 | 2 | 0.001564945 | 0.08376 | 0.84133 |
| MER57B2 | 3 | 1 | 3 | 0.000905797 | 0.08022 | 0.84133 |
| LTR73 | 3 | 1 | 3 | 0.004608295 | 0.07988 | 0.84133 |
| MER21B | 11 | 7 | 1.571428571 | 0.002502681 | 0.09825 | 0.97676 |
| TAR1 | 2 | 1 | 2 | 0.00621118 | 0.26424 | 1 |
| L1MC | 6 | 17 | 0.352941176 | 0.001420692 | 0.99933 | 1 |
| MER5B | 7 | 18 | 0.388888889 | 0.000722109 | 0.99896 | 1 |
| L2a | 112 | 254 | 0.440944882 | 0.001484824 | 1 | 1 |
| L3 | 46 | 50 | 0.92 | 0.001110741 | 0.73326 | 1 |
| MIR | 171 | 175 | 0.977142857 | 0.000996549 | 0.62899 | 1 |
| L2b | 98 | 120 | 0.816666667 | 0.00122529 | 0.98254 | 1 |
| L2c | 151 | 170 | 0.888235294 | 0.001208648 | 0.93492 | 1 |
| AluSp | 15 | 108 | 0.138888889 | 0.002150495 | 1 | 1 |
| MER33 | 0 | 13 | 0 | 0.001355861 | 1 | 1 |
| MIRb | 237 | 223 | 1.062780269 | 0.000990429 | 0.1823 | 1 |
| MER53 | 1 | 5 | 0.2 | 0.000852515 | 0.99328 | 1 |
| MLT1A | 6 | 12 | 0.5 | 0.001323043 | 0.97972 | 1 |
| AluJo | 27 | 120 | 0.225 | 0.001659682 | 1 | 1 |
| L1MB5 | 28 | 24 | 1.166666667 | 0.002431365 | 0.23202 | 1 |
| AluYc | 3 | 8 | 0.375 | 0.000939077 | 0.98628 | 1 |
| L1PA6 | 3 | 50 | 0.06 | 0.008365401 | 1 | 1 |
| L1P1 | 1 | 11 | 0.090909091 | 0.003505417 | 0.99998 | 1 |
| AluJr | 31 | 132 | 0.234848485 | 0.001709734 | 1 | 1 |
| Charlie5 | 2 | 3 | 0.666666667 | 0.001173709 | 0.80103 | 1 |
| MLT1E1A- | 1 | 1 | 1 | 0.011363636 | 0.63422 | 1 |
| MLT1E1A | 4 | 5 | 0.8 | 0.00148721 | 0.73518 | 1 |
| AluSx | 38 | 260 | 0.146153846 | 0.001800093 | 1 | 1 |
| AluSz6 | 17 | 82 | 0.207317073 | 0.001792232 | 1 | 1 |
| LTR16C | 11 | 9 | 1.222222222 | 0.001357261 | 0.29393 | 1 |
| ERVL-E-int | 16 | 19 | 0.842105263 | 0.002128613 | 0.78548 | 1 |
| L1MA8 | 6 | 26 | 0.230769231 | 0.002320393 | 1 | 1 |
| L1M5 | 35 | 94 | 0.372340426 | 0.001460217 | 1 | 1 |
| L1MA9 | 10 | 34 | 0.294117647 | 0.00202973 | 1 | 1 |
| LTR12F | 2 | 2 | 1 | 0.003144654 | 0.59442 | 1 |
| MER45A | 0 | 3 | 0 | 0.000865801 | 1 | 1 |
| MER58A | 13 | 13 | 1 | 0.000963106 | 0.53695 | 1 |
| L1PA14 | 1 | 11 | 0.090909091 | 0.003583062 | 0.99998 | 1 |
| AluY | 25 | 228 | 0.109649123 | 0.001890281 | 1 | 1 |
| L2 | 102 | 90 | 1.133333333 | 0.001584926 | 0.11394 | 1 |
| FLAM_A | 1 | 14 | 0.071428571 | 0.000863558 | 1 | 1 |
| MER47A | 1 | 4 | 0.25 | 0.001324503 | 0.98173 | 1 |

| | | | | | | |
|---|---|---|---|---|---|---|
| AluSc | 10 | 66 | 0.151515152 | 0.001912101 | 1 | 1 |
| L1PA4 | 1 | 78 | 0.012820513 | 0.006543075 | 1 | 1 |
| LTR89 | 1 | 1 | 1 | 0.000865801 | 0.63228 | 1 |
| L1MC4a | 58 | 50 | 1.16 | 0.001801867 | 0.14465 | 1 |
| L1PA7 | 6 | 82 | 0.073170732 | 0.006286897 | 1 | 1 |
| L1PA16 | 7 | 42 | 0.166666667 | 0.002989111 | 1 | 1 |
| L1PA2 | 0 | 39 | 0 | 0.007931666 | 1 | 1 |
| AluSz | 33 | 196 | 0.168367347 | 0.001993024 | 1 | 1 |
| L1M2 | 3 | 23 | 0.130434783 | 0.002517238 | 1 | 1 |
| FLAM_C | 7 | 22 | 0.318181818 | 0.000966736 | 0.99994 | 1 |
| L1PREC2 | 4 | 25 | 0.16 | 0.00317864 | 1 | 1 |
| L1PB | 1 | 4 | 0.25 | 0.002250985 | 0.98177 | 1 |
| Tigger5 | 0 | 2 | 0 | 0.000759878 | 1 | 1 |
| AluSc5 | 1 | 12 | 0.083333333 | 0.001746725 | 0.99999 | 1 |
| L1M4c | 4 | 18 | 0.222222222 | 0.002908386 | 0.99998 | 1 |
| L1MC5 | 20 | 33 | 0.606060606 | 0.001583341 | 0.99407 | 1 |
| MER4D1 | 0 | 4 | 0 | 0.002629849 | 1 | 1 |
| MER4A | 0 | 3 | 0 | 0.00265252 | 1 | 1 |
| L5 | 0 | 1 | 0 | 0.001355014 | 1 | 1 |
| AluSg4 | 1 | 14 | 0.071428571 | 0.001867663 | 1 | 1 |
| AluSx3 | 5 | 59 | 0.084745763 | 0.001992301 | 1 | 1 |
| LTR33 | 7 | 12 | 0.583333333 | 0.001295896 | 0.95428 | 1 |
| MSTB1 | 0 | 8 | 0 | 0.001576976 | 1 | 1 |
| AluSq | 4 | 46 | 0.086956522 | 0.002100457 | 1 | 1 |
| L1ME4a | 17 | 57 | 0.298245614 | 0.001318529 | 1 | 1 |
| MLT1C | 28 | 33 | 0.848484848 | 0.001664649 | 0.83069 | 1 |
| AluJb | 49 | 246 | 0.199186992 | 0.001697195 | 1 | 1 |
| FAM | 1 | 5 | 0.2 | 0.001034768 | 0.99328 | 1 |
| AluSq2 | 13 | 113 | 0.115044248 | 0.002042402 | 1 | 1 |
| L1ME3E | 4 | 10 | 0.4 | 0.001966568 | 0.98972 | 1 |
| LTR33B | 2 | 2 | 1 | 0.001259446 | 0.59416 | 1 |
| MER65D | 2 | 1 | 2 | 0.002109705 | 0.26424 | 1 |
| MLT1D | 8 | 34 | 0.235294118 | 0.001639265 | 1 | 1 |
| L1MA5 | 2 | 12 | 0.166666667 | 0.002674393 | 0.99992 | 1 |
| AluSg | 14 | 88 | 0.159090909 | 0.002114521 | 1 | 1 |
| MER1A | 1 | 8 | 0.125 | 0.002470661 | 0.99967 | 1 |
| MER21C | 17 | 14 | 1.214285714 | 0.002544992 | 0.24386 | 1 |
| LTR41 | 1 | 3 | 0.333333333 | 0.001728111 | 0.95034 | 1 |
| LTR37A | 1 | 3 | 0.333333333 | 0.001525165 | 0.95033 | 1 |
| AluSg7 | 4 | 17 | 0.235294118 | 0.002019482 | 0.99996 | 1 |
| MER11B | 0 | 2 | 0 | 0.003649635 | 1 | 1 |
| AluJr4 | 8 | 29 | 0.275862069 | 0.001639808 | 1 | 1 |
| U6 | 0 | 1 | 0 | 0.000584454 | 1 | 1 |
| MER8 | 0 | 2 | 0 | 0.001027221 | 1 | 1 |
| AluSx1 | 26 | 230 | 0.113043478 | 0.002075232 | 1 | 1 |
| L1MC4 | 37 | 56 | 0.660714286 | 0.001890998 | 0.99714 | 1 |
| MSTD | 9 | 11 | 0.818181818 | 0.001435095 | 0.76821 | 1 |

| | | | | | | |
|---|---|---|---|---|---|---|
| Charlie4z | 7 | 5 | 1.4 | 0.000840195 | 0.23776 | 1 |
| MLT2B3 | 5 | 6 | 0.833333333 | 0.001811047 | 0.71519 | 1 |
| MER31-int | 0 | 5 | 0 | 0.002430724 | 1 | 1 |
| MER34A | 0 | 2 | 0 | 0.001354096 | 1 | 1 |
| L1MC3 | 7 | 31 | 0.225806452 | 0.002315506 | 1 | 1 |
| MER4A1 | 1 | 5 | 0.2 | 0.002236136 | 0.9933 | 1 |
| MER4E | 0 | 2 | 0 | 0.002713704 | 1 | 1 |
| L1M2c | 1 | 2 | 0.5 | 0.00462963 | 0.86529 | 1 |
| THE1D | 15 | 18 | 0.833333333 | 0.001423825 | 0.79211 | 1 |
| LTR64 | 2 | 1 | 2 | 0.002688172 | 0.26424 | 1 |
| MLT1H1-ir | 0 | 1 | 0 | 0.002325581 | 1 | 1 |
| THE1B-int | 1 | 19 | 0.052631579 | 0.004491726 | 1 | 1 |
| L1M7 | 5 | 7 | 0.714285714 | 0.001590186 | 0.82723 | 1 |
| L1MEc | 13 | 43 | 0.302325581 | 0.002287843 | 1 | 1 |
| L1MA4 | 2 | 27 | 0.074074074 | 0.002685765 | 1 | 1 |
| L1M4 | 17 | 36 | 0.472222222 | 0.001966676 | 0.99985 | 1 |
| L1MA7 | 3 | 18 | 0.166666667 | 0.002076604 | 1 | 1 |
| ERVL-int | 2 | 2 | 1 | 0.002531646 | 0.59434 | 1 |
| L1PA5 | 1 | 69 | 0.014492754 | 0.006085729 | 1 | 1 |
| L1MEd | 10 | 17 | 0.588235294 | 0.00158272 | 0.97396 | 1 |
| L1PA10 | 2 | 29 | 0.068965517 | 0.004041249 | 1 | 1 |
| L1PA8 | 0 | 28 | 0 | 0.003444882 | 1 | 1 |
| L1PA8A | 3 | 11 | 0.272727273 | 0.004417671 | 0.99881 | 1 |
| L1P4 | 0 | 7 | 0 | 0.001875167 | 1 | 1 |
| L1PA15-16 | 1 | 5 | 0.2 | 0.003546099 | 0.99332 | 1 |
| LTR22C | 0 | 1 | 0 | 0.002538071 | 1 | 1 |
| L1ME3D | 4 | 9 | 0.444444444 | 0.001974984 | 0.97886 | 1 |
| L1MA2 | 4 | 29 | 0.137931034 | 0.003859462 | 1 | 1 |
| L1ME3B | 3 | 16 | 0.1875 | 0.001892148 | 0.99998 | 1 |
| LTR39 | 2 | 3 | 0.666666667 | 0.002365931 | 0.80121 | 1 |
| MER103C | 12 | 8 | 1.5 | 0.000894454 | 0.11183 | 1 |
| MSTA | 9 | 29 | 0.310344828 | 0.001465979 | 1 | 1 |
| LTR37B | 1 | 2 | 0.5 | 0.000983284 | 0.8648 | 1 |
| MLT1E3 | 1 | 4 | 0.25 | 0.001962709 | 0.98176 | 1 |
| MER4E1 | 1 | 2 | 0.5 | 0.001906578 | 0.86492 | 1 |
| MER5A | 14 | 28 | 0.5 | 0.000803512 | 0.99872 | 1 |
| LTR9B | 0 | 1 | 0 | 0.001319261 | 1 | 1 |
| L1ME1 | 45 | 74 | 0.608108108 | 0.002331884 | 0.99989 | 1 |
| MER39B | 3 | 3 | 1 | 0.002544529 | 0.5771 | 1 |
| L1MD3 | 10 | 9 | 1.111111111 | 0.001696833 | 0.41259 | 1 |
| Tigger4b | 0 | 4 | 0 | 0.001586672 | 1 | 1 |
| Ricksha_0 | 1 | 1 | 1 | 0.002475248 | 0.63258 | 1 |
| Charlie9 | 1 | 2 | 0.5 | 0.001477105 | 0.86486 | 1 |
| MER31B | 2 | 3 | 0.666666667 | 0.002017485 | 0.80115 | 1 |
| MER112 | 6 | 5 | 1.2 | 0.001142857 | 0.38404 | 1 |
| MER124 | 1 | 1 | 1 | 0.001340483 | 0.63237 | 1 |
| MER3 | 5 | 10 | 0.5 | 0.000913743 | 0.9708 | 1 |

| | | | | | | |
|---|---|---|---|---|---|---|
| MER5A1 | 8 | 12 | 0.666666667 | 0.000753154 | 0.91058 | 1 |
| L1ME3 | 2 | 18 | 0.111111111 | 0.001914079 | 1 | 1 |
| L1MEg | 8 | 32 | 0.25 | 0.001765322 | 1 | 1 |
| BLACKJACI | 1 | 3 | 0.333333333 | 0.001407129 | 0.95032 | 1 |
| L1M3 | 5 | 12 | 0.416666667 | 0.001751825 | 0.99244 | 1 |
| LTR7 | 0 | 5 | 0 | 0.002133106 | 1 | 1 |
| MER67C | 0 | 3 | 0 | 0.00182704 | 1 | 1 |
| LTR67B | 6 | 4 | 1.5 | 0.001076137 | 0.21479 | 1 |
| LTR79 | 5 | 4 | 1.25 | 0.00098668 | 0.37116 | 1 |
| Charlie1b | 5 | 4 | 1.25 | 0.001313629 | 0.37116 | 1 |
| MER113A | 0 | 2 | 0 | 0.001252348 | 1 | 1 |
| MER102c | 2 | 3 | 0.666666667 | 0.000861821 | 0.80098 | 1 |
| MER41D | 0 | 2 | 0 | 0.004210526 | 1 | 1 |
| HAL1 | 38 | 49 | 0.775510204 | 0.001794346 | 0.95457 | 1 |
| MLT1A1 | 3 | 9 | 0.333333333 | 0.00133018 | 0.99379 | 1 |
| LTR78 | 7 | 7 | 1 | 0.001452584 | 0.5504 | 1 |
| L1M4b | 5 | 14 | 0.357142857 | 0.002142639 | 0.99821 | 1 |
| MLT1B | 10 | 26 | 0.384615385 | 0.001444124 | 0.99989 | 1 |
| L1MC1 | 15 | 35 | 0.428571429 | 0.002664231 | 0.99995 | 1 |
| L1PA3 | 1 | 77 | 0.012987013 | 0.007233443 | 1 | 1 |
| Charlie19a | 2 | 2 | 1 | 0.001244555 | 0.59416 | 1 |
| LTR2C | 0 | 1 | 0 | 0.003389831 | 1 | 1 |
| FRAM | 4 | 9 | 0.444444444 | 0.001039861 | 0.97882 | 1 |
| HAL1-3A_I | 0 | 1 | 0 | 0.000520291 | 1 | 1 |
| BSR/Beta | 10 | 8 | 1.25 | 0.004032258 | 0.28312 | 1 |
| AluSc8 | 4 | 42 | 0.095238095 | 0.00191405 | 1 | 1 |
| LOR1-int | 2 | 3 | 0.666666667 | 0.002162942 | 0.80118 | 1 |
| LTR57-int | 0 | 1 | 0 | 0.00228833 | 1 | 1 |
| LTR49-int | 0 | 5 | 0 | 0.002295684 | 1 | 1 |
| AluYk4 | 0 | 4 | 0 | 0.002140182 | 1 | 1 |
| HERVK9-in | 0 | 5 | 0 | 0.007473842 | 1 | 1 |
| MER9a2 | 0 | 1 | 0 | 0.003236246 | 1 | 1 |
| MER102a | 1 | 3 | 0.333333333 | 0.001052262 | 0.95029 | 1 |
| MER67A | 1 | 1 | 1 | 0.002087683 | 0.6325 | 1 |
| MER2 | 2 | 13 | 0.153846154 | 0.001402525 | 0.99997 | 1 |
| HERV16-in | 0 | 4 | 0 | 0.002129925 | 1 | 1 |
| MER63B | 2 | 3 | 0.666666667 | 0.001250521 | 0.80104 | 1 |
| MER4A1_ | 0 | 1 | 0 | 0.002487562 | 1 | 1 |
| AluYa5 | 0 | 6 | 0 | 0.001531394 | 1 | 1 |
| L1MEe | 4 | 20 | 0.2 | 0.001678134 | 1 | 1 |
| LTR12C | 6 | 23 | 0.260869565 | 0.008394161 | 0.99999 | 1 |
| L1MD | 5 | 15 | 0.333333333 | 0.001696065 | 0.99915 | 1 |
| L1MEf | 4 | 32 | 0.125 | 0.002241838 | 1 | 1 |
| MER49 | 4 | 4 | 1 | 0.002921841 | 0.56682 | 1 |
| L1ME2 | 18 | 28 | 0.642857143 | 0.00215186 | 0.98218 | 1 |
| MER50 | 0 | 6 | 0 | 0.002332815 | 1 | 1 |
| L1PB1 | 3 | 54 | 0.055555556 | 0.004001186 | 1 | 1 |

| | | | | | | |
|---|---|---|---|---|---|---|
| MER30 | 1 | 5 | 0.2 | 0.001204529 | 0.99328 | 1 |
| LTR46 | 0 | 1 | 0 | 0.003861004 | 1 | 1 |
| MER9a1 | 0 | 1 | 0 | 0.002915452 | 1 | 1 |
| MER54A | 4 | 2 | 2 | 0.002949853 | 0.14261 | 1 |
| MER68 | 4 | 3 | 1.333333333 | 0.002 | 0.35277 | 1 |
| MER4B | 0 | 4 | 0 | 0.002309469 | 1 | 1 |
| MER4B-int | 2 | 4 | 0.5 | 0.002721088 | 0.90872 | 1 |
| AluYf4 | 0 | 2 | 0 | 0.001450326 | 1 | 1 |
| LTR16B1 | 1 | 2 | 0.5 | 0.001677852 | 0.86489 | 1 |
| AluSx4 | 3 | 11 | 0.272727273 | 0.001902456 | 0.9988 | 1 |
| MER63D | 1 | 2 | 0.5 | 0.001158078 | 0.86482 | 1 |
| MER66C | 2 | 1 | 2 | 0.001697793 | 0.26424 | 1 |
| MER66-int | 1 | 3 | 0.333333333 | 0.004016064 | 0.95051 | 1 |
| AluYb8 | 0 | 4 | 0 | 0.001401542 | 1 | 1 |
| AluYk11 | 0 | 1 | 0 | 0.000957854 | 1 | 1 |
| 7SLRNA | 1 | 2 | 0.5 | 0.001350439 | 0.86485 | 1 |
| MER74B | 1 | 2 | 0.5 | 0.002143623 | 0.86495 | 1 |
| LTR5_Hs | 0 | 4 | 0 | 0.00620155 | 1 | 1 |
| L1M3e | 0 | 2 | 0 | 0.002469136 | 1 | 1 |
| LTR2B | 2 | 2 | 1 | 0.006134969 | 0.59483 | 1 |
| Harlequin- | 1 | 5 | 0.2 | 0.009174312 | 0.99342 | 1 |
| LTR75 | 1 | 1 | 1 | 0.003436426 | 0.63275 | 1 |
| LTR10A | 0 | 1 | 0 | 0.003194888 | 1 | 1 |
| HERVIP10F | 0 | 4 | 0 | 0.00845666 | 1 | 1 |
| HERVI-int | 0 | 2 | 0 | 0.013888889 | 1 | 1 |
| HUERS-P1- | 0 | 2 | 0 | 0.003960396 | 1 | 1 |
| LTR7B | 0 | 2 | 0 | 0.002358491 | 1 | 1 |
| Charlie22a | 0 | 1 | 0 | 0.000898473 | 1 | 1 |
| MER65A | 3 | 3 | 1 | 0.002123142 | 0.57705 | 1 |
| MER65-int | 2 | 3 | 0.666666667 | 0.003128259 | 0.80132 | 1 |
| MER52A | 0 | 9 | 0 | 0.004926108 | 1 | 1 |
| MER52C | 1 | 2 | 0.5 | 0.005333333 | 0.86539 | 1 |
| MER83 | 0 | 1 | 0 | 0.002150538 | 1 | 1 |
| MER113 | 2 | 5 | 0.4 | 0.001308901 | 0.95966 | 1 |
| LTR42 | 0 | 1 | 0 | 0.003236246 | 1 | 1 |
| MER4D | 0 | 3 | 0 | 0.002252252 | 1 | 1 |
| Tigger1a_A | 0 | 1 | 0 | 0.003289474 | 1 | 1 |
| MER65B | 1 | 1 | 1 | 0.004854369 | 0.63302 | 1 |
| L1ME2z | 5 | 13 | 0.384615385 | 0.001694915 | 0.99628 | 1 |
| L1P5 | 0 | 1 | 0 | 0.001436782 | 1 | 1 |
| ORSL-2b | 0 | 1 | 0 | 0.001398601 | 1 | 1 |
| Tigger1 | 33 | 45 | 0.733333333 | 0.003716244 | 0.97373 | 1 |
| L1MCa | 3 | 17 | 0.176470588 | 0.002231264 | 0.99999 | 1 |
| MamRep1 | 2 | 2 | 1 | 0.001148106 | 0.59415 | 1 |
| Arthur1A | 0 | 1 | 0 | 0.000866551 | 1 | 1 |
| L1MEb | 3 | 3 | 1 | 0.001649258 | 0.57699 | 1 |
| L3b | 7 | 6 | 1.166666667 | 0.000868056 | 0.3937 | 1 |

| | | | | | | |
|---|---|---|---|---|---|---|
| AluSq4 | 0 | 3 | 0 | 0.002096436 | 1 | 1 |
| AluYc3 | 0 | 1 | 0 | 0.001754386 | 1 | 1 |
| Charlie15a | 1 | 3 | 0.333333333 | 0.000877706 | 0.95028 | 1 |
| AluSq10 | 1 | 4 | 0.25 | 0.001624695 | 0.98174 | 1 |
| Tigger2 | 3 | 8 | 0.375 | 0.002930403 | 0.98634 | 1 |
| MER44A | 2 | 4 | 0.5 | 0.001702852 | 0.90861 | 1 |
| MER119 | 3 | 2 | 1.5 | 0.001658375 | 0.32332 | 1 |
| X7A_LINE | 0 | 1 | 0 | 0.001078749 | 1 | 1 |
| MER1B | 1 | 9 | 0.111111111 | 0.001675354 | 0.99988 | 1 |
| Charlie1 | 4 | 4 | 1 | 0.002080083 | 0.56673 | 1 |
| Charlie1a | 13 | 13 | 1 | 0.002237136 | 0.53702 | 1 |
| LSU-rRNA_ | 1 | 2 | 0.5 | 0.004830918 | 0.86532 | 1 |
| LTR16E1 | 2 | 3 | 0.666666667 | 0.001212121 | 0.80103 | 1 |
| ERV3-16A3 | 9 | 18 | 0.5 | 0.002782931 | 0.993 | 1 |
| MLT2D | 2 | 5 | 0.4 | 0.001104972 | 0.95965 | 1 |
| Charlie2b | 2 | 6 | 0.333333333 | 0.001626457 | 0.98271 | 1 |
| MER89 | 3 | 2 | 1.5 | 0.001437815 | 0.32332 | 1 |
| MER91B | 0 | 1 | 0 | 0.000622278 | 1 | 1 |
| MER31A | 0 | 3 | 0 | 0.002220577 | 1 | 1 |
| MLT1A0 | 15 | 29 | 0.517241379 | 0.001404835 | 0.99841 | 1 |
| MSTB | 6 | 13 | 0.461538462 | 0.001518337 | 0.98931 | 1 |
| LTR18B | 0 | 1 | 0 | 0.002493766 | 1 | 1 |
| MER102b | 6 | 5 | 1.2 | 0.001226392 | 0.38404 | 1 |
| MER46C | 1 | 4 | 0.25 | 0.001420455 | 0.98174 | 1 |
| Charlie8 | 2 | 4 | 0.5 | 0.001364256 | 0.90857 | 1 |
| LTR81B | 0 | 1 | 0 | 0.000862813 | 1 | 1 |
| LTR81 | 1 | 1 | 1 | 0.00308642 | 0.63269 | 1 |
| MER58B | 4 | 8 | 0.5 | 0.001140088 | 0.9577 | 1 |
| MER74A | 2 | 3 | 0.666666667 | 0.001896334 | 0.80114 | 1 |
| LTR52 | 1 | 1 | 1 | 0.000925069 | 0.63229 | 1 |
| Kanga1 | 1 | 1 | 1 | 0.00122399 | 0.63235 | 1 |
| MLT-int | 1 | 1 | 1 | 0.002832861 | 0.63264 | 1 |
| MLT2C1 | 4 | 3 | 1.333333333 | 0.001145038 | 0.35277 | 1 |
| L1ME3F | 5 | 10 | 0.5 | 0.002021427 | 0.97086 | 1 |
| MER91A | 3 | 3 | 1 | 0.000825309 | 0.5769 | 1 |
| HAL1-2a_M | 1 | 1 | 1 | 0.000672043 | 0.63224 | 1 |
| MLT2B1 | 7 | 6 | 1.166666667 | 0.001339286 | 0.3937 | 1 |
| MER47B | 1 | 1 | 1 | 0.00110742 | 0.63232 | 1 |
| LTR81C | 0 | 1 | 0 | 0.001375516 | 1 | 1 |
| Tigger2a | 0 | 5 | 0 | 0.001498801 | 1 | 1 |
| MER34C_ | 1 | 1 | 1 | 0.001526718 | 0.6324 | 1 |
| MER58C | 1 | 2 | 0.5 | 0.000765404 | 0.86477 | 1 |
| MER34 | 0 | 2 | 0 | 0.001506024 | 1 | 1 |
| LTR19A | 0 | 1 | 0 | 0.002232143 | 1 | 1 |
| L1MCc | 5 | 6 | 0.833333333 | 0.00175644 | 0.71518 | 1 |
| MSTA-int | 3 | 11 | 0.272727273 | 0.003482115 | 0.9988 | 1 |
| HERVL-int | 7 | 14 | 0.5 | 0.006151142 | 0.98599 | 1 |

| | | | | | | |
|---|---|---|---|---|---|---|
| LTR50 | 5 | 4 | 1.25 | 0.001526718 | 0.37116 | 1 |
| LTR27 | 0 | 1 | 0 | 0.002427184 | 1 | 1 |
| MER117 | 5 | 3 | 1.666666667 | 0.00066726 | 0.18468 | 1 |
| L4 | 10 | 19 | 0.526315789 | 0.001126928 | 0.99117 | 1 |
| MLT1H-int | 0 | 1 | 0 | 0.001243781 | 1 | 1 |
| MER75 | 0 | 1 | 0 | 0.002105263 | 1 | 1 |
| MLT2B4 | 9 | 6 | 1.5 | 0.001308044 | 0.15263 | 1 |
| L1ME3A | 10 | 29 | 0.344827586 | 0.001801914 | 0.99999 | 1 |
| MER96 | 0 | 1 | 0 | 0.000784314 | 1 | 1 |
| LTR40b | 1 | 1 | 1 | 0.00087108 | 0.63228 | 1 |
| MLT1E2 | 3 | 8 | 0.375 | 0.002002002 | 0.98631 | 1 |
| ERVL-B4-ir | 8 | 11 | 0.727272727 | 0.002832861 | 0.85717 | 1 |
| Arthur1C | 2 | 1 | 2 | 0.001398601 | 0.26424 | 1 |
| MER45C | 1 | 2 | 0.5 | 0.001670844 | 0.86489 | 1 |
| THE1C-int | 2 | 7 | 0.285714286 | 0.003850385 | 0.99278 | 1 |
| L1PA12 | 1 | 7 | 0.142857143 | 0.003930376 | 0.9991 | 1 |
| AluYg6 | 0 | 1 | 0 | 0.001254705 | 1 | 1 |
| MSTB-int | 0 | 3 | 0 | 0.003420753 | 1 | 1 |
| L1MDa | 4 | 20 | 0.2 | 0.002746875 | 1 | 1 |
| L1PA15 | 4 | 31 | 0.129032258 | 0.003670811 | 1 | 1 |
| UCON16 | 0 | 1 | 0 | 0.027027027 | 1 | 1 |
| Eulor10 | 0 | 1 | 0 | 0.029411765 | 1 | 1 |
| L1MA10 | 1 | 7 | 0.142857143 | 0.001371205 | 0.99909 | 1 |
| L1MC2 | 8 | 16 | 0.5 | 0.002484472 | 0.99007 | 1 |
| L1ME3C | 9 | 22 | 0.409090909 | 0.001745616 | 0.99943 | 1 |
| L1MA5A | 4 | 8 | 0.5 | 0.002338498 | 0.95779 | 1 |
| LTR10B1 | 1 | 1 | 1 | 0.004115226 | 0.63288 | 1 |
| LTR21B | 0 | 1 | 0 | 0.016666667 | 1 | 1 |
| LTR62 | 0 | 1 | 0 | 0.003174603 | 1 | 1 |
| L1PA11 | 0 | 18 | 0 | 0.004302103 | 1 | 1 |
| THE1D-int | 0 | 7 | 0 | 0.00372737 | 1 | 1 |
| Charlie4a | 1 | 5 | 0.2 | 0.001435544 | 0.99329 | 1 |
| MER34C2 | 0 | 1 | 0 | 0.001779359 | 1 | 1 |
| MER34B-ir | 1 | 2 | 0.5 | 0.00295858 | 0.86507 | 1 |
| MER39 | 0 | 5 | 0 | 0.001498352 | 1 | 1 |
| MADE2 | 1 | 1 | 1 | 0.000374953 | 0.63219 | 1 |
| MSR1 | 0 | 2 | 0 | 0.006329114 | 1 | 1 |
| Helitron3N | 1 | 1 | 1 | 0.000897666 | 0.63229 | 1 |
| LTR5B | 0 | 3 | 0 | 0.006960557 | 1 | 1 |
| LTR45C | 0 | 2 | 0 | 0.004705882 | 1 | 1 |
| LTR38 | 0 | 1 | 0 | 0.003225806 | 1 | 1 |
| LTR38-int | 0 | 1 | 0 | 0.008130081 | 1 | 1 |
| HERVK13-i | 0 | 1 | 0 | 0.014705882 | 1 | 1 |
| LTR3B_ | 0 | 1 | 0 | 0.003816794 | 1 | 1 |
| MLT1E1 | 0 | 2 | 0 | 0.001917546 | 1 | 1 |
| LTR78B | 3 | 4 | 0.75 | 0.001219141 | 0.76208 | 1 |
| L1MB1 | 7 | 13 | 0.538461538 | 0.002081999 | 0.97422 | 1 |

| | | | | | | |
|---|---|---|---|---|---|---|
| Tigger4a | 1 | 4 | 0.25 | 0.001145803 | 0.98173 | 1 |
| L1MD2 | 25 | 27 | 0.925925926 | 0.002430024 | 0.67609 | 1 |
| MER115 | 1 | 4 | 0.25 | 0.001518027 | 0.98174 | 1 |
| MER77 | 3 | 3 | 1 | 0.002316602 | 0.57707 | 1 |
| LTR3A | 0 | 1 | 0 | 0.006134969 | 1 | 1 |
| LTR33C | 2 | 1 | 2 | 0.001079914 | 0.26424 | 1 |
| LTR85c | 0 | 1 | 0 | 0.00084317 | 1 | 1 |
| Kanga1d | 0 | 1 | 0 | 0.001501502 | 1 | 1 |
| MLT2B2 | 1 | 3 | 0.333333333 | 0.001358081 | 0.95031 | 1 |
| MER85 | 0 | 1 | 0 | 0.001104972 | 1 | 1 |
| MLT1J1 | 8 | 5 | 1.6 | 0.001015228 | 0.13327 | 1 |
| MER92B | 1 | 2 | 0.5 | 0.002239642 | 0.86497 | 1 |
| LTR10E | 0 | 1 | 0 | 0.003436426 | 1 | 1 |
| MER63C | 1 | 2 | 0.5 | 0.002244669 | 0.86497 | 1 |
| LTR16E2 | 4 | 3 | 1.333333333 | 0.001295337 | 0.35277 | 1 |
| L1PA13 | 4 | 30 | 0.133333333 | 0.003299241 | 1 | 1 |
| polypurine | 1 | 1 | 1 | 0.00073692 | 0.63226 | 1 |
| L1ME5 | 4 | 6 | 0.666666667 | 0.001740139 | 0.84903 | 1 |
| MamRep4 | 0 | 2 | 0 | 0.001572327 | 1 | 1 |
| L1P2 | 0 | 10 | 0 | 0.006501951 | 1 | 1 |
| polypyrimi | 0 | 1 | 0 | 0.000766284 | 1 | 1 |
| LTR85a | 0 | 2 | 0 | 0.001050972 | 1 | 1 |
| LTR24C | 1 | 1 | 1 | 0.002617801 | 0.6326 | 1 |
| L1MA1 | 1 | 15 | 0.066666667 | 0.003472222 | 1 | 1 |
| MSTC | 0 | 5 | 0 | 0.001577785 | 1 | 1 |
| MER21A | 3 | 5 | 0.6 | 0.002602811 | 0.87568 | 1 |
| MSTB2 | 3 | 3 | 1 | 0.001871491 | 0.57702 | 1 |
| MLT2A2 | 5 | 7 | 0.714285714 | 0.001795793 | 0.82725 | 1 |
| MER48 | 2 | 1 | 2 | 0.005181347 | 0.26424 | 1 |
| HERVH-int | 0 | 26 | 0 | 0.004400068 | 1 | 1 |
| LTR7A | 0 | 1 | 0 | 0.066666667 | 1 | 1 |
| L1PA17 | 1 | 13 | 0.076923077 | 0.002671599 | 1 | 1 |
| L1PB4 | 3 | 17 | 0.176470588 | 0.002263347 | 0.99999 | 1 |
| LTR53 | 0 | 1 | 0 | 0.00149925 | 1 | 1 |
| MER61B | 0 | 1 | 0 | 0.002320186 | 1 | 1 |
| MLT1A0-ir | 1 | 2 | 0.5 | 0.002159827 | 0.86496 | 1 |
| LTR16A2 | 1 | 2 | 0.5 | 0.001085776 | 0.86481 | 1 |
| MER90 | 2 | 2 | 1 | 0.002849003 | 0.59438 | 1 |
| LTR1 | 1 | 4 | 0.25 | 0.002762431 | 0.98179 | 1 |
| MER11A | 2 | 5 | 0.4 | 0.005186722 | 0.95992 | 1 |
| LTR81AB | 0 | 1 | 0 | 0.003663004 | 1 | 1 |
| LTR1B | 0 | 3 | 0 | 0.002555366 | 1 | 1 |
| MER61D | 0 | 1 | 0 | 0.003676471 | 1 | 1 |
| MER61A | 0 | 1 | 0 | 0.001231527 | 1 | 1 |
| Arthur1 | 3 | 4 | 0.75 | 0.001716002 | 0.76215 | 1 |
| X7B_LINE | 0 | 1 | 0 | 0.001057082 | 1 | 1 |
| L1MA6 | 4 | 14 | 0.285714286 | 0.002560351 | 0.99953 | 1 |

| | | | | | | |
|---|---|---|---|---|---|---|
| MamRep4 | 1 | 2 | 0.5 | 0.0009298 | 0.86479 | 1 |
| L1PB2 | 0 | 9 | 0 | 0.003140265 | 1 | 1 |
| L1P3 | 0 | 11 | 0 | 0.003222027 | 1 | 1 |
| CR1_Mam | 3 | 2 | 1.5 | 0.001156069 | 0.32332 | 1 |
| MER45R | 1 | 1 | 1 | 0.001607717 | 0.63242 | 1 |
| Tigger11a | 0 | 1 | 0 | 0.002314815 | 1 | 1 |
| MER110 | 0 | 1 | 0 | 0.00128041 | 1 | 1 |
| LTR66 | 0 | 1 | 0 | 0.00265252 | 1 | 1 |
| MER54B | 1 | 1 | 1 | 0.002304147 | 0.63254 | 1 |
| MER70A | 1 | 1 | 1 | 0.001869159 | 0.63246 | 1 |
| MLT1E | 0 | 1 | 0 | 0.001101322 | 1 | 1 |
| MamRep5 | 1 | 1 | 1 | 0.001257862 | 0.63235 | 1 |
| L1MD1 | 9 | 21 | 0.428571429 | 0.003022018 | 0.99891 | 1 |
| LTR90B | 1 | 1 | 1 | 0.001858736 | 0.63246 | 1 |
| MER20B | 5 | 6 | 0.833333333 | 0.001444391 | 0.71514 | 1 |
| MER57A1 | 4 | 2 | 2 | 0.001885014 | 0.14271 | 1 |
| LTR10C | 0 | 2 | 0 | 0.003861004 | 1 | 1 |
| MLT1J-int | 2 | 2 | 1 | 0.001386001 | 0.59418 | 1 |
| MADE1 | 0 | 3 | 0 | 0.000383485 | 1 | 1 |
| MLT2A1 | 0 | 5 | 0 | 0.001322751 | 1 | 1 |
| LTR48B | 0 | 2 | 0 | 0.001958864 | 1 | 1 |
| MER34-int | 1 | 1 | 1 | 0.002525253 | 0.63259 | 1 |
| MER70B | 0 | 1 | 0 | 0.002087683 | 1 | 1 |
| LFSINE_Ve | 0 | 1 | 0 | 0.001039501 | 1 | 1 |
| Kanga2_a | 0 | 2 | 0 | 0.001301236 | 1 | 1 |
| MamGypL' | 1 | 1 | 1 | 0.00091659 | 0.63229 | 1 |
| MamRep3 | 0 | 2 | 0 | 0.000940734 | 1 | 1 |
| L1PB3 | 2 | 11 | 0.181818182 | 0.003037835 | 0.9998 | 1 |
| LTR23 | 0 | 1 | 0 | 0.001177856 | 1 | 1 |
| LTR23-int | 1 | 1 | 1 | 0.003584229 | 0.63278 | 1 |
| MER34B | 1 | 1 | 1 | 0.001494768 | 0.6324 | 1 |
| LTR54 | 4 | 2 | 2 | 0.001784121 | 0.14272 | 1 |
| MER34C | 0 | 1 | 0 | 0.005434783 | 1 | 1 |
| MER75A | 0 | 1 | 0 | 0.010869565 | 1 | 1 |
| LTR16A | 13 | 9 | 1.444444444 | 0.00129199 | 0.12409 | 1 |
| L1M6 | 6 | 9 | 0.666666667 | 0.001306051 | 0.88447 | 1 |
| MER34A1 | 0 | 1 | 0 | 0.001302083 | 1 | 1 |
| MamRep1 | 1 | 1 | 1 | 0.000674764 | 0.63224 | 1 |
| MER94 | 4 | 3 | 1.333333333 | 0.000625261 | 0.35277 | 1 |
| LTR16B2 | 1 | 3 | 0.333333333 | 0.001851852 | 0.95035 | 1 |
| Zaphod3 | 0 | 1 | 0 | 0.000806452 | 1 | 1 |
| MER58D | 0 | 1 | 0 | 0.000866551 | 1 | 1 |
| LTR33A | 2 | 3 | 0.666666667 | 0.001475652 | 0.80107 | 1 |
| MER41A | 5 | 6 | 0.833333333 | 0.002352941 | 0.71526 | 1 |
| MER5C | 1 | 2 | 0.5 | 0.001488095 | 0.86487 | 1 |
| MLT1C-int | 1 | 1 | 1 | 0.002169197 | 0.63252 | 1 |
| Tigger13a | 7 | 5 | 1.4 | 0.001603592 | 0.2377 | 1 |

| | | | | | | |
|---|---|---|---|---|---|---|
| LTR16D2 | 0 | 1 | 0 | 0.003558719 | 1 | 1 |
| ORSL | 0 | 1 | 0 | 0.000823723 | 1 | 1 |
| HAL1b | 5 | 6 | 0.833333333 | 0.001243266 | 0.71511 | 1 |
| MLT1G1-ir | 0 | 1 | 0 | 0.003257329 | 1 | 1 |
| LTR1C | 0 | 1 | 0 | 0.004032258 | 1 | 1 |
| THE1A-int | 0 | 8 | 0 | 0.005464481 | 1 | 1 |
| LTR41B | 4 | 2 | 2 | 0.002014099 | 0.14269 | 1 |
| MLT1B-int | 1 | 1 | 1 | 0.001798561 | 0.63245 | 1 |
| MER105 | 1 | 1 | 1 | 0.001385042 | 0.63238 | 1 |
| LTR19B | 0 | 1 | 0 | 0.003558719 | 1 | 1 |
| HERVIP10F | 3 | 3 | 1 | 0.004405286 | 0.5773 | 1 |
| Charlie16a | 0 | 2 | 0 | 0.001010101 | 1 | 1 |
| FordPrefec | 0 | 1 | 0 | 0.003546099 | 1 | 1 |
| MER11C | 0 | 4 | 0 | 0.004618938 | 1 | 1 |
| LTR71B | 1 | 1 | 1 | 0.004201681 | 0.63289 | 1 |
| LTR16D1 | 0 | 1 | 0 | 0.004878049 | 1 | 1 |
| L1MA3 | 4 | 35 | 0.114285714 | 0.003875111 | 1 | 1 |
| Charlie18a | 0 | 3 | 0 | 0.001208216 | 1 | 1 |
| MER51E | 0 | 1 | 0 | 0.004016064 | 1 | 1 |
| LTR86A1 | 2 | 1 | 2 | 0.001904762 | 0.26424 | 1 |
| U2 | 0 | 1 | 0 | 0.000918274 | 1 | 1 |
| LTR75_1 | 0 | 1 | 0 | 0.0078125 | 1 | 1 |
| MER72 | 1 | 2 | 0.5 | 0.002557545 | 0.86501 | 1 |
| SVA_D | 0 | 13 | 0 | 0.009110021 | 1 | 1 |
| AluYa8 | 0 | 1 | 0 | 0.00295858 | 1 | 1 |
| MER52D | 2 | 3 | 0.666666667 | 0.004594181 | 0.80154 | 1 |
| LTR12E | 2 | 1 | 2 | 0.008695652 | 0.26424 | 1 |
| HERVFH21 | 0 | 1 | 0 | 0.008474576 | 1 | 1 |
| LTR14B | 0 | 1 | 0 | 0.002754821 | 1 | 1 |
| Cheshire | 3 | 2 | 1.5 | 0.001970443 | 0.32332 | 1 |
| LTR12 | 2 | 2 | 1 | 0.002583979 | 0.59434 | 1 |
| MER101 | 0 | 1 | 0 | 0.002724796 | 1 | 1 |
| X7C_LINE | 1 | 1 | 1 | 0.001607717 | 0.63242 | 1 |
| LTR36 | 0 | 1 | 0 | 0.00204499 | 1 | 1 |
| MER126 | 1 | 1 | 1 | 0.004694836 | 0.63299 | 1 |
| UCON22 | 0 | 1 | 0 | 0.028571429 | 1 | 1 |
| U13_ | 0 | 1 | 0 | 0.003257329 | 1 | 1 |
| Charlie7a | 0 | 1 | 0 | 0.000626566 | 1 | 1 |
| MamRep1 | 4 | 2 | 2 | 0.000848896 | 0.1428 | 1 |
| MER94B | 0 | 1 | 0 | 0.001607717 | 1 | 1 |
| Eulor2A | 0 | 1 | 0 | 0.014925373 | 1 | 1 |
| X9_LINE | 0 | 1 | 0 | 0.011904762 | 1 | 1 |
| UCON20 | 0 | 1 | 0 | 0.022727273 | 1 | 1 |
| LTR86A2 | 0 | 1 | 0 | 0.003194888 | 1 | 1 |
| X5B_LINE | 0 | 1 | 0 | 0.01010101 | 1 | 1 |
| U1 | 0 | 1 | 0 | 0.004878049 | 1 | 1 |
| Mam_R4 | 0 | 1 | 0 | 0.001798561 | 1 | 1 |

| | | | | | | |
|---|---|---|---|---|---|---|
| MLT2C2 | 4 | 2 | 2 | 0.001511716 | 0.14274 | 1 |
| Ricksha_c | 2 | 1 | 2 | 0.000807754 | 0.26424 | 1 |
| MER96B | 0 | 2 | 0 | 0.000933707 | 1 | 1 |
| MER131 | 0 | 1 | 0 | 0.002352941 | 1 | 1 |
| MER91C | 0 | 1 | 0 | 0.000614251 | 1 | 1 |
| Charlie24 | 2 | 1 | 2 | 0.000753012 | 0.26424 | 1 |
| Tigger16a | 0 | 1 | 0 | 0.001414427 | 1 | 1 |
| MER75B | 0 | 1 | 0 | 0.008849558 | 1 | 1 |
| 5S | 0 | 1 | 0 | 0.000784314 | 1 | 1 |
| MER90a | 3 | 2 | 1.5 | 0.001603849 | 0.32332 | 1 |
| L1MA4A | 4 | 21 | 0.19047619 | 0.003341289 | 1 | 1 |
| Zaphod | 2 | 4 | 0.5 | 0.001887683 | 0.90863 | 1 |
| SVA_E | 0 | 2 | 0 | 0.008474576 | 1 | 1 |
| MER84 | 1 | 1 | 1 | 0.002754821 | 0.63263 | 1 |
| MER84-int | 0 | 1 | 0 | 0.006896552 | 1 | 1 |
| HY1 | 0 | 1 | 0 | 0.002252252 | 1 | 1 |
| Charlie10 | 3 | 2 | 1.5 | 0.00177305 | 0.32332 | 1 |
| LTR9 | 1 | 5 | 0.2 | 0.002486325 | 0.9933 | 1 |
| Charlie26a | 1 | 1 | 1 | 0.001941748 | 0.63248 | 1 |
| MamGypL | 0 | 1 | 0 | 0.000948767 | 1 | 1 |
| LTR29 | 0 | 1 | 0 | 0.001424501 | 1 | 1 |
| MamGypL | 1 | 1 | 1 | 0.000823723 | 0.63227 | 1 |
| Charlie10a | 0 | 1 | 0 | 0.003322259 | 1 | 1 |
| LTR86C | 0 | 1 | 0 | 0.003508772 | 1 | 1 |
| U5 | 0 | 1 | 0 | 0.004651163 | 1 | 1 |
| L1M | 0 | 1 | 0 | 0.001004016 | 1 | 1 |
| Charlie2a | 1 | 3 | 0.333333333 | 0.001449976 | 0.95032 | 1 |
| SVA_F | 0 | 5 | 0 | 0.004878049 | 1 | 1 |
| MER104 | 3 | 2 | 1.5 | 0.000907441 | 0.32332 | 1 |
| Kanga11a | 2 | 1 | 2 | 0.001129944 | 0.26424 | 1 |
| MER5C1 | 1 | 1 | 1 | 0.001623377 | 0.63242 | 1 |
| Charlie23a | 0 | 1 | 0 | 0.000672948 | 1 | 1 |
| Tigger9a | 1 | 1 | 1 | 0.00097371 | 0.6323 | 1 |
| L1MCb | 2 | 4 | 0.5 | 0.001870033 | 0.90863 | 1 |
| LTR47A | 0 | 1 | 0 | 0.001103753 | 1 | 1 |
| MER9a3 | 0 | 2 | 0 | 0.003448276 | 1 | 1 |
| LTR82A | 0 | 1 | 0 | 0.000959693 | 1 | 1 |
| LTR21A | 0 | 1 | 0 | 0.010752688 | 1 | 1 |
| LTR77 | 1 | 1 | 1 | 0.006756757 | 0.63337 | 1 |
| LTR61 | 1 | 1 | 1 | 0.005181347 | 0.63308 | 1 |
| MER106B | 0 | 2 | 0 | 0.00241838 | 1 | 1 |
| Charlie7 | 6 | 5 | 1.2 | 0.001353546 | 0.38404 | 1 |
| LTR56 | 0 | 2 | 0 | 0.002040816 | 1 | 1 |
| MER4C | 3 | 2 | 1.5 | 0.001592357 | 0.32332 | 1 |
| MER41E | 1 | 1 | 1 | 0.002785515 | 0.63263 | 1 |
| AluYf5 | 0 | 1 | 0 | 0.005555556 | 1 | 1 |
| HERVFH19 | 0 | 1 | 0 | 0.008547009 | 1 | 1 |

| | | | | | | |
|---|---|---|---|---|---|---|
| MER110A | 1 | 1 | 1 | 0.001572327 | 0.63241 | 1 |
| AluYd8 | 0 | 1 | 0 | 0.004444444 | 1 | 1 |
| MER11D | 0 | 1 | 0 | 0.003846154 | 1 | 1 |
| MamRep4 | 0 | 1 | 0 | 0.002087683 | 1 | 1 |
| MLT2B5 | 1 | 1 | 1 | 0.003030303 | 0.63268 | 1 |
| LTR85b | 0 | 1 | 0 | 0.000851064 | 1 | 1 |
| LTR60 | 1 | 1 | 1 | 0.003649635 | 0.63279 | 1 |
| LTR2 | 0 | 3 | 0 | 0.003382187 | 1 | 1 |
| MER57C1 | 0 | 1 | 0 | 0.005263158 | 1 | 1 |
| MER21-int | 1 | 3 | 0.333333333 | 0.002747253 | 0.95042 | 1 |
| Tigger4 | 3 | 3 | 1 | 0.002061856 | 0.57704 | 1 |
| MER47C | 0 | 1 | 0 | 0.001303781 | 1 | 1 |
| ORSL-2a | 0 | 1 | 0 | 0.002747253 | 1 | 1 |
| 7SK | 1 | 1 | 1 | 0.001371742 | 0.63237 | 1 |
| LTR1D | 0 | 3 | 0 | 0.003018109 | 1 | 1 |
| LTR51 | 1 | 1 | 1 | 0.002392344 | 0.63256 | 1 |
| Kanga1c | 0 | 1 | 0 | 0.00120919 | 1 | 1 |
| X7D_LINE | 0 | 1 | 0 | 0.017857143 | 1 | 1 |
| Tigger10 | 1 | 1 | 1 | 0.000976563 | 0.6323 | 1 |
| MER106A | 0 | 1 | 0 | 0.001338688 | 1 | 1 |
| LTR84a | 0 | 1 | 0 | 0.001506024 | 1 | 1 |
| Tigger16b | 0 | 1 | 0 | 0.000951475 | 1 | 1 |
| AmnSINE2 | 0 | 1 | 0 | 0.006711409 | 1 | 1 |
| UCON14 | 0 | 1 | 0 | 0.010204082 | 1 | 1 |
| UCON29 | 1 | 1 | 1 | 0.004149378 | 0.63289 | 1 |
| Charlie13a | 2 | 1 | 2 | 0.001801802 | 0.26424 | 1 |
| Zaphod2 | 0 | 1 | 0 | 0.001360544 | 1 | 1 |
| MER2B | 0 | 2 | 0 | 0.001250782 | 1 | 1 |
| AluYh9 | 0 | 1 | 0 | 0.003676471 | 1 | 1 |
| SVA_C | 0 | 3 | 0 | 0.01 | 1 | 1 |
| LTR43B | 1 | 1 | 1 | 0.004310345 | 0.63291 | 1 |
| LTR47B | 2 | 1 | 2 | 0.002421308 | 0.26424 | 1 |
| HY3 | 0 | 1 | 0 | 0.001834862 | 1 | 1 |
| HSMAR1 | 0 | 3 | 0 | 0.004092769 | 1 | 1 |
| L1P4a | 0 | 3 | 0 | 0.005093379 | 1 | 1 |
| LTR75B | 0 | 1 | 0 | 0.004854369 | 1 | 1 |
| SVA_B | 0 | 3 | 0 | 0.006302521 | 1 | 1 |
| MER67B | 0 | 1 | 0 | 0.002066116 | 1 | 1 |
| U3 | 0 | 1 | 0 | 0.004366812 | 1 | 1 |
| MER97c | 0 | 1 | 0 | 0.004608295 | 1 | 1 |
| MLT2E | 0 | 1 | 0 | 0.002557545 | 1 | 1 |
| LTR65 | 0 | 1 | 0 | 0.002178649 | 1 | 1 |
| HSMAR2 | 1 | 6 | 0.166666667 | 0.003380282 | 0.99755 | 1 |
| Tigger8 | 1 | 1 | 1 | 0.001066098 | 0.63232 | 1 |
| MER52-int | 0 | 2 | 0 | 0.004065041 | 1 | 1 |
| HERV4_I-ir | 0 | 1 | 0 | 0.003021148 | 1 | 1 |
| MER68-int | 1 | 1 | 1 | 0.002659574 | 0.63261 | 1 |

| | | | | | | |
|---|---|---|---|---|---|---|
| MER70C | 0 | 1 | 0 | 0.003636364 | 1 | 1 |
| LTR10F | 2 | 1 | 2 | 0.002237136 | 0.26424 | 1 |
| MER6 | 0 | 2 | 0 | 0.002361275 | 1 | 1 |
| U13 | 1 | 1 | 1 | 0.005649718 | 0.63316 | 1 |
| Tigger6b | 1 | 1 | 1 | 0.003389831 | 0.63274 | 1 |
| PRIMA4-in | 1 | 2 | 0.5 | 0.002873563 | 0.86505 | 1 |
| HERVK-int | 0 | 3 | 0 | 0.011764706 | 1 | 1 |
| L1M1 | 10 | 36 | 0.277777778 | 0.003775168 | 1 | 1 |
| MER41G | 0 | 1 | 0 | 0.008849558 | 1 | 1 |
| LTR45B | 0 | 1 | 0 | 0.002252252 | 1 | 1 |
| MER57A-ir | 1 | 5 | 0.2 | 0.002634352 | 0.99331 | 1 |
| LTR45 | 0 | 1 | 0 | 0.005649718 | 1 | 1 |
| Tigger6a | 4 | 2 | 2 | 0.002580645 | 0.14264 | 1 |
| MER101-ir | 1 | 3 | 0.333333333 | 0.002375297 | 0.95039 | 1 |
| LTR7Y | 0 | 1 | 0 | 0.004016064 | 1 | 1 |
| L1M3f | 2 | 3 | 0.666666667 | 0.004273504 | 0.80149 | 1 |
| HERV3-int | 1 | 3 | 0.333333333 | 0.009803922 | 0.95094 | 1 |
| GSATII | 0 | 1 | 0 | 0.005263158 | 1 | 1 |
| HERVE-int | 0 | 4 | 0 | 0.015625 | 1 | 1 |
| GSATX | 0 | 1 | 0 | 0.017857143 | 1 | 1 |
| LTR57 | 0 | 1 | 0 | 0.003745318 | 1 | 1 |
| PRIMAX-in | 2 | 1 | 2 | 0.002680965 | 0.26424 | 1 |
| MER110-ir | 0 | 1 | 0 | 0.00257732 | 1 | 1 |
| LTR12D | 1 | 3 | 0.333333333 | 0.006134969 | 0.95067 | 1 |
| LOR1b | 1 | 2 | 0.5 | 0.001813237 | 0.86491 | 1 |
| MER30B | 0 | 1 | 0 | 0.004405286 | 1 | 1 |
| LTR88a | 1 | 1 | 1 | 0.001090513 | 0.63232 | 1 |
| MER6B | 2 | 1 | 2 | 0.001472754 | 0.26424 | 1 |
| LTR10B | 0 | 1 | 0 | 0.012658228 | 1 | 1 |
| LTR14A | 0 | 1 | 0 | 0.007633588 | 1 | 1 |
| LTR22B | 0 | 1 | 0 | 0.004291845 | 1 | 1 |
| MER6C | 0 | 1 | 0 | 0.006289308 | 1 | 1 |
| LTR22A | 1 | 1 | 1 | 0.005291005 | 0.6331 | 1 |
| MST-int | 0 | 1 | 0 | 0.002785515 | 1 | 1 |
| Ricksha | 0 | 1 | 0 | 0.005988024 | 1 | 1 |
| Ricksha_a | 0 | 1 | 0 | 0.015873016 | 1 | 1 |
| LTR80A | 1 | 1 | 1 | 0.002544529 | 0.63259 | 1 |
| MER113B | 0 | 1 | 0 | 0.001766784 | 1 | 1 |
| LTR86B1 | 0 | 1 | 0 | 0.003355705 | 1 | 1 |
| MamGypL | 1 | 1 | 1 | 0.000877963 | 0.63228 | 1 |
| LTR82B | 0 | 1 | 0 | 0.000863558 | 1 | 1 |
| LTR88b | 4 | 2 | 2 | 0.001215805 | 0.14277 | 1 |
| Helitron2N | 0 | 1 | 0 | 0.002293578 | 1 | 1 |
| MLT1F-int | 0 | 1 | 0 | 0.001953125 | 1 | 1 |
| LTR55 | 0 | 1 | 0 | 0.002457002 | 1 | 1 |
| L1HS | 0 | 13 | 0 | 0.008419689 | 1 | 1 |
| MER6A | 1 | 2 | 0.5 | 0.001960784 | 0.86493 | 1 |

| | | | | | | |
|---|---|---|---|---|---|---|
| MLT1I-int | 1 | 1 | 1 | 0.00125 | 0.63235 | 1 |
| Tigger12 | 0 | 1 | 0 | 0.002159827 | 1 | 1 |
| HERVL74-i | 0 | 2 | 0 | 0.003338898 | 1 | 1 |
| MER135 | 0 | 1 | 0 | 0.001221001 | 1 | 1 |
| LTR32 | 0 | 2 | 0 | 0.001527884 | 1 | 1 |
| X1_LINE | 0 | 1 | 0 | 0.011764706 | 1 | 1 |
| LTR16B | 1 | 1 | 1 | 0.001445087 | 0.63239 | 1 |
| MER127 | 1 | 1 | 1 | 0.004739336 | 0.63299 | 1 |
| LTR10D | 0 | 1 | 0 | 0.005235602 | 1 | 1 |
| PRIMA4_L | 0 | 1 | 0 | 0.002777778 | 1 | 1 |
| MamGypL | 1 | 1 | 1 | 0.001331558 | 0.63237 | 1 |
| LTR81A | 0 | 1 | 0 | 0.001029866 | 1 | 1 |
| DNA1_Ma | 0 | 1 | 0 | 0.005847953 | 1 | 1 |
| U7 | 0 | 1 | 0 | 0.004048583 | 1 | 1 |
| MLT1J1-in | 2 | 1 | 2 | 0.002016129 | 0.26424 | 1 |
| MER129 | 0 | 1 | 0 | 0.008196721 | 1 | 1 |
| MERX | 1 | 1 | 1 | 0.002298851 | 0.63254 | 1 |
| LTR4 | 0 | 1 | 0 | 0.00990099 | 1 | 1 |
| MER87 | 0 | 1 | 0 | 0.004405286 | 1 | 1 |
| LTR49 | 0 | 2 | 0 | 0.001386001 | 1 | 1 |
| MER51D | 1 | 1 | 1 | 0.006451613 | 0.63331 | 1 |
| PRIMA41-i | 2 | 2 | 1 | 0.004545455 | 0.59461 | 1 |
| LTR39-int | 0 | 1 | 0 | 0.00617284 | 1 | 1 |
| LTR54B | 1 | 1 | 1 | 0.001182033 | 0.63234 | 1 |
| LTR76 | 0 | 1 | 0 | 0.005319149 | 1 | 1 |
| MamGypL | 0 | 1 | 0 | 0.001121076 | 1 | 1 |
| MER50C | 0 | 1 | 0 | 0.008474576 | 1 | 1 |
| PABL_B | 0 | 1 | 0 | 0.00265252 | 1 | 1 |
| LTR15 | 0 | 1 | 0 | 0.003759398 | 1 | 1 |
| LTR38B | 0 | 1 | 0 | 0.004366812 | 1 | 1 |
| HERV-Fc2_ | 0 | 1 | 0 | 0.25 | 1 | 1 |
| MER61-int | 0 | 4 | 0 | 0.003469211 | 1 | 1 |
| Helitron1N | 0 | 1 | 0 | 0.00308642 | 1 | 1 |
| Helitron1N | 0 | 1 | 0 | 0.002762431 | 1 | 1 |
| UCON11 | 0 | 1 | 0 | 0.022222222 | 1 | 1 |
| X8_LINE | 0 | 1 | 0 | 0.003322259 | 1 | 1 |
| MER4D0 | 0 | 1 | 0 | 0.002309469 | 1 | 1 |
| HUERS-P3- | 1 | 3 | 0.333333333 | 0.004166667 | 0.95052 | 1 |
| LTR19C | 0 | 1 | 0 | 0.004424779 | 1 | 1 |
| HERV35I-in | 0 | 2 | 0 | 0.003590664 | 1 | 1 |
| U4 | 0 | 1 | 0 | 0.00621118 | 1 | 1 |
| LTR28 | 1 | 1 | 1 | 0.001941748 | 0.63248 | 1 |
| Charlie14a | 0 | 1 | 0 | 0.001908397 | 1 | 1 |
| MamRep1 | 0 | 1 | 0 | 0.001457726 | 1 | 1 |
| LTR14 | 0 | 1 | 0 | 0.010309278 | 1 | 1 |
| HERVK14C | 0 | 1 | 0 | 0.012658228 | 1 | 1 |
| MER66A | 0 | 1 | 0 | 0.00304878 | 1 | 1 |

| | | | | | | |
|---|---|---|---|---|---|---|
| MamGyp-i | 1 | 1 | 1 | 0.002923977 | 0.63266 | 1 |
| MLT1A1-ir | 1 | 1 | 1 | 0.002197802 | 0.63253 | 1 |
| LTR46-int | 0 | 1 | 0 | 0.012987013 | 1 | 1 |
| L1M2a | 0 | 2 | 0 | 0.003745318 | 1 | 1 |
| MER4-int | 3 | 9 | 0.333333333 | 0.003079028 | 0.99382 | 1 |
| L1P4e | 0 | 1 | 0 | 0.004975124 | 1 | 1 |
| UCON21 | 0 | 1 | 0 | 0.035714286 | 1 | 1 |
| MLT1A-int | 0 | 1 | 0 | 0.002222222 | 1 | 1 |
| Eulor1 | 0 | 1 | 0 | 0.011627907 | 1 | 1 |
| LTR7C | 1 | 1 | 1 | 0.003144654 | 0.6327 | 1 |
| MER73 | 0 | 1 | 0 | 0.003571429 | 1 | 1 |
| L1PBa | 4 | 10 | 0.4 | 0.004572474 | 0.98978 | 1 |
| LTR48 | 1 | 2 | 0.5 | 0.002051282 | 0.86494 | 1 |
| MER41-int | 0 | 7 | 0 | 0.004076878 | 1 | 1 |
| MLT1E3-in | 0 | 1 | 0 | 0.012048193 | 1 | 1 |
| PABL_A | 0 | 2 | 0 | 0.003824092 | 1 | 1 |
| MLT1K-int | 0 | 1 | 0 | 0.005025126 | 1 | 1 |
| MER57-int | 0 | 4 | 0 | 0.003125 | 1 | 1 |
| X6A_LINE | 0 | 1 | 0 | 0.003649635 | 1 | 1 |
| MER50-int | 1 | 2 | 0.5 | 0.003883495 | 0.86519 | 1 |
| PABL_B-in | 1 | 1 | 1 | 0.004291845 | 0.63291 | 1 |
| Charlie10b | 0 | 1 | 0 | 0.003676471 | 1 | 1 |
| UCON4 | 0 | 1 | 0 | 0.008130081 | 1 | 1 |
| MER66D | 1 | 1 | 1 | 0.004115226 | 0.63288 | 1 |
| HY4 | 0 | 1 | 0 | 0.00617284 | 1 | 1 |
| LTR87 | 0 | 1 | 0 | 0.001430615 | 1 | 1 |
| HERV1_LTI | 0 | 1 | 0 | 0.0625 | 1 | 1 |
| LTR80B | 1 | 1 | 1 | 0.001956947 | 0.63248 | 1 |
| AluYk12 | 0 | 1 | 0 | 0.004464286 | 1 | 1 |
| MLT1E2-in | 0 | 1 | 0 | 0.007518797 | 1 | 1 |
| MER41C | 2 | 2 | 1 | 0.002369668 | 0.59432 | 1 |
| L1M2b | 0 | 1 | 0 | 0.004166667 | 1 | 1 |
| MSTD-int | 0 | 2 | 0 | 0.002710027 | 1 | 1 |
| LTR3B | 0 | 1 | 0 | 0.010869565 | 1 | 1 |
| Looper | 2 | 1 | 2 | 0.001869159 | 0.26424 | 1 |
| Eulor12 | 0 | 1 | 0 | 0.016666667 | 1 | 1 |
| LTR5 | 0 | 1 | 0 | 0.05 | 1 | 1 |
| LTR6A | 0 | 1 | 0 | 0.003472222 | 1 | 1 |
| MLT1F1-in | 0 | 1 | 0 | 0.00621118 | 1 | 1 |
| MER51C | 0 | 1 | 0 | 0.005586592 | 1 | 1 |
| LTR35A | 0 | 1 | 0 | 0.011111111 | 1 | 1 |
| Charlie4 | 1 | 1 | 1 | 0.008264463 | 0.63365 | 1 |
| MER97b | 1 | 1 | 1 | 0.005494505 | 0.63313 | 1 |
| Eulor9A | 0 | 1 | 0 | 0.008130081 | 1 | 1 |
| MLT1J2-in | 0 | 1 | 0 | 0.001686341 | 1 | 1 |
| MER34D | 0 | 1 | 0 | 0.004 | 1 | 1 |
| MER51-int | 0 | 2 | 0 | 0.003584229 | 1 | 1 |

| | | | | | | |
|---|---|---|---|---|---|---|
| MER65C | 2 | 1 | 2 | 0.001464129 | 0.26424 | 1 |
| LTR12B | 0 | 1 | 0 | 0.004739336 | 1 | 1 |
| MER57D | 1 | 1 | 1 | 0.002183406 | 0.63252 | 1 |
| U17 | 0 | 1 | 0 | 0.111111111 | 1 | 1 |
| LTR35 | 0 | 1 | 0 | 0.008474576 | 1 | 1 |
| MER50B | 0 | 2 | 0 | 0.002890173 | 1 | 1 |
| MSTB1-int | 0 | 1 | 0 | 0.003875969 | 1 | 1 |
| UCON19 | 0 | 1 | 0 | 0.035714286 | 1 | 1 |
| L1MDb | 0 | 3 | 0 | 0.002767528 | 1 | 1 |
| UCON7 | 0 | 1 | 0 | 0.007874016 | 1 | 1 |
| MLT1G-int | 0 | 1 | 0 | 0.005524862 | 1 | 1 |
| LTR31 | 0 | 1 | 0 | 0.004149378 | 1 | 1 |
| L1M3c | 0 | 2 | 0 | 0.00181653 | 1 | 1 |
| HERVK22-i | 5 | 3 | 1.666666667 | 0.007025761 | 0.18414 | 1 |
| LTR17 | 0 | 3 | 0 | 0.003575685 | 1 | 1 |
| MER83B | 0 | 1 | 0 | 0.006944444 | 1 | 1 |
| PrimLTR79 | 0 | 1 | 0 | 0.007092199 | 1 | 1 |
| SVA_A | 0 | 2 | 0 | 0.007434944 | 1 | 1 |
| MER74C | 0 | 1 | 0 | 0.004291845 | 1 | 1 |
| MER99 | 1 | 1 | 1 | 0.003378378 | 0.63274 | 1 |
| UCON26 | 2 | 1 | 2 | 0.004854369 | 0.26424 | 1 |
| LTR88c | 2 | 1 | 2 | 0.000958773 | 0.26424 | 1 |
| FordPrefec | 2 | 1 | 2 | 0.00166113 | 0.26424 | 1 |
| SSU-rRNA_ | 2 | 1 | 2 | 0.0125 | 0.26424 | 1 |
| HUERS-P3l | 0 | 2 | 0 | 0.004366812 | 1 | 1 |
| LTR6B | 0 | 1 | 0 | 0.006493506 | 1 | 1 |
| HERVS71-i | 0 | 1 | 0 | 0.005714286 | 1 | 1 |
| HERVK11-i | 0 | 2 | 0 | 0.006802721 | 1 | 1 |
| MLT1F2-in | 1 | 1 | 1 | 0.003333333 | 0.63273 | 1 |
| X5A_LINE | 0 | 1 | 0 | 0.00729927 | 1 | 1 |
| LTR90A | 2 | 1 | 2 | 0.002457002 | 0.26424 | 1 |
| THE1-int | 0 | 1 | 0 | 0.001531394 | 1 | 1 |
| LTR86B2 | 0 | 1 | 0 | 0.003717472 | 1 | 1 |
| LTR69 | 0 | 1 | 0 | 0.007042254 | 1 | 1 |
| ALINE | 0 | 1 | 0 | 0.003623188 | 1 | 1 |
| LTR72B | 2 | 1 | 2 | 0.005882353 | 0.26424 | 1 |
| L1MEg2 | 0 | 1 | 0 | 0.001269036 | 1 | 1 |
| AluYc5 | 0 | 1 | 0 | 0.022222222 | 1 | 1 |
| HY5 | 0 | 1 | 0 | 0.045454545 | 1 | 1 |
| MER89-int | 1 | 1 | 1 | 0.001424501 | 0.63238 | 1 |
| L1MEg1 | 2 | 1 | 2 | 0.001068376 | 0.26424 | 1 |
| MamGypL | 1 | 1 | 1 | 0.00104712 | 0.63231 | 1 |
| L1M3de | 0 | 1 | 0 | 0.002164502 | 1 | 1 |
| UCON12A | 0 | 1 | 0 | 0.038461538 | 1 | 1 |
| Eulor2B | 0 | 1 | 0 | 0.01754386 | 1 | 1 |
| BC200 | 0 | 1 | 0 | 0.005988024 | 1 | 1 |
| LTR43 | 2 | 1 | 2 | 0.002493766 | 0.26424 | 1 |

| | | | | | | |
|---|---|---|---|---|---|---|
| PABL_A-in | 1 | 2 | 0.5 | 0.004750594 | 0.86531 | 1 |
| AluYb9 | 0 | 1 | 0 | 0.003058104 | 1 | 1 |
| MLT1N2-ir | 0 | 1 | 0 | 0.013157895 | 1 | 1 |
| MER70-int | 0 | 1 | 0 | 0.003875969 | 1 | 1 |
| L1M3d | 0 | 1 | 0 | 0.001855288 | 1 | 1 |
| LTR5A | 0 | 2 | 0 | 0.00754717 | 1 | 1 |
| LTR34 | 1 | 1 | 1 | 0.002590674 | 0.6326 | 1 |
| HERV17-in | 2 | 4 | 0.5 | 0.004561003 | 0.90892 | 1 |
| L1PBb | 2 | 1 | 2 | 0.003787879 | 0.26424 | 1 |
| SUBTEL_sa | 0 | 1 | 0 | 0.029411765 | 1 | 1 |
| MER76 | 1 | 1 | 1 | 0.001392758 | 0.63238 | 1 |
| MER97a | 0 | 1 | 0 | 0.003610108 | 1 | 1 |
| MLT1H2-ir | 1 | 1 | 1 | 0.002857143 | 0.63265 | 1 |
| LTR27B | 0 | 1 | 0 | 0.001745201 | 1 | 1 |
| LTR19-int | 0 | 1 | 0 | 0.004975124 | 1 | 1 |
| Merlin1_H | 0 | 1 | 0 | 0.01754386 | 1 | 1 |
| MSTC-int | 0 | 1 | 0 | 0.004926108 | 1 | 1 |
| MER83B-ir | 1 | 1 | 1 | 0.003846154 | 0.63283 | 1 |
| UCON5 | 0 | 1 | 0 | 0.009803922 | 1 | 1 |
| MER61F | 0 | 1 | 0 | 0.00621118 | 1 | 1 |
| GSAT | 0 | 1 | 0 | 0.014925373 | 1 | 1 |
| MLT1-int | 0 | 1 | 0 | 0.004405286 | 1 | 1 |
| Eulor9B | 0 | 1 | 0 | 0.04 | 1 | 1 |
| L1M3a | 0 | 2 | 0 | 0.003139717 | 1 | 1 |
| L1P3b | 0 | 1 | 0 | 0.012195122 | 1 | 1 |
| UCON31 | 0 | 1 | 0 | 0.009090909 | 1 | 1 |
| MER97d | 0 | 1 | 0 | 0.009803922 | 1 | 1 |
| LTR68 | 0 | 1 | 0 | 0.002816901 | 1 | 1 |
| LTR12_ | 0 | 2 | 0 | 0.003603604 | 1 | 1 |
| L1P4d | 0 | 1 | 0 | 0.00625 | 1 | 1 |
| X6B_LINE | 0 | 1 | 0 | 0.002320186 | 1 | 1 |
| UCON6 | 0 | 1 | 0 | 0.011764706 | 1 | 1 |
| LTR40A1 | 1 | 1 | 1 | 0.001760563 | 0.63244 | 1 |
| UCON28b | 0 | 1 | 0 | 0.012987013 | 1 | 1 |
| L1MEa | 2 | 1 | 2 | 0.002645503 | 0.26424 | 1 |
| L1M2a1 | 0 | 1 | 0 | 0.008064516 | 1 | 1 |
| MER132 | 0 | 1 | 0 | 0.027027027 | 1 | 1 |
| Eulor4 | 0 | 1 | 0 | 0.032258065 | 1 | 1 |
| MER125 | 0 | 1 | 0 | 0.007142857 | 1 | 1 |
| MER68B | 1 | 1 | 1 | 0.002222222 | 0.63253 | 1 |
| UCON8 | 0 | 1 | 0 | 0.01010101 | 1 | 1 |
| MER123 | 0 | 1 | 0 | 0.024390244 | 1 | 1 |
| MER95 | 0 | 1 | 0 | 0.006410256 | 1 | 1 |
| Eulor6A | 0 | 1 | 0 | 0.033333333 | 1 | 1 |
| Eulor5A | 0 | 1 | 0 | 0.007874016 | 1 | 1 |
| MER61C | 0 | 1 | 0 | 0.003184713 | 1 | 1 |
| HERVK14-i | 1 | 2 | 0.5 | 0.004975124 | 0.86534 | 1 |

| | | | | | | |
|---|---|---|---|---|---|---|
| MER61E | 0 | 1 | 0 | 0.002985075 | 1 | 1 |
| MER130 | 0 | 1 | 0 | 0.010309278 | 1 | 1 |
| MER133A | 0 | 1 | 0 | 0.018518519 | 1 | 1 |
| LTR18A | 0 | 1 | 0 | 0.003861004 | 1 | 1 |
| HERVL18-i | 0 | 2 | 0 | 0.003690037 | 1 | 1 |
| MER72B | 0 | 1 | 0 | 0.005524862 | 1 | 1 |
| Eulor9C | 0 | 1 | 0 | 0.009345794 | 1 | 1 |
| L1M3b | 0 | 2 | 0 | 0.003590664 | 1 | 1 |
| LTR25-int | 0 | 3 | 0 | 0.005093379 | 1 | 1 |
| HUERS-P2- | 0 | 1 | 0 | 0.003937008 | 1 | 1 |
| MER67D | 0 | 1 | 0 | 0.000987167 | 1 | 1 |
| L1P4b | 0 | 1 | 0 | 0.006578947 | 1 | 1 |
| MER83C | 0 | 1 | 0 | 0.006493506 | 1 | 1 |
| HERVP71A | 1 | 1 | 1 | 0.003984064 | 0.63285 | 1 |
| Charlie3 | 1 | 2 | 0.5 | 0.005847953 | 0.86546 | 1 |
| Eulor3 | 1 | 1 | 1 | 0.027777778 | 0.63729 | 1 |
| UCON18 | 0 | 1 | 0 | 0.055555556 | 1 | 1 |
| MER101B | 1 | 1 | 1 | 0.003861004 | 0.63283 | 1 |
| Eulor6D | 2 | 1 | 2 | 0.013513514 | 0.26424 | 1 |
| HERVL32-i | 0 | 1 | 0 | 0.008695652 | 1 | 1 |
| LTR58 | 0 | 1 | 0 | 0.013157895 | 1 | 1 |
| Eulor8 | 0 | 1 | 0 | 0.006993007 | 1 | 1 |
| Ricksha_b | 1 | 1 | 1 | 0.008333333 | 0.63366 | 1 |
| UCON25 | 0 | 1 | 0 | 0.023255814 | 1 | 1 |
| MER92C | 2 | 1 | 2 | 0.003012048 | 0.26424 | 1 |
| UCON13 | 0 | 1 | 0 | 0.019607843 | 1 | 1 |
| MLT1E-int | 0 | 1 | 0 | 0.055555556 | 1 | 1 |
| MER57C2 | 0 | 1 | 0 | 0.002141328 | 1 | 1 |
| UCON28c | 0 | 1 | 0 | 0.015384615 | 1 | 1 |
| Eulor6C | 0 | 1 | 0 | 0.045454545 | 1 | 1 |
| X2_LINE | 0 | 1 | 0 | 0.015384615 | 1 | 1 |
| LTR35B | 0 | 1 | 0 | 0.004830918 | 1 | 1 |
| L1PBa1 | 0 | 3 | 0 | 0.007389163 | 1 | 1 |
| UCON2 | 0 | 1 | 0 | 0.008403361 | 1 | 1 |
| UCON15 | 0 | 1 | 0 | 0.022727273 | 1 | 1 |
| UCON27 | 0 | 1 | 0 | 0.009708738 | 1 | 1 |
| UCON10 | 0 | 1 | 0 | 0.015625 | 1 | 1 |
| Eulor2C | 0 | 1 | 0 | 0.026315789 | 1 | 1 |
| Eulor11 | 0 | 1 | 0 | 0.012345679 | 1 | 1 |
| ALR/Alpha | 11 | 10 | 1.1 | 0.007686395 | 0.41696 | 1 |
| MLT1G3-ir | 0 | 1 | 0 | 0.004166667 | 1 | 1 |
| HERVH48- | 0 | 1 | 0 | 0.008474576 | 1 | 1 |
| LTR30 | 0 | 1 | 0 | 0.006896552 | 1 | 1 |
| LTR52-int | 0 | 1 | 0 | 0.003731343 | 1 | 1 |
| UCON28a | 0 | 1 | 0 | 0.007751938 | 1 | 1 |
| Eulor7 | 0 | 1 | 0 | 0.090909091 | 1 | 1 |
| MSTB2-int | 0 | 1 | 0 | 0.010989011 | 1 | 1 |

| | | | | | | |
|---|---|---|---|---|---|---|
| MER133B | 0 | 1 | 0 | 0.024390244 | 1 | 1 |
| HERVKC4-i | 0 | 1 | 0 | 0.02173913 | 1 | 1 |
| UCON1 | 0 | 1 | 0 | 0.034482759 | 1 | 1 |
| Eulor6B | 0 | 1 | 0 | 0.016949153 | 1 | 1 |
| LTR11 | 0 | 1 | 0 | 0.0625 | 1 | 1 |
| HERVE_a-i | 1 | 3 | 0.333333333 | 0.015463918 | 0.95137 | 1 |
| LTR22 | 0 | 1 | 0 | 0.009708738 | 1 | 1 |
| MER134 | 0 | 1 | 0 | 0.014925373 | 1 | 1 |
| MER87B | 0 | 1 | 0 | 0.004098361 | 1 | 1 |
| Charlie11 | 0 | 1 | 0 | 0.009259259 | 1 | 1 |
| MER88 | 0 | 1 | 0 | 0.008849558 | 1 | 1 |
| UCON9 | 0 | 1 | 0 | 0.020408163 | 1 | 1 |
| LTR25 | 2 | 1 | 2 | 0.003546099 | 0.26424 | 1 |
| LTR3 | 0 | 1 | 0 | 0.010752688 | 1 | 1 |
| HERVK3-in | 0 | 3 | 0 | 0.009036145 | 1 | 1 |
| L1P | 0 | 1 | 0 | 0.006289308 | 1 | 1 |
| MER83A-ir | 0 | 1 | 0 | 0.009345794 | 1 | 1 |
| MER92A | 1 | 1 | 1 | 0.006578947 | 0.63333 | 1 |
| Charlie6 | 0 | 1 | 0 | 0.003846154 | 1 | 1 |
| LTR72 | 2 | 1 | 2 | 0.006535948 | 0.26424 | 1 |
| CER | 1 | 1 | 1 | 0.013888889 | 0.63469 | 1 |
| HSAT5 | 0 | 1 | 0 | 0.003846154 | 1 | 1 |
| SATR1 | 0 | 4 | 0 | 0.007352941 | 1 | 1 |
| ACRO1 | 0 | 1 | 0 | 0.016393443 | 1 | 1 |
| MLT1E1-in | 0 | 1 | 0 | 0.022222222 | 1 | 1 |
| UCON23 | 0 | 1 | 0 | 0.032258065 | 1 | 1 |
| Tigger2b | 0 | 1 | 0 | 0.009090909 | 1 | 1 |
| MER76-int | 0 | 1 | 0 | 0.006666667 | 1 | 1 |
| LTR14C | 0 | 1 | 0 | 0.00729927 | 1 | 1 |
| Eulor5B | 0 | 1 | 0 | 0.023255814 | 1 | 1 |
| LTR38C | 0 | 1 | 0 | 0.008064516 | 1 | 1 |
| HERV1_LTI | 1 | 1 | 1 | 0.037037037 | 0.63904 | 1 |
| MLT1L-int | 0 | 1 | 0 | 0.009259259 | 1 | 1 |
| UCON24 | 0 | 1 | 0 | 0.045454545 | 1 | 1 |
| LTR10G | 0 | 1 | 0 | 0.008403361 | 1 | 1 |
| LTR59 | 0 | 1 | 0 | 0.008196721 | 1 | 1 |
| MER136 | 0 | 1 | 0 | 0.035714286 | 1 | 1 |
| UCON17 | 0 | 1 | 0 | 0.03030303 | 1 | 1 |
| SATR2 | 0 | 3 | 0 | 0.011450382 | 1 | 1 |
| MER57E3 | 0 | 1 | 0 | 0.004098361 | 1 | 1 |
| LTR70 | 0 | 1 | 0 | 0.006944444 | 1 | 1 |
| HSAT4 | 0 | 1 | 0 | 0.01010101 | 1 | 1 |
| Eulor6E | 1 | 1 | 1 | 0.032258065 | 0.63814 | 1 |
| U8 | 0 | 1 | 0 | 0.034482759 | 1 | 1 |
| L1P4c | 0 | 1 | 0 | 0.022222222 | 1 | 1 |
| HERVK11D | 0 | 1 | 0 | 0.02173913 | 1 | 1 |
| UCON12 | 0 | 1 | 0 | 0.022222222 | 1 | 1 |

| | | | | | | |
|---|---|---|---|---|---|---|
| REP522 | 0 | 3 | 0 | 0.012295082 | 1 | 1 |
| HERV1_I-ir | 0 | 1 | 0 | 0.020408163 | 1 | 1 |
| HSATI | 0 | 1 | 0 | 0.022222222 | 1 | 1 |
| MER9B | 0 | 1 | 0 | 0.025641026 | 1 | 1 |
| Charlie1b_ | 0 | 1 | 0 | 0.5 | 1 | 1 |
| HERV30-in | 0 | 1 | 0 | 0.01369863 | 1 | 1 |
| CheshMITI | 0 | 1 | 0 | 1 | 1 | 1 |
| SST1 | 1 | 4 | 0.25 | 0.006535948 | 0.98192 | 1 |
| Tigger1a_M | 0 | 1 | 0 | 0.022727273 | 1 | 1 |
| LTR43-int | 0 | 1 | 0 | 0.006666667 | 1 | 1 |
| LSAU | 0 | 1 | 0 | 0.007751938 | 1 | 1 |
| HERV1_LTI | 0 | 1 | 0 | 0.032258065 | 1 | 1 |
| HERV-Fc1_ | 0 | 1 | 0 | 0.058823529 | 1 | 1 |
| HERV-Fc1_ | 0 | 1 | 0 | 0.333333333 | 1 | 1 |
| HERV-Fc2- | 0 | 1 | 0 | 0.333333333 | 1 | 1 |
| HERV-Fc1- | 0 | 1 | 0 | 0.142857143 | 1 | 1 |
| U14 | 0 | 1 | 0 | 0.142857143 | 1 | 1 |
| HAL1N1_M | 0 | 1 | 0 | 0.5 | 1 | 1 |
| HERV1_LTI | 0 | 1 | 0 | 0.029411765 | 1 | 1 |
| HERV15-in | 0 | 1 | 0 | 0.010752688 | 1 | 1 |
| HERV-Fc1_ | 0 | 1 | 0 | 0.2 | 1 | 1 |
| HERV1_LTI | 0 | 1 | 0 | 0.1 | 1 | 1 |
| Charlie12 | 0 | 1 | 0 | 0.045454545 | 1 | 1 |
| Cheshire_I | 0 | 1 | 0 | 0.142857143 | 1 | 1 |
| HSAT6 | 0 | 1 | 0 | 0.058823529 | 1 | 1 |
| D20S16 | 0 | 1 | 0 | 0.003937008 | 1 | 1 |
| MLT1M-in | 0 | 1 | 0 | 1 | 1 | 1 |
| SAR | 1 | 1 | 1 | 1 | 1 | 1 |
| Tigger2a_C | 0 | 1 | 0 | 0.5 | 1 | 1 |
