## Supplementary Table 3 for "Transposable elements have contributed human regulatory regions that are activated upon bacterial infection"

Table S3. Mean estimated age of accessible and inaccessible repeats
(PARs belonging to the 14 related TE families included only)

| TE.family | Age.accessible.repeats | Age.inaccessible.repeats | Pvalue |
|---|---|---|---|
| Tigger3b | 62.53846154 | 66.95185075 | 6.60E-05 |
| MER41B | 59.98955068 | 55.55942791 | 0.003597175 |
| Tigger3c | 70.99365751 | 77.24751817 | 0.006424437 |
| MLT1F | 102.3146853 | 106.2044595 | 0.018560594 |
| MER44B | 82.63914657 | 85.50909985 | 0.027565137 |
| Tigger3 | 60.04545455 | 67.15443134 | 0.027712674 |
| MER44D | 80.22727273 | 85.49771458 | 0.031627788 |
| THE1B | 56.44951822 | 57.98926654 | 0.034933169 |
| Tigger3a | 75.45454545 | 77.75395973 | 0.057736504 |
| THE1C | 53.20483314 | 55.39013411 | 0.072078777 |
| MLT1L | 129.4107744 | 125.9295953 | 0.209824478 |
| MLT1I | 126.8734644 | 127.7287996 | 0.373083138 |
| MLT1H | 118.9964581 | 119.0166263 | 0.64382778 |
| MER44C | 83.98989899 | 83.56690376 | 0.767076694 |
| MLT1K | 129.1083916 | 127.9027113 | 0.96671613 |
