## Supplementary Tables 4 and 5 for "Transposable elements have contributed human regulatory regions that are activated upon bacterial infection"

Table S4. TE families near DEG in the S12 samples

| TE.family | Observed.instances | Expected.instances | Fold.enrichment | Prob.success | P.value | Q.value |
|---|---|---|---|---|---|---|
| MIRb | 117 | 49.813 | 2.348784454 | 0.13318984 | 2.27E-19 | 3.45E-17 |
| THE1B | 57 | 15.619 | 3.64940137 | 0.076190244 | 4.96E-18 | 3.77E-16 |
| MIR3 | 71 | 28.514 | 2.49000491 | 0.144741117 | 4.93E-14 | 2.50E-12 |
| L2b | 65 | 29.436 | 2.208180459 | 0.140171429 | 2.60E-10 | 9.88E-09 |
| L2c | 70 | 33.479 | 2.090862929 | 0.124457249 | 1.26E-09 | 3.83E-08 |
| MIR | 71 | 35.006 | 2.028223733 | 0.132598485 | 3.14E-09 | 6.81E-08 |
| MIRc | 64 | 30.24 | 2.116402116 | 0.13380531 | 2.86E-09 | 6.81E-08 |
| L1MC4a | 28 | 9.664 | 2.897350993 | 0.140057971 | 5.57E-08 | 1.06E-06 |
| L2a | 59 | 29.418 | 2.005574818 | 0.121061728 | 1.23E-07 | 2.08E-06 |
| L1M5 | 23 | 7.433 | 3.094309162 | 0.119887097 | 3.40E-07 | 4.69E-06 |
| MER41B | 27 | 9.762 | 2.765826675 | 0.137492958 | 3.15E-07 | 4.69E-06 |
| MLT1K | 27 | 11.028 | 2.448313384 | 0.122533333 | 6.36E-06 | 8.06E-05 |
| MLT1L | 22 | 8.161 | 2.69574807 | 0.118275362 | 8.71E-06 | 0.000102 |
| L3 | 28 | 11.901 | 2.352743467 | 0.11901 | 1.06E-05 | 0.000111 |
| L1ME3A | 13 | 3.386 | 3.839338452 | 0.102606061 | 1.09E-05 | 0.000111 |
| L1MB5 | 22 | 9.091 | 2.4199758 | 0.151516667 | 3.60E-05 | 0.000342 |
| AluSx | 18 | 7.002 | 2.570694087 | 0.189243243 | 4.12E-05 | 0.000368 |
| L1ME1 | 23 | 9.899 | 2.323467017 | 0.111224719 | 8.50E-05 | 0.000718 |
| MER57E1 | 6 | 0.92 | 6.52173913 | 0.092 | 9.18E-05 | 0.000734 |
| L1MB2 | 34 | 17.632 | 1.92831216 | 0.119135135 | 0.00012 | 0.000887 |
| LTR26 | 7 | 1.294 | 5.409582689 | 0.086266667 | 0.000122 | 0.000887 |
| THE1C | 14 | 4.577 | 3.058772121 | 0.071515625 | 0.000143 | 0.000988 |
| ERVL-E-int | 11 | 3.207 | 3.429996882 | 0.094323529 | 0.000191 | 0.001211 |
| L1MB7 | 45 | 26.463 | 1.700487473 | 0.154754386 | 0.000184 | 0.001211 |
| LTR16C | 6 | 1.06 | 5.660377358 | 0.106 | 0.000204 | 0.001239 |
| MLT2B3 | 6 | 1.017 | 5.899705015 | 0.072642857 | 0.000265 | 0.001547 |
| L1M1 | 7 | 1.446 | 4.840940526 | 0.080333333 | 0.00031 | 0.001744 |
| L1ME4a | 14 | 5.299 | 2.642007926 | 0.139447368 | 0.000369 | 0.001991 |
| ERV3-16A3 | 8 | 1.918 | 4.17101147 | 0.079916667 | 0.00038 | 0.001991 |
| L1PB1 | 5 | 0.753 | 6.640106242 | 0.0753 | 0.000442 | 0.002238 |
| L2 | 32 | 17.363 | 1.842999482 | 0.114986755 | 0.000458 | 0.002246 |
| L1MC4 | 16 | 6.708 | 2.385211688 | 0.13416 | 0.000578 | 0.002745 |
| THE1D | 9 | 2.531 | 3.555906756 | 0.07669697 | 0.000647 | 0.00298 |
| AluSx1 | 13 | 5.277 | 2.46352094 | 0.181965517 | 0.000866 | 0.003763 |
| MER117 | 7 | 1.791 | 3.908431044 | 0.127928571 | 0.000843 | 0.003763 |
| MER20 | 12 | 4.613 | 2.601344028 | 0.121394737 | 0.001307 | 0.005517 |
| L1MB8 | 20 | 9.964 | 2.007226014 | 0.134648649 | 0.001493 | 0.006135 |
| MER44B | 19 | 9.141 | 2.078547205 | 0.096221053 | 0.001627 | 0.006509 |
| AluSz | 22 | 12.025 | 1.82952183 | 0.179477612 | 0.00243 | 0.00947 |
| L1M4 | 11 | 4.347 | 2.530480791 | 0.1242 | 0.00258 | 0.009803 |
| MLT1C | 11 | 4.392 | 2.504553734 | 0.122 | 0.002867 | 0.01063 |
| HAL1 | 18 | 9.077 | 1.983034042 | 0.129671429 | 0.002992 | 0.010829 |
| MSTB | 5 | 1.128 | 4.432624113 | 0.094 | 0.003298 | 0.01166 |
| L1MC1 | 8 | 2.656 | 3.012048193 | 0.102153846 | 0.003409 | 0.011777 |
| MLT1I | 13 | 5.654 | 2.299257163 | 0.091193548 | 0.00349 | 0.01179 |

| | | | | | | |
|---|---|---|---|---|---|---|
| MLT1G | 7 | 2.189 | 3.197807218 | 0.10945 | 0.003999 | 0.013215 |
| HERV9-int | 6 | 1.655 | 3.625377644 | 0.091944444 | 0.004233 | 0.013689 |
| L1MC3 | 6 | 1.7 | 3.529411765 | 0.085 | 0.005133 | 0.016253 |
| MER5A | 9 | 3.53 | 2.549575071 | 0.126071429 | 0.005818 | 0.018046 |
| MSTD | 7 | 2.324 | 3.012048193 | 0.105636364 | 0.005954 | 0.0181 |
| AluSq2 | 7 | 2.508 | 2.791068581 | 0.179142857 | 0.006236 | 0.018585 |
| Tigger3c | 10 | 4.299 | 2.326122354 | 0.126441176 | 0.00764 | 0.022333 |
| MLT1E2 | 4 | 0.87 | 4.597701149 | 0.087 | 0.007832 | 0.022461 |
| L1MEg | 5 | 1.377 | 3.631082062 | 0.098357143 | 0.008609 | 0.024232 |
| LTR8 | 9 | 3.78 | 2.380952381 | 0.14 | 0.008889 | 0.024567 |
| MLT1D | 5 | 1.442 | 3.46740638 | 0.110923077 | 0.010034 | 0.027234 |
| ERVL-B4-in | 6 | 1.986 | 3.021148036 | 0.104526316 | 0.010621 | 0.028324 |
| AluY | 8 | 3.316 | 2.412545235 | 0.157904762 | 0.01133 | 0.028701 |
| MLT1H | 12 | 5.792 | 2.071823204 | 0.098169492 | 0.011155 | 0.028701 |
| L1MCa | 4 | 0.945 | 4.232804233 | 0.085909091 | 0.010973 | 0.028701 |
| MER51A | 7 | 2.641 | 2.65051117 | 0.125761905 | 0.011586 | 0.028871 |
| MER103C | 5 | 1.488 | 3.360215054 | 0.106285714 | 0.011905 | 0.029188 |
| Tigger3 | 5 | 1.517 | 3.295978906 | 0.084277778 | 0.014327 | 0.034566 |
| AluJb | 30 | 20.445 | 1.467351431 | 0.185863636 | 0.016283 | 0.037055 |
| MER21C | 9 | 4.052 | 2.22112537 | 0.115771429 | 0.015886 | 0.037055 |
| MLT1J | 12 | 6.155 | 1.949634444 | 0.1231 | 0.016333 | 0.037055 |
| MER82 | 5 | 1.609 | 3.107520199 | 0.123769231 | 0.015821 | 0.037055 |
| Tigger3a | 16 | 9.157 | 1.74729715 | 0.123743243 | 0.017607 | 0.039358 |
| MLT1G3 | 5 | 1.721 | 2.905287623 | 0.114733333 | 0.022189 | 0.04888 |
| LTR33 | 5 | 1.731 | 2.888503755 | 0.101823529 | 0.023777 | 0.051629 |
| MLT1N2 | 6 | 2.353 | 2.549936252 | 0.112047619 | 0.024255 | 0.051927 |
| Plat_L3 | 5 | 1.783 | 2.804262479 | 0.127357143 | 0.024813 | 0.052383 |
| L1ME2 | 7 | 3.029 | 2.310993727 | 0.1165 | 0.02583 | 0.053784 |
| LTR71A | 4 | 1.236 | 3.236245955 | 0.112363636 | 0.027447 | 0.056379 |
| MER68 | 4 | 1.249 | 3.20256205 | 0.113545455 | 0.028419 | 0.057596 |
| AluSq | 5 | 1.924 | 2.598752599 | 0.174909091 | 0.029774 | 0.058761 |
| L1MB4 | 10 | 5.231 | 1.911680367 | 0.130775 | 0.030111 | 0.058761 |
| MER81 | 9 | 4.552 | 1.9771529 | 0.14225 | 0.03054 | 0.058761 |
| MLT1F | 13 | 7.454 | 1.744030051 | 0.128517241 | 0.030444 | 0.058761 |
| MER5A1 | 5 | 1.908 | 2.620545073 | 0.1272 | 0.033243 | 0.06208 |
| L4 | 6 | 2.523 | 2.378121284 | 0.114681818 | 0.033491 | 0.06208 |
| L1MB1 | 3 | 0.74 | 4.054054054 | 0.067272727 | 0.033379 | 0.06208 |
| AluSz6 | 9 | 4.713 | 1.909611712 | 0.174555556 | 0.034613 | 0.06238 |
| L1MEf | 5 | 1.889 | 2.646903123 | 0.09445 | 0.034884 | 0.06238 |
| MLT1A0 | 7 | 3.176 | 2.204030227 | 0.096242424 | 0.034862 | 0.06238 |
| L1MC5 | 7 | 3.301 | 2.120569524 | 0.143521739 | 0.037537 | 0.066345 |
| Tigger15a | 5 | 2.019 | 2.476473502 | 0.118764706 | 0.042899 | 0.07495 |
| MLT1G1 | 5 | 2.002 | 2.497502498 | 0.095333333 | 0.043808 | 0.075668 |
| LTR13 | 6 | 2.82 | 2.127659574 | 0.201428571 | 0.045251 | 0.077283 |
| Tigger3b | 20 | 13.465 | 1.485332343 | 0.114110169 | 0.045903 | 0.077442 |
| MamSINE1 | 4 | 1.43 | 2.797202797 | 0.11 | 0.046363 | 0.077442 |
| MLT2B1 | 3 | 0.875 | 3.428571429 | 0.0875 | 0.050341 | 0.083173 |

| | | | | | | |
|---|---|---|---|---|---|---|
| MLT1A | 4 | 1.499 | 2.66844563 | 0.099933333 | 0.055441 | 0.090614 |
| MLT1B | 4 | 1.619 | 2.470660902 | 0.107933333 | 0.070106 | 0.113363 |
| HERVL-int | 3 | 0.997 | 3.009027081 | 0.058647059 | 0.074054 | 0.118486 |
| MLT1M | 4 | 1.667 | 2.399520096 | 0.119071429 | 0.075589 | 0.119682 |
| MLT1F2 | 10 | 6.106 | 1.637733377 | 0.098483871 | 0.080523 | 0.124893 |
| MSTA | 4 | 1.668 | 2.398081535 | 0.0834 | 0.079803 | 0.124893 |
| AluSp | 6 | 3.169 | 1.893341748 | 0.176055556 | 0.081462 | 0.125073 |
| MLT1J1 | 3 | 1.067 | 2.811621368 | 0.088916667 | 0.084163 | 0.127927 |
| MER57F | 3 | 1.099 | 2.729754322 | 0.1099 | 0.088251 | 0.132813 |
| L1MD3 | 5 | 2.477 | 2.018570852 | 0.12385 | 0.0921 | 0.136385 |
| MLT2B4 | 3 | 1.115 | 2.69058296 | 0.092916667 | 0.093416 | 0.136385 |
| Tigger7 | 10 | 6.284 | 1.591343094 | 0.106508475 | 0.092754 | 0.136385 |
| MER44C | 7 | 3.96 | 1.767676768 | 0.11 | 0.094213 | 0.136385 |
| L3b | 3 | 1.154 | 2.59965338 | 0.1154 | 0.099134 | 0.142155 |
| MER63A | 3 | 1.159 | 2.588438309 | 0.1159 | 0.100153 | 0.142273 |
| MER45B | 4 | 1.89 | 2.116402116 | 0.105 | 0.112696 | 0.158012 |
| LTR16A1 | 3 | 1.216 | 2.467105263 | 0.110545455 | 0.113311 | 0.158012 |
| LTR78 | 3 | 1.285 | 2.33463035 | 0.098846154 | 0.130449 | 0.180256 |
| MLT1H1 | 5 | 2.746 | 1.820830299 | 0.105615385 | 0.132972 | 0.180463 |
| Tigger1 | 7 | 4.303 | 1.62677202 | 0.107575 | 0.132727 | 0.180463 |
| MLT1J2 | 6 | 3.586 | 1.673173452 | 0.102457143 | 0.143151 | 0.190868 |
| MLT1F1 | 7 | 4.398 | 1.59163256 | 0.125657143 | 0.142325 | 0.190868 |
| MamRep6( | 4 | 2.112 | 1.893939394 | 0.117333333 | 0.152498 | 0.201563 |
| L1MB3 | 22 | 17.707 | 1.24244649 | 0.145139344 | 0.16413 | 0.215067 |
| Charlie1b | 3 | 1.511 | 1.985440106 | 0.1511 | 0.182668 | 0.236185 |
| MER20B | 3 | 1.509 | 1.988071571 | 0.137181818 | 0.183354 | 0.236185 |
| MER5B | 3 | 1.533 | 1.956947162 | 0.12775 | 0.190592 | 0.243445 |
| L1MD2 | 5 | 3.123 | 1.601024656 | 0.107689655 | 0.196637 | 0.248292 |
| MER21B | 5 | 3.151 | 1.586797842 | 0.150047619 | 0.197654 | 0.248292 |
| THE1A | 2 | 0.974 | 2.05338809 | 0.074923077 | 0.254107 | 0.316592 |
| L1MEc | 3 | 1.842 | 1.628664495 | 0.096947368 | 0.278173 | 0.343758 |
| L1ME3C | 2 | 1.252 | 1.597444089 | 0.1252 | 0.361875 | 0.443589 |
| MARNA | 2 | 1.363 | 1.467351431 | 0.1363 | 0.404437 | 0.488657 |
| Charlie7 | 2 | 1.367 | 1.463057791 | 0.124272727 | 0.405071 | 0.488657 |
| MER44D | 4 | 3.294 | 1.214329083 | 0.094114286 | 0.42146 | 0.504425 |
| L1MA9 | 2 | 1.502 | 1.331557923 | 0.07152381 | 0.449057 | 0.533255 |
| MLT1H2 | 3 | 2.463 | 1.218026797 | 0.102625 | 0.453086 | 0.533869 |
| L1MC2 | 2 | 1.559 | 1.282873637 | 0.103933333 | 0.471764 | 0.551601 |
| L1MEd | 2 | 1.585 | 1.261829653 | 0.0990625 | 0.480112 | 0.557076 |
| AluJr | 9 | 8.542 | 1.05361742 | 0.174326531 | 0.489724 | 0.558442 |
| MER58A | 2 | 1.603 | 1.247660636 | 0.1145 | 0.48786 | 0.558442 |
| AluSg | 2 | 1.601 | 1.249219238 | 0.1601 | 0.49231 | 0.558442 |
| AluJo | 8 | 7.652 | 1.045478306 | 0.186634146 | 0.507264 | 0.571141 |
| Tigger13a | 2 | 1.854 | 1.078748652 | 0.115875 | 0.568324 | 0.635185 |
| AluSx3 | 2 | 1.934 | 1.034126163 | 0.161166667 | 0.598804 | 0.664367 |
| MER112 | 2 | 2.122 | 0.942507069 | 0.151571429 | 0.649395 | 0.715276 |
| L1M7 | 1 | 1.094 | 0.914076782 | 0.091166667 | 0.682451 | 0.740947 |

| | | | | | | |
|---|---|---|---|---|---|---|
| LTR16A | 2 | 2.287 | 0.874508089 | 0.081678571 | 0.678834 | 0.740947 |
| L1MC | 1 | 1.122 | 0.891265597 | 0.1122 | 0.695805 | 0.750088 |
| AmnSINE1 | 1 | 1.166 | 0.857632933 | 0.083285714 | 0.704011 | 0.753589 |
| MER77B | 1 | 1.249 | 0.800640512 | 0.104083333 | 0.732569 | 0.778675 |
| L1MD1 | 2 | 2.544 | 0.786163522 | 0.106 | 0.738742 | 0.779784 |
| Charlie1 | 1 | 1.386 | 0.721500722 | 0.1386 | 0.775069 | 0.810967 |
| MER41A | 1 | 1.401 | 0.713775874 | 0.1401 | 0.778956 | 0.810967 |
| AluJr4 | 1 | 1.966 | 0.508646999 | 0.163833333 | 0.883181 | 0.913221 |
| LTR8A | 2 | 3.832 | 0.521920668 | 0.132137931 | 0.91114 | 0.935766 |
| Charlie1a | 1 | 3.052 | 0.327653997 | 0.132695652 | 0.962161 | 0.981533 |
| LTR67B | 0 | 1.476 | 0 | 0.113538462 | 1 | 1 |
| ALR/Alpha | 0 | 0.275 | 0 | 0.0171875 | 1 | 1 |
| HSATII | 0 | 0 | NaN | 0 | 1 | 1 |

Table S5. TE families near DEG in the L12 samples

| TE.family | Observed.instances | Expected.instances | Fold.enrichment | Prob.success | P.value | Q.value |
|---|---|---|---|---|---|---|
| L1MC4a | 17 | 3.524 | 4.824063564 | 0.060758621 | 3.71E-08 | 3.52E-06 |
| THE1B | 14 | 2.921 | 4.792879151 | 0.033193182 | 1.25E-06 | 5.95E-05 |
| MLT1K | 12 | 2.924 | 4.103967168 | 0.057333333 | 2.44E-05 | 0.000638 |
| L1MC4 | 10 | 2.08 | 4.807692308 | 0.056216216 | 2.69E-05 | 0.000638 |
| L2c | 22 | 8.461 | 2.600165465 | 0.056033113 | 3.93E-05 | 0.000747 |
| L2 | 15 | 5.259 | 2.85225328 | 0.051558824 | 0.000237 | 0.00375 |
| MIR | 23 | 10.621 | 2.165521137 | 0.062111111 | 0.000412 | 0.004346 |
| MIRb | 29 | 14.576 | 1.989571899 | 0.06150211 | 0.000349 | 0.004346 |
| MLT1L | 9 | 2.324 | 3.872633391 | 0.055333333 | 0.000408 | 0.004346 |
| MIR3 | 18 | 7.589 | 2.371853999 | 0.067758929 | 0.00054 | 0.005127 |
| L1MB3 | 13 | 4.793 | 2.712288754 | 0.066569444 | 0.000866 | 0.00748 |
| AluJr | 8 | 2.399 | 3.334722801 | 0.077387097 | 0.002007 | 0.015891 |
| MLT1H | 7 | 1.963 | 3.565970453 | 0.044613636 | 0.003148 | 0.021363 |
| MER41B | 10 | 3.624 | 2.759381898 | 0.061423729 | 0.002991 | 0.021363 |
| L2a | 14 | 6.176 | 2.266839378 | 0.055142857 | 0.003516 | 0.022226 |
| L1MB2 | 12 | 4.917 | 2.440512508 | 0.04917 | 0.003743 | 0.022226 |
| ERVL-E-int | 4 | 0.71 | 5.633802817 | 0.044375 | 0.004591 | 0.025656 |
| MLT1J | 7 | 2.193 | 3.191974464 | 0.05927027 | 0.005476 | 0.0289 |
| MER5A | 4 | 0.77 | 5.194805195 | 0.055 | 0.005865 | 0.029323 |
| AluSx | 8 | 3.003 | 2.664002664 | 0.079026316 | 0.008663 | 0.041148 |
| L1MB7 | 16 | 8.44 | 1.895734597 | 0.064427481 | 0.010375 | 0.046934 |
| L3 | 7 | 2.541 | 2.754820937 | 0.05523913 | 0.012585 | 0.048719 |
| L2b | 13 | 6.458 | 2.013007123 | 0.065897959 | 0.012218 | 0.048719 |
| LTR16C | 3 | 0.516 | 5.813953488 | 0.046909091 | 0.012821 | 0.048719 |
| THE1D | 3 | 0.495 | 6.060606061 | 0.033 | 0.012139 | 0.048719 |
| AluJo | 6 | 2.12 | 2.830188679 | 0.078518519 | 0.016461 | 0.057938 |
| L1MD2 | 4 | 0.999 | 4.004004004 | 0.03996 | 0.016467 | 0.057938 |
| AluY | 5 | 1.607 | 3.111387679 | 0.06428 | 0.019751 | 0.062545 |
| L1ME4a | 4 | 1.078 | 3.710575139 | 0.063411765 | 0.019748 | 0.062545 |
| MER51A | 3 | 0.593 | 5.059021922 | 0.053909091 | 0.018644 | 0.062545 |
| AluSc | 3 | 0.627 | 4.784688995 | 0.0627 | 0.021187 | 0.064927 |
| L1MB4 | 5 | 1.633 | 3.061849357 | 0.052677419 | 0.021908 | 0.065038 |
| HAL1 | 6 | 2.273 | 2.639683238 | 0.059815789 | 0.024263 | 0.069848 |
| L1M5 | 5 | 1.699 | 2.942907593 | 0.048542857 | 0.025965 | 0.072548 |
| Tigger3b | 7 | 2.994 | 2.338009352 | 0.0499 | 0.029416 | 0.079844 |
| MLT1H2 | 3 | 0.697 | 4.304160689 | 0.041 | 0.030457 | 0.080372 |
| AluJb | 8 | 3.827 | 2.090410243 | 0.078102041 | 0.034937 | 0.089704 |
| MIRc | 12 | 6.752 | 1.777251185 | 0.063102804 | 0.037651 | 0.094127 |
| MER44B | 5 | 1.887 | 2.649708532 | 0.038510204 | 0.039724 | 0.096764 |
| L1M4 | 3 | 0.791 | 3.792667509 | 0.046529412 | 0.042002 | 0.099756 |
| Tigger3c | 4 | 1.393 | 2.871500359 | 0.05572 | 0.047758 | 0.110659 |
| THE1C | 4 | 1.401 | 2.855103498 | 0.029808511 | 0.051098 | 0.11069 |
| AluSx1 | 5 | 2.068 | 2.417794971 | 0.079538462 | 0.051267 | 0.11069 |
| L1ME2 | 3 | 0.847 | 3.541912633 | 0.047055556 | 0.050081 | 0.11069 |
| L1ME1 | 5 | 2.139 | 2.337540907 | 0.047533333 | 0.061392 | 0.129606 |

| | | | | | | |
|---|---|---|---|---|---|---|
| MLT1C | 4 | 1.551 | 2.578981302 | 0.055392857 | 0.066699 | 0.132345 |
| L1ME3A | 2 | 0.431 | 4.64037123 | 0.0431 | 0.066408 | 0.132345 |
| LTR8 | 3 | 0.96 | 3.125 | 0.064 | 0.066869 | 0.132345 |
| AluSq2 | 3 | 0.998 | 3.006012024 | 0.076769231 | 0.072346 | 0.135047 |
| Tigger7 | 4 | 1.589 | 2.517306482 | 0.048151515 | 0.072499 | 0.135047 |
| MLT1F | 4 | 1.583 | 2.526847757 | 0.04946875 | 0.071552 | 0.135047 |
| MLT1G1 | 2 | 0.518 | 3.861003861 | 0.0518 | 0.091567 | 0.167286 |
| MER82 | 2 | 0.542 | 3.6900369 | 0.0542 | 0.098972 | 0.177402 |
| L1MD3 | 2 | 0.579 | 3.454231434 | 0.0579 | 0.110736 | 0.194813 |
| MLT1G | 2 | 0.597 | 3.350083752 | 0.054272727 | 0.117025 | 0.202135 |
| MLT1F2 | 3 | 1.217 | 2.465078061 | 0.045074074 | 0.120164 | 0.20385 |
| MER103C | 2 | 0.628 | 3.184713376 | 0.052333333 | 0.12768 | 0.209681 |
| MLT1N2 | 2 | 0.629 | 3.179650238 | 0.052416667 | 0.128016 | 0.209681 |
| MLT1A0 | 2 | 0.691 | 2.894356006 | 0.046066667 | 0.150036 | 0.241583 |
| MER58A | 2 | 0.721 | 2.773925104 | 0.055461538 | 0.160172 | 0.253605 |
| MER20 | 3 | 1.424 | 2.106741573 | 0.052740741 | 0.168391 | 0.262249 |
| HERV9-int | 2 | 0.786 | 2.544529262 | 0.0655 | 0.183367 | 0.280965 |
| MER44D | 2 | 0.796 | 2.512562814 | 0.034608696 | 0.188422 | 0.284128 |
| MER21C | 2 | 0.84 | 2.380952381 | 0.049411765 | 0.20407 | 0.302917 |
| AluSz | 4 | 2.525 | 1.584158416 | 0.076515152 | 0.243505 | 0.355892 |
| L1M1 | 1 | 0.295 | 3.389830508 | 0.0295 | 0.258766 | 0.372466 |
| MLT1F1 | 2 | 1.043 | 1.917545542 | 0.049666667 | 0.280387 | 0.397563 |
| AluSp | 2 | 1.066 | 1.876172608 | 0.071066667 | 0.289258 | 0.40411 |
| LTR71A | 1 | 0.419 | 2.386634845 | 0.038090909 | 0.34766 | 0.478662 |
| AluSz6 | 2 | 1.248 | 1.602564103 | 0.073411765 | 0.357955 | 0.478954 |
| Tigger3 | 1 | 0.427 | 2.341920375 | 0.0427 | 0.353631 | 0.478954 |
| Tigger3a | 3 | 2.158 | 1.390176089 | 0.056789474 | 0.367194 | 0.484492 |
| THE1A | 1 | 0.5 | 2 | 0.035714286 | 0.398992 | 0.519236 |
| MLT1I | 2 | 1.399 | 1.429592566 | 0.041147059 | 0.410705 | 0.520226 |
| MLT1B | 1 | 0.509 | 1.964636542 | 0.0509 | 0.406911 | 0.520226 |
| L1MEc | 1 | 0.557 | 1.795332136 | 0.042846154 | 0.43407 | 0.535831 |
| Tigger1 | 2 | 1.466 | 1.36425648 | 0.044424242 | 0.434305 | 0.535831 |
| L1MC1 | 1 | 0.683 | 1.464128843 | 0.045533333 | 0.502937 | 0.609801 |
| MER45B | 1 | 0.683 | 1.464128843 | 0.0683 | 0.507098 | 0.609801 |
| MLT1J2 | 1 | 0.708 | 1.412429379 | 0.050571429 | 0.516416 | 0.613244 |
| AluSg | 1 | 0.999 | 1.001001001 | 0.071357143 | 0.645283 | 0.748686 |
| MER44C | 1 | 1.008 | 0.992063492 | 0.059294118 | 0.646234 | 0.748686 |
| L1MC5 | 1 | 1.187 | 0.842459983 | 0.05935 | 0.705855 | 0.807907 |
| LTR8A | 1 | 1.414 | 0.707213579 | 0.061478261 | 0.767609 | 0.868129 |
| L1MB8 | 2 | 2.884 | 0.693481276 | 0.058857143 | 0.791976 | 0.885149 |
| L1MB5 | 1 | 1.74 | 0.574712644 | 0.062142857 | 0.834107 | 0.921398 |
| L1MA9 | 0 | 0.331 | 0 | 0.0331 | 1 | 1 |
| MLT1H1 | 0 | 0.52 | 0 | 0.052 | 1 | 1 |
| L1MEd | 0 | 0.426 | 0 | 0.0426 | 1 | 1 |
| Tigger15a | 0 | 0.564 | 0 | 0.0564 | 1 | 1 |
| Charlie1a | 0 | 0.783 | 0 | 0.060230769 | 1 | 1 |
| MER81 | 0 | 0.787 | 0 | 0.065583333 | 1 | 1 |

| | | | | | | |
|---|---|---|---|---|---|---|
| L4 | 0 | 0.477 | 0 | 0.0477 | 1 | 1 |
| MER21B | 0 | 0.665 | 0 | 0.060454545 | 1 | 1 |
| LTR16A | 0 | 0.574 | 0 | 0.044153846 | 1 | 1 |
