## Supplementary Tables 6 and 7 for "Transposable elements have contributed human regulatory regions that are activated upon bacterial infection"

Table S6. PARs near DEGs in S12 samples (PARs belonging to the 14 related TE families included only)

| TE.chr | TE.start | TE.end | TE.family | Peak.pval | Gene | Distance.to.TSS |
|---|---|---|---|---|---|---|
| chr4 | 74961891 | 74962441 | MER41B | 1.46E-16 | CXCL2 | 309 |
| chr4 | 96169841 | 96170431 | Tigger3c | 5.25E-24 | UNC5C | 744 |
| chr12 | 92815641 | 92815947 | Tigger3a | 6.78E-07 | CLLU1OS | 855 |
| chr21 | 30678601 | 30678997 | MLT1L | 8.01E-15 | BACH1 | 1077 |
| chr2 | 37312845 | 37313364 | MLT1F | 3.59E-06 | GPATCH11 | 1184 |
| chr1 | 79114002 | 79114290 | THE1C | 8.18E-08 | IFI44 | 1189 |
| chr4 | 38823509 | 38823989 | Tigger3b | 1.11E-11 | TLR6 | 1345 |
| chr1 | 167754291 | 167754845 | MLT1H | 1.66E-31 | MPZL1 | 1457 |
| chr3 | 146260166 | 146260295 | Tigger3c | 5.24E-24 | PLSCR1 | 1526 |
| chr12 | 12221637 | 12222168 | MER41B | 8.10E-08 | BCL2L14 | 1699 |
| chr2 | 113814070 | 113814437 | THE1B | 5.98E-29 | IL36RN | 1776 |
| chr8 | 90794231 | 90794648 | MER44D | 1.73E-98 | RIPK2 | 1915 |
| chr14 | 50770677 | 50771020 | Tigger3a | 3.23E-34 | L2HGDH | 1942 |
| chr2 | 37371121 | 37371719 | Tigger3b | 7.01E-16 | EIF2AK2 | 2343 |
| chr1 | 89442051 | 89442703 | Tigger3b | 0.000211334 | RBMXL1 | 2434 |
| chr2 | 6998590 | 6998932 | THE1B | 1.33E-38 | CMPK2 | 2455 |
| chr3 | 148798669 | 148798884 | Tigger3c | 4.62E-10 | HLTF | 2533 |
| chr1 | 236554481 | 236554633 | MLT1K | 1.96E-20 | EDARADD | 2943 |
| chr6 | 160086744 | 160087114 | MLT1K | 1.06E-15 | SOD2 | 2973 |
| chr7 | 89787138 | 89787479 | Tigger3b | 5.14E-17 | STEAP1 | 3099 |
| chr18 | 8605965 | 8606329 | THE1B | 9.81E-23 | RAB12 | 3112 |
| chr4 | 100734383 | 100734746 | THE1B | 3.72E-08 | DAPP1 | 3242 |
| chr12 | 31452667 | 31452995 | Tigger3a | 8.01E-64 | FAM60A | 3432 |
| chr2 | 231195435 | 231196116 | MER44C | 8.44E-33 | SP140L | 3450 |
| chr15 | 49919655 | 49920738 | Tigger3b | 6.40E-07 | DTWD1 | 3479 |
| chr15 | 38822263 | 38822662 | MLT1H | 6.59E-16 | RASGRP1 | 3678 |
| chr2 | 191860018 | 191860372 | THE1B | 1.03E-13 | STAT1 | 3978 |
| chr2 | 152209727 | 152209975 | MLT1I | 1.62E-12 | TNFAIP6 | 4129 |
| chr11 | 32130164 | 32130273 | MLT1I | 1.31E-05 | RCN1 | 4232 |
| chr15 | 65958419 | 65958939 | MER44B | 8.01E-15 | DENND4A | 4260 |
| chr5 | 135175353 | 135175727 | THE1B | 0.000208536 | SLC25A48 | 4560 |
| chr4 | 169162045 | 169162419 | THE1B | 4.10E-08 | DDX60 | 4865 |
| chr5 | 17212220 | 17212797 | MER41B | 1.93E-17 | BASP1 | 4870 |
| chr5 | 16467573 | 16468010 | MLT1F | 3.15E-05 | FAM134B | 5135 |
| chr15 | 40482843 | 40483089 | MER44C | 2.85E-13 | BUB1B | 5170 |
| chr19 | 6536247 | 6536326 | MLT1K | 1.51E-09 | TNFSF9 | 5237 |
| chr16 | 2585558 | 2585999 | MLT1F | 3.23E-05 | CEMP1 | 5241 |
| chr2 | 29330930 | 29331464 | MER44B | 4.82E-05 | CLIP4 | 5396 |
| chr1 | 205876446 | 205876759 | MER44B | 8.10E-08 | SLC26A9 | 5415 |
| chr8 | 23393134 | 23393496 | THE1B | 9.08E-32 | SLC25A37 | 5485 |
| chr8 | 144629438 | 144629744 | MER41B | 1.17E-15 | GSDMD | 5631 |
| chr11 | 35633408 | 35633829 | MLT1L | 1.12E-10 | FJX1 | 5904 |
| chr2 | 37862722 | 37863085 | THE1C | 0.000149796 | CDC42EP3 | 5945 |
| chr5 | 150433555 | 150433851 | MLT1I | 4.32E-42 | TNIP1 | 6030 |
| chr8 | 144628695 | 144628758 | MER41B | 3.34E-20 | GSDMD | 6617 |

| | | | | | | |
|---|---|---|---|---|---|---|
| chr20 | 4767770 | 4768141 | MLT1I | 1.02E-08 | RASSF2 | 7098 |
| chr20 | 49119385 | 49119762 | MLT1K | 3.94E-19 | PTPN1 | 7127 |
| chr4 | 89368434 | 89368722 | THE1B | 9.58E-09 | HERC6 | 7437 |
| chr6 | 111612175 | 111612540 | THE1B | 0.000174193 | REV3L | 7692 |
| chr18 | 8601178 | 8601476 | MLT1L | 4.10E-21 | RAB12 | 7965 |
| chr8 | 22216286 | 22216553 | MER44B | 2.68E-34 | SLC39A14 | 8207 |
| chr4 | 159744755 | 159744946 | MLT1K | 1.07E-146 | FNIP2 | 8284 |
| chr12 | 113426846 | 113427200 | THE1B | 8.10E-08 | OAS2 | 8848 |
| chr12 | 39170521 | 39171577 | Tigger3b | 1.47E-20 | CPNE8 | 9028 |
| chr6 | 160137199 | 160137578 | MLT1L | 9.55E-63 | WTAP | 9037 |
| chr2 | 85539666 | 85539978 | Tigger3b | 3.89E-18 | TCF7L1 | 9150 |
| chr1 | 71481496 | 71481613 | MLT1L | 1.32E-21 | PTGER3 | 9182 |
| chr4 | 159699553 | 159699643 | MER44B | 6.13E-19 | FNIP2 | 9207 |
| chr11 | 32102748 | 32103127 | MLT1H | 5.03E-42 | RCN1 | 9321 |
| chr15 | 56269363 | 56269689 | Tigger3a | 3.61E-06 | NEDD4 | 9716 |
| chr18 | 61455044 | 61456239 | Tigger3b | 7.17E-21 | SERPINB7 | 9833 |
| chr14 | 53336155 | 53336966 | Tigger3b | 2.30E-11 | FERMT2 | 9864 |
| chr7 | 5622220 | 5622480 | MLT1L | 1.91E-26 | FSCN1 | 9957 |
| chr18 | 21488723 | 21488980 | MER44B | 3.27E-05 | LAMA3 | 10022 |
| chr3 | 146222268 | 146222899 | MER41B | 3.18E-15 | PLSCR1 | 10066 |
| chr22 | 29548763 | 29549395 | MER41B | 4.70E-35 | KREMEN1 | 10745 |
| chr6 | 116588043 | 116588850 | Tigger3b | 1.23E-77 | DSE | 11454 |
| chr18 | 55900488 | 55900843 | THE1B | 0.000208536 | NEDD4L | 11625 |
| chr4 | 185296841 | 185297205 | THE1C | 4.01E-21 | IRF2 | 11660 |
| chr5 | 35887443 | 35887607 | MLT1H | 3.89E-19 | IL7R | 11927 |
| chr3 | 38193351 | 38193489 | MLT1H | 8.10E-09 | MYD88 | 12120 |
| chr10 | 60110511 | 60111729 | Tigger3b | 2.65E-13 | UBE2D1 | 12485 |
| chr10 | 90595119 | 90595493 | Tigger3c | 8.74E-08 | ANKRD22 | 12561 |
| chr2 | 218302811 | 218303055 | MLT1L | 8.10E-08 | DIRC3 | 12578 |
| chr9 | 123677429 | 123677742 | Tigger3a | 8.10E-08 | TRAF1 | 12758 |
| chr8 | 70603935 | 70604201 | Tigger3b | 1.12E-10 | SLCO5A1 | 12924 |
| chr3 | 36917830 | 36918132 | MLT1K | 1.14E-14 | TRANK1 | 13185 |
| chr10 | 115455271 | 115455905 | MER41B | 2.98E-15 | CASP7 | 13236 |
| chr11 | 10458212 | 10458576 | THE1B | 1.02E-08 | AMPD3 | 13290 |
| chr2 | 192561577 | 192562205 | Tigger3 | 4.15E-10 | NABP1 | 13346 |
| chr20 | 36308911 | 36309023 | MLT1L | 6.59E-16 | CTNNBL1 | 13383 |
| chr16 | 80704083 | 80704457 | THE1C | 1.12E-10 | CDYL2 | 14042 |
| chr5 | 148190243 | 148192108 | Tigger3 | 1.74E-63 | ADRB2 | 14046 |
| chr9 | 113021117 | 113021387 | MLT1K | 8.35E-108 | TXN | 14818 |
| chr10 | 90597409 | 90597768 | THE1B | 5.98E-29 | ANKRD22 | 14851 |
| chr14 | 105491212 | 105491394 | MLT1H | 7.93E-09 | CDCA4 | 14939 |
| chr1 | 235695520 | 235696039 | MER44B | 1.12E-10 | GNG4 | 14946 |
| chr1 | 145090129 | 145090756 | MER41B | 1.67E-51 | PDE4DIP | 15304 |
| chr10 | 26970664 | 26971209 | MER44B | 2.96E-19 | PDSS1 | 15377 |
| chr1 | 209740923 | 209741264 | MLT1I | 3.77E-07 | CAMK1G | 15796 |
| chr1 | 33385542 | 33385832 | MLT1K | 8.01E-15 | RNF19B | 16212 |
| chr22 | 43259606 | 43259980 | THE1C | 9.88E-15 | ARFGAP3 | 16366 |

| | | | | | | |
|---|---|---|---|---|---|---|
| chr8 | 90809097 | 90809726 | MER41B | 3.48E-33 | RIPK2 | 16781 |
| chr2 | 201287414 | 201287953 | MER44B | 4.27E-38 | SPATS2L | 17417 |
| chrX | 108957156 | 108958109 | Tigger3 | 1.17E-16 | ACSL4 | 18276 |
| chr20 | 39675857 | 39676164 | Tigger3a | 9.58E-09 | TOP1 | 18399 |
| chr13 | 77503520 | 77503877 | THE1B | 6.33E-16 | IRG1 | 18753 |
| chr15 | 100890949 | 100891721 | Tigger3 | 1.97E-05 | ADAMTS1[illegible] | 19730 |
| chr16 | 27386408 | 27386919 | MLT1K | 1.50E-17 | IL4R | 19739 |
| chr17 | 40418984 | 40419694 | Tigger3b | 3.77E-07 | STAT5A | 19869 |
| chr7 | 100513942 | 100514403 | MER41B | 2.03E-10 | ACHE | 20097 |
| chr8 | 17456186 | 17456545 | THE1B | 4.77E-06 | PDGFRL | 21485 |
| chr14 | 71597389 | 71598020 | MER41B | 1.04E-12 | PCNX | 21652 |
| chr10 | 74055674 | 74056082 | MLT1K | 3.89E-28 | DDIT4 | 21897 |
| chr6 | 37207906 | 37208269 | MLT1F | 2.19E-21 | TMEM217 | 21960 |
| chr1 | 65809021 | 65809734 | MER44C | 2.82E-19 | DNAJC6 | 22012 |
| chr1 | 28718549 | 28718911 | MER44C | 2.29E-11 | PHACTR4 | 22294 |
| chr7 | 128761599 | 128761953 | THE1B | 2.93E-51 | TSPAN33 | 22757 |
| chr10 | 91115280 | 91115922 | MER41B | 4.80E-31 | IFIT3 | 23039 |
| chr2 | 16768851 | 16769204 | THE1B | 2.06E-53 | FAM49A | 23559 |
| chr6 | 144495225 | 144495491 | MLT1H | 8.01E-15 | STX11 | 23562 |
| chr4 | 113091403 | 113091944 | MER44B | 5.75E-23 | C4orf32 | 24850 |
| chr17 | 76238467 | 76238881 | MLT1F | 6.23E-08 | BIRC5 | 25959 |
| chr16 | 50700748 | 50701000 | MLT1K | 2.50E-05 | NOD2 | 26512 |
| chr5 | 52056851 | 52057214 | THE1B | 5.36E-24 | ITGA1 | 26514 |
| chr3 | 122464257 | 122464953 | MER44C | 2.04E-60 | PARP14 | 27026 |
| chr7 | 157100461 | 157100841 | MER41B | 6.59E-16 | DNAJB6 | 27232 |
| chr20 | 44135458 | 44135975 | MER44B | 3.89E-18 | WFDC2 | 27379 |
| chr5 | 56572179 | 56572542 | THE1C | 8.77E-81 | GPBP1 | 27490 |
| chr14 | 55278644 | 55279105 | MLT1K | 4.17E-34 | SAMD4A | 27660 |
| chr8 | 80707426 | 80707501 | MLT1L | 2.97E-09 | HEY1 | 28181 |
| chr12 | 6852492 | 6852856 | THE1B | 6.40E-07 | LAG3 | 28820 |
| chr2 | 37940551 | 37940807 | MER44B | 4.17E-34 | CDC42EP3 | 28925 |
| chr3 | 159676088 | 159676430 | THE1B | 8.01E-15 | IL12A | 30105 |
| chr10 | 97833582 | 97833754 | MLT1L | 5.54E-92 | CCNJ | 30242 |
| chr6 | 154444653 | 154445191 | MER41B | 9.01E-07 | IPCEF1 | 30438 |
| chr9 | 136695425 | 136695763 | THE1C | 1.09E-35 | VAV2 | 30569 |
| chr1 | 33515701 | 33516099 | MER44B | 1.35E-23 | ADC | 30604 |
| chr18 | 55950050 | 55950282 | MLT1I | 1.43E-09 | NEDD4L | 31503 |
| chr6 | 2919129 | 2919605 | MER41B | 1.67E-11 | SERPINB9 | 31629 |
| chr9 | 117583234 | 117584446 | Tigger3b | 4.74E-18 | TNFSF15 | 31634 |
| chr11 | 33753396 | 33753761 | THE1B | 3.77E-24 | C11orf91 | 32288 |
| chr5 | 114823291 | 114823711 | MLT1L | 1.52E-09 | FEM1C | 32895 |
| chr11 | 93245374 | 93245966 | Tigger3c | 2.39E-07 | SMCO4 | 33159 |
| chr2 | 197242057 | 197242412 | THE1B | 8.20E-16 | HECW2 | 33630 |
| chr4 | 103548936 | 103549210 | MLT1L | 1.63E-60 | NFKB1 | 34530 |
| chr2 | 224804164 | 224804525 | THE1B | 1.85E-11 | SERPINE2 | 35302 |
| chr16 | 50691133 | 50691397 | MLT1K | 1.89E-14 | NOD2 | 36115 |
| chr2 | 85398107 | 85398451 | THE1B | 8.10E-08 | TCF7L1 | 36616 |

| | | | | | | |
|---|---|---|---|---|---|---|
| chr1 | 207172325 | 207172661 | Tigger3a | 4.77E-06 | FCAMR | 36625 |
| chr1 | 159072793 | 159073149 | THE1C | 2.82E-19 | AIM2 | 36687 |
| chr4 | 78488695 | 78489242 | MER41B | 5.65E-06 | CXCL13 | 37742 |
| chr3 | 172185173 | 172185281 | MLT1L | 1.68E-68 | TNFSF10 | 38015 |
| chr6 | 116652380 | 116652757 | THE1C | 8.10E-08 | DSE | 38251 |
| chr6 | 43344297 | 43344837 | MLT1F | 8.10E-24 | ZNF318 | 40489 |
| chr10 | 86253219 | 86253424 | MER41B | 7.01E-16 | CCSER2 | 41246 |
| chr5 | 17259132 | 17259508 | THE1B | 1.01E-12 | BASP1 | 41463 |
| chr2 | 237434055 | 237434736 | MER44D | 8.18E-08 | ACKR3 | 41692 |
| chr11 | 14358638 | 14359318 | MER44D | 2.17E-107 | RRAS2 | 42632 |
| chr1 | 161085153 | 161085665 | MER44D | 9.58E-09 | PVRL4 | 42956 |
| chr18 | 43633860 | 43634245 | Tigger3a | 5.17E-17 | PSTPIP2 | 43134 |
| chr18 | 21313224 | 21313581 | THE1B | 6.40E-07 | LAMA3 | 43576 |
| chr21 | 30610444 | 30610516 | MLT1I | 2.96E-14 | BACH1 | 43976 |
| chr2 | 224795308 | 224795759 | MLT1H | 3.94E-06 | SERPINE2 | 44068 |
| chr3 | 53272944 | 53273302 | THE1B | 5.22E-08 | DCP1A | 44143 |
| chr3 | 172279540 | 172279935 | Tigger3b | 1.04E-12 | TNFSF10 | 44395 |
| chr8 | 110207785 | 110208132 | THE1B | 1.28E-23 | NUDCD1 | 45014 |
| chr12 | 92490629 | 92490869 | MLT1K | 9.58E-09 | BTG1 | 45415 |
| chr6 | 82247189 | 82247549 | THE1B | 3.25E-25 | FAM46A | 46033 |
| chr10 | 30676299 | 30676667 | MLT1K | 5.32E-25 | MAP3K8 | 46197 |
| chr4 | 111027480 | 111027993 | MER41B | 8.37E-07 | ELOVL6 | 46674 |
| chr3 | 5181859 | 5182234 | THE1B | 3.48E-13 | EDEM1 | 47095 |
| chr3 | 197522085 | 197522538 | MLT1K | 1.26E-26 | KIAA0226 | 47105 |
| chr1 | 247531208 | 247531843 | MER41B | 6.15E-34 | NLRP3 | 47613 |
| chr18 | 20784579 | 20784934 | THE1B | 1.91E-10 | CABLES1 | 48790 |
| chr9 | 15298347 | 15298710 | THE1B | 5.67E-31 | TTC39B | 49307 |
| chr3 | 195851561 | 195851943 | MLT1L | 3.21E-30 | TFRC | 50493 |
| chr19 | 45095760 | 45096307 | MER41B | 0.000208536 | PVR | 50789 |
| chr8 | 104092287 | 104092647 | THE1B | 9.38E-49 | C8orf56 | 52542 |
| chr20 | 39950417 | 39950856 | Tigger3b | 1.63E-24 | ZHX3 | 52787 |
| chr4 | 84066921 | 84067278 | THE1C | 8.28E-41 | PLAC8 | 52907 |
| chr6 | 24743677 | 24744015 | MLT1L | 8.66E-05 | FAM65B | 53584 |
| chr12 | 56174569 | 56174768 | MER44B | 6.81E-12 | MMP19 | 54447 |
| chr19 | 39842152 | 39842526 | MLT1K | 8.10E-08 | ZFP36 | 54925 |
| chr9 | 137353356 | 137353758 | MLT1F | 0.000208536 | RXRA | 54928 |
| chr6 | 53305963 | 53306345 | MLT1I | 6.43E-33 | GCLC | 55792 |
| chr11 | 43890723 | 43891068 | MLT1H | 1.83E-07 | C11orf96 | 55822 |
| chr11 | 102454069 | 102454414 | Tigger3a | 9.16E-08 | MMP7 | 56352 |
| chr22 | 37828492 | 37828997 | MLT1F | 8.10E-08 | CARD10 | 57401 |
| chr1 | 183577563 | 183577810 | MER44B | 4.26E-20 | SMG7 | 57593 |
| chr1 | 68092318 | 68092608 | MLT1I | 1.07E-07 | GADD45A | 58134 |
| chr6 | 134430822 | 134431087 | Tigger3c | 2.98E-29 | SGK1 | 59295 |
| chr1 | 65553215 | 65553573 | THE1B | 2.30E-82 | AK4 | 59657 |
| chr6 | 111559690 | 111560016 | MLT1K | 2.16E-05 | REV3L | 60216 |
| chr6 | 144532384 | 144532642 | MER44B | 4.62E-05 | STX11 | 60721 |
| chr13 | 51223258 | 51223921 | MER41B | 3.25E-25 | DLEU7 | 61221 |

| | | | | | | |
|---|---|---|---|---|---|---|
| chr14 | 100899097 | 100899436 | Tigger3a | 1.14E-13 | WARS | 62381 |
| chr5 | 172257796 | 172258300 | MLT1F | 1.87E-53 | DUSP1 | 62703 |
| chr8 | 123939744 | 123940063 | THE1B | 3.21E-30 | ZHX2 | 64120 |
| chr2 | 143740577 | 143741064 | MLT1H | 3.61E-06 | KYNU | 64329 |
| chr3 | 171959191 | 171959547 | THE1B | 1.12E-10 | FNDC3B | 65177 |
| chr1 | 149746990 | 149747619 | MER41B | 1.04E-12 | HIST2H2A/ | 65884 |
| chr1 | 33468454 | 33468908 | MLT1K | 1.09E-27 | RNF19B | 66185 |
| chr3 | 88244949 | 88245538 | MER41B | 2.61E-12 | RP11-159C | 66868 |
| chr20 | 48317136 | 48317463 | MER44B | 2.10E-06 | B4GALT5 | 67654 |
| chr11 | 36033656 | 36034380 | MER44C | 2.88E-16 | LDLRAD3 | 68007 |
| chr20 | 49058105 | 49058543 | MLT1L | 2.26E-12 | PTPN1 | 68346 |
| chr3 | 172303580 | 172303726 | MLT1K | 1.12E-10 | TNFSF10 | 68435 |
| chrX | 53743336 | 53743795 | MLT1L | 9.45E-62 | HUWE1 | 68952 |
| chr22 | 29264401 | 29264762 | THE1B | 9.58E-09 | XBP1 | 69507 |
| chr14 | 62092066 | 62092523 | MLT1L | 3.25E-25 | HIF1A | 69706 |
| chr1 | 192848509 | 192848879 | MLT1I | 2.48E-23 | RGS2 | 70236 |
| chr17 | 49011003 | 49011278 | Tigger3a | 4.95E-36 | TOB1 | 70273 |
| chr14 | 50531657 | 50532015 | MLT1I | 2.87E-13 | C14orf182 | 73384 |
| chr5 | 172121104 | 172121458 | MLT1K | 8.56E-23 | DUSP1 | 73633 |
| chr2 | 216734784 | 216735282 | MLT1F | 6.62E-14 | MREG | 73929 |
| chr4 | 77029801 | 77030940 | Tigger3b | 1.07E-43 | CXCL11 | 73958 |
| chr3 | 111623754 | 111623828 | Tigger3c | 6.55E-26 | ABHD10 | 74027 |
| chr17 | 13324346 | 13324678 | Tigger3a | 5.01E-06 | HS3ST3A1 | 74326 |
| chr7 | 26255628 | 26255953 | Tigger3a | 4.77E-06 | SNX10 | 75586 |
| chr4 | 141667648 | 141668016 | THE1B | 2.82E-19 | TBC1D9 | 75900 |
| chr9 | 113082479 | 113082828 | THE1B | 4.17E-34 | TXN | 76180 |
| chr13 | 27766592 | 27766941 | THE1B | 8.85E-20 | RASL11A | 77521 |
| chr4 | 110888924 | 110889380 | Tigger3c | 1.20E-05 | ELOVL6 | 77620 |
| chr11 | 34094326 | 34094653 | Tigger3a | 3.77E-07 | ABTB2 | 77880 |
| chr9 | 37840208 | 37841113 | Tigger3b | 8.56E-23 | SHB | 78016 |
| chr12 | 55326947 | 55327537 | MER41B | 1.11E-12 | MUCL1 | 78626 |
| chr1 | 114552129 | 114553228 | Tigger3b | 3.25E-25 | SYT6 | 78683 |
| chr7 | 148315264 | 148315634 | THE1C | 4.57E-05 | CUL1 | 79370 |
| chr9 | 3742764 | 3743008 | THE1B | 7.56E-17 | GLIS3 | 81117 |
| chr10 | 86294655 | 86294885 | THE1B | 2.16E-05 | CCSER2 | 82682 |
| chr11 | 35880831 | 35881068 | MLT1K | 2.29E-68 | LDLRAD3 | 84461 |
| chr1 | 169600737 | 169601100 | THE1B | 0.000208536 | F5 | 85010 |
| chr5 | 133364517 | 133365092 | MLT1F | 9.35E-37 | TCF7 | 85308 |
| chr2 | 224754093 | 224754450 | THE1B | 4.87977727280 | SERPINE2 | 85377 |
| chr3 | 132159324 | 132159642 | Tigger3a | 6.53E-09 | ACPP | 87678 |
| chr5 | 52315110 | 52315361 | MER44B | 1.18E-33 | ITGA1 | 88128 |
| chr7 | 80633297 | 80633790 | MLT1H | 3.48E-20 | SEMA3C | 88292 |
| chr12 | 66606858 | 66607146 | THE1B | 1.91E-10 | HELB | 89177 |
| chr6 | 160270994 | 160271369 | THE1B | 1.32E-21 | SOD2 | 89704 |
| chr10 | 92763086 | 92763450 | THE1B | 3.77E-07 | ANKRD1 | 91233 |
| chr4 | 103605889 | 103606137 | MER44B | 2.19E-189 | NFKB1 | 91483 |
| chr14 | 103893862 | 103894372 | MER41B | 8.28E-41 | CKB | 91622 |

| | | | | | | |
|---|---|---|---|---|---|---|
| chr5 | 150948731 | 150949095 | THE1B | 5.67E-07 | SLC36A1 | 92581 |
| chr2 | 96908087 | 96908411 | MLT1K | 1.30E-14 | NCAPH | 93112 |
| chr10 | 92765034 | 92765396 | THE1B | 1.07E-22 | ANKRD1 | 93181 |
| chr7 | 30227951 | 30228329 | THE1C | 4.35E-25 | ZNRF2 | 95592 |
| chr9 | 102827033 | 102827717 | Tigger3 | 4.45E-34 | STX17 | 97481 |
| chr6 | 144161944 | 144162299 | THE1B | 6.69E-21 | PLAGL1 | 99136 |
| chr2 | 28515344 | 28515668 | MLT1L | 8.10E-08 | FOSL2 | 99645 |
| chr4 | 75130799 | 75131140 | MLT1H | 2.66E-05 | EREG | 99718 |
| chrX | 47608589 | 47609093 | MLT1F | 2.78E-11 | ELK1 | 100303 |
| chr15 | 90886574 | 90887171 | MLT1K | 8.18E-18 | GDPGP1 | 102433 |
| chrX | 48428276 | 48428761 | MLT1F | 4.62E-10 | SLC38A5 | 103874 |
| chr1 | 222950955 | 222951083 | Tigger3c | 1.93E-07 | AIDA | 104239 |
| chr1 | 222951238 | 222951574 | Tigger3a | 9.58E-09 | AIDA | 104522 |
| chr4 | 159917934 | 159918179 | MLT1L | 1.18E-44 | RAPGEF2 | 107149 |
| chr20 | 509969 | 510585 | MER44C | 2.93E-51 | RBCK1 | 107171 |
| chr11 | 4513624 | 4513750 | MLT1I | 9.35E-14 | TRIM21 | 107466 |
| chr1 | 100702528 | 100703026 | MLT1H | 3.77E-07 | CDC14A | 107556 |
| chr1 | 226896112 | 226896546 | MLT1L | 3.23E-34 | C1orf95 | 107822 |
| chr14 | 54923845 | 54924481 | Tigger3b | 8.18E-08 | SAMD4A | 109332 |
| chr7 | 20513939 | 20514223 | MLT1I | 3.59E-42 | ITGB8 | 109671 |
| chr4 | 40317084 | 40317266 | Tigger3c | 1.06E-09 | RHOH | 115119 |
| chr14 | 69377382 | 69377746 | THE1B | 1.07E-22 | ZFP36L1 | 115918 |
| chr15 | 68476052 | 68478019 | Tigger3 | 2.16E-05 | ITGA11 | 116029 |
| chr21 | 30815888 | 30816105 | THE1C | 2.10E-06 | BACH1 | 116291 |
| chr1 | 51635848 | 51636244 | Tigger3a | 1.71E-52 | TTC39A | 116684 |
| chr9 | 95908954 | 95909315 | THE1B | 3.52E-49 | FGD3 | 116798 |
| chr14 | 103867720 | 103868076 | MLT1I | 5.65E-06 | CKB | 117918 |
| chr13 | 36887409 | 36887999 | Tigger3c | 3.89E-18 | CCNA1 | 117966 |
| chr6 | 110831016 | 110831645 | MER41B | 1.53E-16 | DDO | 118042 |
| chr7 | 148276401 | 148276764 | THE1B | 9.58E-09 | CUL1 | 118240 |
| chr5 | 40560874 | 40561232 | THE1B | 1.12E-10 | PTGER4 | 118366 |
| chr9 | 95910955 | 95911480 | MLT1F | 1.04E-48 | FGD3 | 118799 |
| chr16 | 46893821 | 46894151 | MLT1K | 3.92E-25 | MYLK3 | 121871 |
| chr6 | 18242613 | 18243341 | MER44C | 9.06E-58 | NHLRC1 | 121895 |
| chr3 | 141760140 | 141760503 | THE1B | 8.56E-23 | GK5 | 121909 |
| chr1 | 173131116 | 173132035 | Tigger3b | 2.16E-06 | TNFSF18 | 122016 |
| chr6 | 18510255 | 18510653 | MLT1K | 3.89E-18 | RNF144B | 122510 |
| chr1 | 180932720 | 180932932 | MER44B | 3.27E-05 | IER5 | 124704 |
| chr19 | 17125272 | 17125634 | THE1B | 9.35E-14 | F2RL3 | 125271 |
| chr5 | 74058073 | 74058797 | MER44C | 5.52E-09 | ENC1 | 125774 |
| chr4 | 159461224 | 159461798 | MER41B | 1.68E-68 | C4orf46 | 126031 |
| chr6 | 17993785 | 17994045 | MLT1I | 1.30E-23 | NHLRC1 | 126671 |
| chr20 | 23215450 | 23215793 | MLT1H | 8.18E-08 | GZF1 | 126992 |
| chr3 | 190104232 | 190104514 | MER44B | 4.69E-23 | IL1RAP | 127324 |
| chr2 | 74297288 | 74297640 | THE1B | 1.14E-08 | MTHFD2 | 128047 |
| chr20 | 23478177 | 23478514 | THE1B | 1.32E-21 | GZF1 | 128669 |
| chr11 | 17277995 | 17278364 | MER44B | 2.28E-160 | KCNJ11 | 129040 |

| | | | | | | |
|---|---|---|---|---|---|---|
| chr14 | 52315676 | 52316189 | MLT1F | 4.68E-20 | FRMD6 | 131703 |
| chr7 | 93651925 | 93652302 | THE1C | 5.17E-28 | TFPI2 | 132695 |
| chr2 | 143809113 | 143809389 | THE1B | 8.10E-24 | KYNU | 132865 |
| chr9 | 126629862 | 126630439 | MLT1K | 5.26E-17 | LHX2 | 133508 |
| chr2 | 143811044 | 143811385 | THE1B | 0.000323445 | KYNU | 134796 |
| chr15 | 75382673 | 75382965 | MLT1L | 2.85E-13 | RPP25 | 135026 |
| chr5 | 148343033 | 148343463 | MLT1L | 2.85E-27 | ADRB2 | 136877 |
| chr18 | 12973717 | 12973916 | MLT1L | 0.000208536 | PTPN2 | 136894 |
| chr2 | 136458975 | 136459437 | MER44D | 2.03E-10 | MCM6 | 137757 |
| chr5 | 17356208 | 17356254 | THE1B | 8.85E-20 | BASP1 | 138539 |
| chr9 | 33175508 | 33175883 | Tigger3a | 1.09E-27 | DNAJA1 | 141250 |
| chr4 | 123667421 | 123667743 | Tigger3a | 1.37E-33 | NUDT6 | 142107 |
| chr19 | 14700482 | 14700982 | MLT1F | 1.51E-09 | EMR2 | 142221 |
| chr15 | 38962977 | 38963566 | MLT1L | 7.15E-31 | RASGRP1 | 144392 |
| chr8 | 118658698 | 118659069 | MLT1I | 9.58E-09 | EXT1 | 147658 |
| chr17 | 33421929 | 33422202 | MLT1L | 9.58E-09 | SLFN5 | 147851 |
| chr6 | 2684153 | 2684680 | MER44B | 2.77E-12 | SERPINB1 | 147884 |
| chr11 | 9373659 | 9374002 | MLT1K | 6.40E-07 | DENND5A | 148046 |
| chr11 | 105022710 | 105023179 | Tigger3b | 5.15E-20 | CASP5 | 148709 |
| chr20 | 30929042 | 30929387 | Tigger3a | 6.79E-22 | PLAGL2 | 148736 |
| chr3 | 182690860 | 182691221 | MER44B | 9.35E-14 | LAMP3 | 148778 |
| chr1 | 154403854 | 154404497 | MER44D | 2.50E-90 | ADAR | 150039 |
| chr8 | 41880416 | 41881013 | Tigger3b | 8.44E-33 | PLAT | 151221 |
| chr14 | 21712783 | 21712997 | Tigger3b | 1.37E-21 | ZNF219 | 151366 |
| chr3 | 12724169 | 12724414 | MLT1K | 1.46E-19 | TSEN2 | 151822 |
| chr5 | 68763581 | 68763933 | MLT1K | 6.40E-07 | CCDC125 | 154116 |
| chr8 | 105671636 | 105671948 | MLT1K | 2.45E-16 | LRP12 | 155164 |
| chr20 | 61076914 | 61077279 | THE1C | 8.12E-47 | LAMA5 | 155300 |
| chr8 | 8483215 | 8483743 | MLT1H | 4.29E-18 | MFHAS1 | 157119 |
| chr7 | 22925525 | 22925994 | MLT1L | 2.78E-11 | IL6 | 157406 |
| chr14 | 103827940 | 103828503 | MER44D | 8.44E-15 | CKB | 157491 |
| chr12 | 56070634 | 56071291 | Tigger3b | 1.02E-09 | MMP19 | 157924 |
| chr2 | 223448377 | 223448742 | THE1C | 5.36E-24 | SGPP2 | 159141 |
| chr1 | 145600301 | 145600461 | Tigger3c | 9.58E-09 | TXNIP | 159448 |
| chr14 | 103762394 | 103762776 | Tigger3b | 4.77E-06 | TNFAIP2 | 159654 |
| chr12 | 4911986 | 4912391 | MLT1I | 2.85E-13 | AKAP3 | 160834 |
| chr3 | 98681205 | 98681664 | MLT1K | 5.48E-05 | ST3GAL6 | 161056 |
| chr1 | 63087633 | 63087914 | MLT1H | 1.27E-10 | ATG4C | 161890 |
| chr3 | 5063733 | 5064045 | MLT1K | 3.18E-07 | EDEM1 | 165284 |
| chr2 | 47951581 | 47952019 | MLT1K | 1.09E-27 | MSH2 | 168583 |
| chr4 | 68593363 | 68593784 | MLT1L | 1.07E-26 | STAP1 | 168917 |
| chr3 | 183047488 | 183047741 | Tigger3a | 3.27E-05 | LAMP3 | 171587 |
| chr10 | 112085840 | 112085998 | MLT1L | 2.98E-15 | DUSP5 | 171596 |
| chr1 | 51948608 | 51949030 | MER44B | 5.17E-17 | TTC39A | 171679 |
| chr17 | 70988944 | 70989302 | THE1B | 4.38E-34 | SSTR2 | 171847 |
| chr7 | 112546156 | 112546454 | Tigger3a | 9.25E-48 | GPR85 | 171930 |
| chr4 | 5040086 | 5040704 | Tigger3b | 1.73E-64 | MSX1 | 175931 |

| | | | | | | |
|---|---|---|---|---|---|---|
| chr6 | 109153603 | 109153835 | MLT1L | 2.29E-68 | FOXO3 | 176054 |
| chr5 | 114679681 | 114680409 | MER44C | 7.03E-117 | FEM1C | 176197 |
| chr3 | 32671637 | 32671960 | MLT1I | 3.27E-05 | CMTM7 | 179142 |
| chr4 | 76742480 | 76742640 | MLT1L | 5.53E-07 | CXCL9 | 179786 |
| chr4 | 103694206 | 103694525 | Tigger3b | 0.000283054 | NFKB1 | 179800 |
| chr14 | 94762506 | 94762869 | THE1B | 9.35E-14 | IFI27 | 180649 |
| chr10 | 45655447 | 45656107 | MER44D | 2.55E-135 | C10orf10 | 182220 |
| chr2 | 218753221 | 218753583 | THE1B | 1.11E-11 | DIRC3 | 184598 |
| chr5 | 147391432 | 147391514 | MLT1L | 2.42E-08 | SPINK1 | 184783 |
| chr4 | 112880620 | 112881098 | MLT1F | 1.88E-155 | C4orf32 | 185453 |
| chr2 | 228491364 | 228491576 | Tigger3c | 4.56E-13 | CCL20 | 186980 |
| chr17 | 9253347 | 9253724 | MER44D | 7.55E-59 | NTN1 | 187095 |
| chr8 | 41844574 | 41844872 | Tigger3a | 2.98E-15 | PLAT | 187362 |
| chr14 | 52372374 | 52372739 | THE1C | 2.39E-07 | FRMD6 | 188401 |
| chr5 | 156323346 | 156323752 | MLT1K | 5.26E-06 | HAVCR2 | 189089 |
| chr2 | 85170212 | 85170370 | MLT1K | 3.76E-09 | TCF7L1 | 190161 |
| chr1 | 182374947 | 182375396 | Tigger3c | 2.61E-12 | RGS16 | 192360 |
| chr7 | 38229751 | 38229977 | MER44B | 2.82E-19 | AMPH | 193326 |
| chr4 | 14809411 | 14809775 | THE1B | 8.01E-15 | CPEB2 | 194521 |
| chr15 | 88983151 | 88983694 | MLT1K | 3.12E-10 | ISG20 | 195688 |
| chr15 | 44760918 | 44761440 | MER44B | 2.95E-27 | PATL2 | 196488 |
| chr7 | 86833264 | 86833621 | THE1B | 1.64E-10 | ABCB4 | 197390 |
| chr12 | 94167004 | 94167340 | THE1B | 4.59E-82 | SOCS2 | 198170 |
| chr11 | 61872516 | 61873051 | MER41B | 9.35E-14 | RAB3IL1 | 198571 |
| chr3 | 128949653 | 128950508 | Tigger3b | 9.24E-26 | MBD4 | 199277 |
| chr13 | 67079338 | 67079664 | THE1B | 8.10E-08 | PCDH9 | 200602 |
| chr5 | 55462953 | 55463289 | Tigger3a | 0.000392265 | IL6ST | 202916 |
| chr4 | 156571954 | 156572313 | THE1B | 3.27E-05 | TDO2 | 203575 |
| chr5 | 55464046 | 55464530 | MER44B | 1.45E-109 | IL6ST | 204009 |
| chr19 | 17618766 | 17619037 | MER44B | 1.90E-55 | ABHD8 | 206668 |
| chr6 | 25047223 | 25047546 | THE1B | 6.40E-07 | FAM65B | 206670 |
| chr12 | 38831958 | 38832321 | THE1B | 6.40E-07 | CPNE8 | 208301 |
| chr6 | 119188403 | 119188966 | Tigger3c | 9.35E-14 | CEP85L | 210477 |
| chr8 | 21555167 | 21555530 | THE1B | 5.25E-24 | DOK2 | 210852 |
| chr10 | 75782917 | 75783232 | Tigger3a | 9.58E-09 | NDST2 | 212475 |
| chr2 | 85146889 | 85147344 | MER44B | 4.88E-07 | TCF7L1 | 213187 |
| chr10 | 62414628 | 62415857 | Tigger3b | 8.01E-64 | RHOBTB1 | 213337 |
| chr3 | 119580627 | 119580726 | Tigger3c | 1.37E-21 | POPDC2 | 213339 |
| chr3 | 142623575 | 142623917 | MLT1K | 7.91E-23 | CHST2 | 214254 |
| chr1 | 9137900 | 9138512 | MER41B | 1.11E-11 | SPSB1 | 214425 |
| chr4 | 15271914 | 15272273 | THE1B | 8.10E-08 | CPEB2 | 216157 |
| chr1 | 78137598 | 78137797 | MER44B | 1.07E-22 | NEXN | 216399 |
| chr16 | 48355406 | 48355857 | Tigger3b | 1.28E-33 | N4BP1 | 216778 |
| chr2 | 43666528 | 43666941 | MER44B | 1.11E-11 | ZFP36L2 | 216987 |
| chr4 | 169628378 | 169628897 | MLT1H | 1.14E-08 | DDX60L | 218137 |
| chr2 | 231620858 | 231621165 | Tigger3a | 9.35E-14 | SP100 | 218501 |
| chr3 | 45103610 | 45103839 | MER41B | 2.07E-10 | KIF15 | 218995 |

| | | | | | | |
|---|---|---|---|---|---|---|
| chr6 | 12069268 | 12069720 | Tigger3b | 8.01E-15 | EDN1 | 220874 |
| chr3 | 45106144 | 45106761 | MER41B | 9.22E-08 | KIF15 | 221529 |
| chr1 | 184129504 | 184130051 | MLT1H | 1.39E-51 | C1orf21 | 226139 |
| chr7 | 106145071 | 106145435 | THE1B | 8.87E-06 | NAMPT | 227569 |
| chr7 | 22536596 | 22537152 | MLT1K | 7.36E-21 | IL6 | 228349 |
| chr11 | 129013184 | 129013887 | MER44C | 2.30E-59 | KCNJ5 | 232105 |
| chr6 | 16005726 | 16006284 | MLT1L | 9.35E-14 | GMPR | 232525 |
| chr6 | 17887235 | 17887599 | MLT1I | 4.38E-59 | NHLRC1 | 233117 |
| chr7 | 106756170 | 106756533 | MLT1I | 3.05E-14 | PIK3CG | 233561 |
| chr2 | 85125510 | 85126036 | MLT1F | 1.43E-22 | TCF7L1 | 234495 |
| chr6 | 143336497 | 143336859 | THE1B | 1.16E-15 | HIVEP2 | 235817 |
| chr10 | 115911224 | 115911828 | MER44C | 1.14E-08 | AL162407. | 236694 |
| chr3 | 37608553 | 37608911 | THE1B | 1.12E-10 | GOLGA4 | 239297 |
| chr3 | 9050449 | 9050860 | MLT1K | 5.97E-45 | OXTR | 240320 |
| chr13 | 67120269 | 67121207 | Tigger3b | 4.77E-06 | PCDH9 | 241533 |
| chr20 | 6990971 | 6991535 | MER41B | 3.48E-38 | BMP2 | 242660 |
| chr5 | 52471794 | 52472572 | Tigger3 | 0.000208536 | ITGA1 | 244812 |
| chr4 | 24755122 | 24755453 | MLT1I | 5.55E-76 | LGI2 | 245014 |
| chr5 | 149472015 | 149472372 | THE1B | 5.17E-17 | PPARGC1B | 246809 |
| chr3 | 131348910 | 131349157 | MER44B | 6.40E-07 | NUDT16 | 248281 |
| chr3 | 131349154 | 131349234 | MER44D | 6.40E-07 | NUDT16 | 248525 |
| chr9 | 92470228 | 92470587 | THE1B | 1.04E-12 | GADD45G | 250221 |
| chr8 | 80425424 | 80425787 | THE1B | 6.40E-07 | HEY1 | 250456 |
| chr1 | 212484820 | 212485692 | Tigger3b | 2.16E-06 | ATF3 | 252982 |
| chr1 | 24814651 | 24815016 | THE1B | 3.89E-18 | CLIC4 | 256830 |
| chr3 | 129414219 | 129414582 | THE1C | 5.69E-73 | MBD4 | 258329 |
| chr1 | 28437117 | 28437443 | MLT1L | 2.46E-11 | PHACTR4 | 258669 |
| chr1 | 235002028 | 235002197 | MLT1F | 2.76E-09 | IRF2BP2 | 259051 |
| chr1 | 221314446 | 221314811 | THE1B | 1.85E-25 | HLX | 259862 |
| chr8 | 81930682 | 81931045 | THE1B | 2.42E-65 | FABP5 | 261551 |
| chr2 | 200907828 | 200907925 | MLT1K | 4.77E-06 | SPATS2L | 262677 |
| chr12 | 110201106 | 110201640 | MLT1F | 1.44E-05 | C12orf76 | 264230 |
| chr4 | 90431052 | 90431219 | MLT1I | 2.16E-05 | GPRIN3 | 264966 |
| chr14 | 68205748 | 68206146 | MLT1L | 5.94E-05 | TMEM229 | 265273 |
| chr20 | 23076836 | 23077271 | MLT1L | 1.11E-11 | GZF1 | 265514 |
| chr20 | 3401009 | 3401541 | MLT1F | 3.70E-24 | SIGLEC1 | 266074 |
| chr7 | 93786761 | 93787403 | MER41B | 2.18E-69 | TFPI2 | 267531 |
| chr8 | 81923439 | 81923797 | THE1B | 6.71E-20 | FABP5 | 268799 |
| chr9 | 76052776 | 76053462 | MER44C | 3.85E-50 | ANXA1 | 269762 |
| chr12 | 122502048 | 122502474 | MLT1K | 2.99E-11 | RHOF | 270991 |
| chr14 | 21265239 | 21265561 | Tigger3b | 2.22E-28 | ARHGEF40 | 272866 |
| chr14 | 21264657 | 21264942 | Tigger3b | 1.70E-13 | ARHGEF40 | 273485 |
| chr11 | 86926318 | 86926828 | MLT1H | 1.13E-06 | PRSS23 | 274490 |
| chr21 | 16262567 | 16263288 | Tigger3b | 6.40E-07 | AF165138. | 277741 |
| chr10 | 92950777 | 92951311 | MER44B | 1.44E-22 | ANKRD1 | 278924 |
| chr14 | 95603088 | 95603302 | MLT1L | 4.09E-18 | SYNE3 | 280527 |
| chr3 | 57941755 | 57942079 | Tigger3a | 1.07E-77 | ABHD6 | 281152 |

| | | | | | | |
|---|---|---|---|---|---|---|
| chr2 | 224333992 | 224334472 | MLT1H | 5.43E-73 | AP1S3 | 281929 |
| chr11 | 61382315 | 61382683 | THE1B | 2.61E-12 | RAB3IL1 | 282088 |
| chr4 | 141873984 | 141874315 | Tigger3a | 3.89E-18 | TBC1D9 | 282236 |
| chr2 | 128690191 | 128690493 | Tigger3b | 8.01E-15 | LIMS2 | 290491 |
| chr18 | 56349774 | 56350008 | MLT1K | 1.26E-06 | NEDD4L | 292478 |
| chr20 | 3961008 | 3961262 | MER44B | 2.01E-99 | SIGLEC1 | 293389 |
| chr3 | 39518787 | 39519973 | Tigger3b | 8.10E-08 | XIRP1 | 294080 |
| chr9 | 27653449 | 27653790 | Tigger3b | 2.78E-05 | MOB3B | 294427 |
| chr2 | 191533502 | 191534027 | MER44B | 5.17E-17 | STAT1 | 295055 |
| chr2 | 223584511 | 223584841 | THE1B | 9.58E-09 | SGPP2 | 295275 |
| chr12 | 102609162 | 102610373 | Tigger3b | 8.10E-08 | DRAM1 | 295365 |
| chr5 | 149522186 | 149522863 | MER44D | 1.89E-14 | PPARGC1B | 296980 |
| chr2 | 127758611 | 127758872 | Tigger3b | 0.000208536 | MAP3K2 | 297432 |
| chr7 | 51145725 | 51146253 | MLT1L | 2.37E-46 | GRB10 | 298310 |
| chr4 | 154569994 | 154571181 | Tigger3b | 1.53E-77 | MND1 | 298779 |
| chr5 | 157297791 | 157298272 | MER44D | 4.29E-136 | ADAM19 | 300337 |
| chr5 | 58090790 | 58091155 | THE1B | 5.95E-25 | GAPT | 300723 |
| chr5 | 133780443 | 133780914 | MLT1F | 1.06E-09 | TCF7 | 301960 |
| chr20 | 36778587 | 36778819 | MLT1H | 1.96E-20 | CTNNBL1 | 302327 |
| chr14 | 94884277 | 94884632 | THE1B | 2.82E-19 | IFI27 | 302420 |
| chr9 | 107906250 | 107906600 | Tigger3a | 9.25E-24 | FSD1L | 303475 |
| chr2 | 204875869 | 204876273 | MLT1K | 9.58E-09 | CD28 | 304453 |
| chr20 | 17644584 | 17645088 | MER44B | 3.27E-05 | MGME1 | 304466 |
| chr15 | 90325520 | 90325966 | MER41B | 2.02E-17 | RHCG | 304490 |
| chr7 | 80851190 | 80851544 | THE1B | 1.64E-10 | SEMA3C | 306185 |
| chr3 | 153187079 | 153187294 | Tigger3b | 5.67E-07 | RAP2B | 307050 |
| chr3 | 186778574 | 186778735 | MER44B | 5.36E-24 | RTP4 | 307383 |
| chr6 | 44420452 | 44420688 | MER44D | 1.32E-21 | TMEM63B | 313003 |
| chr6 | 11851570 | 11852087 | MLT1H | 1.73E-53 | TMEM170 | 313059 |
| chr10 | 104892747 | 104893030 | THE1B | 8.43E-15 | CALHM2 | 313511 |
| chr6 | 133369450 | 133370298 | Tigger3 | 0.000208536 | VNN3 | 314677 |
| chr4 | 177926806 | 177927315 | MER41B | 7.39E-22 | VEGFC | 316352 |
| chr3 | 46131698 | 46131848 | MLT1F | 5.86E-09 | CCRL2 | 316804 |
| chr16 | 20100621 | 20101025 | MLT1L | 8.10E-08 | ACSM5 | 319829 |
| chr11 | 94182768 | 94183110 | Tigger3a | 5.06E-69 | PANX1 | 320673 |
| chr6 | 167893576 | 167894006 | MER41B | 5.97E-12 | GPR31 | 323817 |
| chr5 | 131084417 | 131085077 | Tigger3b | 2.21E-09 | CSF2 | 324404 |
| chr6 | 37950219 | 37951119 | Tigger3b | 0.000353273 | MDGA1 | 326563 |
| chr2 | 61436547 | 61437714 | Tigger3b | 4.17E-34 | REL | 327756 |
| chr15 | 66665256 | 66665523 | Tigger3a | 8.01E-15 | SMAD6 | 329041 |
| chr15 | 73942987 | 73943366 | MLT1L | 2.46E-11 | STOML1 | 332179 |
| chr12 | 62662886 | 62663204 | Tigger3a | 1.34E-31 | C12orf61 | 332325 |
| chr7 | 32720379 | 32720882 | MER41B | 0.000208536 | NT5C3A | 332858 |
| chr18 | 74356045 | 74356161 | MLT1L | 1.91E-26 | MBP | 334620 |
| chr2 | 159489650 | 159490150 | MER44B | 2.29E-11 | TANC1 | 334994 |
| chr5 | 119027976 | 119028314 | THE1C | 2.61E-12 | TNFAIP8 | 336246 |
| chr2 | 28981894 | 28982099 | Tigger3b | 1.47E-43 | CLIP4 | 338470 |

| | | | | | | |
|---|---|---|---|---|---|---|
| chr12 | 93623055 | 93623613 | MER44B | 1.59E-47 | SOCS2 | 339975 |
| chr7 | 141264041 | 141264412 | THE1C | 1.90E-81 | MGAM | 343199 |
| chr2 | 17093009 | 17093368 | THE1C | 2.55E-09 | FAM49A | 347717 |
| chr1 | 70965013 | 70965905 | Tigger3b | 3.80E-37 | PTGER3 | 352129 |
| chr7 | 102429096 | 102429570 | Tigger3b | 0.000208536 | ORAI2 | 352350 |
| chr4 | 37238928 | 37239294 | THE1B | 8.10E-08 | RELL1 | 353126 |
| chr8 | 125209347 | 125209457 | MLT1K | 3.91E-23 | MTSS1 | 353572 |
| chr14 | 90508841 | 90509208 | THE1B | 9.58E-09 | CALM1 | 353636 |
| chr7 | 92373464 | 92373836 | THE1B | 2.87E-11 | SAMD9 | 354991 |
| chr1 | 185213369 | 185213930 | Tigger3b | 1.19E-06 | FAM129A | 356496 |
| chr1 | 55431747 | 55432187 | MLT1L | 2.10E-06 | FAM151A | 356892 |
| chr6 | 143461092 | 143461453 | THE1B | 2.78E-05 | HIVEP2 | 360412 |
| chr3 | 45245796 | 45245895 | MLT1K | 9.16E-08 | KIF15 | 361181 |
| chr1 | 150895048 | 150895415 | MLT1I | 1.27E-44 | ADAMTSL4 | 361568 |
| chr4 | 78069467 | 78069814 | Tigger3a | 9.58E-09 | CXCL13 | 363091 |
| chr3 | 189306283 | 189306381 | MLT1I | 0.000208536 | LEPREL1 | 368134 |
| chr9 | 96160338 | 96160702 | THE1B | 3.53E-42 | FGD3 | 368182 |
| chr4 | 141172061 | 141172531 | MLT1F | 8.55E-06 | TBC1D9 | 369386 |
| chr8 | 90397888 | 90398411 | MLT1H | 6.40E-07 | RIPK2 | 371562 |
| chr6 | 112265884 | 112266284 | Tigger3c | 1.38E-29 | TRAF3IP2 | 371964 |
| chr2 | 223662426 | 223662798 | Tigger3a | 1.39E-61 | SGPP2 | 373190 |
| chr1 | 202500980 | 202501603 | MER41B | 4.77E-06 | PTPN7 | 373394 |
| chr3 | 112265154 | 112265403 | MER44B | 1.96E-20 | CD200R1 | 374651 |
| chr8 | 94885846 | 94886476 | MER41B | 4.77E-06 | GEM | 375003 |
| chr1 | 155540092 | 155540553 | MER44D | 8.44E-33 | MUC1 | 378099 |
| chr5 | 14202611 | 14203028 | THE1B | 6.26E-07 | FAM105A | 378854 |
| chr1 | 52156453 | 52156945 | Tigger3b | 3.52E-19 | TTC39A | 379524 |
| chr5 | 169278606 | 169278974 | THE1C | 3.81E-08 | C5orf58 | 380475 |
| chr10 | 28580559 | 28581243 | Tigger3b | 1.31E-08 | BAMBI | 385026 |
| chr5 | 74708316 | 74709009 | Tigger3b | 0.00026457 | GCNT4 | 385027 |
| chr11 | 21828549 | 21828906 | THE1C | 2.05E-23 | ANO5 | 385814 |
| chr1 | 229269548 | 229269914 | THE1C | 2.10E-06 | RHOU | 387561 |
| chr2 | 75172216 | 75172603 | MLT1F | 1.88E-05 | DOK1 | 389883 |
| chr2 | 136995651 | 136996006 | MLT1K | 1.08E-41 | MCM6 | 389925 |
| chr3 | 113395117 | 113395486 | THE1C | 1.27E-23 | BOC | 391710 |
| chr1 | 8960144 | 8960226 | MER41B | 3.27E-05 | SPSB1 | 392711 |
| chr9 | 33836197 | 33836410 | Tigger3c | 3.81E-17 | AQP3 | 393818 |
| chr1 | 56566141 | 56566408 | MLT1I | 0.000208536 | PPAP2B | 394009 |
| chr20 | 35851331 | 35851688 | MLT1I | 5.08E-08 | SOGA1 | 394331 |
| chr20 | 47459683 | 47459812 | MLT1K | 3.27E-05 | ZNFX1 | 394669 |
| chr1 | 42995503 | 42996082 | MER44D | 2.68E-34 | SLC2A1 | 394968 |
| chr9 | 26929880 | 26929963 | MLT1I | 2.82E-19 | MOB3B | 395242 |
| chr22 | 37045653 | 37046017 | THE1B | 1.09E-27 | APOL1 | 396443 |
| chrX | 13321357 | 13321723 | THE1B | 0.00012044 | TLR8 | 396599 |
| chr8 | 42442949 | 42443326 | MLT1H | 2.22E-28 | PLAT | 397966 |
| chr1 | 56561925 | 56562346 | MLT1F | 1.45E-41 | PPAP2B | 398071 |
| chr3 | 141405445 | 141406028 | MLT1K | 3.05E-14 | ACPL2 | 399244 |

| | | | | | | |
|---|---|---|---|---|---|---|
| chr12 | 25711495 | 25711854 | THE1B | 3.27E-05 | RASSF8 | 400106 |
| chr16 | 21314215 | 21314566 | THE1B | 2.91E-24 | LYRM1 | 401789 |
| chr3 | 112237842 | 112238202 | THE1B | 4.23E-05 | CD200R1 | 401852 |
| chr1 | 193180251 | 193180975 | MER44C | 5.70E-133 | RGS2 | 401978 |
| chr12 | 109088700 | 109089127 | MLT1F | 9.04E-13 | CMKLR1 | 402390 |
| chr20 | 4070565 | 4070749 | MLT1H | 1.12E-10 | SIGLEC1 | 402946 |
| chr20 | 35916391 | 35916907 | MLT1F | 5.17E-17 | CTNNBL1 | 405499 |
| chr15 | 91191646 | 91192244 | MER41B | 1.85E-09 | GDPGP1 | 407505 |
| chr2 | 65089508 | 65089966 | MLT1H | 0.000215635 | LGALSL | 407832 |
| chr20 | 7156499 | 7156863 | THE1B | 0.00012044 | BMP2 | 408188 |
| chr9 | 73885555 | 73886065 | MLT1F | 8.75E-09 | TMEM2 | 412215 |
| chr2 | 98646267 | 98646784 | MLT1F | 1.19E-08 | INPP4A | 414531 |
| chr15 | 74827692 | 74828192 | MLT1H | 1.14E-08 | RPP25 | 418563 |
| chr5 | 34626246 | 34626916 | Tigger3b | 9.35E-37 | PRLR | 421943 |
| chr20 | 33717389 | 33717820 | Tigger3b | 4.71E-08 | TP53INP2 | 424850 |
| chr6 | 111140861 | 111141427 | MER41B | 3.07E-18 | DDO | 427887 |
| chr20 | 36904153 | 36904552 | Tigger3b | 4.19E-08 | CTNNBL1 | 427893 |
| chr2 | 218997197 | 218997497 | Tigger3a | 7.56E-17 | DIRC3 | 428574 |
| chr3 | 184532149 | 184533290 | Tigger3b | 1.42E-23 | CHRD | 429573 |
| chr7 | 73548589 | 73548934 | MER41B | 2.08E-22 | STX1A | 430079 |
| chr11 | 127896216 | 127896931 | MER44C | 7.43E-189 | ETS1 | 431723 |
| chr10 | 45906248 | 45906876 | MER41B | 5.01E-42 | C10orf10 | 433021 |
| chr14 | 61220499 | 61220960 | MLT1F | 3.56E-06 | PRKCH | 433315 |
| chr7 | 113161161 | 113161869 | MER44D | 1.31E-10 | GPR85 | 436375 |
| chr5 | 40240108 | 40240611 | MLT1F | 1.96E-20 | PTGER4 | 438987 |
| chr4 | 18026721 | 18027243 | MLT1H | 5.59E-51 | LAP3 | 439329 |
| chr4 | 38077665 | 38078029 | THE1C | 1.16E-10 | RELL1 | 441156 |
| chr6 | 119419177 | 119419534 | THE1B | 6.59E-16 | CEP85L | 441251 |
| chr5 | 79423039 | 79423362 | MLT1I | 1.08E-05 | PAPD4 | 445564 |
| chr6 | 71552324 | 71552722 | MLT1L | 0.00012044 | OGFRL1 | 445782 |
| chr1 | 226288636 | 226288846 | Tigger3b | 1.16E-15 | C1orf95 | 447653 |
| chr12 | 102762569 | 102762975 | THE1C | 0.000208536 | DRAM1 | 448772 |
| chr6 | 71549088 | 71549198 | MLT1I | 1.04E-12 | OGFRL1 | 449306 |
| chr2 | 159374558 | 159374942 | MLT1H | 3.77E-07 | TANC1 | 450202 |
| chr1 | 149362059 | 149362689 | MER41B | 1.32E-21 | HIST2H2A/ | 450814 |
| chr12 | 102764965 | 102765326 | THE1B | 8.10E-08 | DRAM1 | 451168 |
| chr2 | 202504028 | 202504236 | MLT1I | 2.32E-08 | CASP10 | 451559 |
| chr5 | 40225286 | 40225620 | THE1B | 6.40E-07 | PTGER4 | 453978 |
| chr18 | 72276498 | 72277071 | Tigger3c | 1.98E-71 | TIMM21 | 454088 |
| chr9 | 91764646 | 91765185 | MER44B | 7.01E-16 | GADD45G | 454741 |
| chr11 | 9682649 | 9683160 | MLT1F | 0.000166119 | DENND5A | 457036 |
| chr3 | 57691946 | 57692435 | MLT1K | 1.48E-48 | HESX1 | 459129 |
| chr5 | 40217301 | 40217985 | MER44C | 3.61E-06 | PTGER4 | 461613 |
| chr7 | 134369674 | 134370259 | Tigger3b | 8.25E-13 | AC083862. | 462536 |
| chr4 | 159122558 | 159123088 | MLT1F | 1.71E-11 | C4orf46 | 464741 |
| chr7 | 111655006 | 111655634 | MER41B | 1.89E-09 | LSMEM1 | 465272 |
| chr15 | 70656384 | 70656740 | THE1B | 4.77E-06 | LARP6 | 467121 |

| | | | | | | |
|---|---|---|---|---|---|---|
| chr13 | 33468942 | 33469237 | MLT1I | 2.16E-06 | N4BP2L1 | 469883 |
| chr6 | 71521621 | 71521865 | MER44B | 8.56E-23 | OGFRL1 | 476639 |
| chr7 | 107086872 | 107087297 | MLT1F | 4.77E-06 | LAMB1 | 476945 |
| chr13 | 78003686 | 78004041 | THE1B | 5.98E-29 | IRG1 | 477074 |
| chr18 | 11925092 | 11925346 | Tigger3 | 5.23E-24 | SLMO1 | 482547 |
| chr3 | 186600403 | 186600858 | MER44B | 2.66E-32 | RTP4 | 485260 |
| chr2 | 61598208 | 61599179 | Tigger3b | 8.18E-08 | REL | 489417 |
| chr4 | 39321395 | 39321663 | MER44B | 1.06E-09 | TLR6 | 490574 |
| chr2 | 144167155 | 144167505 | Tigger3a | 1.11E-11 | KYNU | 490907 |
| chr15 | 55627216 | 55627462 | Tigger3a | 1.04E-12 | NEDD4 | 491656 |
| chr5 | 171702051 | 171702389 | THE1C | 8.43E-15 | DUSP1 | 492702 |
| chr3 | 16946136 | 16946512 | THE1C | 9.58E-09 | RFTN1 | 495156 |
| chr11 | 6226085 | 6226427 | Tigger3a | 1.42E-23 | TRIM22 | 496647 |
| chr9 | 107713202 | 107713390 | Tigger3b | 7.59E-67 | FSD1L | 496685 |
| chr8 | 59480164 | 59481337 | Tigger3b | 5.81E-21 | FAM110B | 497287 |
| chr9 | 107710729 | 107710996 | Tigger3b | 1.07E-07 | FSD1L | 499079 |
| chr2 | 55360932 | 55361440 | MER41B | 7.14E-24 | PNPT1 | 499958 |
| chr16 | 24011193 | 24011482 | MLT1H | 0.000208536 | GGA2 | 504529 |
| chr7 | 32545842 | 32546242 | MER44B | 8.27E-78 | NT5C3A | 507498 |
| chr20 | 36986203 | 36986835 | MER41B | 6.59E-16 | CTNNBL1 | 509943 |
| chr5 | 115367825 | 115367904 | MER44D | 3.41E-07 | FEM1C | 511217 |
| chr9 | 3304492 | 3304842 | THE1B | 1.43E-09 | GLIS3 | 519283 |
| chr1 | 224522131 | 224522488 | THE1B | 3.17E-07 | TP53BP2 | 520253 |
| chr11 | 16886034 | 16886397 | THE1B | 2.22E-28 | KCNJ11 | 521007 |
| chr6 | 27686792 | 27687140 | THE1B | 2.03E-10 | ZKSCAN4 | 525259 |
| chr5 | 57070069 | 57070616 | Tigger3c | 2.82E-19 | GPBP1 | 525380 |
| chr12 | 46943716 | 46943902 | MER44B | 5.72E-15 | AMIGO2 | 525586 |
| chr12 | 58731260 | 58731633 | THE1C | 1.07E-07 | AVIL | 526576 |
| chr11 | 61137004 | 61137430 | MLT1H | 3.86E-06 | RAB3IL1 | 527341 |
| chr6 | 17591533 | 17592003 | MLT1K | 1.62E-13 | NHLRC1 | 528713 |
| chr1 | 117105129 | 117105486 | THE1B | 1.02E-08 | SLC22A15 | 529831 |
| chr1 | 178535424 | 178535792 | MLT1H | 7.63E-100 | ABL2 | 532668 |
| chr5 | 130873915 | 130874212 | Tigger3a | 3.55E-83 | CSF2 | 535269 |
| chr12 | 32012595 | 32012964 | THE1C | 8.79E-37 | AC024940. | 535345 |
| chr7 | 140637528 | 140637884 | THE1B | 3.89E-18 | SLC37A3 | 537392 |
| chr10 | 31274814 | 31275143 | Tigger3a | 1.48E-66 | MAP3K8 | 546708 |
| chr6 | 71450450 | 71450807 | THE1B | 2.30E-90 | OGFRL1 | 547697 |
| chr15 | 64557473 | 64557834 | THE1B | 1.06E-09 | PIF1 | 549995 |
| chr1 | 8800236 | 8801283 | Tigger3b | 0.000208536 | SPSB1 | 551654 |
| chr14 | 68701105 | 68702021 | Tigger3b | 9.58E-09 | ZFP36L1 | 552354 |
| chr1 | 222237942 | 222238295 | THE1B | 2.82E-19 | MIA3 | 553131 |
| chr8 | 24092526 | 24092891 | THE1B | 1.89E-19 | NKX3-1 | 554025 |
| chrX | 119449774 | 119451029 | Tigger3b | 0.000126789 | SOWAHD | 557198 |
| chr18 | 22094389 | 22094602 | MLT1I | 1.74E-21 | LAMA3 | 564532 |
| chr2 | 48347914 | 48348243 | MLT1I | 9.01E-08 | MSH2 | 564916 |
| chr1 | 181625003 | 181626185 | Tigger3b | 0.000208536 | IER5 | 567365 |
| chr16 | 27935934 | 27936193 | MLT1L | 3.25E-25 | IL4R | 569265 |

| | | | | | | |
|---|---|---|---|---|---|---|
| chr20 | 10045746 | 10046138 | MLT1K | 1.11E-11 | JAG1 | 572192 |
| chr3 | 171183740 | 171184023 | MLT1F | 8.10E-08 | FNDC3B | 573393 |
| chr10 | 74607400 | 74607857 | MLT1F | 1.33E-06 | DDIT4 | 573623 |
| chr6 | 71419640 | 71419913 | Tigger3b | 3.33E-31 | OGFRL1 | 578591 |
| chr12 | 26797368 | 26797731 | THE1B | 0.000208536 | RASSF8 | 579407 |
| chr6 | 34173307 | 34173600 | MLT1L | 3.81E-08 | ITPR3 | 584146 |
| chr11 | 44551879 | 44552510 | MER41B | 6.06E-23 | C11orf96 | 587824 |
| chr7 | 26998572 | 26998923 | MER44D | 9.35E-37 | SNX10 | 588564 |
| chr7 | 35595497 | 35595787 | Tigger3a | 0.000103369 | EEPD1 | 596969 |
| chr9 | 118453366 | 118453955 | MLT1L | 1.12E-10 | TNC | 600344 |
| chr4 | 2693818 | 2694170 | THE1C | 8.01E-15 | RGS12 | 600583 |
| chr12 | 58806169 | 58806707 | MER41B | 6.60E-10 | AVIL | 601485 |
| chr12 | 58807823 | 58807914 | MLT1I | 8.10E-08 | AVIL | 603139 |
| chr7 | 22157027 | 22157335 | MLT1I | 2.48E-25 | IL6 | 608166 |
| chr9 | 114903001 | 114903599 | Tigger3c | 9.35E-14 | SNX30 | 609517 |
| chr14 | 56263240 | 56263600 | THE1B | 0.000450516 | DLGAP5 | 612916 |
| chr1 | 226122537 | 226122753 | Tigger3b | 9.35E-14 | C1orf95 | 613746 |
| chr2 | 75397224 | 75397587 | THE1B | 2.61E-12 | DOK1 | 614891 |
| chr22 | 33419167 | 33419526 | THE1B | 5.54E-10 | RTCB | 615301 |
| chr1 | 226120646 | 226121161 | Tigger3b | 9.58E-09 | C1orf95 | 615338 |
| chr6 | 13500542 | 13500857 | MLT1F | 1.44E-66 | CD83 | 617013 |
| chr7 | 37105464 | 37105817 | THE1B | 6.82E-70 | GPR141 | 617580 |
| chr6 | 148449067 | 148449518 | MER44B | 1.06E-09 | UST | 618944 |
| chr1 | 222171932 | 222172302 | THE1C | 1.69E-29 | MIA3 | 619124 |
| chr4 | 40829838 | 40830200 | THE1B | 1.32E-21 | RHOH | 627873 |
| chr14 | 65636383 | 65636533 | MLT1K | 2.11E-56 | HSPA2 | 628924 |
| chr12 | 66059473 | 66059830 | Tigger3a | 1.04E-12 | HELB | 636493 |
| chr2 | 167267636 | 167268318 | MER44C | 1.57E-13 | GALNT3 | 640460 |
| chr10 | 100082876 | 100083409 | MLT1H | 0.000166652 | AVPI1 | 645695 |
| chr10 | 93704559 | 93704893 | Tigger3a | 1.98E-71 | KIF11 | 648148 |
| chr6 | 137539807 | 137540099 | Tigger3a | 1.74E-36 | TNFAIP3 | 648250 |
| chr1 | 32701665 | 32702098 | MER44D | 3.99E-110 | HPCA | 649495 |
| chr8 | 119918964 | 119919321 | THE1B | 6.98E-14 | ENPP2 | 650003 |
| chr8 | 59637162 | 59637525 | THE1B | 1.05E-56 | FAM110B | 654285 |
| chr12 | 30775475 | 30775959 | MLT1F | 4.35E-25 | FAM60A | 657557 |
| chr2 | 240753682 | 240754017 | THE1B | 4.24E-26 | ANKMY1 | 664820 |
| chr7 | 115644880 | 115645224 | THE1B | 1.09E-27 | MET | 667218 |
| chr1 | 110539282 | 110539461 | Tigger3b | 1.32E-21 | KCNA3 | 674847 |
| chr3 | 120042598 | 120042941 | THE1B | 4.29E-11 | POPDC2 | 675310 |
| chr12 | 65200822 | 65201168 | THE1C | 9.35E-37 | SRGAP1 | 679405 |
| chr21 | 45191482 | 45191852 | THE1C | 1.06E-09 | LRRC3 | 683515 |
| chr6 | 12976762 | 12977203 | MER44B | 1.98E-20 | EDN1 | 686166 |
| chr6 | 168256565 | 168257096 | MER44B | 5.01E-42 | GPR31 | 686806 |
| chr18 | 53940782 | 53941149 | THE1B | 8.44E-33 | TCF4 | 688234 |
| chr6 | 150075703 | 150076160 | MER44B | 1.14E-71 | UST | 688391 |
| chr15 | 99822643 | 99822995 | Tigger3a | 5.98E-23 | ADAMTS17 | 688797 |
| chr8 | 6038171 | 6038426 | MLT1L | 7.90E-18 | DEFB1 | 689669 |

| | | | | | | |
|---|---|---|---|---|---|---|
| chr1 | 70622345 | 70622633 | MER44B | 3.21E-30 | PTGER3 | 695401 |
| chr2 | 9878517 | 9878968 | MLT1L | 4.46E-41 | ODC1 | 701124 |
| chr2 | 219580183 | 219580353 | Tigger3a | 2.61E-08 | DES | 702744 |
| chr2 | 114604714 | 114605109 | MLT1H | 2.88E-16 | IL1RN | 719561 |
| chr1 | 166878211 | 166878758 | MLT1K | 1.91E-10 | RCSD1 | 720570 |
| chr3 | 157878892 | 157879247 | MLT1I | 9.05E-53 | PTX3 | 724314 |
| chr1 | 94269884 | 94270329 | MER44B | 6.40E-07 | F3 | 724450 |
| chr20 | 1944637 | 1945268 | MER41B | 8.43E-15 | EBF4 | 728254 |
| chr6 | 83802284 | 83802739 | MER44B | 1.23E-137 | TPBG | 728323 |
| chr5 | 159471241 | 159471874 | MER41B | 8.10E-08 | IL12B | 729450 |
| chr7 | 95956062 | 95956445 | THE1C | 4.68E-22 | PDK4 | 731979 |
| chr15 | 52088706 | 52089340 | MER44D | 0.000125381 | TNFAIP8L3 | 739911 |
| chr5 | 175863711 | 175864398 | MER44C | 3.07E-37 | HRH2 | 755247 |
| chr3 | 4467405 | 4467807 | THE1B | 2.77E-12 | EDEM1 | 761522 |
| chr10 | 116438688 | 116438887 | MLT1H | 1.11E-11 | AL162407. | 764158 |
| chr6 | 34354417 | 34354610 | Tigger3b | 2.07E-19 | ITPR3 | 765256 |
| chr7 | 45774846 | 45775559 | Tigger3 | 1.12E-10 | MYO1G | 766724 |
| chr1 | 94220111 | 94220905 | Tigger3b | 1.12E-10 | F3 | 773874 |
| chr1 | 22374913 | 22375429 | MLT1K | 4.92E-06 | ECE1 | 775585 |
| chr8 | 47870469 | 47871015 | Tigger3b | 4.94E-13 | CEBPD | 778454 |
| chr15 | 57097862 | 57098378 | MER44C | 0.000126789 | GCOM1 | 785726 |
| chr8 | 38970743 | 38970806 | MLT1H | 2.21E-09 | IDO1 | 788986 |
| chr1 | 179895470 | 179895521 | Tigger3c | 1.11E-11 | ABL2 | 795123 |
| chr5 | 119488207 | 119488915 | MER44C | 2.18E-94 | TNFAIP8 | 796477 |
| chr13 | 108106590 | 108106968 | THE1C | 1.12E-10 | TNFSF13B | 796618 |
| chr5 | 173282459 | 173282578 | MLT1L | 4.77E-06 | CREBRF | 798062 |
| chr10 | 12342564 | 12342858 | THE1B | 1.05E-76 | OPTN | 798589 |
| chr5 | 77176576 | 77176796 | MLT1I | 6.40E-07 | ZBED3 | 801710 |
| chr1 | 211934855 | 211935026 | MLT1K | 2.99E-13 | ATF3 | 803648 |
| chr11 | 127521094 | 127521233 | MLT1H | 5.30E-17 | ETS1 | 807421 |
| chr2 | 114708345 | 114708719 | THE1B | 3.49E-56 | IL1RN | 823192 |
| chr10 | 93528037 | 93528279 | Tigger3c | 2.03E-10 | KIF11 | 824762 |
| chr1 | 64786163 | 64786615 | MLT1F | 9.01E-07 | AK4 | 826615 |
| chr11 | 114997176 | 114997522 | THE1B | 9.58E-09 | NNMT | 828303 |
| chr5 | 79812664 | 79813203 | MER44B | 2.00E-36 | PAPD4 | 835189 |
| chr6 | 83909212 | 83909566 | THE1B | 1.13E-06 | TPBG | 835251 |
| chr1 | 221949851 | 221950143 | MER41B | 8.22E-17 | MIA3 | 841283 |
| chr6 | 158215079 | 158215444 | THE1B | 1.11E-11 | DYNLT1 | 842060 |
| chr1 | 203527617 | 203527743 | MLT1K | 4.17E-34 | PPP1R15B | 844770 |
| chr5 | 159592669 | 159593059 | MLT1K | 0.000269873 | IL12B | 850878 |
| chr8 | 96129814 | 96129962 | MLT1I | 7.08E-12 | GEM | 857363 |
| chr5 | 127108175 | 127108582 | MLT1L | 3.27E-05 | 03-Mar | 857552 |
| chr4 | 153407033 | 153408159 | Tigger3b | 2.25E-64 | MND1 | 857640 |
| chr18 | 9494536 | 9495166 | MER41B | 1.60E-74 | RAB12 | 861679 |
| chr5 | 67342275 | 67342514 | MER44C | 9.84E-36 | CD180 | 861925 |
| chr3 | 31514327 | 31514689 | THE1B | 1.07E-22 | TGFBR2 | 866234 |
| chr2 | 46757770 | 46758135 | THE1B | 5.36E-24 | MSH2 | 871971 |

| | | | | | | |
|---|---|---|---|---|---|---|
| chr4 | 153388256 | 153388620 | THE1C | 1.89E-19 | MND1 | 877179 |
| chr10 | 89702037 | 89703160 | Tigger3b | 3.27E-05 | ANKRD22 | 878727 |
| chr5 | 37568829 | 37569375 | MLT1H | 4.77E-06 | SLC1A3 | 888380 |
| chr14 | 78455295 | 78455846 | MLT1H | 6.06E-23 | CIPC | 889776 |
| chr18 | 54142466 | 54143137 | MER44D | 5.50E-14 | TCF4 | 889918 |
| chr1 | 176020471 | 176021507 | Tigger3b | 2.86E-38 | KIAA0040 | 890291 |
| chr15 | 49015537 | 49015796 | MLT1L | 8.40E-12 | DTWD1 | 897379 |
| chr2 | 30407975 | 30408320 | Tigger3a | 2.47E-18 | ALK | 898699 |
| chr14 | 70163515 | 70164158 | MER41B | 5.22E-15 | ZFP36L1 | 902051 |
| chr1 | 115613689 | 115614146 | MLT1L | 6.40E-07 | SLC22A15 | 904971 |
| chr1 | 172089227 | 172089588 | THE1B | 0.000208536 | TNFSF18 | 919510 |
| chr11 | 129700841 | 129701517 | MER44C | 6.40E-07 | KCNJ5 | 919762 |
| chr4 | 79450844 | 79451112 | MER44B | 1.12E-10 | CXCL13 | 923858 |
| chr13 | 96180369 | 96180731 | THE1B | 2.07E-19 | GPR180 | 926212 |
| chr8 | 92936737 | 92937248 | MER41B | 8.10E-08 | TMEM55A | 928698 |
| chr14 | 106875368 | 106875712 | Tigger3a | 6.40E-07 | CRIP2 | 930439 |
| chr14 | 70195221 | 70195661 | MER41B | 0.000208536 | ZFP36L1 | 933757 |
| chr12 | 106682040 | 106682577 | MER44B | 5.31E-06 | C12orf75 | 934753 |
| chr7 | 152041392 | 152041640 | MER44B | 1.51E-85 | WDR86 | 935333 |
| chr2 | 153160280 | 153160910 | MER41B | 2.16E-06 | TNFAIP6 | 937624 |
| chr4 | 186628763 | 186629113 | THE1B | 1.96E-20 | ACSL1 | 939281 |
| chr15 | 48972264 | 48973356 | Tigger3b | 2.00E-35 | DTWD1 | 939819 |
| chr2 | 100404375 | 100405564 | Tigger3b | 1.43E-09 | KIAA1211L | 941181 |
| chr2 | 129343211 | 129343986 | Tigger3b | 1.20E-05 | LIMS2 | 943511 |
| chr6 | 137239875 | 137240009 | MLT1F | 0.000208536 | TNFAIP3 | 948340 |
| chr1 | 211779829 | 211779964 | MLT1L | 4.52E-06 | ATF3 | 958710 |
| chrX | 44774761 | 44775458 | MER44C | 8.10E-08 | NDP | 965833 |
| chr3 | 186119335 | 186119687 | THE1B | 4.17E-34 | RTP4 | 966431 |
| chr3 | 59253983 | 59254491 | MLT1H | 1.96E-20 | ABHD6 | 973893 |
| chr13 | 76542121 | 76542467 | Tigger3a | 7.56E-17 | IRG1 | 980163 |
| chr3 | 128166399 | 128166875 | MLT1F | 1.14E-08 | MBD4 | 982910 |
| chr3 | 127310960 | 127311131 | MLT1I | 2.28E-08 | TXNRD3 | 984606 |
| chr9 | 111239909 | 111240244 | Tigger3a | 4.75E-21 | KLF4 | 988639 |
| chr5 | 59056800 | 59057887 | Tigger3b | 1.05E-27 | ELOVL7 | 989729 |
| chr8 | 109258310 | 109258510 | MER44B | 2.56E-47 | NUDCD1 | 994636 |
| chr12 | 25108302 | 25108675 | THE1B | 1.67E-98 | RASSF8 | 1003285 |
| chr14 | 93193797 | 93193855 | MLT1L | 5.28E-09 | CATSPERB | 1004316 |
| chr5 | 138021155 | 138021244 | MLT1I | 5.36E-24 | CXXC5 | 1005638 |
| chr9 | 114505557 | 114506109 | MER44B | 8.10E-08 | SNX30 | 1007007 |
| chr11 | 85492103 | 85492482 | THE1C | 2.12E-07 | PRSS23 | 1009617 |
| chr5 | 171184826 | 171185374 | MLT1K | 5.95E-25 | DUSP1 | 1009717 |
| chr18 | 7596115 | 7596382 | MER44B | 6.40E-07 | RAB12 | 1013059 |
| chr10 | 80876631 | 80877087 | MLT1H | 3.00E-08 | PLAC9 | 1014349 |
| chr13 | 45899836 | 45900328 | MLT1H | 1.21E-16 | KIAA0226L | 1015809 |
| chr4 | 157855257 | 157856136 | Tigger3b | 1.66E-31 | TDO2 | 1019764 |
| chr4 | 79548362 | 79548602 | MLT1K | 3.14E-11 | CXCL13 | 1021376 |
| chr4 | 88155603 | 88156674 | Tigger3b | 5.67E-07 | PPM1K | 1022096 |

| | | | | | | |
|---|---|---|---|---|---|---|
| chr4 | 184269988 | 184270609 | MER41B | 2.01E-55 | IRF2 | 1038256 |
| chr4 | 121009786 | 121010166 | MLT1F | 9.59E-09 | TNIP3 | 1042395 |
| chr6 | 145520948 | 145521320 | Tigger3a | 3.52E-19 | STX11 | 1049285 |
| chr6 | 4901950 | 4902236 | MLT1K | 1.28E-16 | FAM50B | 1052328 |
| chr17 | 10121202 | 10121555 | MLT1F | 5.01E-06 | NTN1 | 1054950 |
| chr3 | 17520152 | 17520366 | MLT1H | 1.27E-44 | RFTN1 | 1069172 |
| chr17 | 58686803 | 58687261 | MER44B | 1.96E-20 | BRIP1 | 1071364 |
| chr1 | 176210219 | 176210932 | MER44D | 1.04E-68 | KIAA0040 | 1080039 |
| chr12 | 95049333 | 95049760 | MLT1K | 6.23E-08 | SOCS2 | 1080499 |
| chr2 | 161161663 | 161162118 | Tigger3b | 2.07E-19 | TANC1 | 1080759 |
| chr20 | 9536270 | 9536793 | MLT1H | 1.89E-19 | JAG1 | 1081537 |
| chr13 | 45827633 | 45827928 | MER44B | 6.35E-26 | KIAA0226L | 1088209 |
| chr18 | 44689374 | 44689932 | MER44B | 3.64E-166 | PSTPIP2 | 1098648 |
| chr11 | 87750966 | 87751322 | THE1B | 1.07E-54 | PRSS23 | 1099138 |
| chr13 | 96365138 | 96365448 | THE1C | 8.77E-12 | GPR180 | 1110981 |
| chr16 | 70839714 | 70840038 | MER44B | 4.65E-06 | NFAT5 | 1115267 |
| chr2 | 169214672 | 169214862 | MLT1K | 5.17E-17 | BBS5 | 1120824 |
| chr14 | 93312953 | 93313252 | MLT1I | 1.05E-19 | CATSPERB | 1123472 |
| chr11 | 129911866 | 129912146 | Tigger3 | 1.33E-35 | KCNJ5 | 1130787 |
| chr12 | 95100114 | 95100469 | THE1B | 1.85E-12 | SOCS2 | 1131280 |
| chr14 | 89730517 | 89730883 | THE1B | 3.21E-30 | CALM1 | 1131961 |
| chr13 | 45783306 | 45783562 | Tigger3a | 3.70E-43 | KIAA0226L | 1132575 |
| chr2 | 46496184 | 46496485 | MLT1I | 9.58E-09 | MSH2 | 1133621 |
| chr5 | 54095318 | 54095641 | Tigger3a | 4.77E-06 | IL6ST | 1135280 |
| chr4 | 79685954 | 79686210 | MER44B | 8.13E-13 | ANTXR2 | 1136091 |
| chr5 | 7911379 | 7911745 | THE1B | 3.27E-05 | PAPD7 | 1160886 |
| chr7 | 77235910 | 77236362 | MER44B | 2.13E-19 | ZP3 | 1166695 |
| chr14 | 107118966 | 107119320 | THE1B | 2.29E-68 | CRIP2 | 1174037 |
| chr2 | 46451892 | 46452395 | MLT1F | 4.09E-16 | MSH2 | 1177711 |
| chr5 | 80163118 | 80163382 | MLT1L | 1.91E-26 | PAPD4 | 1185643 |
| chr8 | 64314239 | 64314662 | MLT1L | 1.06E-09 | CYP7B1 | 1185656 |
| chr12 | 95155392 | 95156551 | Tigger3b | 8.22E-17 | SOCS2 | 1186558 |
| chr3 | 127527084 | 127527534 | MLT1K | 5.36E-24 | TXNRD3 | 1200730 |
| chr5 | 170989789 | 170990167 | THE1C | 2.88E-16 | DUSP1 | 1204924 |
| chr8 | 121838628 | 121838991 | THE1B | 6.37E-10 | ENPP2 | 1210119 |
| chr4 | 85735537 | 85736076 | MER44B | 6.50E-08 | AGPAT9 | 1215855 |
| chr8 | 117587213 | 117587418 | MLT1I | 5.75E-61 | EXT1 | 1219309 |
| chr13 | 34219154 | 34219636 | MLT1H | 1.13E-06 | N4BP2L1 | 1220095 |
| chr8 | 117583699 | 117584106 | MLT1H | 1.05E-89 | EXT1 | 1222621 |
| chr3 | 150317226 | 150317869 | MER41B | 5.32E-25 | TM4SF1 | 1227667 |
| chr9 | 80304916 | 80305279 | THE1B | 1.96E-20 | GCNT1 | 1230770 |
| chr12 | 95200539 | 95200933 | Tigger3a | 9.58E-09 | SOCS2 | 1231705 |
| chr12 | 24877833 | 24878196 | THE1C | 5.74E-58 | RASSF8 | 1233764 |
| chr17 | 57877969 | 57878160 | Tigger3b | 6.40E-07 | TEX14 | 1234856 |
| chr15 | 52598622 | 52598941 | Tigger3a | 2.96E-19 | TNFAIP8L3 | 1249827 |
| chr12 | 69798953 | 69799660 | MER44C | 1.43E-09 | IFNG | 1250405 |
| chr8 | 98162623 | 98162978 | THE1B | 9.58E-09 | STK3 | 1250651 |

| | | | | | | |
|---|---|---|---|---|---|---|
| chr2 | 18004912 | 18005244 | THE1B | 3.57E-26 | FAM49A | 1259620 |
| chr14 | 60392528 | 60392886 | THE1C | 4.77E-06 | PRKCH | 1261389 |
| chr2 | 62374460 | 62375275 | Tigger3b | 2.46E-72 | REL | 1265669 |
| chr21 | 34179124 | 34179479 | THE1B | 1.36E-28 | MRPS6 | 1266043 |
| chr1 | 246305403 | 246306075 | MER44C | 3.27E-05 | NLRP3 | 1273381 |
| chr8 | 20490055 | 20490396 | THE1B | 0.000208536 | DOK2 | 1275986 |
| chr4 | 85796459 | 85797440 | Tigger3b | 3.27E-05 | AGPAT9 | 1276777 |
| chr6 | 139476143 | 139476699 | Tigger3c | 1.73E-17 | TNFAIP3 | 1279168 |
| chr11 | 72878400 | 72878518 | MLT1K | 5.78E-05 | KCNE3 | 1287366 |
| chr10 | 69923648 | 69923853 | MLT1L | 1.71E-50 | TSPAN15 | 1287374 |
| chr2 | 33751242 | 33751395 | MLT1K | 9.94E-10 | NLRC4 | 1301717 |
| chr3 | 192548787 | 192549270 | MLT1F | 3.48E-20 | HES1 | 1304662 |
| chr4 | 99432421 | 99432749 | Tigger3a | 1.33E-46 | DAPP1 | 1305239 |
| chr16 | 59004686 | 59005237 | MLT1K | 4.82E-05 | GPR56 | 1317543 |
| chr2 | 161412086 | 161412445 | THE1B | 6.40E-07 | TANC1 | 1331182 |
| chr2 | 153554096 | 153554431 | Tigger3a | 1.09E-27 | TNFAIP6 | 1331440 |
| chr16 | 79299353 | 79299867 | MER44C | 1.32E-07 | CDYL2 | 1331934 |
| chr1 | 45738861 | 45739071 | MLT1F | 2.95E-19 | ARTN | 1337382 |
| chr3 | 33833094 | 33833611 | MLT1H | 3.83E-106 | CMTM7 | 1340599 |
| chr6 | 5190816 | 5190982 | Tigger3 | 7.71E-54 | FAM50B | 1341194 |
| chr2 | 8480964 | 8481323 | THE1B | 1.06E-09 | RNF144A | 1343893 |
| chr7 | 24802641 | 24803324 | MER44C | 3.81E-08 | IGF2BP3 | 1349778 |
| chr12 | 24758281 | 24758686 | MLT1I | 1.11E-11 | RASSF8 | 1353274 |
| chr7 | 114950347 | 114950701 | THE1B | 1.32E-93 | MET | 1361741 |
| chr7 | 3954945 | 3955318 | THE1B | 5.98E-29 | BRAT1 | 1373130 |
| chr1 | 170893894 | 170894252 | THE1B | 1.53E-16 | F5 | 1378167 |
| chr5 | 61834548 | 61834663 | Tigger3c | 1.60E-60 | SMIM15 | 1378799 |
| chr6 | 127619741 | 127620024 | Tigger3b | 1.32E-06 | NCOA7 | 1379290 |
| chr12 | 75869849 | 75870155 | THE1B | 7.44E-52 | CSRP2 | 1382338 |
| chr8 | 66893145 | 66893521 | THE1B | 2.10E-06 | CYP7B1 | 1384453 |
| chr6 | 135968670 | 135969029 | THE1B | 4.77E-06 | SGK1 | 1385494 |
| chr13 | 31501404 | 31501763 | THE1C | 0.00012044 | BRCA2 | 1387846 |
| chr2 | 153624477 | 153624831 | THE1C | 6.62E-14 | TNFAIP6 | 1401821 |
| chr4 | 183896738 | 183897307 | MER44B | 3.33E-09 | IRF2 | 1411558 |
| chr12 | 28932919 | 28933620 | MER44D | 5.21E-47 | ARNTL2 | 1411650 |
| chr9 | 106787528 | 106787862 | Tigger3a | 7.90E-18 | FSD1L | 1422213 |
| chr3 | 185525369 | 185525783 | MLT1F | 4.65E-103 | CHRD | 1422793 |
| chr8 | 69155791 | 69155979 | MLT1K | 2.78E-11 | SLCO5A1 | 1423301 |
| chr16 | 82142840 | 82143207 | THE1B | 3.89E-18 | CDYL2 | 1424339 |
| chr6 | 141647318 | 141647999 | MER44D | 9.20E-107 | HIVEP2 | 1424603 |
| chr2 | 110451042 | 110451326 | THE1B | 1.51E-09 | BCL2L11 | 1425627 |
| chr8 | 64067540 | 64068162 | MER41B | 1.50E-17 | CYP7B1 | 1432156 |
| chr2 | 145114029 | 145114323 | MLT1I | 5.12E-07 | KYNU | 1437781 |
| chr2 | 9140834 | 9141679 | Tigger3b | 4.79E-05 | ODC1 | 1438413 |
| chr6 | 141628733 | 141629011 | THE1B | 8.10E-08 | HIVEP2 | 1443591 |
| chr18 | 10094341 | 10094861 | Tigger3b | 2.37E-46 | RAB12 | 1461484 |
| chr1 | 76892266 | 76892604 | Tigger3b | 2.87E-12 | NEXN | 1461592 |

| | | | | | | |
|---|---|---|---|---|---|---|
| chr11 | 7698296 | 7698675 | THE1C | 2.91E-17 | DENND5A | 1461695 |
| chr11 | 30364344 | 30364828 | MLT1K | 1.72E-09 | RCN1 | 1469109 |
| chr10 | 11667180 | 11667579 | MLT1I | 1.13E-17 | OPTN | 1473868 |
| chr11 | 88131388 | 88131901 | MER44B | 1.55E-30 | PRSS23 | 1479560 |
| chr5 | 18697855 | 18698184 | THE1B | 3.11E-84 | BASP1 | 1480186 |
| chr4 | 9902247 | 9902620 | THE1C | 3.15E-05 | ACOX3 | 1490238 |
| chr8 | 108762004 | 108762192 | MLT1I | 0.000136433 | NUDCD1 | 1490954 |
| chr8 | 93503120 | 93503291 | MLT1K | 1.89E-10 | TMEM55A | 1495081 |
| chr3 | 114500498 | 114500854 | THE1B | 5.42E-05 | BOC | 1497091 |
| chr8 | 101465015 | 101465393 | Tigger3 | 8.01E-15 | OSR2 | 1504395 |
| chr5 | 160251479 | 160251815 | Tigger3a | 9.21E-11 | IL12B | 1509688 |
| chr6 | 26351657 | 26352329 | MER44D | 5.49E-44 | FAM65B | 1511104 |
| chr20 | 12133481 | 12134006 | MER44B | 7.61E-44 | JAG1 | 1515074 |
| chr10 | 25211296 | 25211597 | THE1B | 3.62E-12 | APBB1IP | 1515533 |
| chr5 | 108889628 | 108889992 | THE1B | 0.000208536 | TSLP | 1515766 |
| chr17 | 58159968 | 58160458 | MER44B | 2.43E-19 | TEX14 | 1516855 |
| chr16 | 17405540 | 17405902 | THE1B | 1.04E-12 | MYH11 | 1526668 |
| chr3 | 177536676 | 177536981 | MLT1L | 1.04E-12 | MFN1 | 1528497 |
| chr2 | 40629109 | 40629711 | MER41B | 9.58E-09 | DHX57 | 1540181 |
| chr8 | 122252778 | 122253055 | MLT1H | 2.07E-10 | ZHX2 | 1540576 |
| chr9 | 80629734 | 80630315 | Tigger3c | 8.11E-05 | GCNT1 | 1555588 |
| chrX | 39986339 | 39986784 | MLT1H | 7.23E-71 | GPR34 | 1561440 |
| chr7 | 65715566 | 65716087 | MER44B | 2.01E-99 | ZNF107 | 1565833 |
| chr1 | 31778744 | 31779482 | MER44C | 9.08E-32 | HPCA | 1572111 |
| chr3 | 40797042 | 40797206 | THE1C | 5.17E-17 | XIRP1 | 1572335 |
| chr16 | 82294056 | 82294418 | THE1B | 0.00012044 | CDYL2 | 1575555 |
| chr11 | 107526314 | 107526600 | MER41B | 4.71E-08 | KBTBD3 | 1579583 |
| chr3 | 151294317 | 151294829 | MER41B | 1.04E-12 | RAP2B | 1585198 |
| chr3 | 40809947 | 40810474 | MLT1K | 4.91E-23 | XIRP1 | 1585240 |
| chr11 | 58967466 | 58967827 | THE1B | 9.80E-16 | SERPING1 | 1588762 |
| chr5 | 142642019 | 142642268 | MER44C | 5.35E-82 | ARAP3 | 1589684 |
| chr9 | 77374893 | 77375472 | Tigger3c | 8.01E-64 | ANXA1 | 1591879 |
| chr11 | 72570160 | 72570699 | MER44B | 6.40E-07 | KCNE3 | 1595185 |
| chr15 | 85227428 | 85227788 | THE1B | 1.32E-21 | HOMER2 | 1600546 |
| chr8 | 97805451 | 97805794 | THE1B | 8.08E-22 | STK3 | 1607835 |
| chr2 | 198817377 | 198817712 | THE1B | 1.89E-19 | HECW2 | 1608950 |
| chr1 | 30531933 | 30532373 | MLT1K | 2.97E-35 | RAB42 | 1612906 |
| chr12 | 124961656 | 124962027 | THE1C | 2.82E-19 | HIP1R | 1615393 |
| chr4 | 144912764 | 144913142 | THE1C | 0.000208536 | MMAA | 1626271 |
| chr11 | 112498392 | 112498894 | MLT1F | 3.53E-42 | NNMT | 1629613 |
| chr3 | 25862079 | 25862184 | MLT1K | 3.17E-33 | THRB | 1630420 |
| chr2 | 133582152 | 133582553 | MLT1I | 2.29E-11 | TMEM163 | 1630775 |
| chr8 | 117174049 | 117174302 | Tigger3c | 4.51E-20 | EXT1 | 1632425 |
| chr9 | 25681306 | 25681564 | MER44B | 2.68E-32 | MOB3B | 1643641 |
| chr14 | 97645926 | 97646300 | THE1C | 0.000126789 | RP11-1070 | 1663453 |
| chr8 | 134805556 | 134805662 | MLT1K | 1.12E-10 | KHDRBS3 | 1664036 |
| chr5 | 142717602 | 142718759 | Tigger3b | 1.12E-10 | ARAP3 | 1665267 |

| | | | | | | |
|---|---|---|---|---|---|---|
| chr14 | 86797250 | 86797583 | Tigger3b | 1.06E-09 | GPR65 | 1673883 |
| chr13 | 71954366 | 71954712 | THE1B | 0.000126789 | KLF5 | 1674400 |
| chr8 | 68892816 | 68893180 | THE1C | 3.61E-06 | SLCO5A1 | 1686100 |
| chr1 | 91425373 | 91426068 | MER44C | 1.14E-08 | GBP5 | 1696992 |
| chr1 | 19835271 | 19835775 | MER41B | 6.53E-09 | ECE1 | 1707963 |
| chr4 | 87470373 | 87470786 | Tigger3c | 1.07E-07 | PPM1K | 1707984 |
| chr2 | 172260610 | 172261164 | MER44D | 2.50E-07 | PHOSPHO2 | 1708886 |
| chr3 | 18166052 | 18166432 | MLT1I | 2.82E-19 | RFTN1 | 1715072 |
| chr2 | 172271283 | 172271643 | THE1B | 9.58E-09 | PHOSPHO2 | 1719559 |
| chrX | 117159188 | 117160009 | MER44C | 8.10E-08 | SOWAHD | 1732565 |
| chr12 | 45730220 | 45731325 | Tigger3b | 1.48E-66 | AMIGO2 | 1738163 |
| chr15 | 43207265 | 43207801 | MER44B | 2.61E-12 | PATL2 | 1750127 |
| chr13 | 65126103 | 65126409 | MLT1L | 3.25E-25 | PCDH9 | 1750556 |
| chr5 | 108652323 | 108652874 | MLT1K | 1.85E-25 | TSLP | 1752884 |
| chr14 | 83630144 | 83630676 | MER44B | 1.06E-09 | STON2 | 1765427 |
| chr8 | 117031898 | 117032192 | MLT1I | 1.11E-11 | EXT1 | 1774535 |
| chr10 | 69428094 | 69428584 | MER44B | 4.32E-59 | TSPAN15 | 1782643 |
| chr8 | 97630308 | 97630655 | Tigger3a | 9.28E-24 | STK3 | 1782974 |
| chr16 | 71508875 | 71509130 | MLT1K | 3.52E-49 | NFAT5 | 1784428 |
| chr7 | 114525798 | 114526134 | MLT1K | 1.05E-11 | MET | 1786308 |
| chrX | 50561832 | 50562173 | Tigger3b | 9.58E-09 | PIM2 | 1790159 |
| chr1 | 93194243 | 93194948 | MER44C | 1.34E-31 | F3 | 1799831 |
| chr16 | 17684968 | 17685603 | MER41B | 1.12E-10 | MYH11 | 1806096 |
| chr12 | 125166079 | 125166103 | MLT1L | 0.000184349 | HIP1R | 1819816 |
| chr15 | 98669869 | 98670138 | MLT1L | 3.77E-07 | ADAMTS17 | 1841654 |
| chr5 | 112269766 | 112270140 | THE1C | 1.89E-09 | TSLP | 1860754 |
| chr12 | 90674471 | 90674846 | THE1C | 2.47E-18 | BTG1 | 1861438 |
| chrX | 102670606 | 102671800 | Tigger3b | 4.94E-13 | ARMCX1 | 1865092 |
| chr12 | 45602998 | 45603298 | MLT1F | 3.02E-09 | AMIGO2 | 1866190 |
| chr8 | 116934793 | 116935172 | MLT1F | 1.31E-05 | EXT1 | 1871555 |
| chr3 | 42924784 | 42925143 | THE1B | 6.40E-07 | KIF15 | 1878064 |
| chr6 | 157178837 | 157179103 | Tigger3c | 3.21E-30 | DYNLT1 | 1878401 |
| chr6 | 51478534 | 51478857 | Tigger3a | 0.00026788 | GCLC | 1883280 |
| chr6 | 140083604 | 140083963 | MLT1I | 5.54E-44 | TNFAIP3 | 1886629 |
| chr3 | 18343318 | 18343681 | THE1B | 0.000208536 | RFTN1 | 1892338 |
| chr14 | 59744593 | 59745300 | MER44D | 1.37E-30 | PRKCH | 1908975 |
| chr11 | 107859341 | 107859668 | Tigger3 | 4.00E-146 | KBTBD3 | 1912610 |
| chr12 | 14988897 | 14989525 | MER44C | 5.72E-18 | GPRC5A | 1926894 |
| chr2 | 206523273 | 206523684 | MLT1K | 1.06E-09 | CD28 | 1951857 |
| chr8 | 68624955 | 68625298 | MER44B | 3.82E-51 | SLCO5A1 | 1953982 |
| chr10 | 11165839 | 11166190 | THE1B | 8.18E-08 | OPTN | 1975257 |
| chr5 | 145227551 | 145227921 | MLT1I | 1.91E-10 | SPINK1 | 1976208 |
| chr8 | 102165222 | 102165431 | MLT1K | 4.93E-06 | C8orf56 | 1979758 |
| chr8 | 129545447 | 129546288 | Tigger3b | 3.89E-18 | FAM84B | 1980387 |
| chr6 | 156734314 | 156734824 | MER44B | 6.40E-07 | CNKSR3 | 1998457 |
| chr20 | 25353273 | 25353487 | Tigger3c | 1.68E-101 | GZF1 | 2003765 |
| chr8 | 97400698 | 97401051 | THE1B | 1.06E-09 | STK3 | 2012578 |

| | | | | | | |
|---|---|---|---|---|---|---|
| chr8 | 97391150 | 97391725 | Tigger3 | 0.000208536 | STK3 | 2021904 |
| chr1 | 92968602 | 92968880 | MLT1I | 3.52E-78 | F3 | 2025899 |
| chr12 | 90500016 | 90500745 | MER44D | 3.89E-18 | BTG1 | 2035539 |
| chr6 | 20423363 | 20423690 | Tigger3a | 3.27E-05 | RNF144B | 2035618 |
| chr2 | 133168257 | 133168618 | THE1B | 3.25E-25 | TMEM163 | 2044710 |
| chr8 | 129609922 | 129610295 | THE1C | 1.67E-11 | FAM84B | 2044862 |
| chr3 | 34813559 | 34813902 | THE1B | 9.58E-09 | TRANK1 | 2054407 |
| chr5 | 143118439 | 143119603 | Tigger3b | 0.000234401 | ARAP3 | 2066104 |
| chr5 | 50007061 | 50007650 | MLT1L | 3.00E-10 | ITGA1 | 2076078 |
| chr3 | 169669747 | 169670068 | Tigger3a | 3.91E-29 | FNDC3B | 2087348 |
| chr1 | 31251332 | 31251687 | MLT1K | 6.31E-12 | HPCA | 2099906 |
| chr5 | 81078332 | 81078952 | Tigger3b | 4.74E-10 | PAPD4 | 2100857 |
| chr1 | 49318591 | 49319112 | MLT1F | 1.17E-09 | CDKN2C | 2107303 |
| chr6 | 152342805 | 152343177 | THE1B | 9.58E-09 | IPCEF1 | 2132452 |
| chr4 | 115335515 | 115335879 | THE1B | 3.94E-05 | TIFA | 2136600 |
| chr3 | 18588283 | 18588562 | MLT1K | 6.59E-16 | RFTN1 | 2137303 |
| chr2 | 12728355 | 12728905 | MER44B | 1.10E-59 | ODC1 | 2143685 |
| chr1 | 239545966 | 239546304 | THE1B | 1.46E-13 | KMO | 2149128 |
| chr8 | 134315537 | 134315771 | MLT1I | 3.28E-49 | KHDRBS3 | 2153927 |
| chr1 | 243897642 | 243898738 | Tigger3b | 1.84E-24 | KMO | 2165707 |
| chr11 | 71974494 | 71974593 | MLT1H | 2.98E-14 | KCNE3 | 2191291 |
| chr14 | 32184555 | 32185800 | Tigger3b | 1.11E-11 | EGLN3 | 2207635 |
| chr1 | 188857553 | 188858250 | MER44D | 6.59E-16 | PTGS2 | 2210151 |
| chr2 | 24773164 | 24773517 | THE1B | 0.000208536 | CENPA | 2213638 |
| chr2 | 68418273 | 68418838 | Tigger3c | 1.05E-56 | TGFA | 2255572 |
| chr3 | 2971434 | 2971582 | MLT1K | 3.25E-25 | EDEM1 | 2257747 |
| chr2 | 24728504 | 24729034 | MLT1K | 4.17E-34 | CENPA | 2258121 |
| chr11 | 70109178 | 70109279 | MLT1L | 9.35E-14 | CHKA | 2270895 |
| chr1 | 46704521 | 46704878 | THE1B | 1.17E-16 | ARTN | 2303042 |
| chr8 | 134148179 | 134148488 | MLT1L | 1.20E-25 | KHDRBS3 | 2321210 |
| chr10 | 108472570 | 108473036 | MLT1L | 1.96E-20 | CCDC147 | 2335957 |
| chr2 | 58324707 | 58325901 | Tigger3b | 1.89E-10 | BCL11A | 2352399 |
| chr1 | 245156095 | 245156573 | MLT1F | 0.000208536 | NLRP3 | 2422883 |
| chr14 | 59204247 | 59205450 | Tigger3b | 2.68E-27 | PRKCH | 2448825 |
| chr6 | 106414400 | 106414782 | MLT1L | 4.86E-59 | FOXO3 | 2466254 |
| chr10 | 33272226 | 33272510 | MLT1L | 3.24E-36 | MAP3K8 | 2544120 |
| chr4 | 10976052 | 10976424 | THE1C | 1.11E-11 | ACOX3 | 2564043 |
| chr14 | 31806014 | 31806346 | Tigger3a | 1.50E-17 | EGLN3 | 2587089 |
| chr2 | 173146754 | 173147178 | Tigger3 | 4.04E-12 | PHOSPHO2 | 2595030 |
| chr2 | 173147764 | 173147869 | Tigger3b | 2.43E-27 | PHOSPHO2 | 2596040 |
| chr1 | 231511872 | 231512382 | MER44B | 6.59E-16 | RHOU | 2629885 |
| chr2 | 109243351 | 109243744 | Tigger3c | 1.08E-07 | BCL2L11 | 2633209 |
| chr3 | 104273276 | 104273642 | THE1B | 8.10E-08 | NFKBIZ | 2697232 |
| chr6 | 106135731 | 106136095 | THE1C | 6.07E-11 | FOXO3 | 2744941 |
| chr3 | 95642292 | 95642912 | MER41B | 1.96E-20 | ST3GAL6 | 2808166 |
| chr1 | 244545246 | 244545380 | MLT1F | 1.10E-31 | KMO | 2813311 |
| chr12 | 80272299 | 80272673 | THE1C | 0.000208536 | E2F7 | 2822641 |

| | | | | | | |
|---|---|---|---|---|---|---|
| chr2 | 67516940 | 67517291 | THE1C | 1.73E-20 | LGALSL | 2835264 |
| chr11 | 111264988 | 111265352 | THE1B | 1.14E-08 | NNMT | 2863155 |
| chr10 | 109375076 | 109377497 | Tigger3 | 1.47E-23 | DUSP5 | 2880097 |
| chr10 | 109375076 | 109377497 | Tigger3 | 9.14E-05 | DUSP5 | 2880097 |
| chr10 | 109375076 | 109377497 | Tigger3 | 5.71E-18 | DUSP5 | 2880097 |
| chr7 | 17478573 | 17478933 | THE1B | 2.82E-19 | ITGB8 | 2891390 |
| chr2 | 108972474 | 108972550 | Tigger3c | 1.96E-20 | BCL2L11 | 2904403 |
| chr10 | 23806333 | 23806647 | MLT1I | 6.60E-10 | APBB1IP | 2920483 |
| chr5 | 122774053 | 122774142 | MLT1K | 6.97E-44 | GRAMD3 | 2921680 |
| chr10 | 23801349 | 23801836 | MER44B | 1.57E-08 | APBB1IP | 2925294 |
| chr8 | 62569826 | 62570178 | Tigger3a | 2.43E-05 | CYP7B1 | 2930140 |
| chr8 | 62548594 | 62548666 | MLT1L | 5.98E-29 | CYP7B1 | 2951652 |
| chrX | 15896424 | 15896784 | THE1B | 1.91E-26 | TLR8 | 2971666 |
| chrX | 15901230 | 15901608 | MER41B | 4.62E-11 | TLR8 | 2976472 |
| chr10 | 33831344 | 33832064 | MER44C | 1.12E-10 | NAMPTL | 2978583 |
| chr7 | 17371639 | 17372846 | Tigger3b | 5.17E-17 | ITGB8 | 2997477 |
| chr4 | 116200301 | 116200644 | Tigger3a | 3.07E-09 | TIFA | 3001386 |
| chr7 | 17362402 | 17362733 | Tigger3a | 1.96E-20 | ITGB8 | 3007590 |
| chr6 | 21438843 | 21438908 | MLT1I | 1.46E-70 | RNF144B | 3051098 |
| chr18 | 32921150 | 32921701 | Tigger3 | 1.31E-08 | GAREM | 3053311 |
| chr5 | 31975242 | 31975603 | THE1B | 8.10E-08 | PRLR | 3073256 |
| chrX | 16003964 | 16004475 | MLT1F | 1.34E-31 | TLR8 | 3079206 |
| chr10 | 23640993 | 23641157 | MLT1I | 1.12E-10 | APBB1IP | 3085973 |
| chr9 | 89100196 | 89100565 | MLT1I | 1.35E-23 | GADD45G | 3119361 |
| chr2 | 187472101 | 187472375 | MLT1I | 4.77E-06 | OSGEPL1 | 3139009 |
| chr2 | 187441277 | 187441752 | MLT1H | 4.95E-56 | OSGEPL1 | 3169632 |
| chr4 | 174406389 | 174406753 | MLT1H | 8.01E-15 | VEGFC | 3197934 |
| chr12 | 22810307 | 22810804 | MLT1K | 8.01E-15 | RASSF8 | 3301156 |
| chr14 | 31076459 | 31076832 | Tigger3a | 1.49E-05 | EGLN3 | 3316603 |
| chr12 | 116882156 | 116882527 | THE1B | 9.35E-18 | RASAL1 | 3328669 |
| chr3 | 84773822 | 84774447 | MER44D | 2.65E-07 | RP11-159G | 3333975 |
| chr2 | 131872547 | 131872903 | THE1B | 7.14E-07 | TMEM163 | 3340425 |
| chr2 | 124260421 | 124260893 | MER44B | 2.85E-13 | EPB41L5 | 3342278 |
| chr1 | 196121233 | 196121664 | MLT1H | 2.78E-11 | RGS2 | 3342960 |
| chr6 | 47452598 | 47453754 | Tigger3b | 3.61E-06 | TMEM63B | 3345149 |
| chr2 | 148738821 | 148739084 | MER44B | 1.06E-09 | RBM43 | 3365368 |
| chr5 | 122066133 | 122066530 | MLT1K | 1.09E-35 | TNFAIP8 | 3374403 |
| chr2 | 173943736 | 173944948 | Tigger3b | 2.27E-05 | PHOSPHO2 | 3392012 |
| chr6 | 56803463 | 56803830 | MLT1H | 4.84E-143 | GCLC | 3416200 |
| chr1 | 197808967 | 197809330 | THE1B | 1.16E-15 | LAD1 | 3533040 |
| chr2 | 208105747 | 208106278 | MLT1H | 5.19E-66 | CD28 | 3534331 |
| chr18 | 33433137 | 33433496 | THE1B | 0.000208536 | GAREM | 3565298 |
| chr4 | 174038592 | 174038959 | THE1B | 1.02E-15 | VEGFC | 3565728 |
| chr3 | 67436869 | 67437220 | THE1B | 1.97E-09 | FOXP1 | 3566622 |
| chr3 | 105150052 | 105150529 | MLT1H | 6.11E-14 | NFKBIZ | 3574008 |
| chr3 | 105155478 | 105155765 | MLT1K | 6.23E-08 | NFKBIZ | 3579434 |
| chr7 | 69336356 | 69336892 | MER44B | 1.73E-53 | MLXIPL | 3670630 |

| | | | | | | |
|---|---|---|---|---|---|---|
| chr10 | 23026585 | 23026864 | MLT1H | 8.79E-19 | APBB1IP | 3700266 |
| chr12 | 81171801 | 81172264 | MLT1H | 2.43E-19 | E2F7 | 3722143 |
| chr5 | 82702876 | 82704056 | Tigger3b | 1.23E-77 | PAPD4 | 3725401 |
| chr5 | 82702876 | 82704056 | Tigger3b | 4.70E-71 | PAPD4 | 3725401 |
| chr18 | 65434156 | 65434468 | Tigger3b | 4.51E-20 | HMSD | 3806740 |
| chr3 | 106921677 | 106922042 | THE1B | 7.05E-08 | PVRL3 | 3866874 |
| chr3 | 167852729 | 167853040 | THE1B | 9.25E-48 | FNDC3B | 3904376 |
| chr11 | 110208193 | 110208779 | MER41B | 3.52E-49 | NNMT | 3919728 |
| chr8 | 13488552 | 13488915 | THE1B | 2.18E-07 | PDGFRL | 3945025 |
| chr11 | 110182941 | 110183290 | THE1C | 4.45E-34 | NNMT | 3945217 |
| chr7 | 120373372 | 120373583 | MLT1I | 5.00E-06 | MET | 3956143 |
| chr6 | 48103178 | 48103695 | MLT1F | 2.67E-17 | TMEM63B | 3995729 |
| chr4 | 164315758 | 164316450 | MER44D | 8.10E-08 | RAPGEF2 | 4042149 |
| chr7 | 123601758 | 123602120 | THE1B | 1.11E-11 | LRRC4 | 4065002 |
| chr18 | 67711689 | 67712215 | MER44B | 1.47E-215 | TIMM21 | 4103529 |
| chr8 | 132300535 | 132301185 | MER44D | 6.40E-07 | KHDRBS3 | 4168513 |
| chr4 | 164472777 | 164473118 | THE1B | 2.78E-05 | RAPGEF2 | 4199168 |
| chr4 | 164488602 | 164488961 | THE1B | 2.78E-11 | RAPGEF2 | 4214993 |
| chr18 | 48661750 | 48662325 | MER44B | 1.98E-20 | TCF4 | 4227235 |
| chr4 | 164519530 | 164520158 | Tigger3b | 1.20E-05 | RAPGEF2 | 4245921 |
| chrX | 123162151 | 123162400 | MER44B | 4.77E-06 | SOWAHD | 4269575 |
| chr2 | 174843280 | 174843574 | MER44C | 2.98E-14 | PHOSPHO2 | 4291556 |
| chr18 | 48596935 | 48597451 | MER44B | 9.58E-09 | TCF4 | 4292109 |
| chr18 | 48565673 | 48565982 | MLT1K | 3.42E-28 | TCF4 | 4323578 |
| chr18 | 34430381 | 34430769 | MER44B | 3.38E-12 | GAREM | 4562542 |
| chr10 | 120248241 | 120248462 | MLT1F | 3.25E-25 | AL162407. | 4573711 |
| chr2 | 175128725 | 175129193 | MER44D | 3.77E-07 | PHOSPHO2 | 4577001 |
| chr5 | 163321419 | 163321959 | MER44B | 1.96E-47 | IL12B | 4579628 |
| chr5 | 164998513 | 164998787 | THE1C | 3.18E-15 | C5orf58 | 4660662 |
| chr5 | 163429736 | 163430317 | MLT1K | 1.09E-27 | IL12B | 4687945 |
| chr13 | 62105963 | 62106253 | Tigger3a | 8.95E-14 | PCDH9 | 4770712 |
| chr10 | 120553127 | 120553548 | MLT1H | 3.20E-09 | AL162407. | 4878597 |
| chr10 | 120715139 | 120715684 | MLT1K | 1.69E-29 | AL162407. | 5040609 |
| chr21 | 25358586 | 25358956 | THE1B | 6.40E-07 | MAP3K7CL | 5090834 |
| chr10 | 121161352 | 121161643 | MLT1I | 4.62E-10 | FAM53B | 5146216 |
| chrX | 77005115 | 77005806 | MER44D | 1.04E-12 | NHSL2 | 5651115 |
| chr18 | 37418702 | 37419046 | Tigger3a | 0.00012044 | EPG5 | 6008526 |
| chr9 | 85341356 | 85341973 | MER41B | 2.45E-12 | GCNT1 | 6267210 |
| chr9 | 85803975 | 85804348 | THE1C | 8.10E-09 | GADD45G | 6415578 |
| chr2 | 183897090 | 183897403 | Tigger3a | 4.62E-10 | OSGEPL1 | 6713981 |
| chr13 | 60115402 | 60115737 | Tigger3a | 4.77E-06 | PCDH9 | 6761228 |
| chrX | 19728181 | 19728900 | MER44D | 9.71E-31 | TLR8 | 6803423 |
| chr12 | 85410042 | 85410564 | MER44B | 5.49E-26 | BTG1 | 7125720 |
| chr2 | 177789975 | 177790229 | MLT1L | 9.58E-09 | PHOSPHO2 | 7238251 |
| chr5 | 86409730 | 86410279 | MER44B | 2.63E-12 | PAPD4 | 7432255 |
| chr6 | 101256595 | 101257333 | Tigger3b | 3.45E-06 | FOXO3 | 7623703 |
| chr2 | 178181250 | 178181599 | Tigger3a | 3.27E-05 | PHOSPHO2 | 7629526 |

| | | | | | | |
|---|---|---|---|---|---|---|
| chr5 | 102194541 | 102195039 | Tigger3b | 4.83E-24 | TSLP | 8210719 |
| chr4 | 48728289 | 48728396 | MLT1I | 4.94E-13 | RHOH | 8526324 |
| chr2 | 179102850 | 179103547 | MER44C | 1.11E-11 | PHOSPHO2 | 8551126 |
| chr2 | 181902694 | 181903184 | Tigger3b | 3.91E-08 | OSGEPL1 | 8708200 |
| chr2 | 181901734 | 181902425 | Tigger3b | 3.48E-07 | OSGEPL1 | 8708959 |
| chr2 | 181802230 | 181802604 | THE1C | 2.16E-06 | OSGEPL1 | 8808780 |
| chr6 | 62981095 | 62982242 | Tigger3b | 3.89E-18 | OGFRL1 | 9016262 |
| chr6 | 99845302 | 99845615 | Tigger3a | 1.60E-24 | FOXO3 | 9035421 |
| chr5 | 100208148 | 100208501 | THE1C | 4.24E-190 | TSLP | 10197257 |
| chr4 | 57927355 | 57927939 | MER44B | 1.88E-103 | STAP1 | 10496505 |
| chr5 | 90502602 | 90503297 | Tigger3b | 1.11E-11 | PAPD4 | 11525127 |
| chr4 | 54939437 | 54939923 | MLT1F | 3.52E-11 | STAP1 | 13484521 |

Table S7. PARs near DEGs in L12 samples (PARs belonging to the 14 related TE families included only)

| TE.chr | TE.start | TE.end | TE.family | Peak.pval | Gene | Distance.to.TSS |
|---|---|---|---|---|---|---|
| chr4 | 74961891 | 74962441 | MER41B | 2.72E-17 | CXCL2 | 309 |
| chr1 | 167754291 | 167754845 | MLT1H | 1.82E-29 | MPZL1 | 1457 |
| chr3 | 101566285 | 101566626 | Tigger3a | 2.63E-08 | NFKBIZ | 1730 |
| chr2 | 113814070 | 113814437 | THE1B | 1.86E-19 | IL36RN | 1776 |
| chr8 | 90794231 | 90794648 | MER44D | 7.64E-69 | RIPK2 | 1915 |
| chr2 | 6998590 | 6998932 | THE1B | 1.75E-25 | CMPK2 | 2455 |
| chr7 | 89786435 | 89786988 | MLT1K | 2.02E-06 | STEAP1 | 2665 |
| chr6 | 160086744 | 160087114 | MLT1K | 4.85E-05 | SOD2 | 2973 |
| chr2 | 113811677 | 113812041 | THE1B | 9.45E-10 | IL36RN | 4172 |
| chr19 | 6536247 | 6536326 | MLT1K | 1.00E-09 | TNFSF9 | 5237 |
| chr10 | 6239239 | 6239493 | Tigger3c | 3.14E-11 | PFKFB3 | 5334 |
| chr5 | 150433555 | 150433851 | MLT1I | 2.22E-17 | TNIP1 | 6030 |
| chr20 | 49119385 | 49119762 | MLT1K | 7.30E-09 | PTPN1 | 7127 |
| chr8 | 22216286 | 22216553 | MER44B | 2.28E-15 | SLC39A14 | 8207 |
| chr4 | 159744755 | 159744946 | MLT1K | 6.64E-176 | FNIP2 | 8284 |
| chr12 | 113426846 | 113427200 | THE1B | 3.19E-07 | OAS2 | 8848 |
| chr11 | 32102748 | 32103127 | MLT1H | 9.44E-35 | RCN1 | 9321 |
| chr7 | 26421286 | 26421649 | Tigger3c | 1.04E-39 | SNX10 | 11278 |
| chr1 | 79114002 | 79114290 | THE1C | 1.06E-12 | IFI44L | 11625 |
| chr10 | 90595119 | 90595493 | Tigger3c | 9.83E-05 | ANKRD22 | 12561 |
| chr9 | 113021117 | 113021387 | MLT1K | 8.33E-34 | TXN | 14818 |
| chr10 | 90597409 | 90597768 | THE1B | 2.15E-10 | ANKRD22 | 14851 |
| chr10 | 26970664 | 26971209 | MER44B | 3.19E-08 | PDSS1 | 15377 |
| chr8 | 90809097 | 90809726 | MER41B | 6.20E-17 | RIPK2 | 16781 |
| chr20 | 43783363 | 43783738 | THE1C | 2.00E-06 | PI3 | 19777 |
| chr7 | 100513942 | 100514403 | MER41B | 3.31E-06 | ACHE | 20097 |
| chr2 | 224819059 | 224819594 | MLT1F | 1.43E-16 | SERPINE2 | 20233 |
| chr10 | 91115280 | 91115922 | MER41B | 1.60E-13 | IFIT3 | 23039 |
| chr6 | 37210656 | 37210851 | MLT1L | 1.10E-20 | TMEM217 | 24710 |
| chr1 | 66231379 | 66231744 | THE1C | 2.57E-05 | PDE4B | 26451 |
| chr5 | 52056851 | 52057214 | THE1B | 2.71E-09 | ITGA1 | 26514 |
| chr2 | 28587533 | 28587831 | MLT1K | 1.65E-11 | FOSL2 | 27482 |
| chr8 | 80707426 | 80707501 | MLT1L | 1.89E-05 | HEY1 | 28181 |
| chr6 | 154444653 | 154445191 | MER41B | 7.45E-13 | IPCEF1 | 30438 |
| chr6 | 2919129 | 2919605 | MER41B | 4.03E-10 | SERPINB9 | 31629 |
| chr9 | 117583234 | 117584446 | Tigger3b | 2.00E-06 | TNFSF15 | 31634 |
| chr11 | 33753396 | 33753761 | THE1B | 1.69E-22 | C11orf91 | 32288 |
| chr2 | 224804164 | 224804525 | THE1B | 2.80E-11 | SERPINE2 | 35302 |
| chr1 | 111817719 | 111818119 | MLT1H | 8.91E-24 | CHI3L2 | 35791 |
| chr7 | 129134078 | 129134785 | Tigger3b | 1.52E-11 | STRIP2 | 35995 |
| chr1 | 159072793 | 159073149 | THE1C | 8.91E-24 | AIM2 | 36687 |
| chr3 | 172185173 | 172185281 | MLT1L | 2.09E-09 | TNFSF10 | 38015 |
| chr11 | 114210932 | 114211441 | MLT1L | 2.41E-12 | NNMT | 42059 |
| chr11 | 14358638 | 14359318 | MER44D | 7.30E-33 | RRAS2 | 42632 |
| chr2 | 224795308 | 224795759 | MLT1H | 1.81E-16 | SERPINE2 | 44068 |

| | | | | | | |
|---|---|---|---|---|---|---|
| chr3 | 172279540 | 172279935 | Tigger3b | 4.75E-13 | TNFSF10 | 44395 |
| chr10 | 30676299 | 30676667 | MLT1K | 8.17E-07 | MAP3K8 | 46197 |
| chr17 | 39829331 | 39829811 | MER41B | 9.45E-10 | KRT17 | 50206 |
| chr11 | 43890723 | 43891068 | MLT1H | 4.27E-08 | C11orf96 | 55822 |
| chr3 | 119383828 | 119384156 | MLT1L | 2.33E-54 | PLA1A | 56084 |
| chr22 | 37828492 | 37828997 | MLT1F | 2.00E-06 | CARD10 | 57401 |
| chr3 | 39283192 | 39283592 | MLT1K | 6.78E-06 | XIRP1 | 58485 |
| chr1 | 65553215 | 65553573 | THE1B | 3.19E-07 | AK4 | 59657 |
| chr5 | 172257796 | 172258300 | MLT1F | 9.44E-35 | DUSP1 | 62703 |
| chr20 | 48317136 | 48317463 | MER44B | 1.65E-08 | B4GALT5 | 67654 |
| chr11 | 75400899 | 75401184 | MLT1L | 1.87E-11 | DGAT2 | 69371 |
| chr2 | 216734784 | 216735282 | MLT1F | 9.92E-06 | MREG | 73929 |
| chr4 | 77029801 | 77030940 | Tigger3b | 6.22E-20 | CXCL11 | 73958 |
| chr1 | 89442051 | 89442703 | Tigger3b | 6.93E-07 | GBP1 | 75297 |
| chr13 | 27766592 | 27766941 | THE1B | 7.39E-15 | RASL11A | 77521 |
| chr12 | 55326947 | 55327537 | MER41B | 3.69E-08 | MUCL1 | 78626 |
| chr1 | 110539282 | 110539461 | Tigger3b | 1.78E-12 | CSF1 | 81003 |
| chr9 | 34771146 | 34771482 | Tigger3a | 2.72E-14 | CCL19 | 81292 |
| chr2 | 29330930 | 29331464 | MER44B | 4.36E-16 | ALK | 84174 |
| chr2 | 224754093 | 224754450 | THE1B | 2.11E-88 | SERPINE2 | 85377 |
| chr5 | 52315110 | 52315361 | MER44B | 4.76E-21 | ITGA1 | 88128 |
| chr12 | 66606858 | 66607146 | THE1B | 1.15E-11 | HELB | 89177 |
| chr3 | 126332409 | 126333025 | Tigger3b | 1.78E-12 | CHST13 | 89283 |
| chr6 | 160270994 | 160271369 | THE1B | 3.19E-07 | SOD2 | 89704 |
| chrX | 108957156 | 108958109 | Tigger3 | 6.78E-06 | KCNE1L | 90227 |
| chr14 | 103893862 | 103894372 | MER41B | 8.91E-24 | CKB | 91622 |
| chr20 | 48341310 | 48341749 | MLT1K | 1.08E-09 | B4GALT5 | 91828 |
| chr10 | 92765034 | 92765396 | THE1B | 5.68E-11 | ANKRD1 | 93181 |
| chr7 | 24802641 | 24803324 | MER44C | 4.85E-05 | MPP6 | 97339 |
| chr2 | 96908087 | 96908411 | MLT1K | 1.41E-10 | DUSP2 | 98619 |
| chr2 | 28515344 | 28515668 | MLT1L | 2.72E-09 | FOSL2 | 99645 |
| chr4 | 75130799 | 75131140 | MLT1H | 5.35E-09 | EREG | 99718 |
| chr4 | 113091403 | 113091944 | MER44B | 3.15E-17 | TIFA | 103752 |
| chr4 | 159917934 | 159918179 | MLT1L | 7.90E-17 | RAPGEF2 | 107149 |
| chr1 | 226896112 | 226896546 | MLT1L | 7.59E-18 | C1orf95 | 107822 |
| chr7 | 20513939 | 20514223 | MLT1I | 5.65E-16 | ITGB8 | 109671 |
| chr17 | 76238467 | 76238881 | MLT1F | 1.75E-07 | SOCS3 | 113981 |
| chr4 | 40317084 | 40317266 | Tigger3c | 1.78E-12 | RHOH | 115119 |
| chr1 | 51635848 | 51636244 | Tigger3a | 8.31E-22 | TTC39A | 116684 |
| chr10 | 97833582 | 97833754 | MLT1L | 4.92E-52 | BLNK | 117702 |
| chr6 | 18242613 | 18243341 | MER44C | 3.84E-36 | RNF144B | 125436 |
| chr2 | 128528712 | 128529065 | Tigger3a | 2.00E-06 | LIMS2 | 129012 |
| chr1 | 94220111 | 94220905 | Tigger3b | 1.22E-10 | GCLM | 129854 |
| chr6 | 37007299 | 37007440 | MLT1H | 3.72E-06 | PIM1 | 130537 |
| chr8 | 58774567 | 58774929 | THE1C | 8.37E-10 | FAM110B | 132137 |
| chr7 | 93651925 | 93652302 | THE1C | 1.64E-18 | TFPI2 | 132695 |
| chr17 | 13542297 | 13542927 | MER41B | 2.14E-06 | HS3ST3A1 | 142810 |

| | | | | | | |
|---|---|---|---|---|---|---|
| chr15 | 38962977 | 38963566 | MLT1L | 4.21E-10 | RASGRP1 | 144392 |
| chr11 | 105022710 | 105023179 | Tigger3b | 4.84E-06 | CASP5 | 148709 |
| chr5 | 68763581 | 68763933 | MLT1K | 6.47E-05 | CCDC125 | 154116 |
| chr12 | 105411670 | 105412004 | Tigger3a | 1.92E-07 | APPL2 | 155068 |
| chr8 | 105671636 | 105671948 | MLT1K | 3.08E-18 | LRP12 | 155164 |
| chr20 | 61076914 | 61077279 | THE1C | 7.13E-38 | LAMA5 | 155300 |
| chr7 | 22925525 | 22925994 | MLT1L | 2.80E-11 | IL6 | 157406 |
| chr14 | 103827940 | 103828503 | MER44D | 3.28E-06 | CKB | 157491 |
| chr1 | 63087633 | 63087914 | MLT1H | 1.06E-29 | ATG4C | 161890 |
| chr1 | 149746990 | 149747619 | MER41B | 6.24E-11 | OTUD7B | 162084 |
| chr4 | 89368434 | 89368722 | THE1B | 6.94E-09 | PPM1K | 168715 |
| chr1 | 51948608 | 51949030 | MER44B | 2.00E-06 | TTC39A | 171679 |
| chr17 | 70988944 | 70989302 | THE1B | 3.10E-10 | SSTR2 | 171847 |
| chr7 | 112546156 | 112546454 | Tigger3a | 5.17E-33 | GPR85 | 171930 |
| chr10 | 13952034 | 13952557 | MLT1H | 2.50E-05 | FRMD4A | 172387 |
| chr4 | 5040086 | 5040704 | Tigger3b | 9.64E-28 | MSX1 | 175931 |
| chr1 | 65809021 | 65809734 | MER44C | 8.03E-32 | AK4 | 177406 |
| chr22 | 30838382 | 30838659 | MER41B | 0.000296 | OSM | 178318 |
| chr1 | 33515701 | 33516099 | MER44B | 5.99E-14 | FNDC5 | 179226 |
| chr10 | 45655447 | 45656107 | MER44D | 5.42E-100 | C10orf10 | 182220 |
| chr11 | 102454069 | 102454414 | Tigger3a | 3.72E-06 | MMP10 | 186818 |
| chr17 | 9253347 | 9253724 | MER44D | 1.92E-25 | NTN1 | 187095 |
| chr8 | 41844574 | 41844872 | Tigger3a | 5.56E-07 | PLAT | 187362 |
| chr14 | 52372374 | 52372739 | THE1C | 6.78E-15 | FRMD6 | 188401 |
| chr1 | 182374947 | 182375396 | Tigger3c | 1.55E-05 | RGS16 | 192360 |
| chr12 | 94167004 | 94167340 | THE1B | 7.00E-59 | SOCS2 | 198170 |
| chr8 | 21555167 | 21555530 | THE1B | 2.15E-27 | DOK2 | 210852 |
| chr3 | 142623575 | 142623917 | MLT1K | 1.12E-18 | CHST2 | 214254 |
| chr2 | 43666528 | 43666941 | MER44B | 9.45E-10 | ZFP36L2 | 216987 |
| chr8 | 99739225 | 99739459 | MLT1K | 8.00E-09 | OSR2 | 217170 |
| chr3 | 134313911 | 134314284 | THE1C | 1.22E-07 | AMOTL2 | 223734 |
| chr7 | 22536596 | 22537152 | MLT1K | 1.74E-27 | IL6 | 228349 |
| chr4 | 159461224 | 159461798 | MER41B | 3.93E-97 | FNIP2 | 228490 |
| chr2 | 85125510 | 85126036 | MLT1F | 0.000268 | TCF7L1 | 234495 |
| chr9 | 102827033 | 102827717 | Tigger3 | 1.52E-24 | NR4A3 | 238024 |
| chr10 | 91397049 | 91397822 | Tigger3b | 1.55E-16 | IFIT1 | 238487 |
| chr20 | 6990971 | 6991535 | MER41B | 7.98E-15 | BMP2 | 242660 |
| chr10 | 104962238 | 104962368 | MLT1L | 7.59E-09 | CALHM2 | 244173 |
| chr4 | 90414080 | 90414187 | Tigger3b | 5.35E-31 | GPRIN3 | 247994 |
| chr1 | 212484820 | 212485692 | Tigger3b | 0.000235 | ATF3 | 252982 |
| chr1 | 221314446 | 221314811 | THE1B | 0.000384 | HLX | 259862 |
| chr3 | 171959191 | 171959547 | THE1B | 1.22E-10 | TNFSF10 | 263749 |
| chr6 | 111612175 | 111612540 | THE1B | 1.08E-11 | TRAF3IP2 | 265115 |
| chr9 | 33175508 | 33175883 | Tigger3a | 1.52E-11 | AQP3 | 265267 |
| chr7 | 93786761 | 93787403 | MER41B | 2.07E-36 | TFPI2 | 267531 |
| chr10 | 92950777 | 92951311 | MER44B | 9.45E-10 | ANKRD1 | 278924 |
| chr2 | 224333992 | 224334472 | MLT1H | 1.23E-33 | AP1S3 | 281929 |

| | | | | | | |
|---|---|---|---|---|---|---|
| chr3 | 148798669 | 148798884 | Tigger3c | 1.34E-07 | TM4SF1 | 287923 |
| chr8 | 121270299 | 121270844 | MLT1K | 1.40E-08 | DEPTOR | 288271 |
| chr5 | 157297791 | 157298272 | MER44D | 5.97E-37 | ADAM19 | 300337 |
| chr7 | 90092429 | 90092799 | THE1C | 1.67E-05 | STEAP1 | 301849 |
| chr9 | 107906250 | 107906600 | Tigger3a | 2.82E-10 | FSD1L | 303475 |
| chr15 | 90325520 | 90325966 | MER41B | 6.21E-14 | RHCG | 304490 |
| chr11 | 35633408 | 35633829 | MLT1L | 6.94E-09 | SLC1A2 | 305668 |
| chr7 | 128761599 | 128761953 | THE1B | 1.63E-09 | STRIP2 | 312319 |
| chr10 | 104892747 | 104893030 | THE1B | 1.01E-27 | CALHM2 | 313511 |
| chr4 | 112880620 | 112881098 | MLT1F | 1.55E-93 | TIFA | 314598 |
| chr19 | 17618766 | 17619037 | MER44B | 1.09E-21 | JAK3 | 316550 |
| chr6 | 111559690 | 111560016 | MLT1K | 1.71E-12 | TRAF3IP2 | 317639 |
| chr20 | 30929042 | 30929387 | Tigger3a | 1.67E-05 | CCM2L | 322021 |
| chr5 | 131084417 | 131085077 | Tigger3b | 1.11E-10 | CSF2 | 324404 |
| chr12 | 62662886 | 62663204 | Tigger3a | 5.35E-31 | C12orf61 | 332325 |
| chr12 | 121122160 | 121122431 | Tigger3a | 4.23E-08 | OASL | 335662 |
| chr11 | 129013184 | 129013887 | MER44C | 1.05E-25 | FLI1 | 339413 |
| chr2 | 17093009 | 17093368 | THE1C | 5.18E-05 | FAM49A | 347717 |
| chr8 | 81930682 | 81931045 | THE1B | 7.33E-24 | ZNF704 | 347854 |
| chr1 | 70965013 | 70965905 | Tigger3b | 1.14E-12 | PTGER3 | 352129 |
| chr9 | 136695425 | 136695763 | THE1C | 2.80E-11 | SLC2A6 | 354897 |
| chr2 | 223662426 | 223662798 | Tigger3a | 5.57E-43 | SGPP2 | 373190 |
| chr6 | 17993785 | 17994045 | MLT1I | 1.50E-07 | RNF144B | 374732 |
| chr1 | 155540092 | 155540553 | MER44D | 2.28E-15 | MUC1 | 378099 |
| chr1 | 52156453 | 52156945 | Tigger3b | 1.15E-15 | TTC39A | 379524 |
| chr4 | 185296841 | 185297205 | THE1C | 2.14E-06 | ACSL1 | 379542 |
| chr2 | 46757770 | 46758135 | THE1B | 6.94E-09 | TTC7A | 385159 |
| chr1 | 203527617 | 203527743 | MLT1K | 4.81E-08 | MYBPH | 390678 |
| chr3 | 113395117 | 113395486 | THE1C | 1.52E-16 | BOC | 391710 |
| chr9 | 33836197 | 33836410 | Tigger3c | 6.90E-05 | AQP3 | 393818 |
| chr1 | 42995503 | 42996082 | MER44D | 1.10E-26 | SLC2A1 | 394968 |
| chr21 | 48047113 | 48047490 | THE1C | 6.25E-05 | LSS | 399880 |
| chr17 | 40418984 | 40419694 | Tigger3b | 8.67E-07 | PLEKHH3 | 400236 |
| chr22 | 37045653 | 37046017 | THE1B | 2.29E-16 | APOL2 | 412643 |
| chr2 | 127758611 | 127758872 | Tigger3b | 4.81E-08 | PROC | 417129 |
| chr15 | 74827692 | 74828192 | MLT1H | 5.69E-09 | RPP25 | 418563 |
| chr1 | 182140287 | 182140513 | MLT1L | 8.23E-30 | RGS16 | 427243 |
| chr3 | 184532149 | 184533290 | Tigger3b | 3.85E-15 | CHRD | 429573 |
| chr3 | 184532149 | 184533290 | Tigger3b | 5.82E-35 | CHRD | 429573 |
| chr11 | 127896216 | 127896931 | MER44C | 4.39E-130 | ETS1 | 431723 |
| chr10 | 45906248 | 45906876 | MER41B | 1.08E-64 | C10orf10 | 433021 |
| chr7 | 113161161 | 113161869 | MER44D | 5.58E-08 | GPR85 | 436375 |
| chr6 | 11851570 | 11852087 | MLT1H | 2.02E-13 | EDN1 | 438507 |
| chr2 | 74206785 | 74207160 | MLT1I | 8.48E-13 | RTKN | 445801 |
| chr1 | 226288636 | 226288846 | Tigger3b | 2.19E-11 | C1orf95 | 447653 |
| chr3 | 149547557 | 149547921 | THE1B | 2.71E-09 | TM4SF1 | 457998 |
| chr6 | 17887235 | 17887599 | MLT1I | 2.48E-30 | RNF144B | 481178 |

| | | | | | | |
|---|---|---|---|---|---|---|
| chr18 | 11925092 | 11925346 | Tigger3 | 1.12E-18 | SLMO1 | 482547 |
| chr1 | 247531208 | 247531843 | MER41B | 1.12E-22 | TRIM58 | 488656 |
| chr5 | 150948731 | 150949095 | THE1B | 1.59E-07 | TNIP1 | 490767 |
| chr13 | 52288880 | 52289228 | Tigger3b | 7.33E-13 | FAM124A | 492367 |
| chr5 | 171702051 | 171702389 | THE1C | 6.53E-22 | DUSP1 | 492702 |
| chr9 | 107713202 | 107713390 | Tigger3b | 4.45E-48 | FSD1L | 496685 |
| chr12 | 46943716 | 46943902 | MER44B | 1.08E-09 | AMIGO2 | 525586 |
| chr5 | 130873915 | 130874212 | Tigger3a | 1.61E-50 | CSF2 | 535269 |
| chr5 | 17356208 | 17356254 | THE1B | 3.96E-22 | MYO10 | 541582 |
| chr20 | 11163266 | 11163716 | MLT1H | 1.55E-13 | JAG1 | 544859 |
| chr10 | 31274814 | 31275143 | Tigger3a | 8.21E-31 | MAP3K8 | 546708 |
| chr1 | 149362059 | 149362689 | MER41B | 4.51E-42 | OTUD7B | 547014 |
| chr6 | 71450450 | 71450807 | THE1B | 1.21E-12 | OGFRL1 | 547697 |
| chr2 | 192561577 | 192562205 | Tigger3 | 1.25E-05 | STAT4 | 548723 |
| chr2 | 192562518 | 192562795 | Tigger3 | 1.42E-05 | STAT4 | 549664 |
| chr11 | 35880831 | 35881068 | MLT1K | 4.10E-101 | SLC1A2 | 553091 |
| chr20 | 10045746 | 10046138 | MLT1K | 0.000331 | JAG1 | 572192 |
| chr8 | 90195310 | 90195825 | MLT1F | 9.45E-10 | RIPK2 | 574148 |
| chr18 | 21313224 | 21313581 | THE1B | 2.96E-06 | CABLES1 | 577435 |
| chr15 | 51926741 | 51927266 | MER41B | 0.000119 | TNFAIP8L3 | 577946 |
| chr6 | 71419640 | 71419913 | Tigger3b | 8.15E-28 | OGFRL1 | 578591 |
| chr11 | 44551879 | 44552510 | MER41B | 2.80E-13 | C11orf96 | 587824 |
| chr7 | 26998572 | 26998923 | MER44D | 2.29E-16 | SNX10 | 588564 |
| chr12 | 46872659 | 46872860 | THE1C | 0.000331 | AMIGO2 | 596628 |
| chr3 | 57691946 | 57692435 | MLT1K | 2.55E-36 | SPATA12 | 597477 |
| chr9 | 114903001 | 114903599 | Tigger3c | 2.00E-06 | SNX30 | 609517 |
| chr6 | 13500542 | 13500857 | MLT1F | 1.67E-22 | CD83 | 617013 |
| chr2 | 197827715 | 197828050 | MLT1I | 5.85E-05 | HECW2 | 619288 |
| chr2 | 240753682 | 240754017 | THE1B | 1.55E-05 | GPC1 | 621069 |
| chr1 | 32701665 | 32702098 | MER44D | 3.84E-36 | FNDC5 | 625769 |
| chr4 | 40829838 | 40830200 | THE1B | 1.78E-12 | RHOH | 627873 |
| chr1 | 184129504 | 184130051 | MLT1H | 5.98E-24 | FAM129A | 629805 |
| chr2 | 37371121 | 37371719 | Tigger3b | 6.78E-13 | CRIM1 | 630288 |
| chr7 | 127036229 | 127036712 | MLT1H | 4.55E-20 | LRRC4 | 630410 |
| chr5 | 149522186 | 149522863 | MER44D | 6.78E-06 | SMIM3 | 634643 |
| chr11 | 9682649 | 9683160 | MLT1F | 1.95E-05 | AMPD3 | 646698 |
| chr6 | 137539807 | 137540099 | Tigger3a | 6.00E-35 | TNFAIP3 | 648250 |
| chr5 | 40027554 | 40027918 | THE1B | 6.25E-05 | PTGER4 | 651680 |
| chr12 | 56070634 | 56071291 | Tigger3b | 5.48E-08 | IL23A | 661370 |
| chr17 | 36162041 | 36162289 | MLT1L | 4.58E-09 | C17orf96 | 665670 |
| chr2 | 152899228 | 152899857 | MER41B | 2.05E-24 | TNFAIP6 | 676572 |
| chr2 | 47951581 | 47952019 | MLT1K | 1.50E-05 | TTC7A | 678127 |
| chr3 | 15651558 | 15652008 | MLT1H | 7.86E-05 | FGD5 | 679288 |
| chr7 | 30227951 | 30228329 | THE1C | 1.30E-23 | CHN2 | 683540 |
| chr6 | 12976762 | 12977203 | MER44B | 4.42E-12 | EDN1 | 686166 |
| chr1 | 70622345 | 70622633 | MER44B | 5.13E-26 | PTGER3 | 695401 |
| chr1 | 154403854 | 154404497 | MER44D | 2.00E-63 | EFNA1 | 695437 |

| | | | | | | |
|---|---|---|---|---|---|---|
| chr1 | 192848509 | 192848879 | MLT1I | 9.66E-09 | RGS18 | 720703 |
| chr3 | 157878892 | 157879247 | MLT1I | 2.49E-20 | PTX3 | 724314 |
| chr6 | 83802284 | 83802739 | MER44B | 2.21E-52 | TPBG | 728323 |
| chr5 | 159471241 | 159471874 | MER41B | 1.21E-12 | IL12B | 729450 |
| chr7 | 95956062 | 95956445 | THE1C | 2.16E-30 | PDK4 | 731979 |
| chr17 | 33421929 | 33422202 | MLT1L | 2.29E-16 | CCL1 | 734582 |
| chr3 | 128949653 | 128950508 | Tigger3b | 8.04E-13 | GATA2 | 743802 |
| chr6 | 34354417 | 34354610 | Tigger3b | 5.50E-20 | ITPR3 | 765256 |
| chr1 | 22374913 | 22375429 | MLT1K | 5.85E-16 | ECE1 | 775585 |
| chr22 | 41064963 | 41065111 | MLT1L | 6.94E-09 | ENTHD1 | 775744 |
| chr6 | 17591533 | 17592003 | MLT1K | 1.33E-20 | RNF144B | 776774 |
| chr2 | 73867752 | 73867830 | MLT1H | 7.17E-10 | RTKN | 785131 |
| chr8 | 38970743 | 38970806 | MLT1H | 1.00E-05 | IDO1 | 788986 |
| chr10 | 69923648 | 69923853 | MLT1L | 2.49E-32 | DDX21 | 792029 |
| chr7 | 106756170 | 106756533 | MLT1I | 4.76E-21 | LAMB1 | 807709 |
| chr2 | 114708345 | 114708719 | THE1B | 6.51E-05 | IL1RN | 823192 |
| chr6 | 83909212 | 83909566 | THE1B | 7.86E-05 | TPBG | 835251 |
| chr3 | 152041267 | 152041654 | Tigger3a | 0.000331 | RAP2B | 838373 |
| chr3 | 57941755 | 57942079 | Tigger3a | 5.37E-28 | SPATA12 | 847286 |
| chr1 | 68092318 | 68092608 | MLT1I | 2.39E-19 | TCTEX1D1 | 850480 |
| chr5 | 159596353 | 159596707 | MLT1K | 5.28E-12 | IL12B | 854562 |
| chr12 | 110201106 | 110201640 | MLT1F | 8.69E-07 | HVCN1 | 864004 |
| chr19 | 16085681 | 16086056 | THE1C | 3.72E-06 | SYDE1 | 864289 |
| chr9 | 116669029 | 116669589 | MLT1K | 3.69E-08 | TNFSF15 | 877324 |
| chr2 | 30407975 | 30408320 | Tigger3a | 9.74E-09 | ALK | 898699 |
| chr7 | 101705188 | 101705485 | MLT1I | 1.26E-25 | VGF | 899391 |
| chr9 | 36560693 | 36561098 | MER41B | 1.92E-07 | SIT1 | 911269 |
| chr5 | 175863711 | 175864398 | MER44C | 3.96E-22 | RGS14 | 920438 |
| chr2 | 153160280 | 153160910 | MER41B | 1.00E-11 | TNFAIP6 | 937624 |
| chr11 | 9373659 | 9374002 | MLT1K | 4.81E-08 | AMPD3 | 955856 |
| chr14 | 36829366 | 36829514 | MLT1H | 5.35E-06 | NFKBIA | 956412 |
| chr10 | 89604848 | 89605975 | Tigger3b | 0.000172 | ANKRD22 | 975912 |
| chr5 | 148190243 | 148192108 | Tigger3 | 1.29E-60 | SPINK1 | 983594 |
| chr5 | 59056800 | 59057887 | Tigger3b | 6.40E-11 | ELOVL7 | 989729 |
| chr15 | 50343215 | 50343594 | THE1C | 7.14E-05 | TNFAIP8L3 | 1005199 |
| chr1 | 183577563 | 183577810 | MER44B | 3.23E-12 | RGS16 | 1009805 |
| chr15 | 74212762 | 74213204 | MER41B | 1.57E-08 | RPP25 | 1033551 |
| chr3 | 171183740 | 171184023 | MLT1F | 4.05E-22 | TNFSF10 | 1039273 |
| chr7 | 30589650 | 30589998 | THE1B | 1.16E-07 | CHN2 | 1045239 |
| chr6 | 110831016 | 110831645 | MER41B | 3.69E-08 | TRAF3IP2 | 1046010 |
| chr2 | 37788862 | 37789173 | MLT1I | 1.70E-10 | CRIM1 | 1048029 |
| chr1 | 193180251 | 193180975 | MER44C | 4.56E-34 | RGS18 | 1052445 |
| chr20 | 9536270 | 9536793 | MLT1H | 3.42E-20 | JAG1 | 1081537 |
| chr10 | 74055674 | 74056082 | MLT1K | 7.64E-11 | UNC5B | 1082990 |
| chr22 | 29548763 | 29549395 | MER41B | 1.85E-18 | LIF | 1087039 |
| chr1 | 222171932 | 222172302 | THE1C | 3.72E-06 | HLX | 1117348 |
| chr17 | 53847123 | 53847282 | MLT1K | 2.91E-16 | TRIM25 | 1117986 |

| | | | | | | |
|---|---|---|---|---|---|---|
| chr2 | 37862722 | 37863085 | THE1C | 2.90E-08 | CRIM1 | 1121889 |
| chr14 | 93312953 | 93313252 | MLT1I | 7.94E-08 | CATSPERB | 1123472 |
| chr16 | 27386408 | 27386919 | MLT1K | 2.80E-13 | IL27 | 1123762 |
| chr9 | 73172791 | 73173067 | MLT1K | 2.47E-07 | TMEM2 | 1125213 |
| chr1 | 93219965 | 93220441 | MLT1F | 8.79E-09 | GCLM | 1130318 |
| chr12 | 95100114 | 95100469 | THE1B | 7.04E-15 | SOCS2 | 1131280 |
| chr5 | 148343033 | 148343463 | MLT1L | 5.52E-16 | SPINK1 | 1136384 |
| chr1 | 93194243 | 93194948 | MER44C | 3.77E-24 | GCLM | 1155811 |
| chr22 | 29479836 | 29480396 | MER41B | 6.21E-12 | LIF | 1156038 |
| chr4 | 79685954 | 79686210 | MER44B | 4.23E-06 | CXCL13 | 1158968 |
| chr11 | 4070425 | 4070554 | MLT1K | 2.75E-08 | SLC22A18/ | 1161098 |
| chr7 | 77235910 | 77236362 | MER44B | 5.66E-18 | ZP3 | 1166695 |
| chr15 | 91191646 | 91192244 | MER41B | 2.26E-25 | RHCG | 1170616 |
| chr14 | 50770677 | 50771020 | Tigger3a | 1.46E-37 | FRMD6 | 1184796 |
| chr12 | 95155392 | 95156551 | Tigger3b | 4.50E-23 | SOCS2 | 1186558 |
| chr2 | 37940551 | 37940807 | MER44B | 8.97E-47 | CRIM1 | 1199718 |
| chr4 | 141172061 | 141172531 | MLT1F | 8.67E-07 | CCRN4L | 1207835 |
| chr3 | 129414219 | 129414582 | THE1C | 1.23E-30 | GATA2 | 1208368 |
| chr17 | 49011003 | 49011278 | Tigger3a | 2.93E-17 | FAM117A | 1211134 |
| chr18 | 56349774 | 56350008 | MLT1K | 4.34E-22 | PMAIP1 | 1217170 |
| chr9 | 80291393 | 80291778 | MLT1H | 1.02E-06 | GCNT1 | 1217247 |
| chr3 | 150317226 | 150317869 | MER41B | 1.44E-07 | TM4SF1 | 1227667 |
| chr5 | 67342275 | 67342514 | MER44C | 4.13E-22 | CCDC125 | 1233486 |
| chr11 | 129911866 | 129912146 | Tigger3 | 1.22E-30 | FLI1 | 1238095 |
| chr17 | 65279774 | 65280418 | MER41B | 3.31E-06 | FAM20A | 1250834 |
| chr2 | 62374460 | 62375275 | Tigger3b | 4.76E-42 | REL | 1265669 |
| chr21 | 34179124 | 34179479 | THE1B | 2.72E-17 | SLC5A3 | 1266389 |
| chr13 | 35736768 | 35737028 | MLT1L | 2.82E-07 | CCNA1 | 1268937 |
| chr8 | 122252778 | 122253055 | MLT1H | 2.11E-06 | DEPTOR | 1270750 |
| chr10 | 89305726 | 89306256 | MER41B | 3.00E-08 | ANKRD22 | 1275631 |
| chr6 | 139476143 | 139476699 | Tigger3c | 4.65E-10 | TNFAIP3 | 1279168 |
| chr10 | 69428094 | 69428584 | MER44B | 4.42E-35 | DDX21 | 1287298 |
| chr2 | 33751242 | 33751395 | MLT1K | 1.89E-88 | NLRC4 | 1301717 |
| chr9 | 111239909 | 111240244 | Tigger3a | 2.82E-17 | AKAP2 | 1302523 |
| chr15 | 73942987 | 73943366 | MLT1L | 1.94E-07 | RPP25 | 1303389 |
| chr3 | 192548787 | 192549270 | MLT1F | 1.14E-14 | HES1 | 1304662 |
| chr1 | 93032668 | 93033299 | Tigger3b | 1.18E-05 | GCLM | 1317460 |
| chr11 | 15658162 | 15658580 | MLT1H | 2.71E-09 | RRAS2 | 1342156 |
| chr10 | 12342564 | 12342858 | THE1B | 3.30E-17 | FRMD4A | 1342846 |
| chr2 | 31089920 | 31090492 | MLT1L | 5.50E-22 | NLRC4 | 1359028 |
| chr7 | 114950347 | 114950701 | THE1B | 2.18E-14 | MET | 1361741 |
| chr6 | 134430822 | 134431087 | Tigger3c | 3.77E-24 | VNN3 | 1376049 |
| chr1 | 92968602 | 92968880 | MLT1I | 1.95E-45 | GCLM | 1381879 |
| chr8 | 66893145 | 66893521 | THE1B | 9.00E-09 | CYP7B1 | 1384453 |
| chr6 | 144495225 | 144495491 | MLT1H | 9.90E-08 | HIVEP2 | 1394545 |
| chr2 | 153624477 | 153624831 | THE1C | 1.21E-12 | TNFAIP6 | 1401821 |
| chr4 | 184269988 | 184270609 | MER41B | 4.58E-25 | ACSL1 | 1406138 |

| | | | | | | |
|---|---|---|---|---|---|---|
| chr3 | 185525369 | 185525783 | MLT1F | 6.15E-55 | CHRD | 1422793 |
| chr6 | 141647318 | 141647999 | MER44D | 1.32E-70 | HIVEP2 | 1424603 |
| chr3 | 12723442 | 12723986 | MLT1F | 2.73E-08 | HRH1 | 1429057 |
| chr3 | 12724169 | 12724414 | MLT1K | 7.89E-24 | HRH1 | 1429784 |
| chr3 | 141405445 | 141406028 | MLT1K | 9.89E-07 | CHST2 | 1432143 |
| chr8 | 64067540 | 64068162 | MER41B | 8.28E-09 | CYP7B1 | 1432156 |
| chr11 | 17277995 | 17278364 | MER44B | 2.00E-57 | TMEM86A | 1436303 |
| chr2 | 145114029 | 145114323 | MLT1I | 6.80E-17 | KYNU | 1437781 |
| chr14 | 50498142 | 50498511 | MLT1I | 1.40E-36 | FRMD6 | 1457305 |
| chr1 | 170893894 | 170894252 | THE1B | 4.45E-08 | SLC19A2 | 1460727 |
| chr12 | 65200822 | 65201168 | THE1C | 3.19E-07 | HELB | 1495155 |
| chr20 | 12133481 | 12134006 | MER44B | 1.78E-05 | JAG1 | 1515074 |
| chr10 | 25211296 | 25211597 | THE1B | 1.46E-17 | APBB1IP | 1515533 |
| chr17 | 58159968 | 58160458 | MER44B | 2.46E-17 | TEX14 | 1516855 |
| chr16 | 17405540 | 17405902 | THE1B | 3.19E-07 | MYH11 | 1526668 |
| chr2 | 131872547 | 131872903 | THE1B | 3.35E-09 | LYPD1 | 1529521 |
| chr14 | 90508841 | 90509208 | THE1B | 9.90E-08 | CATSPERB | 1537830 |
| chr5 | 133364517 | 133365092 | MLT1F | 1.36E-21 | IRF1 | 1540154 |
| chr11 | 58967466 | 58967827 | THE1B | 2.80E-13 | YPEL4 | 1551533 |
| chrX | 39986339 | 39986784 | MLT1H | 9.02E-23 | GPR34 | 1561440 |
| chr14 | 107118966 | 107119320 | THE1B | 2.39E-23 | GPR132 | 1597238 |
| chr2 | 198817377 | 198817712 | THE1B | 8.41E-62 | HECW2 | 1608950 |
| chr7 | 102429096 | 102429570 | Tigger3b | 4.81E-08 | VGF | 1623299 |
| chr10 | 74607400 | 74607857 | MLT1F | 1.76E-11 | UNC5B | 1634716 |
| chr3 | 40869783 | 40870229 | MLT1H | 0.000425 | XIRP1 | 1645076 |
| chr9 | 77374893 | 77375472 | Tigger3c | 4.12E-21 | GCNT1 | 1659278 |
| chr6 | 158425180 | 158425805 | MER41B | 1.34E-11 | SOD2 | 1664282 |
| chr13 | 71954366 | 71954712 | THE1B | 1.14E-10 | KLF5 | 1674400 |
| chr18 | 55879512 | 55879859 | THE1B | 9.35E-06 | PMAIP1 | 1687319 |
| chr11 | 21828549 | 21828906 | THE1C | 7.59E-09 | NAV2 | 1692620 |
| chr12 | 45730220 | 45731325 | Tigger3b | 2.33E-54 | AMIGO2 | 1738163 |
| chr8 | 93503120 | 93503291 | MLT1K | 4.76E-21 | GEM | 1758188 |
| chr12 | 124961656 | 124962027 | THE1C | 1.17E-36 | HCAR3 | 1762353 |
| chr5 | 61834548 | 61834663 | Tigger3c | 1.67E-60 | ELOVL7 | 1774460 |
| chr4 | 183896738 | 183897307 | MER44B | 9.09E-08 | ACSL1 | 1779440 |
| chr7 | 141264041 | 141264412 | THE1C | 1.84E-31 | FAM131B | 1786079 |
| chr11 | 16886034 | 16886397 | THE1B | 2.80E-13 | TMEM86A | 1828270 |
| chr4 | 157855257 | 157856136 | Tigger3b | 5.17E-33 | FNIP2 | 1834152 |
| chr14 | 22681084 | 22681289 | Tigger3c | 3.00E-14 | LRRC16B | 1839915 |
| chr15 | 34869762 | 34870253 | Tigger3c | 1.18E-05 | GREM1 | 1846870 |
| chr6 | 110018497 | 110018941 | MLT1F | 2.82E-10 | TRAF3IP2 | 1858714 |
| chr5 | 18697855 | 18698184 | THE1B | 1.15E-34 | MYO10 | 1883229 |
| chr6 | 140083604 | 140083963 | MLT1I | 1.62E-11 | TNFAIP3 | 1886629 |
| chr6 | 16005726 | 16006284 | MLT1L | 6.47E-05 | CD83 | 1887854 |
| chr2 | 172260610 | 172261164 | MER44D | 1.63E-16 | KLHL41 | 1889838 |
| chr12 | 14988897 | 14989525 | MER44C | 9.13E-11 | GPRC5A | 1926894 |
| chr20 | 4810318 | 4810839 | MLT1H | 1.75E-07 | BMP2 | 1937470 |

| | | | | | | |
|---|---|---|---|---|---|---|
| chr12 | 109088700 | 109089127 | MLT1F | 0.000235 | HVCN1 | 1976517 |
| chr20 | 4767770 | 4768141 | MLT1I | 0.000254 | BMP2 | 1980168 |
| chr10 | 11667180 | 11667579 | MLT1I | 3.40E-21 | FRMD4A | 2018125 |
| chr8 | 129609922 | 129610295 | THE1C | 1.01E-11 | FAM84B | 2044862 |
| chr5 | 50007061 | 50007650 | MLT1L | 3.22E-06 | ITGA1 | 2076078 |
| chr1 | 31251332 | 31251687 | MLT1K | 2.80E-11 | FNDC5 | 2076180 |
| chr9 | 110437670 | 110438280 | MER41B | 1.70E-10 | AKAP2 | 2104487 |
| chr5 | 71784745 | 71784920 | MLT1I | 8.80E-08 | ENC1 | 2138312 |
| chr8 | 97805451 | 97805794 | THE1B | 8.62E-13 | OSR2 | 2150835 |
| chr8 | 134315537 | 134315771 | MLT1I | 2.61E-15 | KHDRBS3 | 2153927 |
| chr3 | 146260166 | 146260295 | Tigger3c | 8.46E-16 | AGTR1 | 2155274 |
| chr4 | 111027480 | 111027993 | MER41B | 2.51E-21 | TIFA | 2167703 |
| chr8 | 72922500 | 72922862 | THE1B | 8.62E-47 | SLCO5A1 | 2177542 |
| chr9 | 37840208 | 37841113 | Tigger3b | 3.19E-07 | SIT1 | 2190784 |
| chr3 | 146222268 | 146222899 | MER41B | 1.13E-07 | AGTR1 | 2192670 |
| chr1 | 224522131 | 224522488 | THE1B | 3.01E-12 | C1orf95 | 2214011 |
| chr6 | 43344297 | 43344837 | MLT1F | 9.90E-08 | TREML1 | 2226755 |
| chr2 | 68418273 | 68418838 | Tigger3c | 2.30E-27 | TGFA | 2255572 |
| chr15 | 55627216 | 55627462 | Tigger3a | 5.13E-26 | MYZAP | 2256675 |
| chr7 | 73548589 | 73548934 | MER41B | 8.73E-23 | SRRM3 | 2282280 |
| chr1 | 46704521 | 46704878 | THE1B | 1.14E-10 | ARTN | 2303042 |
| chr6 | 5190816 | 5190982 | Tigger3 | 1.34E-30 | SERPINB9 | 2303316 |
| chr18 | 10094341 | 10094861 | Tigger3b | 1.78E-12 | SLMO1 | 2313032 |
| chr14 | 89730517 | 89730883 | THE1B | 6.94E-09 | CATSPERB | 2316155 |
| chr8 | 134148179 | 134148488 | MLT1L | 1.54E-15 | KHDRBS3 | 2321210 |
| chr8 | 97630308 | 97630655 | Tigger3a | 1.32E-28 | OSR2 | 2325974 |
| chr15 | 49015537 | 49015796 | MLT1L | 0.000322 | TNFAIP8L3 | 2332997 |
| chr6 | 168256565 | 168257096 | MER44B | 2.01E-11 | DLL1 | 2334196 |
| chr21 | 45191482 | 45191852 | THE1C | 5.53E-34 | FTCD | 2364322 |
| chr1 | 224329192 | 224329593 | Tigger3c | 1.24E-07 | C1orf95 | 2406906 |
| chr7 | 140637528 | 140637884 | THE1B | 3.19E-07 | FAM131B | 2412607 |
| chr3 | 186600403 | 186600858 | MER44B | 4.03E-10 | CHRD | 2497827 |
| chr11 | 61872516 | 61873051 | MER41B | 2.06E-18 | NRXN2 | 2500593 |
| chr2 | 231195435 | 231196116 | MER44C | 3.04E-07 | CCL20 | 2515350 |
| chr10 | 33272226 | 33272510 | MLT1L | 1.34E-07 | MAP3K8 | 2544120 |
| chr3 | 169669747 | 169670068 | Tigger3a | 5.48E-39 | TNFSF10 | 2553228 |
| chr5 | 137796925 | 137797466 | MLT1K | 1.29E-06 | SLC25A48 | 2581207 |
| chr14 | 31806014 | 31806346 | Tigger3a | 9.86E-09 | EGLN3 | 2587089 |
| chr11 | 7698296 | 7698675 | THE1C | 1.44E-09 | AMPD3 | 2631183 |
| chr14 | 94840431 | 94840538 | MLT1I | 1.18E-05 | CATSPERB | 2650950 |
| chr11 | 107526314 | 107526600 | MER41B | 9.00E-09 | CASP5 | 2652313 |
| chr11 | 107552989 | 107553307 | MLT1H | 2.85E-13 | CASP5 | 2678988 |
| chr6 | 109153603 | 109153835 | MLT1L | 2.18E-14 | TRAF3IP2 | 2723820 |
| chr5 | 65847829 | 65848117 | MLT1H | 6.05E-09 | CCDC125 | 2727883 |
| chr2 | 9878517 | 9878968 | MLT1L | 7.17E-10 | RNF144A | 2741446 |
| chr10 | 17118583 | 17118924 | MLT1I | 6.78E-13 | FRMD4A | 2750409 |
| chr2 | 173147764 | 173147869 | Tigger3b | 6.99E-34 | KLHL41 | 2776992 |

| | | | | | | |
|---|---|---|---|---|---|---|
| chr2 | 58324707 | 58325901 | Tigger3b | 4.04E-07 | REL | 2782753 |
| chr20 | 3961008 | 3961262 | MER44B | 9.61E-29 | BMP2 | 2787047 |
| chr1 | 30531933 | 30532373 | MLT1K | 9.97E-14 | FNDC5 | 2795494 |
| chr5 | 138020959 | 138021094 | MLT1I | 1.24E-07 | SLC25A48 | 2805241 |
| chr14 | 21712783 | 21712997 | Tigger3b | 2.80E-11 | LRRC16B | 2808207 |
| chr11 | 111264988 | 111265352 | THE1B | 4.63E-05 | NNMT | 2863155 |
| chr4 | 84066921 | 84067278 | THE1C | 6.47E-05 | FGF5 | 2877502 |
| chr10 | 109378085 | 109378419 | Tigger3 | 1.25E-63 | DUSP5 | 2879175 |
| chr18 | 32512896 | 32513233 | MLT1I | 2.38E-09 | RNF125 | 2890854 |
| chr1 | 24814651 | 24815016 | THE1B | 6.94E-09 | GPR3 | 2904130 |
| chr18 | 9494536 | 9495166 | MER41B | 2.64E-28 | SLMO1 | 2912727 |
| chr12 | 89888321 | 89888821 | MLT1H | 1.56E-147 | CLLU1OS | 2925047 |
| chr10 | 23801349 | 23801836 | MER44B | 5.94E-07 | APBB1IP | 2925294 |
| chr8 | 62569826 | 62570178 | Tigger3a | 2.86E-34 | CYP7B1 | 2930140 |
| chr5 | 43638030 | 43638402 | THE1C | 5.07E-05 | PTGER4 | 2956226 |
| chr22 | 43259606 | 43259980 | THE1C | 3.87E-12 | ENTHD1 | 2970387 |
| chr11 | 107859341 | 107859668 | Tigger3 | 9.69E-124 | CASP5 | 2985340 |
| chr2 | 24728504 | 24729034 | MLT1K | 5.69E-09 | FNDC4 | 2985714 |
| chr11 | 61382315 | 61382683 | THE1B | 9.45E-10 | NRXN2 | 2990961 |
| chr7 | 32545842 | 32546242 | MER44B | 2.65E-54 | CHN2 | 3001431 |
| chr2 | 136458975 | 136459437 | MER44D | 0.000235 | LYPD1 | 3038376 |
| chr2 | 167267636 | 167268318 | MER44C | 6.78E-16 | KLHL41 | 3097892 |
| chr20 | 33717389 | 33717820 | Tigger3b | 1.64E-06 | CCM2L | 3110368 |
| chr9 | 89100196 | 89100565 | MLT1I | 3.22E-10 | GADD45G | 3119361 |
| chr1 | 98120304 | 98120674 | THE1B | 1.81E-07 | F3 | 3121642 |
| chr3 | 122464257 | 122464953 | MER44C | 4.15E-19 | PLA1A | 3136513 |
| chr6 | 26351657 | 26352329 | MER44D | 1.04E-33 | UBD | 3170961 |
| chr3 | 33833094 | 33833611 | MLT1H | 6.81E-111 | TGFBR2 | 3185001 |
| chr10 | 115455271 | 115455905 | MER41B | 4.10E-08 | DUSP5 | 3192766 |
| chr3 | 18166052 | 18166432 | MLT1I | 1.74E-08 | FGD5 | 3193782 |
| chr11 | 61137004 | 61137430 | MLT1H | 7.26E-10 | NRXN2 | 3236214 |
| chr5 | 55464046 | 55464530 | MER44B | 1.49E-76 | ITGA1 | 3237064 |
| chr12 | 116882156 | 116882527 | THE1B | 1.66E-22 | IQCD | 3244306 |
| chr8 | 87522783 | 87522990 | MLT1F | 2.52E-10 | RIPK2 | 3246983 |
| chr2 | 201287414 | 201287953 | MER44B | 6.94E-09 | CD28 | 3283243 |
| chr8 | 117583699 | 117584106 | MLT1H | 1.14E-41 | DEPTOR | 3301849 |
| chr11 | 6226085 | 6226427 | Tigger3a | 7.99E-46 | SLC22A18A | 3316758 |
| chrX | 122221168 | 122221424 | Tigger3c | 4.81E-08 | SOWAHD | 3328592 |
| chr20 | 3401009 | 3401541 | MLT1F | 1.15E-21 | BMP2 | 3346768 |
| chr14 | 95603088 | 95603302 | MLT1L | 8.31E-12 | CATSPERB | 3413607 |
| chr2 | 200636551 | 200637094 | MER44B | 3.70E-05 | HECW2 | 3428124 |
| chr4 | 85735537 | 85736076 | MER44B | 9.75E-08 | PPM1K | 3442694 |
| chr16 | 80704083 | 80704457 | THE1C | 1.97E-15 | DNAAF1 | 3474406 |
| chr2 | 148738821 | 148739084 | MER44B | 1.59E-07 | TNFAIP6 | 3475020 |
| chr5 | 56572179 | 56572542 | THE1C | 2.06E-18 | ELOVL7 | 3475074 |
| chr1 | 244545246 | 244545380 | MLT1F | 1.11E-31 | TRIM58 | 3475119 |
| chr5 | 77844789 | 77845275 | MLT1H | 8.79E-08 | GCNT4 | 3521500 |

| | | | | | | |
|---|---|---|---|---|---|---|
| chr1 | 197808967 | 197809330 | THE1B | 1.29E-05 | LAD1 | 3533040 |
| chr2 | 136995651 | 136996006 | MLT1K | 6.15E-14 | LYPD1 | 3575052 |
| chr18 | 53940782 | 53941149 | THE1B | 1.78E-12 | PMAIP1 | 3626029 |
| chr15 | 85227428 | 85227788 | THE1B | 6.85E-25 | IL16 | 3629341 |
| chr3 | 18616854 | 18617304 | MLT1L | 3.07E-08 | FGD5 | 3644584 |
| chr8 | 131232913 | 131233305 | THE1C | 3.77E-06 | FAM84B | 3667853 |
| chr3 | 197522085 | 197522538 | MLT1K | 5.03E-10 | HES1 | 3668148 |
| chr2 | 108972474 | 108972550 | Tigger3c | 2.29E-16 | MERTK | 3683504 |
| chr9 | 88530701 | 88531067 | THE1B | 3.19E-07 | GADD45G | 3688859 |
| chr9 | 95908954 | 95909315 | THE1B | 3.87E-23 | GADD45G | 3688947 |
| chr9 | 95910955 | 95911480 | MLT1F | 9.44E-35 | GADD45G | 3690948 |
| chr1 | 173131116 | 173132035 | Tigger3b | 3.72E-06 | SLC19A2 | 3697949 |
| chr8 | 117174049 | 117174302 | Tigger3c | 2.66E-27 | DEPTOR | 3711653 |
| chr8 | 109258310 | 109258510 | MER44B | 0.000115 | LRP12 | 3741838 |
| chr3 | 190104232 | 190104514 | MER44B | 1.21E-09 | HES1 | 3749418 |
| chr8 | 74497991 | 74498329 | THE1C | 3.28E-06 | SLCO5A1 | 3753033 |
| chrX | 37731208 | 37731641 | MLT1H | 1.29E-05 | GPR34 | 3816583 |
| chr20 | 39950417 | 39950856 | Tigger3b | 3.58E-32 | PI3 | 3852659 |
| chr8 | 117031898 | 117032192 | MLT1I | 1.78E-12 | DEPTOR | 3853763 |
| chr16 | 70839714 | 70840038 | MER44B | 3.59E-06 | RRAD | 3883979 |
| chr5 | 84109437 | 84109800 | THE1B | 1.67E-05 | MEF2C | 3904173 |
| chr11 | 110208193 | 110208779 | MER41B | 4.81E-08 | NNMT | 3919728 |
| chr9 | 96160338 | 96160702 | THE1B | 2.22E-17 | GADD45G | 3940331 |
| chr11 | 110182941 | 110183290 | THE1C | 3.49E-45 | NNMT | 3945217 |
| chr1 | 59255903 | 59256211 | MLT1I | 4.10E-08 | ATG4C | 3993593 |
| chr2 | 12728355 | 12728905 | MER44B | 7.96E-50 | FAM49A | 4001820 |
| chr1 | 178535424 | 178535792 | MLT1H | 2.06E-42 | RGS16 | 4031964 |
| chr3 | 45103610 | 45103839 | MER41B | 1.17E-28 | KLHDC8B | 4105203 |
| chr1 | 243897642 | 243898738 | Tigger3b | 5.02E-11 | TRIM58 | 4121761 |
| chr5 | 142642019 | 142642268 | MER44C | 3.76E-41 | DPYSL3 | 4128104 |
| chr5 | 122066133 | 122066530 | MLT1K | 1.34E-30 | 03-Mar | 4136874 |
| chr11 | 109968459 | 109969611 | Tigger3b | 1.24E-05 | NNMT | 4158896 |
| chr10 | 116470008 | 116470366 | THE1B | 3.36E-26 | DUSP5 | 4207503 |
| chr20 | 2506234 | 2506760 | MER41B | 6.94E-09 | BMP2 | 4241549 |
| chr5 | 114679681 | 114680409 | MER44C | 1.21E-41 | TSLP | 4270669 |
| chr3 | 4467405 | 4467807 | THE1B | 8.28E-09 | OXTR | 4324285 |
| chr10 | 86253219 | 86253424 | MER41B | 3.12E-28 | ANKRD22 | 4328463 |
| chr3 | 167852729 | 167853040 | THE1B | 3.87E-23 | TNFSF10 | 4370256 |
| chr6 | 150075703 | 150076160 | MER44B | 1.01E-49 | IPCEF1 | 4399469 |
| chr14 | 59744593 | 59745300 | MER44D | 2.91E-16 | GCH1 | 4434879 |
| chr2 | 187441277 | 187441752 | MLT1H | 3.50E-28 | STAT4 | 4452548 |
| chr6 | 25047223 | 25047546 | THE1B | 4.81E-08 | UBD | 4475744 |
| chr6 | 45649527 | 45649902 | MLT1H | 1.55E-06 | TREML1 | 4531985 |
| chr5 | 163321419 | 163321959 | MER44B | 8.54E-18 | IL12B | 4579628 |
| chr7 | 37105464 | 37105817 | THE1B | 2.41E-05 | INHBA | 4618893 |
| chr1 | 243380222 | 243380503 | Tigger3a | 0.000235 | TRIM58 | 4639996 |
| chr5 | 163429736 | 163430317 | MLT1K | 2.77E-19 | IL12B | 4687945 |

| | | | | | | |
|---|---|---|---|---|---|---|
| chr8 | 110207785 | 110208132 | THE1B | 2.38E-09 | LRP12 | 4691313 |
| chr6 | 116588043 | 116588850 | Tigger3b | 4.52E-10 | TRAF3IP2 | 4694123 |
| chr6 | 116589209 | 116589641 | Tigger3b | 1.61E-07 | TRAF3IP2 | 4695289 |
| chr20 | 25353273 | 25353487 | Tigger3c | 1.62E-52 | REM1 | 4709607 |
| chr6 | 24743677 | 24744015 | MLT1L | 9.56E-10 | UBD | 4779275 |
| chr1 | 56561925 | 56562346 | MLT1F | 4.76E-19 | TTC39A | 4784996 |
| chr3 | 25862079 | 25862184 | MLT1K | 9.61E-29 | TGFBR2 | 4785808 |
| chr20 | 1944637 | 1945268 | MER41B | 1.94E-07 | BMP2 | 4803041 |
| chr1 | 117105129 | 117105486 | THE1B | 1.44E-06 | FAM212B | 4815798 |
| chr1 | 145090129 | 145090756 | MER41B | 1.22E-40 | OTUD7B | 4818947 |
| chr21 | 30610444 | 30610516 | MLT1I | 1.76E-46 | SLC5A3 | 4835352 |
| chr13 | 46923035 | 46923249 | THE1B | 1.52E-37 | FAM124A | 4873252 |
| chr1 | 145015160 | 145015649 | MER44B | 7.79E-46 | OTUD7B | 4894054 |
| chr14 | 86797250 | 86797583 | Tigger3b | 1.06E-10 | STON2 | 4932533 |
| chrX | 53743336 | 53743795 | MLT1L | 8.72E-20 | PIM2 | 4971663 |
| chr9 | 27448315 | 27448676 | THE1B | 4.81E-08 | DDX58 | 5006622 |
| chr7 | 134369674 | 134370259 | Tigger3b | 6.21E-19 | STRIP2 | 5271591 |
| chr20 | 35916391 | 35916907 | MLT1F | 7.04E-15 | CCM2L | 5309370 |
| chr5 | 82702876 | 82704056 | Tigger3b | 3.49E-45 | MEF2C | 5309917 |
| chr16 | 21314215 | 21314566 | THE1B | 1.99E-14 | MYH11 | 5435343 |
| chr6 | 106414400 | 106414782 | MLT1L | 4.15E-17 | TRAF3IP2 | 5462873 |
| chr5 | 79812664 | 79813203 | MER44B | 1.92E-07 | GCNT4 | 5489375 |
| chr6 | 148778812 | 148779118 | THE1C | 2.75E-11 | HIVEP2 | 5678132 |
| chr20 | 36308911 | 36309023 | MLT1L | 1.34E-30 | CCM2L | 5701890 |
| chr1 | 100702528 | 100703026 | MLT1H | 1.00E-09 | F3 | 5703866 |
| chr2 | 55360932 | 55361440 | MER41B | 5.99E-52 | REL | 5747214 |
| chr10 | 64965375 | 64965583 | Tigger3a | 1.10E-06 | DDX21 | 5750299 |
| chr10 | 120553127 | 120553548 | MLT1H | 2.31E-09 | FAM53B | 5754311 |
| chr21 | 21749695 | 21749998 | Tigger3c | 0.000288 | AF165138. | 5764869 |
| chr9 | 15298347 | 15298710 | THE1B | 1.50E-05 | IFNB1 | 5778392 |
| chr14 | 70195221 | 70195661 | MER41B | 9.89E-17 | BATF | 5793105 |
| chr14 | 70163515 | 70164158 | MER41B | 4.26E-18 | BATF | 5824608 |
| chr2 | 55275052 | 55275707 | Tigger3b | 0.000239 | REL | 5832947 |
| chr5 | 80163118 | 80163382 | MLT1L | 2.20E-40 | GCNT4 | 5839829 |
| chr13 | 45899836 | 45900328 | MLT1H | 4.12E-14 | FAM124A | 5896173 |
| chr3 | 95642292 | 95642912 | MER41B | 3.19E-07 | NFKBIZ | 5903921 |
| chr16 | 50691133 | 50691397 | MLT1K | 6.32E-14 | MT2A | 5950712 |
| chr13 | 45827633 | 45827928 | MER44B | 9.34E-11 | FAM124A | 5968573 |
| chr13 | 45783306 | 45783562 | Tigger3a | 3.04E-46 | FAM124A | 6012939 |
| chr18 | 67711689 | 67712215 | MER44B | 2.07E-157 | HMSD | 6084273 |
| chr8 | 111663983 | 111664221 | MLT1I | 3.81E-06 | LRP12 | 6147511 |
| chr20 | 509969 | 510585 | MER44C | 8.98E-30 | BMP2 | 6237724 |
| chr5 | 164998513 | 164998787 | THE1C | 6.64E-07 | IL12B | 6256722 |
| chr20 | 36904153 | 36904552 | Tigger3b | 1.72E-06 | CCM2L | 6297132 |
| chr4 | 153388256 | 153388620 | THE1C | 4.87E-06 | FNIP2 | 6301668 |
| chr5 | 165050936 | 165051391 | MER44B | 2.54E-11 | IL12B | 6309145 |
| chr1 | 176210219 | 176210932 | MER44D | 3.46E-28 | RGS16 | 6356824 |

| | | | | | | |
|---|---|---|---|---|---|---|
| chr20 | 36986203 | 36986835 | MER41B | 1.36E-21 | CCM2L | 6379182 |
| chr1 | 176020471 | 176021507 | Tigger3b | 9.53E-16 | RGS16 | 6546249 |
| chr20 | 23478177 | 23478514 | THE1B | 6.94E-09 | REM1 | 6584580 |
| chr5 | 119488207 | 119488915 | MER44C | 6.62E-57 | 03-Mar | 6714489 |
| chr5 | 81078332 | 81078952 | Tigger3b | 9.59E-07 | GCNT4 | 6755043 |
| chr6 | 48103178 | 48103695 | MLT1F | 1.67E-05 | TREML1 | 6985636 |
| chr2 | 159374558 | 159374942 | MLT1H | 8.67E-07 | TNFAIP6 | 7151902 |
| chr5 | 102841802 | 102841973 | MLT1L | 3.72E-06 | TSLP | 7563785 |
| chr4 | 177926806 | 177927315 | MER41B | 1.35E-39 | ACSL1 | 7749432 |
| chr14 | 68205748 | 68206146 | MLT1L | 1.67E-05 | BATF | 7782620 |
| chr4 | 130019709 | 130020237 | MER41B | 8.06E-06 | TNIP3 | 7944316 |
| chr11 | 94238084 | 94239279 | Tigger3b | 7.14E-05 | BIRC3 | 7948934 |
| chr2 | 183897090 | 183897403 | Tigger3a | 1.61E-05 | STAT4 | 7996897 |
| chr11 | 94182768 | 94183110 | Tigger3a | 1.06E-34 | BIRC3 | 8005103 |
| chr14 | 67838180 | 67838490 | Tigger3a | 7.88E-07 | BATF | 8150276 |
| chr2 | 162085058 | 162085553 | Tigger3c | 2.96E-06 | KLHL41 | 8280657 |
| chr12 | 39170521 | 39171577 | Tigger3b | 2.82E-07 | AMIGO2 | 8297911 |
| chr10 | 62414628 | 62415857 | Tigger3b | 2.06E-18 | DDX21 | 8300025 |
| chr4 | 48728289 | 48728396 | MLT1I | 5.68E-11 | RHOH | 8526324 |
| chr2 | 161161663 | 161162118 | Tigger3b | 2.71E-09 | TNFAIP6 | 8939007 |
| chr4 | 169225873 | 169226443 | Tigger3b | 3.37E-17 | RAPGEF2 | 8952264 |
| chr6 | 62981095 | 62982242 | Tigger3b | 6.94E-09 | OGFRL1 | 9016262 |
| chr4 | 99432421 | 99432749 | Tigger3a | 1.06E-34 | GPRIN3 | 9266335 |
| chr7 | 51145725 | 51146253 | MLT1L | 3.50E-39 | INHBA | 9416593 |
| chr4 | 103605889 | 103606137 | MER44B | 1.49E-48 | TIFA | 9589559 |
| chr4 | 103548936 | 103549210 | MLT1L | 1.21E-30 | TIFA | 9646486 |
| chr8 | 8691693 | 8692247 | Tigger3c | 2.54E-11 | PSD3 | 9692562 |
| chr16 | 46893821 | 46894151 | MLT1K | 7.76E-12 | MT2A | 9747958 |
| chr15 | 99822643 | 99822995 | Tigger3a | 1.18E-15 | RHCG | 9801613 |
| chr8 | 8483215 | 8483743 | MLT1H | 4.35E-09 | PSD3 | 9901066 |
| chr11 | 87750966 | 87751322 | THE1B | 4.30E-10 | KCTD14 | 10023327 |
| chr6 | 99845302 | 99845615 | Tigger3a | 1.96E-26 | SRSF12 | 10030102 |
| chr7 | 65715566 | 65716087 | MER44B | 6.01E-34 | SRRM3 | 10115127 |
| chr5 | 100208148 | 100208501 | THE1C | 1.40E-19 | TSLP | 10197257 |
| chr14 | 65678086 | 65678426 | MER44D | 9.63E-07 | BATF | 10310340 |
| chr14 | 65647200 | 65647579 | MLT1K | 2.03E-28 | GCH1 | 10337486 |
| chr11 | 88131388 | 88131901 | MER44B | 5.46E-10 | KCTD14 | 10403749 |
| chr18 | 72276498 | 72277071 | Tigger3c | 1.52E-54 | HMSD | 10649082 |
| chr4 | 174038592 | 174038959 | THE1B | 1.34E-07 | ACSL1 | 11637788 |
| chr12 | 24758281 | 24758686 | MLT1I | 4.83E-08 | GPRC5A | 11696278 |
| chr12 | 24877833 | 24878196 | THE1C | 2.08E-38 | GPRC5A | 11815830 |
| chr12 | 25062633 | 25063025 | MLT1F | 1.87E-16 | GPRC5A | 12000630 |
| chr12 | 25108302 | 25108675 | THE1B | 1.03E-25 | GPRC5A | 12046299 |
| chr15 | 102165386 | 102165762 | THE1C | 4.65E-10 | RHCG | 12144356 |
| chr8 | 6038171 | 6038426 | MLT1L | 1.71E-12 | PSD3 | 12346383 |
| chr18 | 44689374 | 44689932 | MER44B | 9.56E-15 | PMAIP1 | 12877246 |
| chrX | 16003964 | 16004475 | MLT1F | 1.71E-31 | P2RY8 | 14418634 |

| | | | | | | |
|---|---|---|---|---|---|---|
| chr4 | 54939437 | 54939923 | MLT1F | 4.87E-06 | RHOH | 14737472 |
| chr12 | 32576552 | 32576801 | MER41B | 2.08E-05 | AMIGO2 | 14892687 |
| chr4 | 55123212 | 55123663 | MER41B | 0.000331 | RHOH | 14921247 |
| chr6 | 56803463 | 56803830 | MLT1H | 1.04E-39 | OGFRL1 | 15194674 |
| chr13 | 96180369 | 96180731 | THE1B | 9.90E-08 | LINC00346 | 15340845 |
| chr4 | 24755122 | 24755453 | MLT1I | 7.37E-26 | RHOH | 15437218 |
| chr12 | 32012595 | 32012964 | THE1C | 1.43E-13 | AMIGO2 | 15456524 |
| chr12 | 28932919 | 28933620 | MER44D | 1.11E-67 | GPRC5A | 15870916 |
| chr4 | 56178114 | 56178526 | MLT1K | 6.12E-07 | RHOH | 15976149 |
| chr3 | 84773822 | 84774447 | MER44D | 2.63E-18 | NFKBIZ | 16772386 |
| chrX | 19728181 | 19728900 | MER44D | 1.64E-05 | P2RY8 | 18142851 |
