## Supplementary Tables 8 and 9 for "Transposable elements have contributed human regulatory regions that are activated upon bacterial infection"

Table S8. PARs in QTL-regions in S12 samples

(Ordered by decreasing peak signal, PARs belonging to the 14 related TE families included only)

| TE.chr | TE.start | TE.end | TE.family | Peak.pval | QTL.chr | QTL.start | QTL.end |
|---|---|---|---|---|---|---|---|
| chr2 | 224754093 | 224754450 | THE1B | 4.88E-310 | chr2 | 2.25E+08 | 224811722 |
| chr6 | 71450450 | 71450807 | THE1B | 2.30E-90 | chr6 | 71424364 | 71524118 |
| chr3 | 57941755 | 57942079 | Tigger3a | 1.07E-77 | chr3 | 57934626 | 57953782 |
| chr4 | 5040086 | 5040704 | Tigger3b | 1.73E-64 | chr4 | 5017508 | 5042928 |
| chr7 | 128761599 | 128761953 | THE1B | 2.93E-51 | chr7 | 1.29E+08 | 128769369 |
| chr14 | 50770677 | 50771020 | Tigger3a | 3.23E-34 | chr14 | 50699868 | 50792273 |
| chr9 | 15298347 | 15298710 | THE1B | 5.67E-31 | chr9 | 15287121 | 15304782 |
| chr10 | 90597409 | 90597768 | THE1B | 5.98E-29 | chr10 | 90592544 | 90608480 |
| chr3 | 128949653 | 128950508 | Tigger3b | 9.24E-26 | chr3 | 1.29E+08 | 129016398 |
| chr6 | 71521621 | 71521865 | MER44B | 8.56E-23 | chr6 | 71424364 | 71524118 |
| chr7 | 95956062 | 95956445 | THE1C | 4.68E-22 | chr7 | 95920516 | 95980767 |
| chr5 | 17356208 | 17356254 | THE1B | 8.85E-20 | chr5 | 17351738 | 17372997 |
| chr10 | 26970664 | 26971209 | MER44B | 2.96E-19 | chr10 | 26958858 | 26991179 |
| chr4 | 38823509 | 38823989 | Tigger3b | 1.11E-11 | chr4 | 38806827 | 38905731 |
| chr2 | 224804164 | 224804525 | THE1B | 1.85E-11 | chr2 | 2.25E+08 | 224811722 |
| chr1 | 28718549 | 28718911 | MER44C | 2.29E-11 | chr1 | 28654521 | 28749014 |
| chr4 | 39321395 | 39321663 | MER44B | 1.06E-09 | chr4 | 39303925 | 39395734 |
| chr11 | 30364344 | 30364828 | MLT1K | 1.72E-09 | chr11 | 30311493 | 30376608 |
| chr10 | 90595119 | 90595493 | Tigger3c | 8.74E-08 | chr10 | 90592544 | 90608480 |
| chr17 | 57877969 | 57878160 | Tigger3b | 6.40E-07 | chr17 | 57825572 | 57920532 |
| chr12 | 92815641 | 92815947 | Tigger3a | 6.78E-07 | chr12 | 92794173 | 92827601 |
| chr6 | 127619741 | 127620024 | Tigger3b | 1.32E-06 | chr6 | 1.28E+08 | 127712911 |
| chr20 | 48317136 | 48317463 | MER44B | 2.10E-06 | chr20 | 48242799 | 48322497 |
| chr1 | 212484820 | 212485692 | Tigger3b | 2.16E-06 | chr1 | 2.12E+08 | 212551650 |
| chr2 | 224795308 | 224795759 | MLT1H | 3.94E-06 | chr2 | 2.25E+08 | 224811722 |
| chr15 | 68476052 | 68478019 | Tigger3 | 2.16E-05 | chr15 | 68468695 | 68546275 |
| chr9 | 80629734 | 80630315 | Tigger3c | 8.11E-05 | chr9 | 80600487 | 80690409 |
| chr6 | 24743677 | 24744015 | MLT1L | 8.66E-05 | chr6 | 24729115 | 24754859 |
| chr5 | 159592669 | 159593059 | MLT1K | 0.00026987 | chr5 | 1.6E+08 | 159606199 |

| Ensembl.ID | Gene.name | reQTL.rsID | reQTL.pval | Is.DEG |
|---|---|---|---|---|
| ENSG00000135900 | MRPL44 | rs36053433 | 1.45E-06 | FALSE |
| ENSG00000112305 | SMAP1 | rs9455220 | 1.37E-08 | FALSE |
| ENSG00000136068 | FLNB | rs268774 | 1.53E-06 | FALSE |
| ENSG00000170891 | CYTL1 | rs9998195 | 2.00E-08 | FALSE |
| ENSG00000158457 | TSPAN33 | rs1835489 | 2.18E-06 | TRUE |
| ENSG00000087299 | L2HGDH | rs1465159 | 5.14E-09 | TRUE |
| ENSG00000155158 | TTC39B | rs677622 | 2.50E-12 | TRUE |
| ENSG00000152766 | ANKRD22 | rs11202868 | 2.90E-07 | TRUE |
| ENSG00000183624 | HMCES | rs4927956 | 1.31E-07 | FALSE |
| ENSG00000112305 | SMAP1 | rs9455220 | 1.37E-08 | FALSE |
| ENSG00000004864 | SLC25A13 | rs757332 | 6.09E-09 | FALSE |
| ENSG00000176788 | BASP1 | rs298549 | 3.62E-07 | TRUE |
| ENSG00000148459 | PDSS1 | rs17558723 | 1.66E-11 | TRUE |
| ENSG00000174125 | TLR1 | rs56054004 | 4.44E-07 | FALSE |
| ENSG00000135900 | MRPL44 | rs36053433 | 1.45E-06 | FALSE |
| ENSG00000204138 | PHACTR4 | rs114805497 | 1.42E-27 | TRUE |
| ENSG00000035928 | RFC1 | rs12642089 | 1.50E-06 | FALSE |
| ENSG00000152219 | ARL14EP | rs1933337 | 7.92E-07 | FALSE |
| ENSG00000152766 | ANKRD22 | rs11202868 | 2.90E-07 | TRUE |
| ENSG00000062716 | VMP1 | rs2665405 | 5.48E-11 | FALSE |
| ENSG00000205057 | CLLU1OS | rs61935212 | 1.36E-10 | TRUE |
| ENSG00000093144 | ECHDC1 | rs11556354 | 1.18E-10 | FALSE |
| ENSG00000158470 | B4GALT5 | rs6019957 | 7.08E-09 | TRUE |
| ENSG00000066027 | PPP2R5A | rs792433 | 1.63E-08 | FALSE |
| ENSG00000135900 | MRPL44 | rs36053433 | 1.45E-06 | FALSE |
| ENSG00000129007 | CALML4 | rs12438246 | 1.38E-08 | FALSE |
| ENSG00000156052 | GNAQ | rs1930541 | 3.69E-07 | FALSE |
| ENSG00000112308 | C6orf62 | rs2328831 | 4.58E-15 | FALSE |
| ENSG00000170234 | PWWP2A | rs2546372 | 7.47E-08 | FALSE |

Table S9. PARs in QTL-regions in L12 samples

(Ordered by decreasing peak signal, PARs belonging to the 14 related TE families included only)

| TE.chr | TE.start | TE.end | TE.family | Peak.pval | QTL.chr | QTL.start | QTL.end |
|---|---|---|---|---|---|---|---|
| chr2 | 224754093 | 224754450 | THE1B | 2.11E-88 | chr2 | 2.25E+08 | 224811722 |
| chr6 | 137539807 | 137540099 | Tigger3a | 6.00E-35 | chr6 | 1.38E+08 | 137546106 |
| chr4 | 5040086 | 5040704 | Tigger3b | 9.64E-28 | chr4 | 5017508 | 5042928 |
| chr2 | 68418273 | 68418838 | Tigger3c | 2.30E-27 | chr2 | 68396495 | 68468287 |
| chr2 | 224819059 | 224819594 | MLT1F | 1.43E-16 | chr2 | 2.25E+08 | 224833564 |
| chr2 | 224795308 | 224795759 | MLT1H | 1.81E-16 | chr2 | 2.25E+08 | 224811722 |
| chr2 | 224795308 | 224795759 | MLT1H | 1.81E-16 | chr2 | 2.25E+08 | 224833564 |
| chr2 | 224804164 | 224804525 | THE1B | 2.80E-11 | chr2 | 2.25E+08 | 224811722 |
| chr2 | 224804164 | 224804525 | THE1B | 2.80E-11 | chr2 | 2.25E+08 | 224833564 |
| chr6 | 24743677 | 24744015 | MLT1L | 9.56E-10 | chr6 | 24729115 | 24754859 |
| chr6 | 160270994 | 160271369 | THE1B | 3.19E-07 | chr6 | 1.6E+08 | 160285712 |

| Ensembl.ID | Gene.name | reQTL.rsID | reQTL.pval | Is.DEG |
|---|---|---|---|---|
| ENSG00000085449 | WDFY1 | rs36053433 | 3.08E-07 | FALSE |
| ENSG00000027697 | IFNGR1 | rs12529779 | 5.93E-07 | FALSE |
| ENSG00000170891 | CYTL1 | rs9998195 | 3.11E-15 | FALSE |
| ENSG00000221823 | PPP3R1 | rs75483778 | 2.30E-09 | FALSE |
| ENSG00000135900 | MRPL44 | rs13020280 | 9.87E-08 | FALSE |
| ENSG00000085449 | WDFY1 | rs36053433 | 3.08E-07 | FALSE |
| ENSG00000135900 | MRPL44 | rs13020280 | 9.87E-08 | FALSE |
| ENSG00000085449 | WDFY1 | rs36053433 | 3.08E-07 | FALSE |
| ENSG00000135900 | MRPL44 | rs13020280 | 9.87E-08 | FALSE |
| ENSG00000112308 | C6orf62 | rs2328831 | 5.18E-09 | FALSE |
| ENSG00000112110 | MRPL18 | rs9457747 | 6.03E-07 | FALSE |
